# The EN-TEx resource of multi-tissue personal epigenomes & variant-impact models

**DOI:** 10.1101/2021.04.26.441442

**Authors:** Joel Rozowsky, Jorg Drenkow, Yucheng T Yang, Gamze Gursoy, Timur Galeev, Beatrice Borsari, Charles B Epstein, Kun Xiong, Jinrui Xu, Jiahao Gao, Keyang Yu, Ana Berthel, Zhanlin Chen, Fabio Navarro, Jason Liu, Maxwell S Sun, James Wright, Justin Chang, Christopher JF Cameron, Noam Shoresh, Elizabeth Gaskell, Jessika Adrian, Sergey Aganezov, François Aguet, Gabriela Balderrama-Gutierrez, Samridhi Banskota, Guillermo Barreto Corona, Sora Chee, Surya B Chhetri, Gabriel Conte Cortez Martins, Cassidy Danyko, Carrie A Davis, Daniel Farid, Nina P Farrell, Idan Gabdank, Yoel Gofin, David U Gorkin, Mengting Gu, Vivian Hecht, Benjamin C Hitz, Robbyn Issner, Melanie Kirsche, Xiangmeng Kong, Bonita R Lam, Shantao Li, Bian Li, Tianxiao Li, Xiqi Li, Khine Zin Lin, Ruibang Luo, Mark Mackiewicz, Jill E Moore, Jonathan Mudge, Nicholas Nelson, Chad Nusbaum, Ioann Popov, Henry E Pratt, Yunjiang Qiu, Srividya Ramakrishnan, Joe Raymond, Leonidas Salichos, Alexandra Scavelli, Jacob M Schreiber, Fritz J Sedlazeck, Lei Hoon See, Rachel M Sherman, Xu Shi, Minyi Shi, Cricket Alicia Sloan, J Seth Strattan, Zhen Tan, Forrest Y Tanaka, Anna Vlasova, Jun Wang, Jonathan Werner, Brian Williams, Min Xu, Chengfei Yan, Lu Yu, Christopher Zaleski, Jing Zhang, Kristin Ardlie, J Michael Cherry, Eric M Mendenhall, William S Noble, Zhiping Weng, Morgan E Levine, Alexander Dobin, Barbara Wold, Ali Mortazavi, Bing Ren, Jesse Gillis, Richard M Myers, Michael P Snyder, Jyoti Choudhary, Aleksandar Milosavljevic, Michael C Schatz, Roderic Guigó, Bradley E Bernstein, Thomas R Gingeras, Mark Gerstein

**Author notes:** co-first authors (Joel Rozowsky, Jorg Drenkow, Yucheng T Yang, Gamze Gursoy, Timur Galeev, Beatrice Borsari, Charles B Epstein, Kun Xiong, Jinrui Xu, Jiahao Gao). co-senior authors (Michael C Schatz, Roderic Guigó, Bradley E Bernstein, Thomas R Gingeras, Mark Gerstein).

## Abstract

Understanding how genetic variants impact molecular phenotypes is a key goal of functional genomics, currently hindered by reliance on a single haploid reference genome. Here, we present the EN-TEx resource of personal epigenomes, for ∼25 tissues and >10 assays in four donors (>1500 open-access functional genomic and proteomic datasets, in total). Each dataset is mapped to a matched, diploid personal genome, which has long-read phasing and structural variants. The mappings enable us to identify >1 million loci with allele-specific behavior. These loci exhibit coordinated epigenetic activity along haplotypes and less conservation than matched, non-allele-specific loci, in a fashion broadly paralleling tissue-specificity. Surprisingly, they can be accurately modelled just based on local nucleotide-sequence context. Combining EN-TEx with existing genome annotations reveals strong associations between allele-specific and GWAS loci and enables models for transferring known eQTLs to difficult-to-profile tissues. Overall, EN-TEx provides rich data and generalizable models for more accurate personal functional genomics.

## INTRODUCTION

The Human Genome Project assembled one representative haploid sequence 20 years ago (Collins et al., 2003; Venter et al., 2001). Since then, many individual genomes have been sequenced (Stephens et al., 2015). Compared to the reference, an individual’s personal genome typically contains ∼4.5 million variants (The 1000 Genomes Project Consortium et al., 2015). The vast majority of these are located in non-coding regions and are most often present in the heterozygous state (French and Edwards, 2020; Hindorff et al., 2009; Zhang and Lupski, 2015). A goal of functional genomics is to assess the impact of these variants on molecular endophenotypes (e.g., epigenetic activity, RNA expression, or protein levels) and relate these to cell, tissue, and organismal traits, including disease phenotypes (Civelek and Lusis, 2014; Knight, 2014; Manning and Cooper, 2017).

To this end, researchers have conducted many genome-wide association studies (GWAS) and expression quantitative trait loci (eQTL) analyses associating genetic variants with phenotypic traits and changes in gene expression. In particular, the Genotype-Tissue Expression (GTEx) project has performed RNA sequencing (RNA-seq) experiments on >40 human tissues from nearly 1,000 individuals, allowing for the identification of >175K eQTLs (GTEx Consortium, 2013; 2015; 2017; 2020). In complementary fashion, the Encyclopedia of DNA Elements (ENCODE) project and the Roadmap Epigenomics Project were initiated to annotate functional genomic regions throughout the human genome (Encode Project Consortium et al., 2007; Encode Project Consortium et al., 2020; Kundaje et al., 2015).

However, these studies have been carried out using the generic reference genome, not directly using the variations observed in an individual’s diploid sequence. By using a diploid genome, heterozygous loci can distinguish sequences from each of the two parental chromosomes (haplotypes) and give rise to distinct molecular signals from each (e.g, RNA expression or transcription factor [TF] binding). The imbalance of expression or epigenetic activity between the haplotypes can be accurately measured by taking the reference allele as a baseline, avoiding biological and technical biases. If the imbalance is significant, the heterozygous variant is termed allele-specific (AS). AS variants have been identified in numerous previous studies and are implicated in several diseases (Baran et al., 2015; Castel et al., 2020; Chen et al., 2016; Chen et al., 2012; Do et al., 2020; Liu et al., 2018; Maurano et al., 2015; Onuchic et al., 2018; Pirinen et al., 2015; Robles-Espinoza et al., 2021). (Note that only some AS variants are thought to be causal for the observed changes.)

Here, to better connect personal genomes and functional genomics, we created the EN-TEx resource (ENCODE assays applied to GTEx samples). In particular, this resource comprises a uniformly processed dataset of >10 functional genomic assays, consistently collected for ∼25 tissues from four individuals. It is coupled with long-read genome assemblies, containing comprehensive sets of structural variants (SVs). Compared to what was previously possible, mapping reads from the assays directly to diploid genomes allows for more precise quantification of differential expression and regulatory-element activity and for directly visualizing the impact of SVs on chromatin; the uniform nature of the dataset makes possible more precise ascertainment of inter-individual vs. inter-tissue differences; and the scale of the resource enables the creation of the largest catalog of non-coding AS variants, an order-of-magnitude more than what was available previously. We leveraged this catalog to build generalized models of variant impact. In particular, we created a model that predicts the AS imbalance resulting from a single-nucleotide variant (SNV) just from the extended sequence context around a site (i.e. within a ∼250 bp window around an SNV with activity in a particular assay). It highlights the importance of ∼100 TF motifs we term AS-sensitive. Moreover, we can use these to disentangle the complicated relationship between the AS expression of a gene and the AS activity of its promoter. In addition, we can relate the EN-TEx resource to external genome annotations -- eQTLs and regulatory elements already known for the human genome. We built generalized models that transfer eQTLs from a source tissue to a different target one, leveraging the fact that EN-TEx represents a uniform collection of epigenetics data in hard-to-obtain tissues. This is practically quite useful given it is typically much easier to measure eQTLs directly in blood than other tissues, such as the heart, especially when using large cohorts of individuals. We also show that data from the EN-TEx resource can “decorate” existing regulatory elements, identifying subsets that are much more highly enriched with eQTL and GWAS SNVs than had been previously possible and illuminating broad relationships between conservation, AS activity and tissue-specificity. Overall, EN-TEx provides a powerful resource of multi-omic information and describes a more accurate approach for future functional genomics. Such information will be crucial for the interpretation of whole genomes and for future applications of them in precision medicine.

## RESULTS

### Improvements in Genome Analysis from Uniform Multi-tissue Data Collection &Diploid Genome Mapping

We sequenced and phased the genomes of four individuals from the GTEx cohort (identified as individuals 1 through 4) using various complementary sequencing technologies (i.e., short-read Illumina, linked-read 10x Genomics, long-read PacBio, and Nanopore, see STAR Methods). After identifying SNVs, small insertions and deletions (indels), and SVs, we phased the haplotypes of the assembled genomes using linked-reads and proximity ligation sequencing (Hi-C; Fig. S1.1) (Edge et al., 2017). This step generated long sequence blocks of phased variants extending across each chromosome, forming phased personal genomes for each of the four individuals (Fig. 1). The paternal/maternal origins of many of the phased blocks were determined by comparing the AS expression levels with known imprinted loci (Fig. 1B, S1.1B, S1.1C, and STAR Methods).

**Fig. 1.**
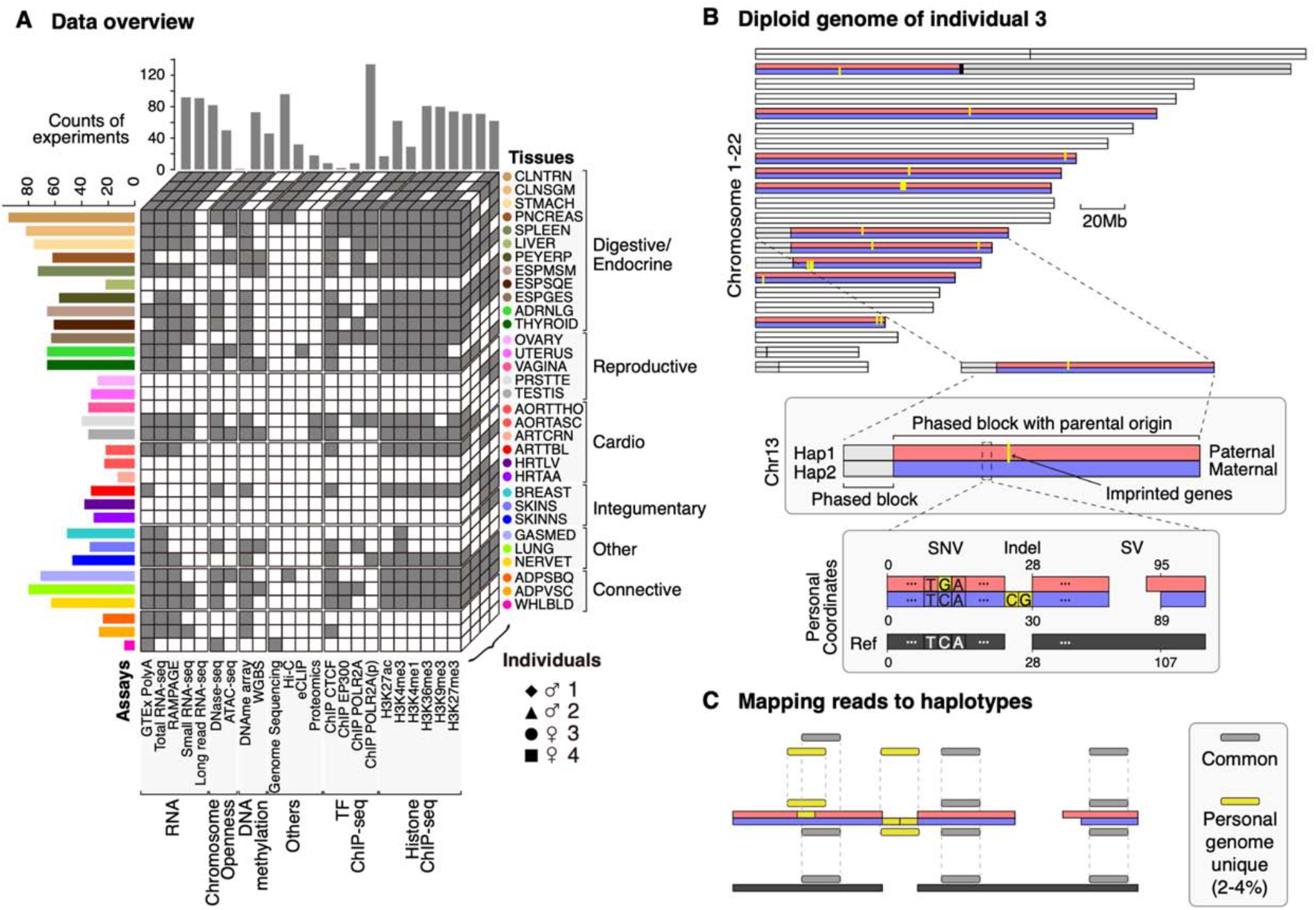
Personal Genome and Data Matrix. **(A) Data matrix for the EN-TEx resource**. Each voxel in the cube stack corresponds to a functional genomic assay for a particular tissue from one of the four individuals. A dark gray voxel indicates that the functional genomic data has been generated. The total experimental counts per assay across the tissues and individuals are shown above the cube stack. The total experimental counts per tissue type are shown on the left. The color and acronym scheme of the tissues is adopted from GTEx (GTEx Consortium, 2020). **(B) Illustration of the personal genome in contrast to the reference genome**. The personal diploid genome of individual 3 is shown at the top of the panel. Its construction explicitly considers SNVs, indels, and SVs. The chromosomes are phased with long-read sequencing to form phased blocks. With known imprinting events (yellow) in the human genome, the maternal (blue) or paternal (red) origin of many of the phased haplotype blocks can be identified (see STAR Methods). As an example, chromosome 13 (chr13) contains two phased blocks, one of which has its maternal and paternal haplotypes identified based on an imprinted gene’s AS expression patterns. A schematic diagram of a genomic region in chr13 shows the detailed differences between the personal diploid genome and the reference haploid genome. Because of the genetic variants in the personal genome, the diploid representation and the reference have different coordinate systems and sequences. **(C) Mapping reads to personal and reference genomes**. Due to the genetic variants, some reads can be mapped only to the personal genome but not to the reference genome (and to a lesser extent just to the reference). Such reads (yellow) are referred to as “personal genome unique,” whereas the reads that can be mapped to both genomes (gray) are referred to as “common.” Compared with the reference genome, using the personal genome results in an average of ∼2.5% more mapped reads for the various functional genomic experiments. The percentage ranges from ∼1% for RNA-seq to ∼4% for Hi-C data (more details in Fig. S1 and STAR Methods). Note that the numbers for these differences only apply to the high-stringency mapping (STAR Methods).

In parallel, we carried out 1,635 experiments from >10 different epigenome, transcriptome, and proteome assays on ∼25 tissues obtained from each of four individuals (i.e. ChIP-seq, ATAC-seq, Hi-C, DNase-seq, whole-genome bisulfite sequencing [WGBS], short and long-read RNA-seq, eCLIP, and labeled proteomic mass-spectrometry; Fig. 1A, S1.2A, and STAR Methods). All datasets in the EN-TEx resource were processed using the personal phased diploid and reference genomes, giving rise to three mappings and three corresponding signal tracks for each assay (maternal and paternal haplotypes and the reference; Fig. 1C and S1.2).

We found ∼2.5% more reads mapped to the personal genomes than the reference (for strict mapping criteria; Fig. 1C and STAR Methods). This mapping had a measurable effect (>2 fold) on the expression levels of >200 genes across the four individuals, a change comparable in magnitude to the differential expression often found in comparing healthy to disease states (Fig. 2A and S1.3) (Shang et al., 2020; Su et al., 2019; Vennou et al., 2020; Zhong et al., 2019). A similar fraction of regulatory regions exhibited a significant change in activity levels when using the personal compared to the reference genome (specifically, comparing the H3K27ac signal level on ENCODE candidate cis-regulatory regulatory elements [cCREs], Fig. 2A and S1.3) (Encode Project Consortium *et al*., 2020).

**Fig. 2.**
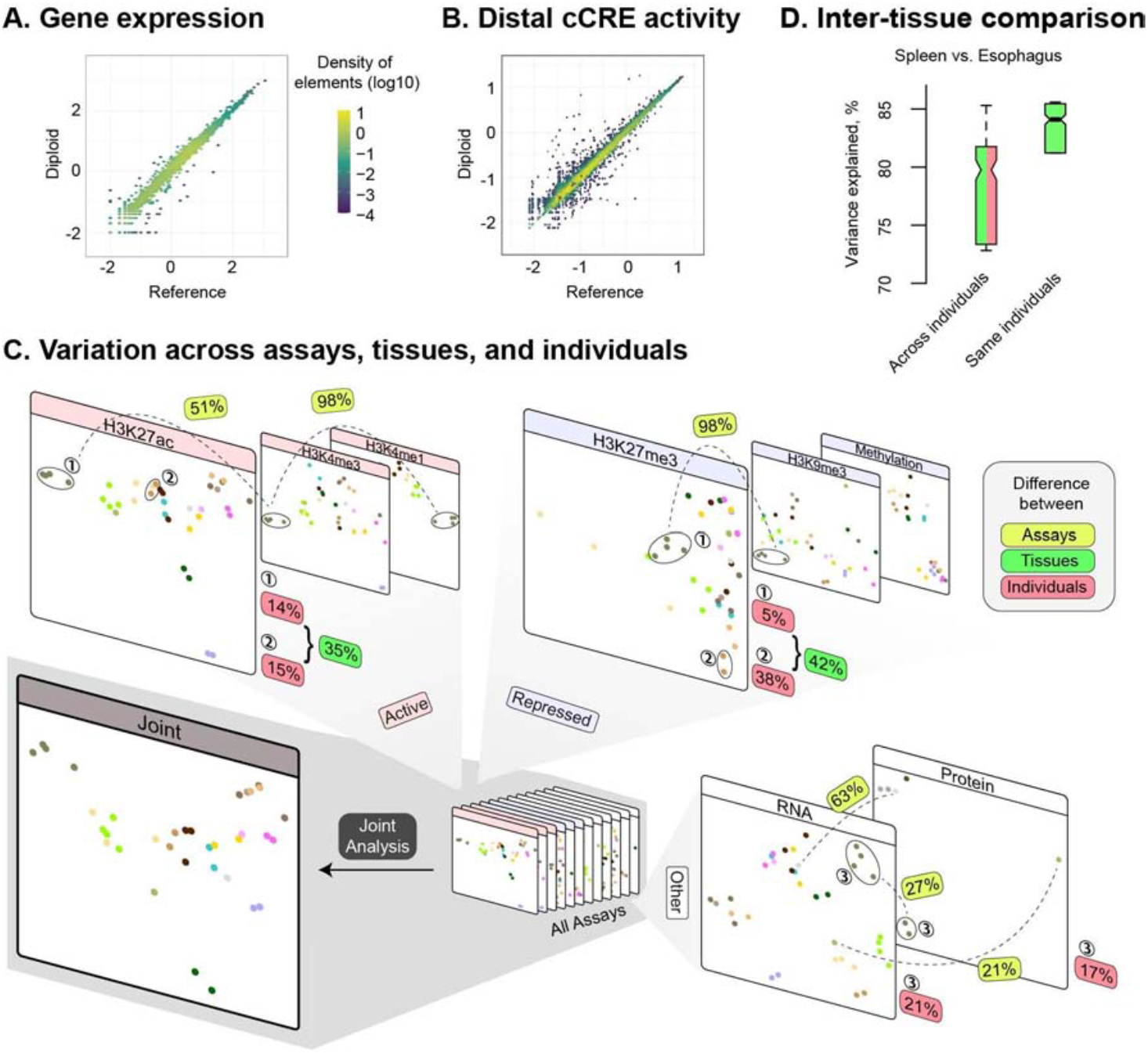
Improvements in Genome Analysis from Uniform Multi-tissue Data Collection and Diploid Genome Mapping. **(A) Comparing the quantification of gene expression using personal diploid genomes and the reference haploid genome**. This panel shows the gene expression levels for individual 3 calculated using the reads mapped to the reference genome (x axis) and to the personal diploid genomes (y axis). The expression values of the genes (TPM) are log-transformed, i.e., log10(TPM + 0.001). The color scale indicates the density of the genes. **(B) Comparing the ascertainment of regulatory element activity using personal diploid genomes and the reference haploid genome**. This panel focuses on ENCODE distal cCREs. For H3K27ac, we calculated its activity at cCREs using a similar formula as the expression levels of genes. An analogous scatter plot for proximal cCREs is shown in Fig. S1.3B. **(C) Consistently analyzing functional genomic data across individuals, tissues, and assays**. Joint analysis of variation (JIVE) (Hellton and Thoresen, 2016) was used to project all of the functional genomic signals from different samples onto two-dimensional planes (one for each assay, e.g., ChIP-seq, ATAC-seq, DNase-seq). For each assay type (e.g., H3K27ac), the samples are represented as dots on the plane. Two dots that are close to each other have similar levels of functional genomic signals. Combining all of the types of assays, we projected the samples onto a single two-dimensional plane. The tissues are colored according to the GTEx convention from Fig. 1. For each tissue, the number of colored dots indicates the number of individuals, up to four. In addition, a linear-regression-based approach was used to measure the difference between two samples in functional genomic signals (see details in STAR Methods). The variance explained is used to indicate the similarity between two samples. Note that the value of variance explained is equal to the squared correlation coefficient. The difference is measured by one minus the explained variance of the regression. Taking H3K27ac as an example, on average, only 14% of the variation in the functional signals between the spleens of two individuals is unexplained, indicating that the spleen samples have very similar H3K27ac signal patterns. The dissimilarity is also small (15%) between the transverse colons of two individuals. In contrast, the average dissimilarity between these two types of tissues of the same individuals increases to 35%. For H3K27me3, the dissimilarity between different tissues is also high (42%). The dissimilarities between different types of functional signals are even higher. In particular, the dissimilarity between H3K27ac and H3K4me3 is 51%. See STAR Methods and Fig. S1.4 for more details. **(D) An example of more accurate inter-tissue comparisons possible with matched samples**. For histone modification signals at cCREs, the spleen and esophagus of the same individual are more similar (i.e., median 0.92 correlation, 84% variance explained [or 16% dissimilarity]) than those of different individuals (0.89 or 79%). This difference indicates the genetic and environmental difference between individuals and its influence on inter-tissue comparisons.

Because of its uniform data collection and processing, EN-TEx provides an ideal platform to consistently measure inter-individual, inter-tissue, and inter-assay variability (Fig. 2B and S1.4). In particular, we can explore all the sources of variation to place each EN-TEx sample in a high-dimensional space. It is readily apparent, as expected, that inter-individual variation is less than inter-tissue variation, which is less than inter-assay variation. We can also see how many well-known inter-assay comparisons can be placed in a consistent framework. For instance, the variation of RNA vs. protein level can be described consistently in relation to RNA vs. H3K27ac. Finally, for the specific situation of comparing between tissues, the EN-TEx resource allows us to determine inter-tissue differences with greater accuracy than for equivalent data not matched across individuals (∼4%, see Fig. 2C).

### Large-scale Determination of AS SNVs & Construction of the AS Catalog

We investigated AS behavior on a large scale using EN-TEx. For most assays, we performed these calculations uniformly using a standardized pipeline that dealt with various technical issues such as the reference and ambiguous mapping biases (for RNA/ChIP/ATAC/DNase-seq assays; Fig. S2.1 and STAR Methods) (Chen *et al*., 2016; Degner et al., 2009; Encode Project Consortium, 2012; Encode Project Consortium *et al*., 2007; Gerstein et al., 2012; Khurana et al., 2013; Rozowsky et al., 2011). Overall, we ran the pipeline on 28 tissues, 13 assays, and four individuals (encompassing ∼1,000 samples). On average, we detected ∼800 AS events in each sample, representing about ∼4% of the total number of accessible SNVs. (An accessible SNV is a heterozygous SNV with sequencing depth sufficient to detect AS behavior.) We were also able to group the AS SNVs within a genomic element together to determine its overall AS status; on average, we found ∼200 AS elements in each sample.

We had to use a more specialized approach for some of the assays, in particular, WGBS, Hi-C, and mass-spec proteomics. For instance, for Hi-C, we constructed haplotype-resolved contact matrices and then identified haplotype-specific AS interactions (Fig. S2.1H). Overall, per sample, we found ∼0.5M Hi-C interactions out of a total of ∼6.5M (Fig. S2.1I). We also identified AS peptides exhibiting significant imbalance corresponding to 696 unique genes (STAR Methods).

After determining the AS SNVs in each sample, we combined them across all tissues, individuals, and assays (Fig. 3A and S2.3). There are different strategies possible for combining, and we found that pooling the reads across tissues dramatically increased our detection power (by ∼5X), making it possible to identify ∼27K AS SNVs for an assay in an individual (for RNA/ChIP/ATAC assays, Fig. 3A, S2.3 and STAR Methods). We then combined the AS SNVs across all the assays and found ∼365K AS SNVs per individual (now including WGBS and DNase). Finally, when we combined these data across all four individuals, we reached a total of about ∼1.3M AS SNVs, which constitutes our AS catalog (with 516K coming from RNA-seq, ChIP-Seq and ATAC-seq, see Fig. 3A and S2.3).

**Fig. 3.**
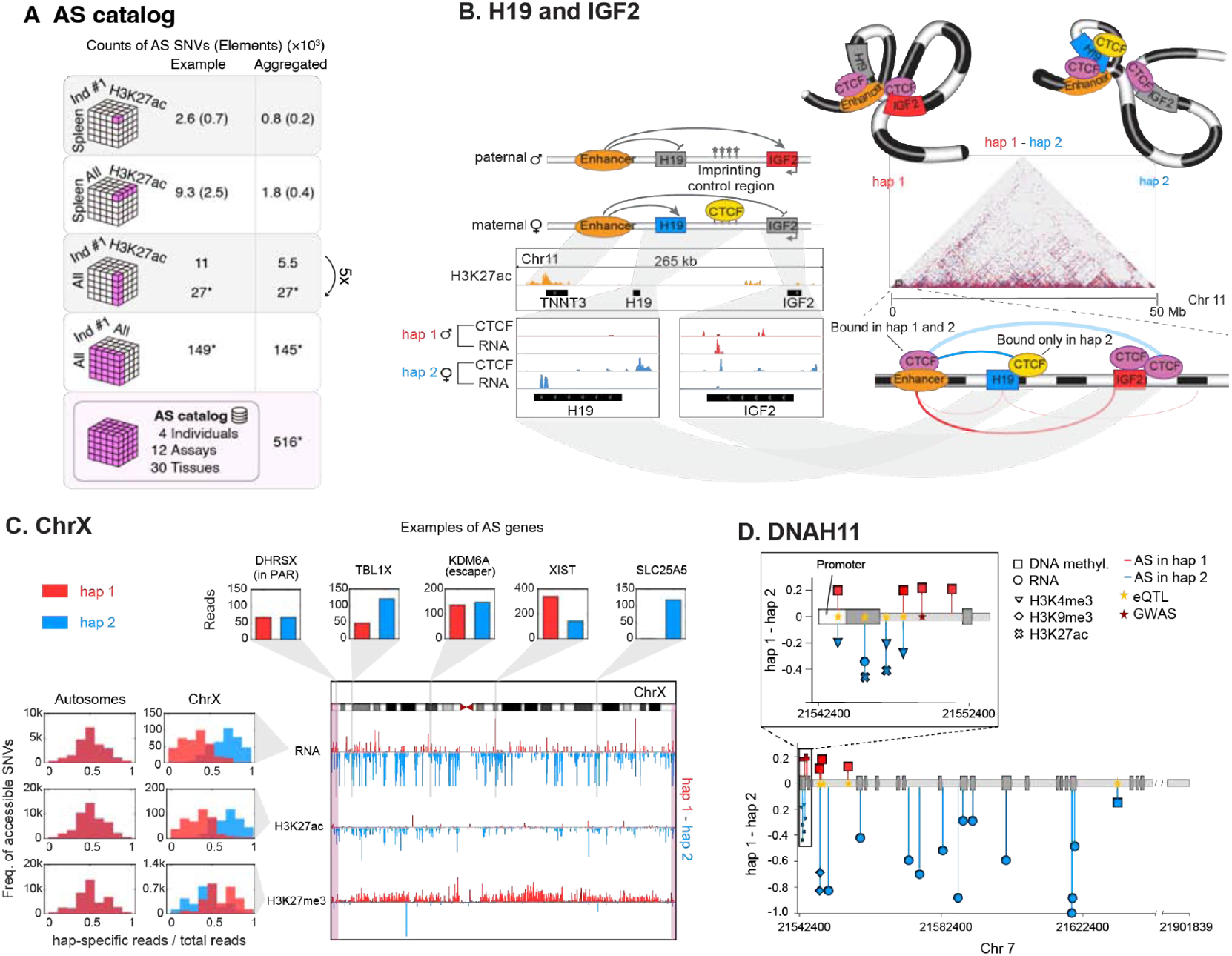
Building the AS Catalog & Examples of Coordinated AS Activity. **(A) Detecting AS events and using them to construct the catalog**. For a functional genomics experiment, if the numbers of reads mapped to the two haplotypes at a heterozygous SNV position are significantly different, the SNV is referred to as an AS SNV. We used two approaches to aggregate the information across samples to increase the detection power. In the “union” approach, any SNV observed as AS in any sample is included in our AS catalog. In the “pooling” approach, because a given functional genomic assay, e.g., RNA-seq, is usually conducted on many samples, for each SNV, we pool the mapped reads from all the samples together to determine its AS state (STAR Methods). This approach increases the statistical power of the AS test, and thus may be able to identify a SNV as AS, even though the SNV is not detected as AS in any sample. In the figure, the asterisk indicates when the aggregation involves the pooling approach. Genomic elements (e.g., cCREs) with a single AS SNV are identified as AS elements. For a given element with multiple heterozygous SNVs, we pool the reads of all the SNVs to determine its AS state. Here, we show more details about building the AS catalog: In the first row, using the H3K27ac assay of the spleen in individual 1 as an example, we identified 2.6K AS SNVs and 0.7K AS elements. Averaged over all individuals, tissues, and assays, we detected 0.8K AS SNVs and 0.2K AS elements. In the second row, using the union approach for the spleen H3K27ac data from the four individuals (union across individuals) resulted in 9.3K AS SNVs and 2.5K AS elements. The average numbers became 1.8K and 0.4K. In the third row, using the union approach for the H3K27ac data of all the available tissues of one individual (union across different tissues) resulted in 11K AS SNVs. Compared with the union approach, the pooling approach led to 27K AS SNVs. These two approaches’ average numbers were 5.5K and 27K, respectively. In the fourth row, we aggregated all AS SNVs for all the assays of all the different tissues from one individual (individual 1), resulting in 149K AS SNVs (using pooling across tissues and union across assays). The average of this number over all four individuals is 145K. Finally, we aggregated all the available assays and tissues from all the individuals and got 516K AS SNVs. Note that this result is only calculated using only RNA/ChIP/ATAC-seq assays. When we added in DNase and methylation, which require somewhat different approaches, the total increased to 1.3M. More details are shown in Fig. S2.3. **(B) Detecting AS events at a known imprinted locus: *H19*/*IGF2***. Functional genomic signals are measured in the gastrocnemius medialis of individual 2. In agreement with the known imprinting mechanism (top), we detected AS expression of *H19* on haplotype 2 (maternally expressed) and AS expression of *IGF2* on haplotype 1 (paternally expressed). Consistent with the expression data, we found an AS Hi-C interaction between an upstream cCRE and *H19* on haplotype 2 (the bold blue arc), and an AS interaction between the same cCRE and *IGF2* on haplotype 1 (the bold red arc). Other neighboring AS Hi-C interactions (which have fewer read counts or contact irrelevant genes) are also depicted with shaded arcs. **(C) Detecting coordinated AS activity across a chromosome**. Haplotype (Hap) 1 of chrX is inactivated in individual 3. The signal tracks show that, in the tibial nerve, hap1 chrX generally has lower expression levels, lower H3K27ac levels, and higher H3K27me3 levels than hap2 chrX. As a summary, such differentiations between two haplotypes are shown in histograms on the left. In contrast, the histogram of autosomes displays balanced gene expression and histone modification levels between the two haplotypes. The top inset bar-graphs show the expression of five genes as examples. The patterns between haplotypes are consistent with prior knowledge. *DHRSX*, located in the pseudoautosomal region (pink bars at the chrX ends of the signal track), and *KDM6A*, known to escape chrX-inactivation, show balanced expression between haplotypes. In contrast, *TBL1X* and *SLC25AC* fall in the inactivated region of chrX, showing lower hap1 expression. Similar differences in expression levels and histone modifications levels between chrX are also observed in the other female, i.e., individual 4 (Fig. S2.8D). **(D) Detecting novel AS events at an uncharacterized, disease-associated locus: the *DNAH11* gene associated with ciliary dyskinesia**. Because DNA methylation tends to repress gene expression, the polarity (direction of AS imbalance) of the AS DNA methylation in the promoter region is in the opposite direction to that of the AS expression and chromatin active state in the gene body. As expected, the active epigenetic marks H3K4me3 and H3K27ac demonstrate consistent AS imbalances, and most of the AS SNVs associated with *DNAH11* are known eQTLs from GTEx. One such SNV (rs11760336) lies within the *DNAH11* promoter, likely changing the gene expression directly. In addition, some of the AS SNVs overlap with known GWAS variants as indicated in the figure.

The AS catalog has a number of unique aspects. It is much larger than collections of AS chromatin events compiled by previous studies (see STAR Methods for a detailed comparison) (Chen *et al*., 2016; Leung et al., 2015; Onuchic *et al*., 2018). Moreover, we estimated that the AS SNVs detected in the four EN-TEx individuals cover 76% of common AS SNVs in the European population, suggesting that the catalog includes a majority of the AS events at common SNV loci in Europeans (STAR Methods). Because of its size, we can leverage the catalog to determine AS SNVs in an entirely new sample with increased sensitivity (Fig. S2.5). (In addition, using a related strategy, we can develop alternate, “high-power” AS assignments from joint calling across samples; Fig. S2.6.)

Another important aspect of the AS catalog is that most variants are in noncoding regions of the genome and are determined using non-RNA-based assays. In particular, only ∼2.5% of the AS variants in the catalog are uniquely detected by RNA-seq, and 95% are only detected by assays other than RNA-seq (Fig. S2.4). Finally, in addition to the common AS variants we detected, some AS sites correspond to rare SNVs; in total, we found that 101K of the 1.3M AS variants were rare (Fig. S7.2N). We were also able to cross-reference these rare SNVs with known pathogenic and deleterious variants, including 14 in ClinVar (STAR Methods) (Harrison et al., 2016).

Using the catalog, we found several examples of coordinated AS activity across different assays. First, we surveyed known imprinted loci, finding that AS activity is fairly consistent across tissues (Fig. S2.1F and S1.1C). A good example is the classic case of the *IGF2* and *H19* genes.

As expected, in several tissues, we observed that *H19* is expressed only maternally, and *IGF2*, only paternally, due to AS CTCF binding at the imprinting control region (Fig. 3B) (Autuoro et al., 2014). Going beyond this, haplotype-resolved Hi-C showed that, on the maternal haplotype, a cCRE upstream of *H19* interacts with this gene but not with *IGF2*. In contrast, on the paternal haplotype, the same cCRE only interacts with *IGF2*, suggesting a potential mechanism for this locus.

A second example is in the coordinated activity over chromosome X. On this chromosome, we observed gene expression, active histone marks, and POL2R and CTCF binding skewed toward one haplotype, with repressive marks biased to the other (Fig. 3C, S2.8A, and S2.8B). There are notable exceptions, including genes in pseudoautosomal regions (e.g., *DHRSX*) and documented “escaper” genes (e.g., *KDM6A* (Itoh et al., 2019)). In addition, haplotype-specific Hi-C manifested great differences in AS interactions on chromosome X at some loci (e.g., *XACT*; Fig. S2.8C and STAR Methods). Finally, a third and novel example of coordinated AS activity is *DNAH11, a* gene associated with ciliary dyskinesia (OMIM #611884). We observed AS methylation in the promoter regions on the opposite haplotype to the AS expression and activity of H3K4me3 and H3K27ac, consistent with transcriptional downregulation. In fact, some of the AS SNVs in the promoter have previously been identified as GTEx eQTLs (Fig. 3D).

### Interrelating SVs & Chromatin Modifications

We identified ∼18K SVs in each of the four individuals (>50 bp in length; Fig. 4A, S3.1, and STAR Methods). The majority of the SVs tended to be depleted in most functional regions (e.g., exons and cCREs) and to have typical allele-frequency spectra, consistent with previous findings (Fig. S3.1) (Audano et al., 2019; Sudmant et al., 2015). Furthermore, on a large scale, we found that the SVs were distributed over the diploid genome unevenly with notably different associations with chromatin on different haplotypes (Fig. 4B).

**Fig. 4.**
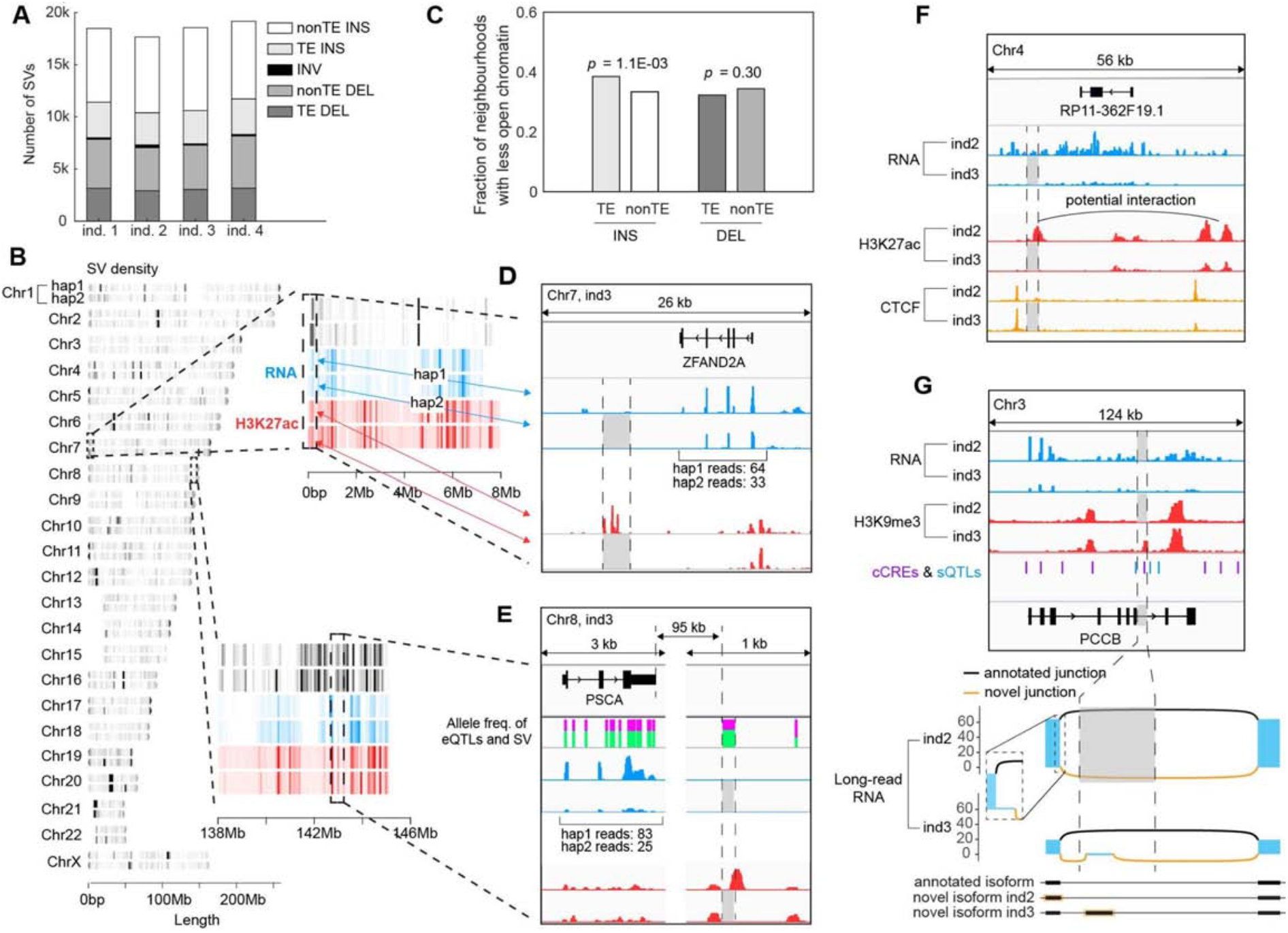
Relating SVs to Chromatin & Expression. **(A) Classification and frequency of SVs**. The fractions of insertions (INSs), deletions (DELs), and inversions (INVs) in each individual are shown. INSs and DELs are further classified based on whether they are related to transposable elements (TEs). See Fig. S3.1 for a detailed breakdown of SV composition. **(B) The chromosomal distribution of SVs on the diploid genome**. Genomic regions of chr7 and chr8 (in individual 3) are enlarged to show details of the positions of detected SVs and the levels of H3K27ac and RNA expression obtained from the transverse colon. The colors indicate the density of SVs. **(C) The effect of TEs on chromatin**. The genomic regions neighboring the TE insertions show reduced chromatin accessibility more often than those of the non-TE insertions. This difference is not observed between TE deletions and non-TE deletions The change in accessibility is determined by comparing the accessibility (from ATAC-seq) between the two haplotypes of each individual, taking the comparison of the haplotype without the SV as a reference (Fig. S3.3A and STAR Methods). P-values are based on the Chi-squared test. **(D) The effect of a 2.6 kb deletion**. The deletion in hap2 removed several H3K27ac peaks and reduced *ZFAND2A* expression from hap2 in the thyroid gland, consistent with previous work (more details shown in Fig. S3.2D) (Chiang *et al*., 2017). **(E) The effect of a 98-bp deletion**. The deletion in hap2 removed a H3K27ac peak downstream of the *PSCA gene*, potentially contributing to its reduced expression. This observation is in the transverse colon of individual 3. The heights of the green bars indicate the allele frequencies of the deletion and the surrounding GTEx eQTLs. The deletion and the eQTLs have similar frequencies and potentially are in linkage disequilibrium. Note, the heights of a green bar plus its corresponding magenta bar sum to 1. Similar results were observed in two other tissues from individual 3 (Fig. S3.2E). **(F) The effect of a 2.3-kb homozygous deletion**. This SV removed an H3K27ac peak downstream of lncRNA *RP11-362F19*.*1* in individual 3, which has lower expression in the spleen sample. In individual 2 without the SV, there are CTCF peaks that point to a potential interaction between the H3K27ac peaks downstream and upstream of the lncRNA. These observations together suggest that the loss of the H3K27ac peak due to the SV contributes to the reduction of lncRNA gene expression. See Fig. S3.2F for more details. **(G) The effect of a 5.2-kb homozygous deletion**. This SV removed an H3K9me3 peak in individual 2, potentially resulting in an increase in *PCCB* expression in the spleen. The sashimi plots, below the signal graphs, show examples of novel splicing isoforms identified by long-read RNA-seq in the adrenal gland of individual 2 and in the heart left ventricle of individual 3 (without the SV). Fig. S3.2G shows more detail on the splicing isoforms. The differences in splicing between individuals 2 and 3 potentially reflect the SV disrupting regions important to splicing, as suggested by the known GTEx sQTL sites nearby (Garrido-Martín et al., 2021). Alternatively, the differences in splicing between the two samples may be due to tissue specificity. Fig. S3.2H provides evidence of novel splicing variants that are potentially associated with an SV.

Overall, a significant fraction of genes with AS expression were associated with nearby SVs or indels (1.5% for SV deletions and 13.6% for small deletions, as shown in Fig. S3.2A; see Fig. S3.2B and S3.2C for additional analyses). Furthermore, many of these expression changes were also coupled to chromatin changes, as expected (Spielmann et al., 2018). In particular, we assessed whether chromatin significantly changes around heterozygous SVs by calculating a disruption score (Fig. S3.3A). We found that transposable element (TE) insertions were associated with a reduction in nearby open chromatin (compared with non-TE ones, Fig. 4C and S3.3B). We observed similar results when comparing the chromatin near SVs between EN-TEx individuals (both heterozygous and homozygous SVs; Fig. S3.3B). These results agree with findings that cells repress active chromatin in order to suppress TEs (Goodier and Kazazian, 2008; Levin and Moran, 2011; Zamudio and Bourc’his, 2010).

We found many specific examples of SVs impacting chromatin and nearby gene expression, potentially in a causal fashion. For instance, Fig. 4D shows a well-supported example: a heterozygous deletion, overlapping a known SV eQTL, removing an activating region, and a matching decrease in expression of a nearby gene (specifically, an H3K27ac peak near *ZFAND2A*)(Chiang et al., 2017). Fig. 4E shows a similar example: a heterozygous deletion removing an activating region near *PSCA*. Here, the deletion is not known to be associated with an eQTL but has a similar allele frequency to nearby eQTL SNVs and thus might represent the causal variant associated with them. On average, we identified ∼300 potential SV eQTLs in each individual (See STAR Methods S3.2; Fig. S3.2E shows additional documented examples of SV-eQTL associations, and Fig. S3.2I shows examples of whole-exon deletions).

Figures 4F and 4G show the analogous situation for homozygous deletions in comparing individuals. The first example shows an SV removing an active region and the corresponding downregulation of a nearby long non-coding RNA (lncRNA) (see Fig. S3.2J for a similar example). Fig. 4G shows an SV removing a likely repressive region in an intron of *PCCB* (a H3K9me3 peak). Moreover, this SV is adjacent to several GTEx splicing QTL sites (sQTLs; Fig. S3.2G), and long-read RNA-seq indicates that both individuals have different splice isoforms near the SV location. Notably, the EN-TEx resource enables direct comparison between SVs, determined by long-read DNA sequencing, with their impact on transcript structure, determined by long-read RNA-seq.

### Generalized Application #1: Predicting AS ChIP-seq Activity from Nucleotide Sequence

Up to this point, we have focused on the four EN-TEx individuals; now we turn to leveraging the resource to create generalized knowledge beyond them, broadly applicable in many contexts. In particular, we demonstrate four applications, focused on predictive modeling of AS behavior and approaches to “decorating” existing genome annotations.

The first application is modeling the impact of a variant to determine whether or not it will give rise to AS behavior. In particular, the ability of an SNV to disrupt a TF-binding motif suggests a direct relationship to the AS imbalance for a sequence-specific TF. Furthermore, given the importance of TFs in modulating open and closed chromatin, there is also a relationship, though less direct, to AS histone modifications. To study this, we cross-referenced all the genomic AS sites in the EN-TEx ChIP-seq data with 660 known human TF motifs and then ranked the motifs based on enrichment of AS SNVs (Fig. 5A)(Weirauch et al., 2014). Overall, we identified 194 TF motifs that were significantly enriched in AS SNVs and selected a “top 100” subset (STAR Methods).

**Fig. 5.**
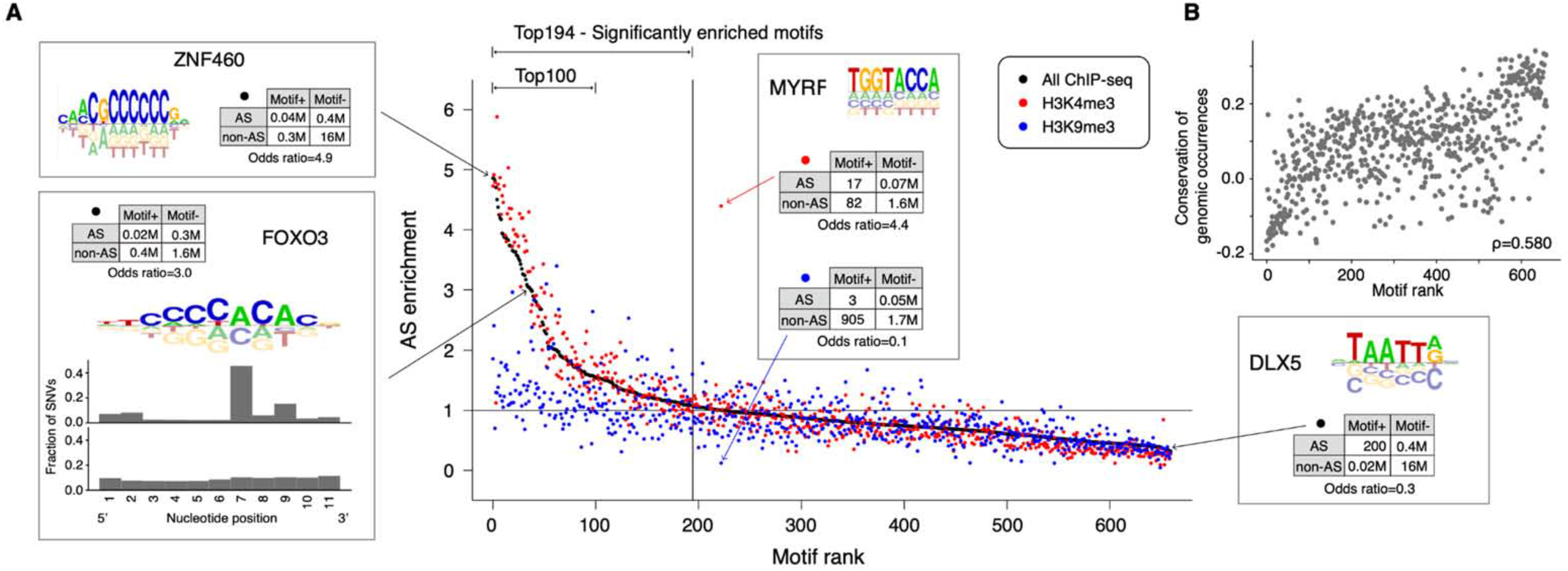
Relating TF Motifs to AS Activity. **(A) Motifs ranked by enrichment of AS SNVs**. We calculated the enrichment of AS SNVs for the motif of each TF. Taking FOXO3 as an example for the calculation, we made a 2-by-2 contingency table: motif or non-motif regions and AS SNVs or non-AS SNVs. The odds ratio of the table indicates the enrichment of AS SNVs for the FOXO3 motif. In the scatter plot, each dot corresponds to the motif of a TF, and the dots are ranked by their AS enrichment. Different colors indicate that the AS enrichments are calculated using different histone modification assays: using all accessible SNVs of all ChIP-seq (black), using accessible SNVs from H3K4me3 ChIP-seq (red), and using H3K9me3 ChIP-seq (blue). **(B) Motif ranking is correlated with conservation**. To calculate the conservation of a motif, we first calculated the phyloP score of each motif region in the genome, and then averaged these scores over the many genomic regions of this motif.

These top-ranked motifs represented TF binding sites particularly “sensitive” to mutations and more likely to give rise to AS behavior. They were enriched in C2H2 zinc-finger motifs, while the bottom-ranked motifs were more likely to have a homeobox domain (e.g. FOXO3 and ZNF460 vs DLX5 highlighted in Fig. 5A). FOXO3, in particular, represents well how AS SNVs affect the zinc-finger motif; the AS SNVs occurred at a distinct nucleotide position in the motif, known to modulate binding, while non-AS SNVs occurred more uniformly (STAR Methods)(Najafabadi et al., 2017). For many motifs, the enrichment associated with activating and repressing histone marks followed opposite trends (e.g. MYRF). Additionally, we found that the enrichment rank correlates with the conservation of the motif regions in the genome but is not correlated with a motif’s complexity (i.e., “PWM entropy”; Fig. 5B, S4.1, and STAR Methods).

The enrichment for AS SNVs in sensitive TF motifs suggests that we might be able to predict whether an SNV would be associated with AS behavior by whether or not it overlaps such a motif. To investigate this, we focused on CTCF: We started with constructing a simple model that predicted whether an SNV would be AS for CTCF binding based on whether it overlapped with a CTCF motif in regions showing regulatory activity (STAR Methods). This model had slight predictive performance (Fig. 6B). We then surmised that we could achieve better performance by including the sequence context surrounding the CTCF motif. To do this, we built progressively more complicated models, culminating in a deep-learning transformer model that allowed us to take into account the sequence in a 250-bp window around the SNV (Fig. 6A and STAR Methods) (Ji et al., 2021). The transformer model achieved surprisingly good performance (cross-validated AUROC of 0.69 using EN-TEx samples, for predicting whether or not an accessible SNV for CTCF binding in any tissue would be AS purely based on the sequence characteristics of the surrounding window; Fig. 6B). We were also able to build a similar model for POLR2A and H3K27ac; in the latter case, we validated the model, trained on EN-TEx, on an external benchmark dataset (AUROC of 0.74; Fig. 6B and S4.2B). Our transformer model predicts whether an SNV would be AS in a tissue-independent fashion. We next tried to enhance it in a tissue-specific fashion, by including additional epigenetic information; this only marginally improved the model, underscoring the overwhelming importance of sequence context in assessing the impact of a variant (Fig. 6C).

**Fig. 6.**
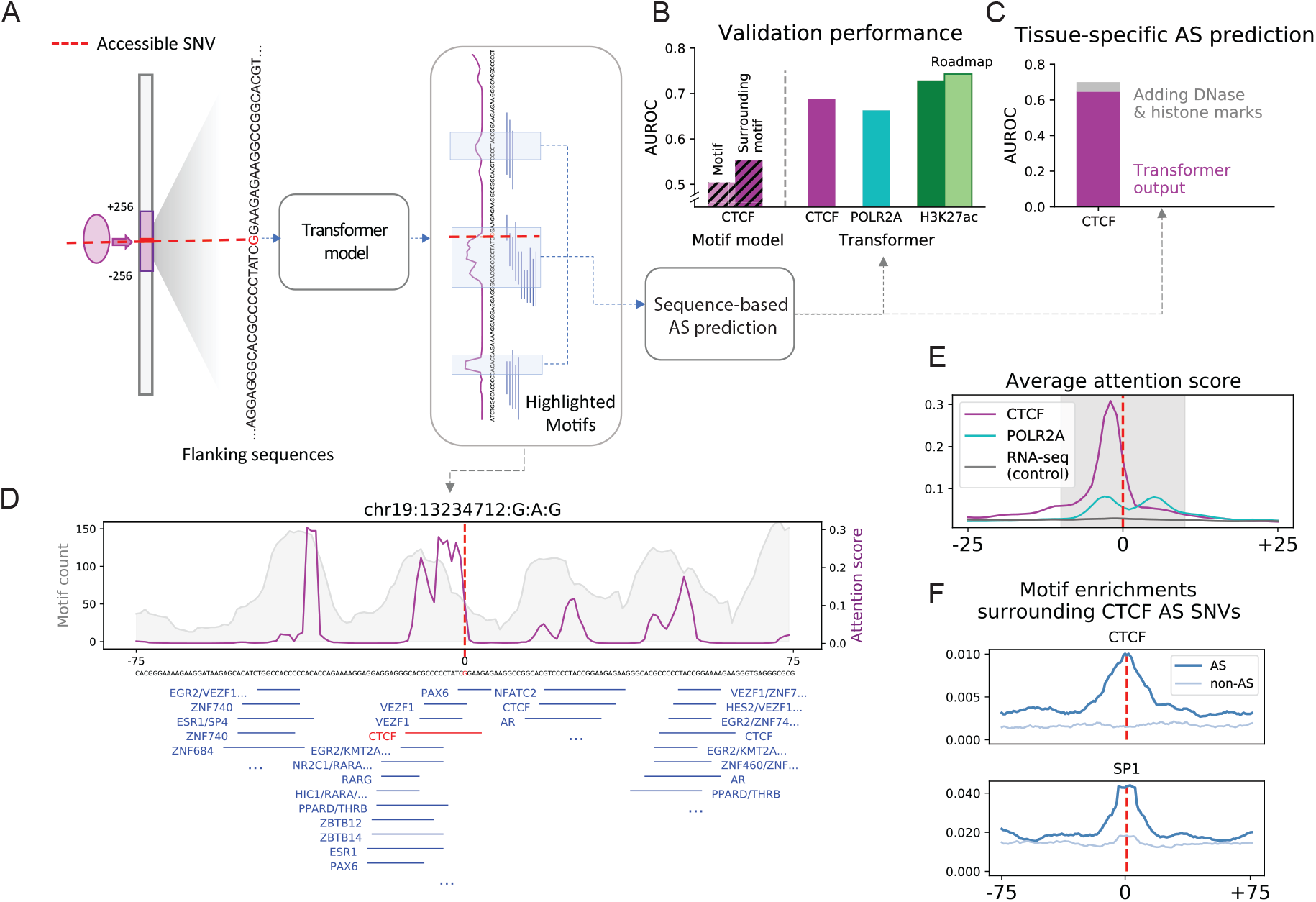
Deep-learning Model Predicting AS Activity from Sequence. **(A) Schema of the sequence-based predictive model**. A sequence-based transformer model was first trained on the flanking regions (128 bp) of accessible SNVs (for CTCF binding) to predict whether or not the SNVs are AS. The SNVs are from the AS catalog. The attention score (magenta lines) reflects the weights that the model attaches to different nucleotide positions in the input sequences. **(B) Average performance of the sequence-based model predicting AS effects**. The model was trained on AS SNVs from individual 3 and then tested on non-overlapping SNVs in other EN-TEx individuals. The model for AS H3K27ac prediction was also validated with an external dataset from the Roadmap Epigenomics Projects (Roadmap) (Kundaje *et al*., 2015). The model achieved comparable performance in both the internal and external validation. As a reference, we evaluated a simple logistic regression model with only overlapping information with CTCF motifs (motif overlap) as a covariate. In addition to this information, we included motif instances in the neighborhood (motifs in the window) to build another model as a reference. As a result, the transformer models outperformed the motif models. See STAR Methods for more details. **(C) Performance of a tissue-specific model using sequence features and epigenomic signals**. A random forest model was trained with the prediction score of the sequence-based transformer model and the tissue-specific epigenomic information (DNase and histone marks but not CTCF) to predict the tissue-specific AS effects. The performance was averaged across all tissues of the same individual. Adding epigenomic features only marginally improved the tissue-specific AS prediction performance. **(D) Example of attention patterns learned by the model**. Attention patterns on the flanking regions of a selected CTCF AS SNV (magenta) show strong consistency with high enrichment of motifs (gray). Interestingly, the regions with high attention scores (attention peaks) tend to contain several TF binding motifs that might interact with CTCF. The central peak surrounding the hetSNV position contains a central CTCF motif, which is highlighted in red. (Other examples of this attention score are shown in Fig. S4.2). **(E) Average attention pattern of the sequence-based model**. The attention score is averaged over 2,000 randomly selected test samples. As expected, the attention score peaks around the AS SNV for CTCF binding, whereas there are no such strong peaks for POLR2A. The probable reason is that POLR2A works in coordination with many different regulatory factors instead of particular TFs, and these factors are not necessarily close to POLR2A binding, leading to a less peaked attention score pattern. RNA-seq is used as a control because the AS SNV for expression is usually present as a tag and is not the causal SNV. **(F) Motif enrichment surrounding the AS CTCF SNV**. In line with the average attention patterns, some TF motifs, including the CTCF motif itself and SP1, show stronger enrichment within the proximity of the AS SNV of CTCF binding as compared to SNVs that are accessible to CTCF binding but do not show AS behavior. This result also agrees with the attention pattern of the model, indicating potentially discriminating motifs.

In order to better understand the sequence context that gives rise to AS behavior for our CTCF model we explored one characteristic of transformer models; they direct attention to specific sequence positions, often corresponding to known motifs. An example is shown in Fig. 6D; one can see the attention paid to the CTCF motif at the center, but many other locations with known motif clusters are also flagged as important. The attention score from the model averaged over many positions clearly shows that it is more focused on the central SNV than other “control” models (Fig 6E). This averaged attention score is ideal for comparing to overall motif occurrence: as expected, we observed a central enrichment for CTCF, but we also saw an enrichment for other TFs, such as SP1 (Fig. 6F and S4.2D).

### Generalized Application #2: Models Interrelating AS Activity at Promoters and Genes

We next investigated how the AS behavior of neighboring genomic elements are interrelated and potentially predictive of each other. We focused on the AS activity at the promoter using histone modifications and AS expression of the downstream gene (Fig. 7A). Simple statistics revealed that the relationship is more complicated than one would naively expect (Fig. 7B). Even though promoters of genes with AS expression are significantly more likely to have AS histone modifications, we found many examples of AS gene expression with no AS behavior at the promoter as well as many examples of the converse (see Fig. 7B and S5.1E). The complexity is potentially due to a number of factors including alternative distal regulation or redundancy of regulatory sites.

**Fig. 7.**
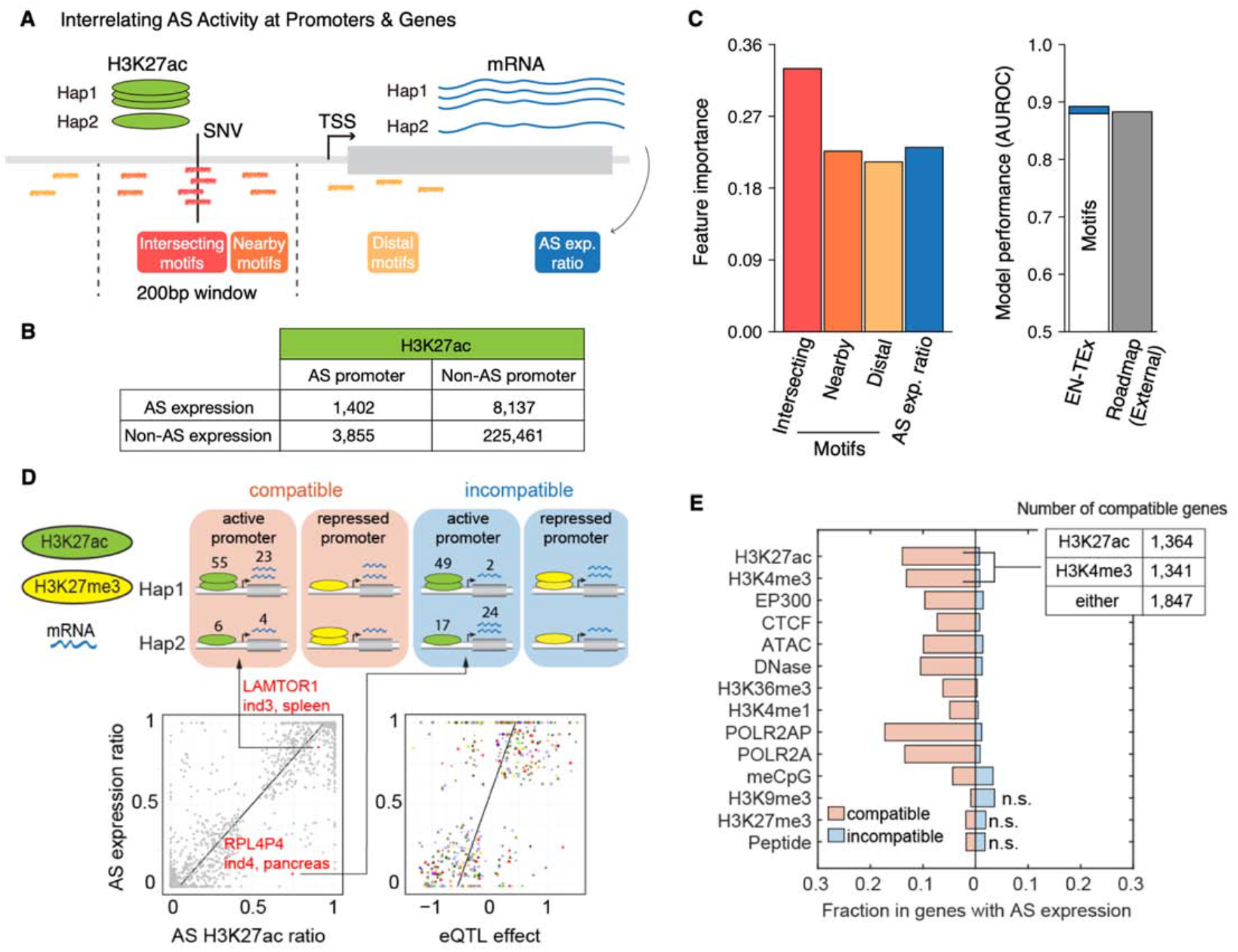
Interrelating AS Activity at Promoters & Genes. **(A) A schematic description of the statistical model predicting AS bound promoters**. The model attempts to predict whether the promoter will be AS (Details of the model are in the STAR Methods.). Motifs (colored short lines) of ranked TFs were used as the main features of the model. The first feature is the number of the top 100 AS sensitive TFs’ motifs that intersect the SNV. The second feature is those within 100 bp of the SNV but do not intersect. The third feature is the number of any of 660 TF motifs distal to the SNV (i.e. >100 bp away). AS bound promoters have significantly more motifs than non-AS bound promoters (also see Fig. S5.1D). The fourth feature AS imbalance ratio between transcripts from Hap1 and Hap2. **(B) Contingency table for AS genes and AS promoters**. We made a 2-by-2 contingency table: AS genes vs non-AS genes (in terms of AS gene expression) and AS promoters vs non-AS promoters (in terms of H3K27ac activity). The overall odds ratio of the table indicates the association between AS activity in genes and promoters. Tables for other assays are shown in Fig. S5.1E. **(C) Model performance and feature importance**. Gini impurity-based importance scores are shown for the four features in the model. We then used the different combinations of the four features to train and test our models. The left-hand-side of the panel shows that the sequence-based features (i.e., the three motif features) are the main contributors to good performance. Adding other features on top of these provides little improvement, which is indicated by the small blue line on the right-hand panel which indicates the incremental improvement of the AS expression ratio. The overall performance of the model is shown by the bars on the right-hand side (cross-validated performance on the EN-TEx individuals and then independent validation on the external Roadmap individuals -- note that the Roadmap result includes all 4 model features.) Performance for assays other than H3K27ac is shown in Fig. S5.1A. **(D) Compatibility between AS expression and AS promoter state**. As shown in the examples, a high active histone modification signal (e.g., H3K27ac) of a haplotype should correspond to high gene expression of the same haplotype. Note that for a repressive histone modification, this trend is the opposite. For AS genes with promoters accessible to H3K27ac, there is indeed a strong and positive correlation. The exact read counts of H3K27ac and gene expression, respectively, from haplotype 1 and 2 for *LAMTOR1* and *RPL4P4* are indicated in the examples. In addition, we used the slope (beta coefficient) of the leading eQTL of an AS gene (GTEx Consortium, 2020). The slope is positively correlated with the AS ratio (overall Pearson’s correlation coefficient of 0.6 with p = 0.01). Note that genetic variants with slopes around 0 are unlikely to have the statistical significance to be identified as eQTLs. This also holds for SNVs with read fractions of approximately 0.5. Therefore, there are no data points observed at the center of the plot. **(E) Overall compatibility between AS expression & promoter AS chromatin activity**. The compatibility is measured by the fraction of genes for which AS expression that are compatible with promoter AS chromatin state or AS peptide expression (Fig. S5.2 and STAR Methods). As a null model, we randomly paired genes with promoters (and peptides) to calculate z-scores (more details in the STAR Methods). Compatibility between AS expression and AS methylation (meCpG), H3K9me3, and H3K27me3 is weak, potentially because these marks of repressed chromatin can also be associated with genes poised for transcription or genes that are actively transcribed (Berger, 2007; Suzuki and Bird, 2008).

Given this, we constructed statistical models to predict the AS behavior of promoters and genes (Fig. 7A, 7C, S5.1B, and S5.1F). Successful models for the promoter, remarkably, needed very few features, and most of these related to the sequence context, viz: (features 1 & 2) the number of AS-sensitive TF motifs near the central SNV in the promoter (overlapping or nearby), (3) any TF motif, sensitive or not, distal to the SNV, and (4) the AS expression imbalance of the downstream gene (Fig. S5.1B). Interestingly, other features that one might have expected to be important -- including the overall expression level of the gene or the eQTL status of the SNV in the promoter -- were not very informative (Fig. S5.1D). The model achieved strong performance on the EN-TEx individuals and could be validated on an external dataset (cross-validated AUROCs of 0.81 and 0.88 within EN-TEx and on external data, respectively; Fig. 7C, S5.1A and STAR Methods). Finally, given that RNA-seq data is much more readily available than ChIP-seq data, the model can be readily applied in a practical context, e.g., to predict AS promoter activity throughout the 838-individual GTEx cohort, using just RNA-seq data and genotypes (STAR Methods).

Next, we considered the particular case with known AS behavior in both the promoter and the associated gene. Here, we found quantitative agreement for the magnitude and direction of the AS imbalance for many different epigenetic and proteomic assays (Fig. 7E). As expected, we observed positive correlations between AS expression and AS imbalance for many different chromatin signals (e.g., H3K27ac and ATAC-seq) and negative correlations for repressive marks and DNA methylation (Fig. 7E and S5.2A). Fig. 7E summarizes the overall degree of compatibility (with an associated list of strongly compatible gene-promoter pairs, see STAR Methods). Finally, the association between AS behavior at the promoter and the downstream gene can be extended to include eQTLs: a positive correlation is evident between the GTEx eQTL effect size and the AS imbalance of the associated gene (both in terms of expression and promoter activity; Fig. 7D and S5.2A).

### Generalized Application #3: Using the EN-TEx Resource to Extend eQTL Annotations to Hard-to-obtain Tissues

The above correlation between AS activity and eQTLs is an example of how the EN-TEx resource can be integrated with external annotation (e.g., from GTEx). This integration can go further by applying the EN-TEx chromatin and AS information to help prioritize GTEx eQTLs for prioritizing putative causal variants, in line with previous findings (Fig. S5.2A and S6.1A) (Brown et al., 2017). More importantly, because EN-TEx includes ChIP-seq data, which is relatively more difficult to obtain than RNA-seq data and is uniformly generated from hard-to-obtain tissues (e.g., heart), we can use EN-TEx to extend existing eQTL annotations to additional tissues.

We start with the observation that eQTL SNVs have stronger chromatin signals in the tissues in which they are active than in the tissues in which they are not (Fig. S6.1B), suggesting that the chromatin around an SNV may influence its chance of being an eQTL in a tissue. Then, by combining the EN-TEx chromatin data and the GTEx eQTL catalog, we developed a random-forest statistical model that transfers the activity of an eQTL from a given donor tissue (the source, e.g., skin) to another target tissue (e.g., tibial artery) by considering the chromatin profile in the target tissue (from EN-TEx, Fig. 8A and S6.2A). Overall, when compared with known GTEx eQTLs, our predictions are highly accurate, independent of which donor or target tissues are employed (balanced accuracy of 0.86; Fig. 8B, S6.3B and S6.3C). Our model tends to transfer stronger GTEx eQTLs to the target (Fig. 8C); conversely, it also identifies “likely” eQTLs, not quite reaching the “official” GTEx significance threshold (likely due to sample size) but still achieving greater significance than those not transferred.

**Fig. 8.**
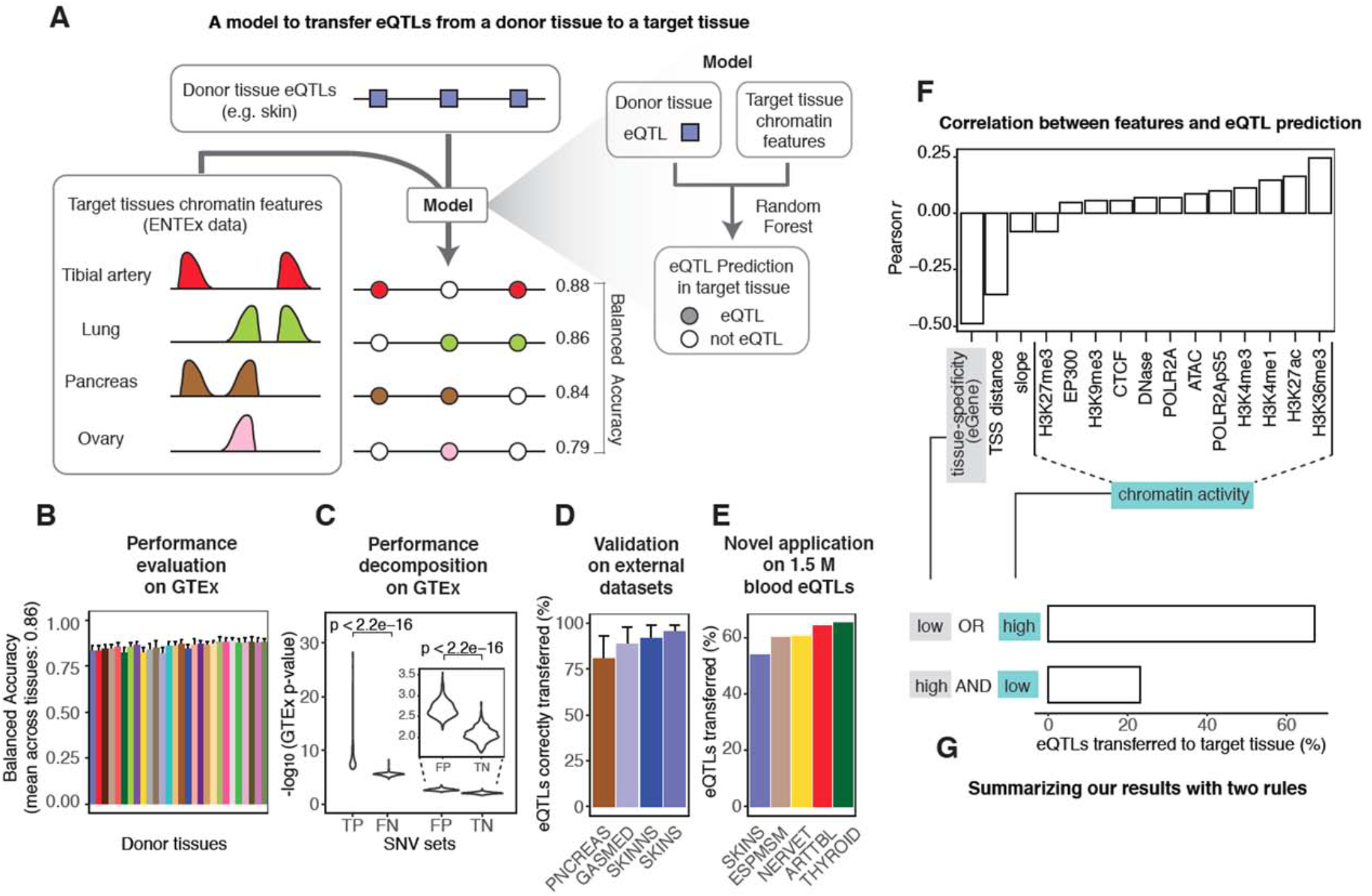
Transferring eQTLs across Tissues. **(A) Schema of the predictive model**. We employed a random forest algorithm to predict eQTLs using EN-TEx chromatin features (histone modifications, TF ChIP-seq, DNase-seq, and ATAC-seq). For a catalog of eQTLs active in a source tissue (donor), we aim to transfer them to another tissue (target) by leveraging the EN-TEx chromatin features in the target tissue (see also Fig. S6.2A). In addition, we consider some properties of the eQTLs in the donor tissue: tissue-specificity of the associated eGene, distance from the TSS of the eGene, and slope of the association with the eGene (not shown in the figure; Fig. S6.2C). Taking several target tissues (tibial artery, lung, pancreas, and ovary), we can transfer skin eQTLs with high accuracy (the balanced accuracy obtained for some examples of target tissues is shown). **(B) Performance evaluation of the model**. The predictive models have high performance (balanced accuracy), and the performance is robust to the donor tissues used. The X-axis indicates the tissues used as the donors, colored according to the GTEx convention, and the Y-axis indicates the average performance (balanced accuracy) across the target tissues that are available. The whiskers indicate standard deviations. The average balanced accuracy across all donor-target tissue pairs is 0.86. See also Fig. S6.3B and S6.3C. **(C)Performance decomposition of the model**. Our results can be classified into four groups: (1) the true-positive (TP) set of SNVs, in which the SNVs are predicted as eQTLs in the target tissue by both our model (model eQTL) and GTEx (GTEx eQTL); (2) the false-negative (FN) set, in which the SNVs are not eQTLs according to our model (model not eQTL), but are GTEx eQTLs; (3) the false-positive (FP) set, which contains the SNVs that are model eQTLs but are not eQTLs according to GTEx (GTEx not eQTL); (4) and the true-negative (TN) set, which contains SNVs that are model not_eQTL and GTEx not eQTL. We compared the p-values of the SNVs calculated by GTEx in the relevant target tissue. As a result, TP SNVs have more significant p-values than FN SNVs. We observed the same result when comparing FP and TN SNVs. This means that eQTLs predicted by our model in a given target tissue, but not part of the GTEx catalog in that tissue (FP), are more significant than eQTLs (from the donor) neither predicted by our models nor part of the GTEx catalog in that target tissue (TN). **(D) External validation of the model**. We have validated our predictions against four eQTLs catalogs other than GTEx, i.e., pancreas (PNCREAS), skeletal muscle (GASMED), suprapubic (SKINNS), and lower-leg (SKINS) skin (see also STAR Methods S6.3) (Kerimov *et al*., 2021). Our model has high performance for these four tissues, i.e., the majority of the eQTLs contained in these external catalogs for the four tissues are correctly predicted (mean sensitivity across all donor-tissue submodels for a given target tissue is reported; the whiskers indicate standard deviations). **(E) Large-scale application of the model**. We next applied the model to a novel set of roughly 1.5 M eQTLs, obtained from a large cohort (> 30,000 donors) study of blood (Võsa *et al*., 2021). We were able to predict a large proportion of these eQTLs in each of the EN-TEx target tissues. The bar plot shows the top five tissues with the largest fractions of predicted eQTLs. Aorta artery was used as the source donor tissue. See also Fig. S6.3D and S6.3E. **(F) Relative importance of the features in the model**. We computed the correlation between 15 selected predictive features (e.g., signal vector across SNVs) and the model’s probability of classifying donor-tissue eQTLs as eQTLs also in the target tissue. For instance, donor-tissue eQTLs marked by H3K36me3 in the target tissue are more likely to be classified as “eQTLs” (Pearson’s *r* = 0.25) in the target, while donor-tissue eQTLs associated with tissue-specific genes are less likely to be classified as “eQTLs” (Pearson’s r = -0.49). The bar plot shows, for each feature, the strongest Pearson’s correlation coefficient observed across all the 756 donor-target tissue pairs. See also Fig. S6.4A for a full description of these correlation patterns involving all the predictive features employed by the model. **(G) Schematic showing how two simple rules help to predict eQTLs in a target tissue**. To summarize the results in (F), we have found that two observations help partially define this model: i) if the eGene is not tissue-specific or the donor-tissue eQTL has high chromatin activity in the target tissue, then the donor-tissue eQTL is more likely to be an eQTL also in the target tissue; and ii) if the eGene shows some degree of tissue-specificity and the donor-tissue eQTL has low chromatin activity in the target tissue, then the eQTL is less likely to be transferable to the target tissue (see also Fig. S6.4B). Here, “chromatin activity” is defined as the sum of peaks of all chromatin features (histone modification ChIP-seq, TF ChIP-seq, DNase-seq, and ATAC-seq peaks at the SNV, described in Fig. S6.2C). As an example in the panel, we show the results obtained when transferring eQTLs from testis (donor tissue) to the thyroid (target tissue). See STAR Methods S6.4 and Fig. S6.4B for more details on the calculation of these results, as well as to how they compare to those obtained with other donor-target tissue pairs.

We further validated our model, trained on GTEx, against other “external” eQTL catalogs (Kerimov et al., 2021). In particular, it correctly identified >75% of the eQTLs reported by external catalogs for pancreas, skeletal muscle, and skin (Fig. 8D). Finally, to showcase the value of our approach to enhance existing eQTL catalogs, we applied it to a set of 1.5M blood eQTLs from a recent large cohort study; we were able to transfer up to 60% of these, enhancing the GTEx catalog with ∼500K new potential eQTLs per tissue (Fig. 8E, S6.3D and S6.3E) (Võsa et al., 2021). Note the utility of this application: up to now, large-cohort, high-power eQTLs studies so far have been conducted mostly on a few readily available tissues, such as blood or skin (Võsa *et al*., 2021). Thus, the uniformly collected EN-TEx chromatin data allow us to leverage these existing annotations to other, more difficult-to-secure tissues.

Finally, we evaluated the contribution of the different genomic features to the model (Fig. 8F and S6.4A). As expected, we found that SNVs with chromatin activity, especially H3K36me3, were more likely to be transferred. We observed the opposite for SNVs associated with genes that are tissue-specific or have distant transcription start sites (TSS) (Fig. 8F and S6.4A). Given this, we can summarize the main features of our model in a simple two-rule heuristic (Fig. 8G and S6.4B).

### Generalized Application #4: Decorating ENCODE Regulatory Elements with EN-TEx Tissue & AS Information

The ENCODE encyclopedia annotations were constructed using a disparate collection of cell lines and tissues; they are also devoid of variant annotations. Similar to the application extending GTEx eQTLs above, we can layer the results from EN-TEx onto these annotations, “decorating” them and extending their utility. In particular, we can combine them with the AS catalog, highlighting subsets exhibiting AS activity (Fig. S7.1B). Next, for each EN-TEx tissue we consistently determined whether each ENCODE element is active, repressed, or bivalent (Fig. 9A and S7.1F). (Note that repressive and bivalent categories are not included in the current ENCODE encyclopedia.) Overall, 97% of the ∼1M cCREs in the ENCODE encyclopedia can be decorated, and we validated our decorations using data from other studies with tissue-matched Hi-C (Fig. S7.1G).

**Fig. 9.**
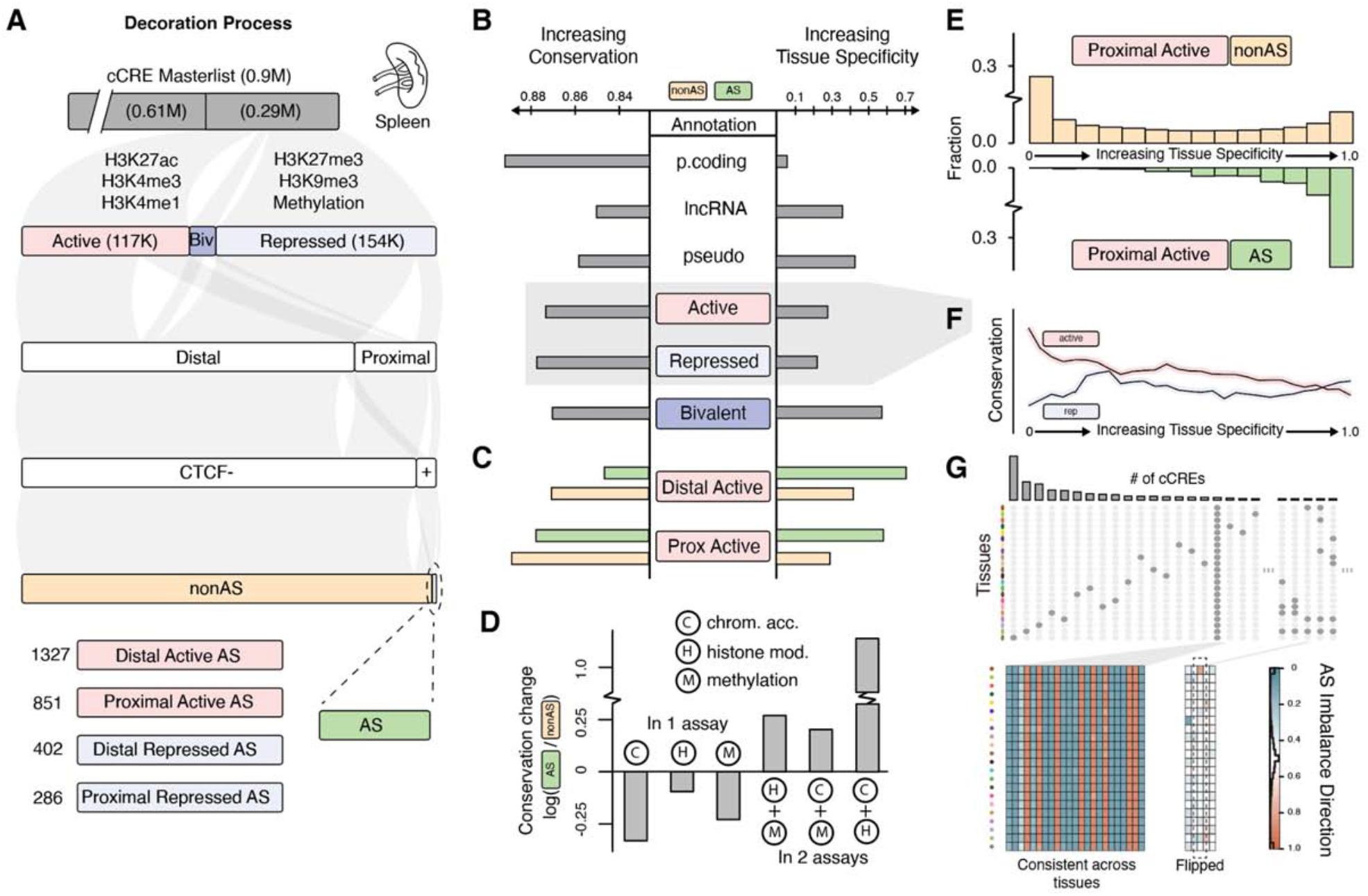
Decorating the ENCODE Registry and Relating Tissue Specificity & Conservation. **(A) Workflow for decorating cCREs by personal functional genomic data**. The workflow starts with the master list of cCREs from the ENCODE registry. These 0.9 million cCREs currently have no tissue-specific information. Spleen is used as an example to illustrate the tissue-specific decoration of these cCREs using EN-TEx data. In this tissue, 290k of the cCREs have functional genomic signals and can be categorized as active (117k), repressed (154k), or bivalent (19k). Then, for each category, the cCREs can be consecutively classified according to their genomic locations (proximal or distal), CTCF binding, and allele-specificity. (This classification can occur in any order.) As a result, the spleen has 2,866 AS cCREs with various genomic activities and locations. More details on the workflow are described in the STAR Methods. The numbers of active and repressed cCREs are comparable in each tissue (i.e., on average ∼202k active and ∼166k repressed cCREs). Only a small subset of the active cCREs exhibit AS activity (∼2.5% or 1,750 cCREs, averaged across tissues; see Fig. S7.1D for all available tissues). **(B) Tissue specificity and conservation of various genomic annotations**. Taking the distal active cCREs as an example, this annotation category contains all distal active cCREs observed in all available tissues. The tissue specificity of this annotation category is simply the fraction of the cCREs observed as distal and active in only one of the available tissues (see Fig. S7.2 and STAR Methods for more details). This fraction is referred to as fractional uniqueness, and is determined for each of the annotation categories. A smaller uniqueness indicates that the members of the category are more ubiquitous across tissues. The plot shows uniqueness values for many annotation categories. Additionally, conservation (degree of purifying selection) for each annotation category is estimated by the fraction of rare variants from gnomAD in the annotated genomic regions of the annotation category. **(C) Tissue specificity and conservation of annotations separated by allelic activity**. For the genomic regions of each annotation category, we separated them into AS or non-AS according to their AS states. Using the same method as in (B), we calculated the tissue specificity and conservation, respectively, for the two groups. As shown in the panel, the AS group and non-AS group of the same annotation category tend to have distinct patterns in terms of conservation and tissue specificity. Taking the distal active category as an example, the AS group is more tissue specific and less conserved than the non-AS group. **(D) AS events occurring in 1 or 2 assays and their relationship to purifying selection**. We identify AS events for chromatin accessibility (C, DNAse-seq and ATAC-seq), histone modification (H), methylation (M). Specifically, we compare AS events occurring in a single assay vs. AS events occurring in two assays. The genomic variants that are not AS are referred to as non-AS. The change in conservation between an AS group and the non-AS group is calculated as the log ratio of their conservation scores (fraction of rare variants, see previous panels). This ratio is negative for AS events in one assay and positive for AS events in two assays, suggesting that an AS locus (e.g. an SNV) with multiple events is more conserved. **(E) Comparing the tissue distribution of proximal active cCREs, AS & non-AS** (Top) Non-AS categories show a “U” shape trend, indicating that there are many non-AS cCREs either extremely tissue specific or ubiquitous, whereas (Bottom) AS categories demonstrate an “L” shape trend, indicating that there are many AS cCREs extremely tissue specific but not ubiquitous. **(F) Relationship between tissue specificity and conservation for active and repressed elements**. This panel shows a more in-depth comparison (in relation to panel B) of active and repressed cCREs and their relationship to conservation. As tissue specificity increases, active cCREs demonstrate less conservation, but repressed cCREs increase in conservation. **(G) Direction of imbalance for AS cCREs**. Most of these AS cCREs are detected (using H3K27ac) only in a single tissue of individual 3. However, a few AS cCREs are observed across many tissues. The heatmap shows the direction of the allelic imbalance across the most ubiquitous AS cCREs. The imbalance between the two haplotypes is measured by the fraction of unique reads mapped to each haplotype. Here, the imbalance direction is consistent across tissues. However, a few tissue-specific cCREs show directional flips between tissues.

Given our decoration strategy, we used a straightforward approach to measure tissue-specificity, which can be consistently applied to many different types of annotations (STAR Methods). Briefly, the tissue specificity of a given annotation subset (e.g., gene-proximal cCREs) is given as the fraction of elements only present in one tissue (Fig. 9B, S7.2, and STAR Methods). As expected, by this measure, only a small percentage of protein-coding genes were tissue-specific (∼8%; by either RNA-seq or mass spectrometry) (Mele et al., 2015; Wang et al., 2019), and, in comparison, pseudogenes, lncRNAs, and active regulatory elements exhibited higher tissue specificity (Fig. 9B). More notably, AS genes and regulatory elements were more tissue-specific than the corresponding non-AS ones (Fig. 9C and S7.2D). Moreover, we observed that unlike many genomic elements that mostly fall into two distinct categories, tissue-specific or ubiquitously active across all tissues (giving rise to the characteristic “U-shaped” histogram in Fig. 9E), AS elements are only tissue-specific in many different assays (an “L-shaped” histogram, depleted in “housekeeping behavior” as shown in Fig. 9E; also see Fig. S8.6A and S7.2N). Finally, for the few elements that are AS across all available tissues, we found the haplotype direction of the AS imbalance to be consistent (43 elements; Fig. 9G, S7.2I and S7.2J) (Hounkpe et al., 2021). This finding, plus the fact that we did not observe many loci where the imbalance direction flipped across tissues, supports our aggregation strategy for identifying AS events (Fig. 3A and STAR Methods).

We examined the relationship between tissue specificity and conservation (Fig. 9B and 9F). Notably, we found that annotations with higher tissue specificity have lower purifying selection (Fig. 9F; as expected, the opposite trend for repressed annotations). Consistent with previous studies, we found that AS elements are under less purifying selection than non-AS ones (Fig. 9C and S7.2N) (Chen *et al*., 2016; Fu et al., 2014; Gerstein *et al*., 2012; Onuchic *et al*., 2018). (This result is further bolstered by Fig. 5B, which shows that more AS-sensitive TF motifs are less conserved.) Conversely, we detected an increase in purifying selection for loci AS in more than one assay, perhaps reflecting their greater functional importance (e.g., methylation and histone modifications; Fig. 9D). In summary, we found that loci demonstrating more activity across tissues, haplotypes, or functional assays showed increased conservation.

Next, we analyzed the relationship between decorated regulatory elements and eQTL or GWAS SNVs. First, we systematically estimated the enrichment of eQTL and sQTL variants in cCREs active in the matched tissue type (Fig. 10A and S7.3A). The enrichment was somewhat stronger than previous studies and showed greater magnitude for proximal vs distal cCREs, especially, as expected, for sQTLs (Fig. S7.3C) (Whalen and Pollard, 2019). Notably, we compared eQTL/sQTL enrichment in AS elements with non-AS ones, finding substantially higher enrichment in AS subsets (Fig. 8A). In particular, for distal active cCREs, the AS subset showed significantly higher enrichment across all tissues, with some tissues showing considerably larger improvement (>2X improvement for those cCREs containing CTCF binding sites). Furthermore, we found that the AS subset has more TF-binding motifs than other cCREs, and these, in turn, are enriched in “AS-sensitive” motifs, suggesting greater regulatory importance (Fig. S8.6B and S7.3J).

**Fig. 10.**
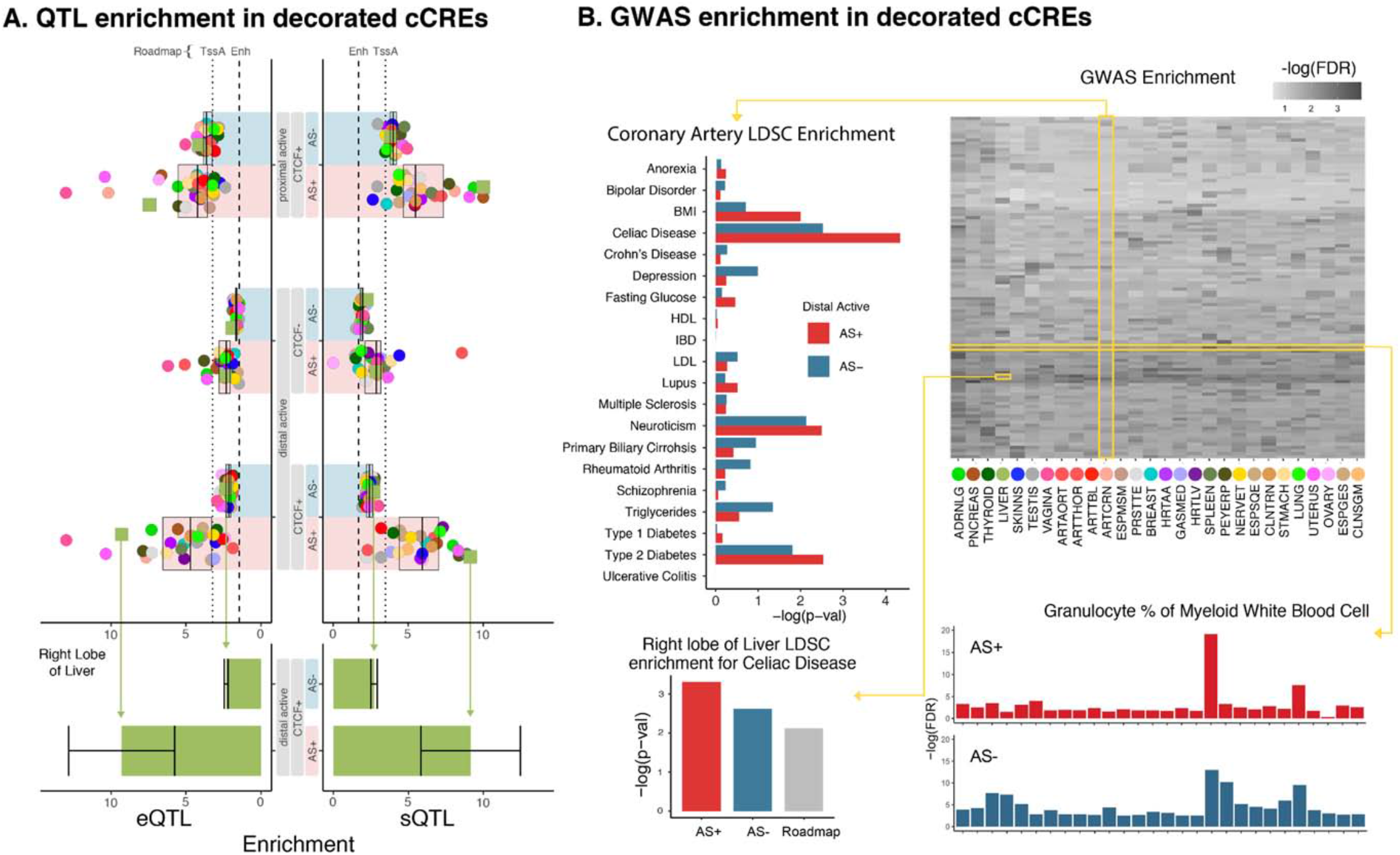
Relating Encyclopedia Decorations to QTLs and GWAS loci. **(A) QTL enrichment in decorated cCREs**. Both the expression quantitative trait loci (eQTLs) and splicing QTLs (sQTLs) are from GTEx. Colored dots show the enrichment for each tissue (using the GTEx colors from Fig. 1). Each bar shows the median enrichment over all tissues for a given decorated annotation subset. Overall, AS elements show stronger eQTL and sQTL enrichments compared to non-AS ones. As a reference, median enrichment of Roadmap “Enh” and “TssA” annotations are shown as dashed and dotted lines, respectively. The enrichments for the liver are highlighted. To estimate the robustness of this particular calculation, we resampled the genetic variants as described in Fig. S7.3, estimating a range of enrichments, shown with whiskers (see Fig. S7.3A for enrichment results for all tissues and STAR Methods for more details). **(B) GWAS enrichment in decorated cCREs**. The heatmap shows the GWAS tag-SNP enrichment of distal active CTCF cCREs. The tissues (colored according to the GTEx convention from Fig. 1A) are denoted at the bottom of the heatmap, and phenotypes (usually diseases, not explicitly labeled) run along the vertical axis. Around the heatmap, we show zoomed-in views highlighting various comparisons. At the bottom, we show that the tissue specificity of the enrichment for the trait Granulocyte % of Myeloid White Blood Cells is much stronger for AS vs. non-AS cCREs. On the left, we show higher LDSC enrichment for AS elements compared to the corresponding non-AS ones for one tissue (coronary artery) across many associated traits. We then extend this comparison (for a single tissue-trait pair) to include Roadmap annotations, shown at the bottom left.

Like eQTLs, decorated AS elements produced significantly better GWAS enrichment for disease traits (Fig. 10B and S7.3D). As a baseline, we estimated the GWAS SNV enrichment in tissue-specific regulatory elements across diverse traits (using tag-SNP and LDSC approaches; Fig. S7.3E and STAR Methods) (Bulik-Sullivan et al., 2015). We found that the subsets of these cCREs that were AS showed substantially greater enrichment than those not AS. For example, cCREs that exhibited AS activity in the coronary artery had higher enrichment for cardiovascular disease GWAS SNVs as compared to non-AS ones (Einarson et al., 2018; Emilsson et al., 2015; Khan et al., 2018; Terracciano et al., 2008). Also, for immune-associated traits, we found that enriched AS cCREs manifest better tissue specificity for the correct tissue (spleen) as compared to non-AS ones (Fig. 10B and S7.3G).

### Accessing the EN-TEx Resource

All data contained in the EN-TEx resource are fully open-consented and accessible without registration, and all of the raw sequencing data, annotations and decorations from downstream analyses and the associated tools are directly downloadable from entex.encodeproject.org (STAR Methods). Overall, they form an integrated resource including the diploid and reference signal tracks, topologically associating domain annotations, a complete catalog of AS events, lists of tissue-specific active and repressive regulatory elements, and applications of models to larger cohorts (Fig. S1.2 and STAR Methods). We have developed two novel interactive applications for visualizing the resource (Fig. S8.3, S8.4, and STAR Methods). The first performs a variety of dimensionality reductions (e.g., VAE and UMAP). The second is a chromosome “painting” tool for visualizing large-scale diploid maps of functional genomics signals. The EN-TEx cCRE decorations can also be accessed via the interactive SCREEN browser (Fig. S8.5 and STAR Methods) (Encode Project Consortium *et al*., 2020).

## DISCUSSION

The main contribution of EN-TEx is the creation of a broadly useful and accessible resource of hard-to-obtain tissues from a comprehensive catalogue of personal epigenomes and the corresponding annotations and decorations. We envision the EN-TEx resource enabling additional analyses outside of the scope of discussion here. Vignettes of methylation data related to aging or the cross-tissue epigenetics of genes associated with COVID-19 provide hints of what is possible (Fig. S8.7 and S8.8).

A key aspect of the EN-TEx resource is that it can be easily connected with other human-genome annotation resources, potentially extending them. In particular, by training on the GTEx eQTL catalog, we show how to build a model that can transfer eQTLs from an easily obtained tissue to one that is harder to get. With this approach, we leverage the fact that EN-TEx represents a uniform collection of epigenetics data from hard-to-obtain tissues. We also show that EN-TEx can decorate the ENCODE regulatory elements to give a unified view of tissue specificity and conservation and provide subsets of elements that are particularly enriched in GWAS variants. We imagine that in the future, one could apply these strategies to connect with and extend other genome resources beyond ENCODE and GTEx.

The second aspect of EN-TEx is that we can leverage the scale of the AS catalog to develop models illuminating the biological impact of variants. These models suggest that the local sequence context around a variant is the dominating factor in determining its impact, with certain TF motifs being particularly sensitive to mutations. That said, it is not just the TF motif right at the SNV position that is relevant, but the surrounding sequence (within a ∼250 bp window). This suggests that determining whether or not a particular site is AS may have to do with other, potentially interacting, TFs binding nearby. For instance, a particular TF-binding site could be stabilized from mutational impact (and AS behavior) by being only one of the many DNA-binding sites of a large hetero-oligomeric complex. Alternatively, redundant binding sites for a single factor may act as “backup” against the effects of one mutation (Payne and Wagner, 2015). The concept of “buffering” posits a mechanism for this (Maurano *et al*., 2015).

Although the four individuals were considered healthy, their genomes contain many rare and potentially deleterious variants. These are not normally accessible by traditional QTL studies, which are best powered for common variants. In contrast, our AS analysis and models can provide information on rare variants, and, in this regard, the EN-TEx resource is particularly informative to precision medicine. Moreover, the approach piloted by EN-TEx could be scaled in a straightforward fashion to more individuals in the future, providing information on additional rare variants. This contrasts to the situation for common variants, where a scale-up would provide diminishing amounts of additional information (STAR Methods S2.4).

A final contribution of the EN-TEx resource is demonstrating how the diploid genome is important for future human functional genomics. In particular, we show that diploid genomes provide more accurate quantification of differential expression and regulatory activity, which is essential for disease studies. Furthermore, the matching of individuals and tissues in EN-TEx allows a precise ascertainment of the relative contribution of inter-tissue and inter-individual variation. We envision that in the near future, with the decreased cost of sequencing, generating a matched personal genome sequence as an accompaniment to each functional genomics experiment will become the norm for large-scale studies. Thus, the EN-TEx personalized epigenomics approach for analyzing the impact of genome variation will necessarily become commonplace, potentially providing the benefits described here towards more exact precision medicine.

## Notes

### Competing Interest Statement

The authors have declared no competing interest.

### Summary of Updates

Revised text, figures and supplement.

