## Supplementary Methods for "The EN-TEx resource of multi-tissue personal epigenomes & variant-impact models"

STAR★Methods for

**Table of Contents**

List of Supplementary Files and Corresponding Sections 11

Glossary 13

Overview 15

S1. Supp. Content for Main Text Section "Improvements in Genome Analysis from Uniform Multi-tissue Data Collection & Diploid Genome Mapping" 16

S1.1. Personal Genome Construction 16

S1.1.1. Sequencing of the Personal Genome 16

Long Read Sequencing 16

Short Read Illumina Sequencing 16

S1.1.2. Personal Genome Construction and Variant Calling 17

S1.1.3. Refining Novel Insertion Sequences with Iris 18

S1.1.4. Assigning Parental Origin by Imprinted Genes 19

S1.2. Functional Genomics Data Stack 20

S1.2.1. RNA Sequencing 20

S1.2.2. RAMPAGE 20

S1.2.3. eCLIP 20

S1.2.4. Histone ChIP-seq 21

S1.2.5. Transcription Factors ChIP-seq 21

S1.2.6. ATAC-seq 22

S1.2.7. DNase-seq 22

S1.2.8. WGBS 22

S1.2.9. DNAme Array 22

S1.2.10. Hi-C 23

Hi-C Data Processing Details 23

S1.2.11. Proteomics 24

Liquid Chromatography with Tandem Mass Spectrometry (LC-MS/MS) Analysis 24

Personal Proteome Database 25

MS Identification and Quantification 25

Peptide and Gene Identification and Quantification Results 26

RNA-seq Comparison 26

Novel Gene Discovery 26

S1.3. Mapping Functional Genomics Data to the Personal Genomes 27

S1.4. Variation Analysis of cCRE Activity 28

S1.4.1. Visualizing the Variation of cCRE Activity with JIVE 28
S1.4.2. Using a Regression-based Approach to Quantify Activity Variation 28
S2. Supp. Content for Main Text Section “Large-scale Determination of AS SNVs & Construction of the AS Catalog” 30
S2.1. Calling AS events 30
S2.1.1. Allele-specific Expression (ASE), Binding (ASB), and Chromatin Accessibility (ASCA) 30
Overview 30
Mapping 30
Filtering and Assessment of Read Imbalance at Heterozygous SNVs (hetSNVs) 31
S2.1.2. Allele-specific Methylation (ASM) 32
S2.1.3. Hi-C - Allele-Specific Interactions 33
Creating Haplotype-specific Interaction Matrices 33
AS Interactions 33
S2.1.4. Proteomics - Allele specific Peptide (ASP) Analysis 33
S2.2. Allele-specific Functional Elements 34
S2.2.1. Genes and cCREs 34
S2.2.2. Correlation Between AS Genes and Diseases 34
S2.2.3. Gene Ontology Enrichment Analysis of AS Genes 34
S2.3. Aggregation Across Tissues and Assays 35
S2.3.1. AS Expression and Binding 35
S2.3.2. Methylation 35
S2.4. Generalizability of the AS catalog 35
S2.5. Calling AS Events in External Datasets Used for Validation (Roadmap and NA12878) 36
S2.6. High-confidence and High-power Call Sets 37
S2.7. Integration with the ClinGen Allele Registry 38
S2.8. Examples of Coordinated AS Activity Across Assays 38
S3. Supp. Content for Main Text Section “Interrelating SVs & Chromatin Modifications” 39
S3.1. Analysis of Structural Variants 39
S3.2. Associating SVs with Expression Quantitative Trait Loci (eQTLs) 40
S3.3. Aggregating the Impact of SVs on Neighboring Chromatin 42
S4. Supp. Content for Main Text Section “Generalized Application #1: Predicting AS ChIP-seq Activity from Nucleotide Sequence” 43

S4.1. Relation between AS SNPs and Transcription Factor (TF) Motifs 43
S4.2. Allele-Specific Effect Prediction with the BERT Model 44
S5. Supp. Content for Main Text Section “Generalized Application #2: Models Interrelating AS Activity at Promoters and Genes” 45
S5.1. Prediction of Promoter AS Activity With a Random Forest Model 45
S5.2. Compatibility Between Assays 46
S5.2.1. Compatible and Incompatible: Single Chromatin Mark vs. Gene Expression 46
S5.2.2. Compatibility with AS Proteomics 47
S5.2.3. Enrichment of ASE Genes Near ASM Promoters 47
S6. Supp. Content for Main Text Section “Generalized Application #3: Using the EN-TEx Resource to Extend eQTL Annotations to Hard-to-obtain Tissues” 48
S6.1. Correlation Between Chromatin Features and eQTL Activity 48
S6.2. Building a Predictive Model That Transfers eQTLs From a Donor Tissue to a Target Tissue 48
S6.3. Model Performance, Validation, and Novel Application 49
S6.4. Model Interpretation 50
S7. Supp. Content for Main Text Section “Generalized Application #4: Decorating ENCODE Regulatory Elements with EN-TEx Tissue & AS Information” 51
S7.1. Decoration of Candidate Cis Regulatory Elements (cCREs) from ENCODE Encyclopedia 51
S7.1.1. Signal Normalization Method 51
S7.1.2. Decoration of Regulatory Annotations 52
S7.1.3. Repressed Regions 53
S7.1.4. Validating Annotations Using 3D Genome Organization 54
S7.2. Tissue Specificity 55
S7.2.1. Tissue-specific Results 55
S7.2.2. Tissue Specificity of ASB and ASE 56
S7.2.3. The Effect of Tissue Specificity on Conservation 56
S7.2.4. The Relationship Between Purifying Selection and Regions Exhibiting Allele Specificity 56
S7.3. Relating Encyclopedia Decorations to QTLs & GWAS Loci 57
S7.3.1. QTL Enrichment Analysis 57
S7.3.2. GWAS Enrichment Analysis 58
S8. Supp. Content for Main Text Section “Discussion” 59
S8.1. The EN-TEx Supplemental Data Repository 59

S8.2. Open-consent of Data 59
S8.3. Explorer Tool 59
S8.4. The EN-TEx Chromosome-Level Data Visualization Tool 60
S8.5. Additional Data Exploration with SCREEN 61
S8.6. Providing Evidence for the Buffering Hypothesis Using AS cCREs and Housekeeping Genes 61
Supplementary Figures 63
Figure S1. Supp. Figures for Main Text Section “Improvements in Genome Analysis from Uniform Multi-tissue Data Collection & Diploid Genome Mapping” 63
Figure S1.1a. Personal genome construction 65
Figure S1.1b. Phase blocks of personal genomes with parental origins 66
Figure S1.1c. Consistency of ASE imbalance direction in known imprinted genes across tissue samples (individual 3) 67
Figure S1.2a. Functional genomics data 69
Figure S1.2b. Example of reference-aligned genome-wide Hi-C maps for the skeletal muscle tissue of two individuals 70
Figure S1.2c. Number of paired reads and number of contacts from reference-aligned genome-wide Hi-C contact maps 71
Figure S1.2d. A/B compartment annotation of four individuals and two tissues for chromosome 1 72
Figure S1.2e. A/B compartments cluster based on tissue in autosomes or sex in chromosome X 73
Figure S1.2f. Summary of significant interactions determined by FitHiC2 74
Figure S1.2g. Comparison of TopDom TAD calls for EN-TEx individuals and available Hi-C tissues 75
Figure S1.2h. Observed personal peptide summary 76
Figure S1.2i. Novel peptides 77
Figure S1.3a. Mapping to personal genomes 78
Figure S1.3b. Gene expression and cCRE activity obtained from mapping to personal genomes 80
Figure S1.3c. Differential gene expression and cCRE activity between mapping to reference and personal genomes 81
Figure S1.4a. Variation explained between two experiments corrected by replicates 82
Figure S1.4b. Similarity between the signals of two functional genomic experiments 83

Figure S1.4c. Variation explained between proteomics and RNA-seq data 84
Figure S1.4d. Comparing the explained variances calculated respectively using matched and unmatched data 85
Figure S2. Supp. Figures for Main Text Section “Large-scale Determination of AS SNVs & Construction of the AS Catalog” 86
Figure S2.1a. Workflow of the AlleleSeq2 pipeline 87
Figure S2.1b. Generating haplotype-specific signal tracks 89
Figure S2.1c. Data used to generate signal tracks 90
Figure S2.1d. Distribution and fraction of the number of hetSNVs associated with AS behavior across different EN-TEx donors, tissues, and assays 91
Figure S2.1e. Schematic showing how AS methylation is calculated 92
Figure S2.1f. ASM calls in known imprinting control regions 93
Figure S2.1g. Generating haplotype-specific Hi-C 94
Figure S2.1h. Haplotype-specific contact maps for Chr20 generated using the personal genome coordinates 95
Figure S2.1i. Number of Hi-C contacts obtained from haplotype specific Hi-C contact maps 96
Figure S2.2a. Distribution and fractions of the number of genomic elements (genes and cCREs) associated with AS behavior across different EN-TEx donors, tissues, and assays 97
Figure S2.2b. Gene ontology enrichment analysis of AS genes 98
Figure S2.3. Summary of AS catalog 99
Figure S2.4. Numbers of AS hetSNVs detected in different EN-TEx assays 101
Figure S2.5. Construction of validation dataset from AS events in non-EN-TEx datasets 104
(A) Distribution and fraction of the number of hetSNVs associated with AS behavior detected in NA12878 and Roadmap individuals STL002 and STL003. (B) Datasets used for calling AS events in the Roadmap individuals. 104
Figure S2.6a. Numbers of AS hetSNVs detected from RNA-seq in different tissues from individual 3 105
Figure S2.6b. Validation of high-power AS calling methods 106
Figure S2.8a. Heatmap to show haplotype specificity of Chr X for all assays and tissues from individual 3 107
Figure S2.8b. Chromosome painting of ChrX using RNA-Seq and ChIP-Seq in both haplotypes of individual 3 in two tissues 108
Figure S2.8c. XACT locus on ChrX is shown to have haplotype-specific chromatin interactions with an upstream region. 109

Figure S2.8d. Coordinated AS activity in X chromosomes. 110

Figure S3. Supp. Figures for Main Text Section “Interrelating SVs & Chromatin Modifications” 111

Figure S3.1. Analysis of SVs 112

Figure S3.2a. ASE events associated with indels and SVs. 114

Figure S3.2b. Indels and SVs associated with ASE 115

Figure S3.2c. An indel that potentially changes gene expression 116

Figure S3.2d. Shadow figure associated with Figure 4D 117

Figure S3.2e. SVs potentially linked to eQTLs 118

Figure S3.2f. Shadow figure associated with Figure 4F 120

Figure S3.2g. Novel splicing variants of *PCCB* 122

Figure S3.2h. Novel splicing variants of *TRDN-AS1* 123

Figure S3.2i. Null alleles potentially caused by deletion of entire exons. 124

Figure S3.2j. Homozygous deletions in individual 2 but not individual 1. 125

Figure S3.3a. Calculating changes in the chromatin state in the SV neighborhood 126

Figure S3.3b. Changes in the chromatin state of SV neighborhoods 127

Figure S4. Supp. Figures for Main Text Section “Generalized Application #1: Predicting AS ChIP-seq Activity from Nucleotide Sequence” 128

Figure S4.1a. AS enriched motif ranking 129

Figure S4.1b. Fraction of AS/non-AS SNVs overlapping with the motif 130

Figure S4.2a. Performance of trained allelic effect prediction models 131

Figure S4.2b. Tissue-specific performance in H3K27ac (with Roadmap external validation) 132

Figure S4.2c. Additional examples of attention patterns learned by the model 133

Figure S4.2d. Motifs that peak in the proximity of the AS CTCF SNPs 134

Figure S5. Supp. Figures for Main Text Section “Generalized Application #2: Models Interrelating AS Activity at Promoters and Genes” 135

Figure S5.1a. Performance validation of models predicting allele-specific bound promoters for each assay 136

Figure S5.1b. Association analysis between features and the promoter allele-specific binding events 137

Figure S5.1c. Scatter plots for the association analysis between features and the promoter ASB events 138

Figure S5.1d. Features tested but not informative for the prediction of ASB promoters 139

Figure S5.1e. Contingency table for ASE genes and ASB promoters 140

Figure S5.1f. Features and performance of a model to predict ASE for a gene from ASB on the associated promoter 141

Figure S5.2a. Compatibility between allelic events 143

Figure S5.2b. Flow chart of filtering and AS expression/AS Proteomics comparison 144

Figure S5.2c. Compatibility between AS mRNA and AS peptide calculations 145

Figure S5.2d. Enrichment of ASE genes near ASM promoters 146

Figure S6. Supp. Figures for Main Text Section "Generalized Application #3: Using the EN-TEx Resource to Extend eQTL Annotations to Hard-to-obtain Tissues" 147

Figure S6.1a. Chromatin features can help prioritize causal eQTLs 148

Figure S6.1b. Chromatin-marked loci associated with eQTL activity 149

Figure S6.2a. Schema of the predictive model using skin as the donor tissue and tibial artery as the target tissue 150

Figure S6.2b. Number of eQTLs available from each donor tissue 151

Figure S6.2c. List of predictive features employed by the random forest model to predict eQTL activity in a target tissue 153

Figure S6.3a. Description of the metrics used to evaluate the random forest model 154

Figure S6.3b. Performance of the random forest submodels by donor tissue 155

Figure S6.3c. Performance of the random forest submodels by target tissue 156

Figure S6.3d. Proportion (%) of blood eQTLs from Vosa et al. 2021 that can be transferred to each EN-TEx tissue 157

Figure S6.3e. Number of potentially novel eQTLs predicted for each EN-TEx tissue that are not present in the GTEx catalog 158

Figure S6.4a. Dissecting the contribution of features to predicting tissue-specific eQTL activity 159

Figure S6.4b. By analyzing the chromatin activity of donor-tissue eQTLs in the target tissue and the tissue specificity of their eGenes, we can identify eQTLs active in the target tissue 160

Figure S7. Supp. Figures for Main Text Section "Generalized Application #4: Decorating ENCODE Regulatory Elements with EN-TEx Tissue & AS Information" 161

Figure S7.1a. Data preprocessing 162

Figure S7.1b. Framework of cCRE decoration 163

Figure S7.1c. Framework of cCRE decoration in the spleen 164

Figure S7.1d. Number of cCREs in various tissues 165

Figure S7.1e. cCRE decoration results matrix 166
Figure S7.1f. Identifying fully repressed elements independent of cCREs 167
Figure S7.1g. cCRE enrichment with respect to A/B compartments 168
Figure S7.2a. The number of transcribed genes in tissues 169
Figure S7.2b. Tissue specificity of transcribed genes 170
Figure S7.2c. Gini index of gene expression level across tissues 171
Figure S7.2d. Tissue specificity of different subgroups of cCREs vs. genes and epigenomic peaks 172
Figure S7.2e. Tissue specificity of different subgroups of cCREs 173
Figure S7.2f. Tissue specificity of RAMPAGE data at TSSs of protein-coding genes 174
Figure S7.2g. ASE genes across different tissues of individual 3 176
Figure S7.2h. Haplotype 1 allele ratios (number of haplotype 1 reads over the total number of reads) for expression of genes that are accessible across all tissues and AS in at least one tissue of individual 3 178
Figure S7.2i. Annotation of pan-tissue H3K27ac AS+ cCREs of individual 3 179
Figure S7.2j. Annotation of pan-tissue ASE genes of individual 3 180
Figure S7.2k. Rare DAF for active, bivalent, and repressive cCREs in increasing tissue count 181
Figure S7.2l. Conservation of enhancer decorations 182
Figure S7.2m. Conservation of active and repressed cCREs for tissue-specific and ubiquitous categories 183
Figure S7.2n. Conservation of regions exhibiting AS activity 184
Figure S7.3a. eQTL and sQTL enrichment in cCREs 185
Figure S7.3b. Roadmap annotations 187
Figure S7.3c. QTL enrichment in cCREs: EN-TEx vs. Roadmap 188
Figure S7.3d. Framework of GWAS enrichment analysis 189
Figure S7.3e. Stratified LDSC enrichment: Comparing EN-TEx AS, non-AS, and Roadmap annotations 190
Figure S7.3f. GWAS enrichment: cCREs vs. cCREs with 500 bp extensions 191
Figure S7.3g. GWAS enrichment across tissues 192
Figure S7.3h. GWAS enrichment for Roadmap annotations 193
Figure S7.3i. GWAS enrichment for AS+ vs. AS- cCREs 194
Figure S7.3j. Enrichment of AS sensitive or AS non-sensitive motifs in cCREs 195
Figure S8. Supp. Figures for Main Text Section “Discussion” 196

Figure S8.3. Explorer tool 198
Figure S8.4a. Screenshot of the EN-TEx chromosome painting tool 199
Figure S8.4b. Examples of the chromosome painting tool 201
Figure S8.5. Viewing EN-TEx decoration on the SCREEN website 202
Figure S8.6a. Allelic specificity of housekeeping genes 203
Figure S8.6b. Enrichment of TF motifs in CTCF+ cCREs 204
Figure S8.7. Predicting the ages of tissues from their DNA methylation 205
Figure S8.8. Histone ChIP-seq data for COVID19-related genes 206
Figure S8.9. Coverage of EN-TEx AS hetSNVs in GTEx corresponding individual tissue 207
References 208

#### List of Supplementary Files and Corresponding Sections

| | |
|---|---|
| phased_block.tar.gz | S1.1.4 |
| fithic2_out.tar.gz | S1.2.10 |
| TopDomTADcalls.tar.gz | S1.2.10 |
| Supp_data_proteomics.xlsx | S1.2.11 |
| Supp_DE_genes.tsv | S1.3 |
| Similarity_of_functional_genomic_activities_of_cCREs.xlsx | S1.4.2 |
| normalized_proteomics_RNA-seq.dat | S1.4.2 |
| sample_signal_track.tar.gz | S2.1.1 |
| AlleleSeq2_workflow_examples.tar.gz | S2.1.1 |
| hetSNVs_default_AS.tsv | S2.1.1 |
| ENTEx.TissueStacked.phased.final.txt | S2.1.2 |
| hic_files.tar.gz | S2.1.3 |
| genes_default_AS.tsv | S2.2.1 |
| cCREs_default_AS.tsv | S2.2.1 |
| Associated_AS_Disease_Genes.xlsx | S2.2.2 |
| hetSNVs_pooled_AS.tsv | S2.3.1 |
| ENTEx.TissueAggregated.final.txt | S2.3.2 |
| pgenome_NA12878.tar.gz | S2.5 |
| pgenome_STL002.tar.gz | S2.5 |
| pgenome_STL003.tar.gz | S2.5 |
| hetSNVs_default_AS_validation.tsv | S2.5 |
| hetSNVs_high-confidence_AS.tsv | S2.6 |
| hetSNVs_high-power_AS.tsv | S2.6 |
| Supp_Data_SVs_associated_with_eQTL.xlsx | S3.2 |
| motif_ranking.tsv | S4.1 |
| ASB-predictions-on-GTEx-cohort.tsv | S5.1 |
| AS_allhets_alt_allele_ratio_eqtl_intersect.tsv | S5.2.1 |
| Supp_Data_compatibility.xlsx | S5.2.2 |
| predictions.blood.eQTLs.tar.gz | S6.3 |
| cCRE_histoneSignals_qnorm.tar.gz | S7.1.1 |
| cCRE_Decoration.matrix | S7.1.2 |
| active.combined_set.txt.zip | S7.1.2 |
| bivalent.combined_set.txt.zip | S7.1.2 |
| repressed.combined_set.txt.zip | S7.1.2 |
| stringent.regions.MF.hg38.bed | S7.1.2 |
| ENTEx_fully_repressed_regions_independent_of_cCREs.bed | S7.1.3 |
| Tissue_Specificity.zip | S7.2.1 |
| QTL_enrichment.zip | S7.3.1 |
| GWAS_enrichment.zip | S7.3.2 |
| ENTEx.Explorer.cCRE.Combined.zip | S8.3 |

| | |
|---|---|
| ENTEx.Explorer.Expression.Combined.zip | S8.3 |
| ENTEx.Proteomics.cCRE.Combined.zip | S8.3 |

#### Glossary

Below are terms and acronyms that are frequently used in the manuscript.

| | |
|---|---|
| Assay for Transposase-Accessible Chromatin using sequencing, ATAC-seq | A method to determine genomic regions with open chromatin (i.e., sequences not wrapped in nucleosomes). The method uses transposase to insert tags into open chromatins, allowing these regions to be identified by DNA sequencing. |
| Allele specific, AS | Most genomic regions, e.g., an open reading frame, exist on both sister chromosomes. In genetics, the two copies of the same genomic regions are called alleles of the genomic regions. The biological activity, such as gene expression and histone modification, from one allele can be much higher than that from the other allele. In such a case, the biological activity of the particular genomic region is allele specific, or imbalanced between the two alleles. |
| Allele-specific binding, ASB | The expression level of a gene from one allele is significantly higher than that from the other allele. |
| Allele-specific expression, ASE | The histone modification level (quantified by the number of reads from ChIP-seq) of one allele is significantly higher than that of the other allele. |
| Candidate cis regulatory element, cCRE | Genomic region that potentially regulates the expression of nearby and/distal genes. Most of the sequences are non-coding sequences. |
| Chromatin immunoprecipitation followed by sequencing, ChIP-seq | A method to determine the DNA sequences bound by a given protein. Proteins, along with the DNA bound by the protein, are isolated through immuno-precipitation, and then the identity of the DNA is revealed by sequencing. Even proteins that carry post-translational modifications, e.g., acetylation, can be pulled down by special antibodies. |
| DNase-seq | Another method to determine open genomic regions. DNA that is not wrapped in nucleosomes is easily degraded by DNase, and generates less reads when sequenced. |
| Quantitative trait locus, QTL | The levels of a quantitative trait, e.g., the expression levels of a particular gene, of an individual often correlate with the specific alleles that the individual carries at particular loci. These loci are called QTLs. If the trait of interest is the expression level of a gene, the loci are called eQTLs. |
| Haplotype | All DNA sequences that are inherited from one parent. |
| Maternal/Paternal | Genetic materials inherited from the mother/father. |

| | |
|---|---|
| Phase | When variants are identified from genomic sequencing results, it is possible to determine whether they come from the same sister chromosomes. Variants that come from a continuous block of the same chromosome are in the same phase. |
| RNA sequencing, RNA-seq | A method to determine the levels of RNA expressed. Total RNA (or a subset, e.g., mRNA) is isolated from samples, converted to cDNA, and sequenced. The last step determines the amount and sequence of each RNA, which allows the RNA to be mapped to specific loci (e.g., a gene) in the genome. |
| Structural variant, SV | Large (>= 50 bp) mutations, such as insertions, deletions, and inversions |
| Transposable element, TE | DNA sequences that can relocate in the genome. The relocation can create structural variants at the original and the new locations of the TEs. TEs can be classified based on their sequences. Examples of TE families are SINE (short interspersed nuclear elements), LINE (long interspersed nuclear elements), and SVA (SINE-VNTR-Alus) retrotransposon. |

#### Overview

This document provides a comprehensive and organized reference to all datasets, methods, and analyses associated with the EN-TEx project. The structure of this document is presented in a parallel fashion to the main text. Results and figures in each section of the main text are documented in the relevant subsections titled "Supp. content for 'Main-text-section-title'" within this supplement. Supplementary figures are numbered based on the secondary heading of their relevant supplementary text section (e.g., Figure S1.1a-f are associated with Section S1.1). Unless specified, we refer to the supplementary text in a general manner (e.g., "see Supplement") in the main text. However, we explicitly refer to the supplementary figures with exact figure numbers. All supplementary figures are attached at the end of this document. A list of supplementary figures can be found as part of the table of contents.

The large dataset produced under the EN-TEx project includes more than 25 different functional genomic assays sampled in more than 30 tissues of four individuals (Individual 1-4, also referred as ENC-001 to ENC-004 interchangeably). These assays include genotyping, RNA-seq, transcription factor (TF) chromatin immunoprecipitation sequencing (ChIP-seq), histone ChIP-seq, DNase I hypersensitive sites sequencing (DNase-seq), Assay for Transposase-Accessible Chromatin using sequencing (ATAC-seq), *in-situ* high-throughput chromosome conformation capture (Hi-C), and more. The EN-TEx dataset also includes processed data, such as an allele-specific (AS) heterozygous single-nucleotide polymorphism (SNP) catalog and candidate cis-regulatory elements (cCRE) annotation. Together, this dataset provides a valuable resource for studies of gene regulation and precision medicine. However, due to the richness of the data, it is challenging to include all of the details within the main text of this paper.

The data resource is organized in a hierarchical pyramid-like structure, where the raw data files are located at the base. On top of those are the processed data, and finally the high-level summaries lie at the top. Specifically, the main text summarizes everything in a broad manner, providing a macroscopic view of this study. Raw data, the cornerstone of this study, are hosted online at the ENCODE portal. Links to the metadata, bam files, and other raw data can be found at the "Raw Data" link on the EN-TEx data portal website (ENTEx.encodeproject.org). The processed data files, located in the middle of the pyramid, are described in detail in their corresponding sections in this document. These files are referred to as "File: file_name" in the text and are hosted on the EN-TEx data portal website with the same file names. Additionally, on the website, each file is followed by the supplement text section number that contains the description of that file.

### S1. Supp. Content for Main Text Section “Improvements in Genome Analysis from Uniform Multi-tissue Data Collection & Diploid Genome Mapping”

#### S1.1. Personal Genome Construction

##### S1.1.1. Sequencing of the Personal Genome

Figure S1.1a summarizes the technologies used to sequence the whole genomes of the four individuals.

**Long Read Sequencing**
To prepare samples for PacBio sequencing, genomic DNA was isolated as previously described (Ardui et al., 2018; Nattestad et al., 2018) and evaluated for purity and quantity using UV-Vis (Nanodrop 1000, Thermo Fisher) and fluorometric (Qubit, Thermo Fisher) assays. DNA sizing was checked on the Pippin Pulse (Sage Science). Samples all exhibited a mode size above 50 kbp (most above 100 kbp) and were considered good candidates for PacBio sequencing. DNA was sheared using gTUBEs (Covaris) to a mode size of ~15 kbp using 3,200 RPM for four passes on an Eppendorf 5424R centrifuge. The sheared material was subjected to SMRTbell library preparation using the Template Prep Kit v1 (PacBio). After checking for size and quantity, the material was size fractionated on the Blue Pippin instrument (Sage Science) using the protocol “0.75% 1-18kb v2”, size-based separation mode, and target value 3,400 in well #12. Fractions were checked via fluorometric quantitation (Qubit) and pulse-field sizing (FEMTO Pulse). For CLR sequencing, isolated gDNA was SMRTbell library prepared using the Express Kit v2 (PacBio) and subjected to size selection on a Blue Pippin instrument (Sage Science) with a 40 kbp size cutoff. Libraries were loaded on a Sequel II using v2.0 binding and v2.0 sequencing kits, no pre-extension, and 15-hour movie times.

For nanopore sequencing, samples were sheared to approximately 60 kb and size selected by SRE (Circulomics). Fragmented DNA was prepared for sequencing following the manufacturer's instructions with the SQK-LSK110 kit (Oxford Nanopore). Prepared libraries were sequenced on a PromethION 24 with PROM0002 flowcell for 72 hours. One nuclease flush and reload was performed at 24 hours. Live high accuracy base calling was used.

**Short Read Illumina Sequencing**
We generated and analyzed Illumina whole-genome sequencing (WGS) data for each of the four human genome samples. WGS libraries were prepared using the TruSeq DNA PCR-Free Library Preparation Kit (Illumina) in accordance with the manufacturer’s instructions. Briefly, 1 µg of DNA was sheared using a Covaris LE220 sonicator (adaptive focused acoustics). DNA fragments underwent bead-based size selection and were subsequently end-repaired, adenylated, and ligated to Illumina sequencing adapters. Final libraries were evaluated using fluorescent-based assays, including qPCR with the Universal KAPA Library Quantification Kit and Fragment Analyzer (Advanced Analytics) or BioAnalyzer (Agilent 2100). Libraries were

sequenced on an Illumina NovaSeq 6000 sequencer using 2 x 150 bp cycles to a minimum depth of 30X.

##### S1.1.2. Personal Genome Construction and Variant Calling

Personal genomes were constructed from a combination of long-range Hi-C reads, 10x Genomics linked reads, and long reads (Pacbio reads for individuals 2 and 3 and Oxford Nanopore reads base called with Guppy v4 for individuals 1 and 4) using the reference-guided assembler CrossStitch (Figure S1.1a) (https://github.com/schatzlab/crossstitch) (Alonge et al., 2020). This pipeline has been used in several other studies of human and non-human genomes with as many as 100 different genomes at once (Aganezov et al., 2020; Alonge *et al.*, 2020; Chen et al., 2019) for comprehensive single-nucleotide variant (SNV), insertion and deletion (indel), and structural variant (SV) calling. Notably, previous studies have shown that it is possible to accurately identify and phase SVs with variants identified from 10x linked reads and Hi-C data using the approach in CrossStitch to near chromosome-level resolution (Cretu Stancu et al., 2017).

Specifically, the following preprocessing steps were performed:

1. Align all reads (Hi-C, 10X, PacBio) to the human reference (GRCh38).
2. Call small variants from the linked reads with LongRanger (ver. 2.1.2) (https://support.10xgenomics.com/genome-exome/software/pipelines/latest/what-is-long-ranger).
3. Phase small variants with HapCUT2 (ver. 1.1) (Edge et al., 2017) using HiC and 10X data.
4. Call large SVs with Sniffles (ver. 1.0.11) (Sedlazeck et al., 2018) using default parameters in samples sequenced with Pacbio and using --min_homo_af 0.93 in samples sequenced with Oxford Nanopore. Additionally, in samples sequenced with Pacbio, call SVs with pbsv (ver. 2.2.1) (https://github.com/PacificBiosciences/pbsv) and merge the call sets with SURVIVOR (ver. 1.0.6) (Jeffares et al., 2017), discarding SVs that were only identified by pbsv.
5. Filter SVs with low read support (fewer than 10 reads in samples 2 and 3, fewer than 3 reads in sample 1, and fewer than 4 reads in sample 4). Additionally, in samples 2 and 3, filter SVs labeled by Sniffles with the IMPRECISE INFO flag.

Then, the CrossStitch software (commit 53f64af) performed the following steps to obtain a personal genome:

6. Refine SVs with Iris (ver. 1.0) (Chen *et al.*, 2019).
7. Phase long reads using the phased small variants with which they overlap using an analogous approach to the NanoSV algorithm (Cretu Stancu *et al.*, 2017).
8. Phase large SVs based on the phasing of the reads supporting them (Figure S1.1a).
9. Integrate (“splice”) the phased variants into two copies of each human chromosome to produce personal diploid chromosome sequences using vcf2diploid (ver. 1.0) (Rozowsky et al., 2011).

10. Assign one sequence of each chromosome to pseudo-haplotype 1 and the other to pseudo-haplotype 2.

Note that each chromosome was phased independently from the other chromosomes, so that pseudo-haplotype 1 of one chromosome may correspond to pseudo-haplotype 2 of another chromosome. Unfortunately, the available data are insufficient to distinguish such cases and assemble full haplotypes genome wide. However, using maternally and paternally imprinted genes we were able to fully resolve the parent of origin for most haplotypes of most chromosomes (see S1.1.4 below).

In all four samples, the use of 10x and Hi-C data resulted in chromosome-arm-length phase blocks for all autosomes (Figure 1B and Figure S1.1a). Specifically, the N50 of the phase blocks were 133.65 Mb, 133.68 Mb, 134.99 Mb, and 135.00 Mb for the four individuals, respectively. In addition, in both samples for which long reads were used, more than 90% of large indels were able to be confidently phased with CrossStitch.

For all four individuals, variant call format (VCF) files containing the SNVs and indels are accessible from the ENCODE portal (Jou et al., 2019) (see Figure S1.1a for accession numbers).

We adopted the reference-guided approach over alternative *de novo* assembly-based approaches because it gave more accurate and comprehensive results for the genome data available. For example, for individual 2 we also applied the leading PacBio-based *de novo* assembly algorithm FALCON-unzip (Chin et al., 2016) to assemble the genome *de novo*, but this resulted in a contig N50 of only 7.0 Mbp. Aligning the FALCON-unzip contigs to GRCh38 using MUMmer (Kurtz et al., 2004) /Assemblytics (Nattestad and Schatz, 2016) identified only <13,000 SVs compared with >18,000 for our reference-guided approach, with thousands of variants, especially heterozygous variants, unresolved. *De novo* assembly of the 10X Genomics linked reads or Illumina paired-end reads was even more limited, with contig N50 values of only 72 kbp and 13 kbp using the 10X Genomics Supernovo (Weisenfeld et al., 2017) and Illumina Megahit (Li et al., 2015) assemblers, respectively. The 10x Genomics *de novo* assembly was particularly problematic for SV identification, as we observed an enrichment for ~200 bp insertions not observed with other sequencing technologies. In communication with 10X Genomics, we found these to be false positives derived from their assembly algorithm (Aganezov *et al.*, 2020). However, we and others found the SNV and indel calls to be highly accurate, especially within repetitive elements that could not be mapped using standard short-read paired-end sequencing.

##### S1.1.3. Refining Novel Insertion Sequences with Iris

Iris is an established method for refining the breakpoints and sequences of insertion variants (Chen *et al.*, 2019). It has been used in a number of contexts (Aganezov *et al.*, 2020; Alonge *et al.*, 2020). Each of the calls, when taken directly from the variant caller, consists of an insertion sequence obtained from the alignment of a single representative read, and Iris improves upon this sequence by integrating all of the reads that support the variant's presence. The tool gathers the sequences of all of the reads listed in the RNAMES INFO field output by Sniffles, extracts the original insertion sequence with the surrounding context from the reference genome, and uses the gathered reads to polish this sequence with racon (ver. 1.4.0) (Vaser et al., 2017). Then, this polished sequence is aligned back to the reference with minimap2 (ver. 2.17) (Li, 2018), and a refined insertion sequence is obtained. If no insertion is found from this alignment, which has a similar length to that of the original variant call, Iris falls back on the original sequence to ensure it does not mask variants in more difficult-to-map regions.

We benchmarked the performance of Iris using data from HG002, a sample sequenced as part of the Genome in a Bottle release (ftp://ftp-trace.ncbi.nlm.nih.gov/giab/ftp/data/AshkenazimTrio/HG002_NA24385_son/). In this individual, we called SVs separately using Oxford Nanopore (ONT) data and Pacbio Circular Consensus Sequencing (CCS) data, both sequenced to ~50x coverage using the ngmlr aligner (Sedlazeck *et al.*, 2018) and the Sniffles variant caller. Because of the high accuracy of the CCS reads, we used the insertion sequences obtained from these calls as a proxy for the ground truth to evaluate the accuracy of the ONT calls. We compared the CCS and ONT call sets before and after refining the ONT calls with Iris. In each comparison, we evaluated all of the variant calls in the CCS dataset, which had an ONT variant call within 10 kbp in both the refined and unrefined call sets. Among these 14,001 variants, we measured the average sequence similarity between the CCS call and the ONT call, with the similarity of two strings S and T measured as [1 - edit_distance(S, T)] / max[length(S), length(T)]. Using the unrefined calls, the average similarity was 0.854, while the refined calls gave an average similarity of 0.94, demonstrating the ability of Iris to obtain more accurate insertion breakpoints and sequences. Panel G in Figure S1.1a shows the distribution of sequence similarities before and after refinement.

Iris is available as a stand-alone method at the following link: https://github.com/mkirsche/Iris

##### S1.1.4. Assigning Parental Origin by Imprinted Genes

The list of known human imprinted genes was downloaded from the Imprinted Gene Database (geneimprint.com). For known imprinted genes that showed allele-specific expression (ASE) in tissues from each individual, the haplotype-specific read counts were combined from these tissues and the potential parental origin of the haplotype blocks was determined based on the direction of the imbalance (haplotype 1 or haplotype 2) and the known expressed allele of the imprinted gene (maternal or paternal allele) (Figure S1.1c).

The parental origin results of individual 3 are shown in Figure 1B and are available in the file phased_block_ind3.txt within File: phased_block.tar.gz, where each line is a phased block. The first three columns are genomic coordinates of the phased block. The fourth and fifth columns are the parental origins of haplotype 1 and haplotype 2, respectively. 'NoInfo' indicates that there are no imprinted genes in that phased block. 'Contradict' indicates that there is at least one AS gene-imprinted gene pair that has a different imbalance direction compared to the other AS gene-imprinted gene pairs, and thus contradictory conclusions are reached for the same phased block. A similar approach can be used for the other EN-TEx individuals (See Figure S1.1b and File: phased_block.tar.gz).

#### S1.2. Functional Genomics Data Stack

The functional genomics data processing pipelines used in the EN-TEx project are described below. In total, EN-TEx includes more than 25 different biochemical assays performed on multiple (30+) tissues from four individuals (Figure 1A and Figure S1.2a). The tissues and legend of Figure 1A are detailed in panel B of Figure S1.2a.

##### S1.2.1. RNA Sequencing

Multiple RNA-seq experiments were performed in ENCODE Phase III on the 30+ tissue samples sourced from GTEx and included in EN-TEx, including: 1) long RNA-seq, i.e., RNA with a length greater than 200 nt, and total RNA-seq, 2) small RNA-seq, i.e., RNA with a length less than 200 nt, and 3) microRNA-seq, i.e., RNA with a length less than 30 nt. More information about each RNA-seq protocol and data processing pipeline can be found at the ENCODE Portal:
1) https://www.encodeproject.org/data-standards/rna-seq/long-rnas/, 2) https://www.encodeproject.org/data-standards/rna-seq/small-rnas/ and 3) https://www.encodeproject.org/microrna/microrna-seq/. RNA-seq data quality was calculated using the number of aligned reads and replicate concordance (as described in the ENCODE pipelines linked above).

##### S1.2.2. RAMPAGE

RNA annotation and mapping of promoters for analysis of gene expression (RAMPAGE) is a biochemical assay that captures 5′-complete cDNA to identify and quantify transcriptional start sites (TSSs), and characterize transcripts. The assay is described in detail at the ENCODE Portal: https://www.encodeproject.org/data-standards/rampage/. The ENCODE RAMPAGE data processing pipeline was developed for RAMPAGE libraries containing cDNA sequences longer than 200 nt. The pipeline takes cDNA sequences as input (in FASTQ format) and outputs alignments normalized for both positive and negative strands of the genome. Reproducible peaks between replicates were identified using the irreproducible discovery rate (IDR). The quality of the RAMPAGE data was determined using read depth and replicate concordance with respect to peaks in the data.

##### S1.2.3. eCLIP

Enhanced crosslinking and immunoprecipitation (eCLIP) is a biochemical assay that identifies RNA-binding protein (RBP) occupancy sites across the transcriptome. The eCLIP experimental protocol is available at the ENCODE Portal:
https://www.encodeproject.org/documents/842f7424-5396-424a-a1a3-

3f18707c3222/@@download/attachment/eCLIP_SOP_v1.P_110915.pdf. Additional assay details are available at https://www.encodeproject.org/eclip/. All eCLIP antibodies were required to undergo primary and secondary characterizations. RBP antibody standards are available at the ENCODE Portal: https://www.encodeproject.org/documents/fb70e2e7-8a2d-425b-b2a0-9c39fa296816/@@download/attachment/ENCODE_Approved_Nov_2016_RBP_Antibody_Characterization_Guidelines.pdf. The quality of the eCLIP data was determined using the number of unique RNA fragments, IDR, and the fraction of reads in peaks (FRiP).

S1.2.4. Histone ChIP-seq

Histone ChIP-seq is a biochemical assay that observes interactions between histone proteins and DNA. This assay selects for a specific histone protein variant or post-translational modification using immunoprecipitation followed by DNA sequencing. The histone ChIP-seq experimental protocol is available at the ENCODE Portal: https://www.encodeproject.org/documents/be2a0f12-af38-430c8f2d-57953baab5f5/@@download/attachment/Epigenomics_Alternative_Mag_Bead_ChIP_Protocol_v1.1_exp.pdf. Additional assay details are available at https://www.encodeproject.org/chip-seq/histone/. All commercial histone antibodies were validated by at least two independent experiments. Histone mark antibody standards are available at the ENCODE Portal: https://www.encodeproject.org/documents/4bb40778-387a-47c4-ab24-cebe64ead5ae/@@download/attachment/ENCODE_Approved_Oct_2016_Histone_and_Chromatin_associated_Proteins_Antibody_Characterization_Guidelines.pdf. The quality of the histone ChIP-seq data was determined using read depth, the number of uniquely mapping reads over the total number of reads (i.e., non-redundant fraction, NRF) and two PCR bottlenecking coefficients (PBC1 and PBC2).

S1.2.5. Transcription Factors ChIP-seq

ChIP-seq captures DNA and DNA-binding protein (e.g., CTCF, EP300, and Pol II) interactions through immunoprecipitation, pulldown, and DNA sequencing. All ChIP-seq protocols involved in the generation of data included in EN-TEx are available at the ENCODE Portal: 1) https://www.encodeproject.org/documents/20ebf60b-4009-4a57-a540-8fd93407eccc/@@download/attachment/Epigenomics_CR_ChIP_Protocol_v1.0.pdf, 2) https://www.encodeproject.org/documents/6ecd8240-a351-479b-9de6-f09ca3702ac3/@@download/attachment/ChIP-seq_Protocol_v011014.pdf, 3) https://www.encodeproject.org/documents/a59e54bc-ec64-4401-8cf6-b60161e1eae9/@@download/attachment/EN-TEx%20ChIP-seq%20Protocol%20-%20Myers%20Lab.pdf, and 4) https://www.encodeproject.org/documents/f2aa60f2-90a6-4e4b-863a-c6831be371a2/@@download/attachment/ChIP-Seq%20Biorupter%20Pico%20TruSeq%20protocol%20for%20Syapse-c5bdc444fe0511e69d6a06346f39f379.pdf. Additional ChIP-seq protocol details are available here: https://www.encodeproject.org/chip-seq/transcription_factor/. The quality of the ChIP-seq data was determined using read depth, NRF, two PCR bottlenecking coefficients (PBC1 and PBC2), replicate concordance (i.e., IDR), and FRiP.

S1.2.6. ATAC-seq

ATAC-seq identifies accessible regions of DNA by inserting primers into open chromatin regions via transposase, followed by DNA sequencing. The ATAC-seq experimental protocol is available at the ENCODE Portal: https://www.encodeproject.org/documents/404ab3a6-4766-45ca-af80-878a344f07b6/@@download/attachment/ATAC-Seq%20protocol.pdf. Additional details about the ATAC-seq protocol can be found at https://www.encodeproject.org/atac-seq/. The quality of the ATAC-seq data was determined using the number of non-duplicate, non-mitochondrial aligned reads, IDR, NRF, two PCR bottlenecking coefficients (PBC1 and PBC2), the number of resulting peaks in the data, DNA fragment length distribution, FRiP, and TSS enrichment.

S1.2.7. DNase-seq

DNase-seq is a biochemical method that identifies open regions of chromatin. These regions are identified by performing enzyme digests using endonuclease DNase I, which inserts itself into open regions, followed by DNA sequencing. DNase-seq experimental protocols are available at the ENCODE Portal: https://www.encodeproject.org/documents/c6ceebb6-9a7a-4277-b7be-4a3c1ce1cfc6/@@download/attachment/08112010_nuclei_isolation_human__tissue_V6_3.pdf. Additional protocol information can be found here: https://www.encodeproject.org/data-standards/dnase-seq/. The quality of the DNase-seq data was determined using the number of uniquely mapped reads, fraction of mitochondrial reads, and signal portion of tags score.

S1.2.8. WGBS

Whole-genome bisulfite sequencing (WGBS) was used to identify DNA methylation. WGBS converts unmethylated cytosine (C) into uracil (U), leaving methylated C unchanged. DNA sequencing followed by read alignment to a genome results in CpG island, CHG, and CHH methylation levels being observed. The WGBS experimental protocol is available at the ENCODE Portal: https://www.encodeproject.org/documents/9d9cbba0-5ebe-482b-9fa3-d93a968a7045/@@download/attachment/WGBS_V4_protocol.pdf. Additional WGBS assay details are available at https://www.encodeproject.org/data-standards/wgbs/. The quality of the WGBS data was determined from genomic read coverage, C-to-T conversion rate, and correlation of CpG methylation levels between replicates.

S1.2.9. DNAme Array

DNA methylation profiling by array assay (DNAme) measures CpG island methylation. Similar to WGBS, DNA is treated with bisulfite converting unmethylated C to U. After library amplification and purification, DNA fragments are hybridized to a microarray (Illumina Infinium Methylation

EPIC BeadChip) that probes for both methylated and unmethylated states. DNA methylation is then quantified by comparing the signal between the two DNA microarray probes. Illumina Genomestudio (v2011.1) was used to calculate the fraction of methylated reads at each CpG site from the raw microarray output.

S1.2.10. Hi-C

High-quality Hi-C data were generated from the four EN-TEx donors using samples collected from the gastrocnemius medialis and transverse colon tissues. The *in-situ* Hi-C protocol used to produce Hi-C libraries was described previously by Rao et al. (2014) (Rao et al., 2014). A detailed protocol document is provided with each dataset at the ENCODE Portal website: https://www.encodeproject.org/documents/e1ef20c9-7539-40bc-bdbf-a4deab7f72c7/. Approximately 20 mg of tissue was used for each Hi-C experiment, and the MboI restriction enzyme was used for restriction digests. All sequencing was performed on an Illumina 4,000 platform. The data was processed twice, separately utilizing a reference genome or personal genomes constructed for each individual's tissue (see S1.1).

**Hi-C Data Processing Details**

(a) Hi-C interaction matrices: Interaction matrices were generated using the Juicer pipeline (Durand et al., 2016), an open-source tool for analyzing large Hi-C libraries. We utilized BWA-MEM (Li and Durbin, 2010) to align individual reads to the hg38 reference genome, which was obtained from the ENCODE data portal. For each paired-end read, the two individual sequences were first separately aligned to the reference genome before being paired based on their read names. Chimeric reads and PCR duplicates were removed prior to the creation of an interaction matrix for each tissue of each individual (Figure S1.2b). Figure S1.2c provides information on the number of reads and number of contacts per sample utilized to create the matrices.

(b) A/B compartments: Determination of the A and B compartments was done using the Juicer pipeline (Durand *et al.*, 2016) at a 1 MB resolution. In detail, the observed/expected interaction matrices were normalized using the Knight-Ruiz matrix-balancing algorithm (KR) (Knight and Ruiz, 2012). A correlation matrix from these interaction matrices was calculated, with the first eigenvector of the matrix corresponding to A/B compartments. The positive values of the vector indicate genomic regions belonging to the A compartment, while negative values correspond to the B compartment (Figure S1.2d-e).

(c) Significant Hi-C interactions: Significant intrachromosomal Hi-C interactions were identified with FitHiC2 (v2.0.7) (Ay et al., 2014; Kaul et al., 2020). Preprocessing of EN-TEx Hi-C interaction matrices followed the author's instructions in the FitHiC2 GitHub repository's README (https://github.com/ay-lab/fithic). Matrices were binned at a resolution of 50 kb and bin biases were generated using the author's provided software (HiCKRy.py — with percentOfSparseToRemove set to 0.1). FitHiC2 output for each sample can be found in the compressed folder File: fithic2_out.tar.gz on the EN-TEx resource website. See Figure S1.2f for the number of all vs. significant interactions for each sample.

(d) TAD annotations: Topologically associating domains (TADs) were identified using TopDom (v0.9.0) (Shin et al., 2016) with a window_size parameter of 3. Before running TopDom, EN-TEx Hi-C libraries were binned at a resolution of 100 kb and normalized using the Knight-Ruiz matrix-balancing algorithm (Knight and Ruiz, 2012) implemented by Juicer (Durand *et al.*, 2016). TopDom's window_size parameter was optimized for the known enrichment of CTCF motif directionality at TAD boundaries (Rao *et al.*, 2014) and visual consistency/fit with Hi-C interaction matrices. CTCF directionality was identified using paired EN-TEx ChIP-seq peak (narrowPeak format) files and the 'CTCF_known1' motif as described by Cameron et al. (2020) (Cameron et al., 2020). TAD calls across EN-TEx individuals for the same tissue remained consistent (based on the frequency of TADs for a given size) and showed slight differences between the two tissues observed by Hi-C (Figure S1.2g), supporting their cell-type-specific nature. TAD boundary similarity was calculated by overlapping TAD annotations between individuals/tissues with a buffer of three bins when considering two boundaries to be the same.

TAD annotations were generated by applying TopDom (Shin *et al.*, 2016) to the Hi-C data from the two tissues of all four donors. Annotations are available for download via the ENTEx.encodeproject.org portal. These annotations are found within a compressed folder named File: TopDomTADcalls.tar.gz.

##### S1.2.11. Proteomics

**Liquid Chromatography with Tandem Mass Spectrometry (LC-MS/MS) Analysis**
For 10 mg tissue, 200 µl lysis buffer [50 mM Tris-HCl pH8.5, 50 mM NaCl, 8 M urea, 4% SDS, and Halt protease inhibitor (Thermo)] was added. After the tissue was homogenized by a pestle/mortar, a Dounce homogenizer, or similar device, the sample was heated at 95°C for 10 min, followed by probe sonication until the viscosity was reduced. The sample was then centrifuged at 13,000 rpm for 15 min, and supernatant was collected. RNA was first extracted from samples (see RNA-seq method).

Protein concentration was measured by the Pierce 660 nm Protein Assay (Thermo). For each sample, 100 µg proteins were taken, and volumes were equalized by 100 mM TEAB to 100 µl, reduced by 20 mM TCEP (Sigma), and then alkylated by 40 mM iodoacetamide (Sigma). Proteins were purified by 20% TCA precipitation. A total of 100 mM TEAB was added to the sample, followed by digestion with trypsin (MS grade, Thermo) at 37°C for 18 hours. The peptides were labeled by TMT10plex as per the manufacturer's instruction, and the labeled samples were pooled and SpeedVac dried. Samples of 300 µg peptides were fractionated on a U3000 HPLC system (Thermo Fisher) using an XBridge BEH C18 column (2.1 mm id x 15 cm, 130 Å, 3.5 µm, Waters) at pH 10, at 200 µl/min on a 30 min linear gradient from 5-35% acetonitrile/$NH_4OH$. The fractions were collected every 30 sec into a 96-well plate, and these were concatenated to 35 fractions and dried.

The peptides were resuspended in 0.5% formic acid (FA) and 50% was injected for LC-MS/MS analysis on an Orbitrap Fusion Tribrid mass spectrometer coupled with a U3000 RSLCnano UHPLC system (Thermo Fisher). The peptides were loaded onto a PepMap C18 trap (100 µm i.d. x 20 mm, 100 Å, 5 µm) for 10 min at 10 µl/min with 0.1% FA/$H_2O$, and then separated on a PepMap C18 column (75 µm i.d. x 500 mm, 100 Å, 2 µm) at 300 nl/min and a linear gradient of 4-33.6% ACN/0.1% FA in 90 min/cycle at 120 min, or 4-32% ACN/0.1% FA in 150 min or 180 min with a cycle time of 180 min or 210 min for each fraction. For data acquisition, we used the SPS10-MS3 method with the top speed set at 3 s per cycle time. The full MS scans (m/z 380-1,500) were acquired at a 120,000 resolution at m/z 200, and the automatic gain control (AGC) was set at 400,000 with a 50 ms maximum injection time. The most abundant multiply-charged ions (z = 2-6, above 5,000 counts) were subjected to MS/MS fragmentation by collision-induced dissociation (35% CE) and detected in an ion trap for peptide identification. The isolation window by quadrupole was set at m/z 1.0, and the AGC at 10,000 with a 35 ms maximum injection time. The dynamic exclusion window was set at ±7 ppm with a duration of 60 seconds. Following each MS2, the 10-notch MS3 was performed on the top 10 most abundant fragments isolated by synchronous precursor selection. The precursors were fragmented by higher-energy collisional dissociation at 60% CE, and then detected in the Orbitrap at m/z 110-400 at a 50,000 resolution for peptide quantification. The AGC was set at 50,000 with a maximum injection time of 86 ms.

**Personal Proteome Database**

GENCODE v27 (Frankish et al., 2019) annotation was lifted over from GRCh38 to each EN-TEx donor's personal genome, to generate eight sets of general feature format (GFF) annotations. The GFFRead utility (Pertea and Pertea, 2020) was used to extract the nucleic acid sequence for all protein-coding transcripts. An in-house Python script was then applied to translate each protein-coding transcript into its amino acid sequence. All protein sequences from the eight genomes were combined with the GENCODE v27 reference, redundant sequences were removed, and each unique protein sequence was given a unique accession ID that included the genomes that contain the protein. The final database contained 128,063 unique protein sequences, 82,136 (64%) from the GENCODE reference and 45,927 (36%) unique to the EN-TEx donors. A total of 6,344 protein sequences from GENCODE (8% of the reference proteome) were not matched to any of the alleles in the four individuals. File: Supp_data_proteomics.xlsx on the EN-TEx resource website provides cross-mapping of protein accessions to ENSEMBL transcript and gene IDs. Decoy protein sequences were generated using the DecoyPYrat tool (Wright and Choudhary, 2016).

**MS Identification and Quantification**

Spectra were processed using ProteomeDiscoverer v2.4 (Thermo Scientific) and searched against the personal proteome database using both Mascot v2.4 (Matrix Science) and SequestHT with target-decoy scoring evaluated using Percolator (Spivak et al., 2009). The precursor tolerance was set at 30 ppm and the fragment tolerance was set at 0.5 Da; spectra were matched with fully tryptic peptides with a maximum of two missed cleavages. Fixed modifications included carbamidomethyl [C] and TMT6plex [N-Term]. Variable modifications included TMT6plex [K], oxidation [M], carbamyl [K], methyl [DE], deamidation [NQ], and acetyl [N-term]. The carbamyl and methyl modifications were included due to their high incidence after samples were exposed to high concentrations of urea during the RNA extraction process. Peptide results were initially filtered to a 1% false discovery rate (FDR; 0.01 q-value). The reporter ion quantifier node included a TMT-11-plex quantification method with an integration window tolerance of 15 ppm and integration method based on the most confident centroid peak at the MS3 level. Protein quantification was performed using unique peptides only, with protein groups considered for peptide uniqueness. Peptides were quantified and normalized using tandem mass tags (TMTs) for isobaric labeling. Peptide results from ProteomeDiscoverer were remapped to the protein database and marked as reference, genome, or allele-specific (see File: Supp_data_proteomics.xlsx). Gene-level quantification of proteins was conducted by summing normalized unambiguous peptide TMT intensities.

**Peptide and Gene Identification and Quantification Results**

At a 1% FDR we report 256,512 peptide-to-spectrum matches and 117,934 distinct peptide sequences (0.01 q-value at the peptide level), of which 45,276 were quantified using TMT isobaric labels. Personal peptides were further filtered to unambiguously match one gene and have a posterior error probability below 0.699. These peptides did not map to the reference genome, only matching personal protein sequences. The 4,489 peptides identified were not present in all eight genomes across the four donors, 830 of these peptides were missing in one or more of the donors completely, and 4,334 were only present on a single allele in at least one of the donors. This corresponds to 13% coverage of the possible observable personal peptides across all protein-coding genes in the personal genomes, and a 1% increase in the number of significant distinct peptide sequences quantified (Figure S1.2h).

Gene quantification was conducted using only unambiguous peptides summing the peptide isobaric tag intensities. A total of 9,242 genes were quantified, 540 genes had non-reference peptides, 1,333 genes had peptides not present in all eight genomes (personal peptides), 518 genes had peptides absent in at least one donor, and 1,260 genes had peptides specific to a single allele in at least one donor.

**RNA-seq Comparison**

For comparison between proteomic and RNA-seq abundances, a paired set of samples and confidently identified genes matching between the proteomic and RNA-seq datasets were extracted. For each dataset, the values were normalized and then scaled to the maximum value across the samples/tissues. A Pearson correlation was then used to test the similarity between the two sets across the samples.

**Novel Gene Discovery**

All spectra were also processed via the ICR GENCODE OpenMS novel peptide discovery proteomics pipeline (Weisser et al., 2016) against a database containing GENCODE v27 reference proteins and a set of potential novel protein-coding sequences, including many unannotated PhyloCSF conserved regions (Mudge et al., 2019). Novel peptide results were filtered according to high-stringency criteria (Wright et al., 2016). This resulted in 291 novel peptides, which were further filtered to remove peptides that could be explained by semi-tryptic cleavage or single amino acid variants. The 27 remaining peptides were assessed, validating eight novel protein models, which have all now been annotated in the GENCODE reference set (Figure S1.2i).

All spectra, results, and supporting files, including the personal proteome database, have been deposited in the PRIDE (Perez-Riverol et al., 2019) proteomic repository (https://www.ebi.ac.uk/pride/) under project accession number PXD022787.

#### S1.3. Mapping Functional Genomics Data to the Personal Genomes

Typically, the analysis of next-generation sequencing data makes use of a current version of the reference human genome; this includes SNV detection, DNA methylation, transcriptional analysis (RNA-seq), identification of TF binding sites and histone modifications (ChIP-seq), and analysis of three-dimensional (3D) chromosomal looping interactions (Hi-C). Depending on the application, this process usually includes selection of either more lenient or more stringent mapping criteria, for example allowing reads to map to single or multiple genomic regions with varying numbers of permitted mismatches (Huang et al., 2014; Munger et al., 2014). Using the personal genome to map functional genomics reads becomes particularly important if the mapping requires more stringent criteria, which is necessary when the goal is precision and individualization.

We used DNA from transverse colon tissues to construct both haplotype sequences for each individual. Mapping sequences to the derived haplotypes, rather than to the reference genome, resulted in an overall improvement in mapping accuracy across different assays (RNA-seq, DNA-seq, Hi-C, and ChIP-seq). By applying conventional mapping criteria, we observed an increase in the number of mapped reads of about 0.5-1%. When we applied more stringent filtering criteria to select for high-quality, uniquely mapping sequences, we observed a much larger improvement, reaching an increase of 2-4% across assays over the four individuals (Figure 1C). We also generated a list of genes differentially expressed between mapping to the personal vs. reference genome (see File: Supp_DE_genes.tsv). Figure S1.3a-c summarizes the numbers of reads and percentages for precision mapping across the four individuals for DNA-seq, CHIP-seq, Hi-C, and RNA-seq. Mapping categories include mapping to haplotype 1 (Hap.1), haplotype 2 (Hap.2), union of Hap.1 and Hap.2 (Hap1&Hap2), reference (ref), intersection between Hap.1 and Hap.2 but not ref (Hap1&&Hap2¬Ref), and improvement as a measure of Gain=[(Hap.1 ∪ Hap.2)-Ref]/Ref.

For all assays, we excluded counting reads that mapped to the X, Y, and M chromosomes for all individuals. In general, to ensure high-quality mapping we selected reads with at most two mismatches and unique mapping. We used raw reads from transverse colon, publicly available at the ENCODE portal, with the exception of DNA-seq. For DNA-seq mapping, we used reads from blood samples that we obtained from GTEx to avoid any bias deriving from the construction of haplotypes using DNA sequences. For DNA-seq and RNA-seq mapping we used paired-end reads. For RNA-seq, to account for gene splicing, we used *.gtf files with transcript genomic coordinations and STAR Aligner v2.7. For DNA-seq, Hi-C, and ChIP-seq we used BWA v0.7.17 and selected reads with at most two mismatches and quality Q>30. For RNA-seq we used sequences with quality mapping Q=255.

#### S1.4. Variation Analysis of cCRE Activity

##### S1.4.1. Visualizing the Variation of cCRE Activity with JIVE

To visualize the relationship among the functional genomic data across the tissues, we used a dimension-reduction approach, namely Joint and Individual Variance Explained (JIVE) (Hellton and Thoresen, 2016). For each functional genomic experiment of histone modifications, we calculated its signals at the cCREs using the UCSC Genome Browser bigWig tools (Kent et al., 2010). For proteomics and RNA-seq experiments, we simply used the normalized protein abundance and RNA abundance of each gene from the experiments. For each type of assay, we generated a data matrix in which the columns are the tissues from the four individuals and the rows are cCREs or genes, and each element is the signal of the functional genomic activity measured by the assay. For each assay type, we quantile-normalized the signals. For the joint analysis of the different experimental assays, we combined these matrices by column to form a meta-matrix. In each separate data matrix, some columns in each(Ay *et al.*, 2014, 2014) data matrix are not shared by all the assays, and thus these columns are excluded from the meta-matrix.

To reduce computation burdens, we removed the rows that have low standard deviation. From this informative meta-matrix, we applied the JIVE algorithm to project the columns into a two-dimensional (2D) space (Figure 2B). As expected, this projection used all the information of the matrix. In addition, from the matrix of each assay, the JIVE algorithm excluded the information that can be explained by the other matrices, and then projected the matrix containing the information unique to the assay into a 2D space (Figure 2B). For example, in the 2D space of RNA-seq, the same tissues from different individuals are well clustered, and the different tissues are well separated. This tendency is weaker for other assays. Taken together, this observation indicates that RNA-seq likely captures the most unique signatures of different tissues.

##### S1.4.2. Using a Regression-based Approach to Quantify Activity Variation

With a linear regression approach, we used the explained variation of the regression to measure the similarity between two experiments. A larger explained variation of the regression indicated a higher similarity between the two experiments. To elaborate on the variation, we use a concrete example: the H3K27ac signals of cCREs from the spleens of two individuals. In this example, each of these individuals had two technical replicates of the H3K27ac signals measured by ChIP-seq. In each replicate, the signal at a cCRE was the fold change of reads between the IP experiment and the control experiment. For each cCRE, we first calculated the percentage difference of the signals between the two replicates. We focused on the cCREs with differences smaller than a certain cutoff so that the signals of these selected cCREs in one replicate can be largely explained by their counterparts in the other replicate using linear regression (i.e., $R^2 > 0.95$). To compare the two individuals, we used the common set of the selected cCREs with low technical noise. For each of the two individuals, we averaged the signals of the two replicates for the common cCREs. Therefore, we generated two sets of cCREs with H3K27ac signals having little noise, respectively, for the two individuals. Again, using a simple linear regression, we calculated the variance in one of the sets explained by the other. A high value indicates that the two sets of H3K27ac signals are very similar in terms of a linear relationship. As an example, the explained variation between replicates and the explained variation between experiments for different types of histone modifications in spleen is demonstrated in Figure S1.4a.

The aforementioned calculation was used for all the available histone modifications and samples (examples shown in Figure S1.4b) as well as normalized protein and RNA abundances (Figure S1.4c). For each modification, we estimated the variance explained between individuals (i.e., the same tissues of different individuals) and between tissues (i.e., different tissues of the same individual). In addition, we estimated the variance explained between two different histone modifications (i.e., the same tissue of an individual). For mass spectrometry (MS), to make the protein abundances of different genes comparable across different tissues, we normalized the protein abundances of each gene across tissues so that the highest and lowest protein abundances are one and zero, respectively. The MS approach we used pooled and labeled multiple samples together to determine protein abundances in a batch, resulting in little technical noise across the samples. To be comparable, we also normalized the RNA-seq data of the samples in the same way.

In general, histone modifications showed high similarity between the same tissue of two individuals; as expected, this number was smaller when comparing different tissues of the same individuals (Figure S1.4b). The similarity between different types of functional genomic activities from the same tissue was extremely low (Figure S1.4b). For example, H3K27ac between individuals was very similar in spleen and in transverse colon. However, the H2K27ac similarity between the two tissues was substantially reduced (Figure S1.4b). In line with this disparity across tissues, the similarity between normalized gene expression and protein abundance also varied substantially across tissues. The lower similarity in prostate is consistent with previous observations (Kosti et al., 2016). The similarities between all the available histone modifications are reported in File: Similarity_of_functional_genomic_activities_of_cCREs.xlsx. The normalized proteomics and RNA-seq data of genes are in File: normalized_proteomics_RNA-seq.dat.

Prior to the development of the EN-TEx resource, the similarities among assays were usually calculated from unmatched data. For example, a large number of histone modification signals were detected from many different human individuals in Roadmap (Roadmap Epigenomics et al., 2015). The intrinsic difference between two individuals due to genetic and environmental factors is expected to bias the similarity of two histone modifications. For the histone modifications that are positively correlated to each other, their similarity is expected to be underestimated, whereas for the negatively correlated ones, the similarity may be overestimated. The degree of such bias due to unmatched data has not been investigated for the many types of functional genomic data generated from numerous human samples. With the EN-TEx data, we can finally estimate such bias quantitatively and reliably. For example, we measured both the H3K27ac and H3K4me3 signals from the spleens of two individuals, individual 1 and 2, and the average similarity between the two signals from the same individuals was 80% in terms of variance explained, but comparing the two signals from different individuals, the number decreased to 70%. We used this approach for all the EN-TEx histone modifications, and thus estimated the influence of unmatched data on the similarity between different types of assays (Figure S1.4d). The difference varies with the explained variance. For the two signals with high similarity (large explained variance), using unmatched data results in about 10% smaller explained variance than using matched data. As expected, this trend was the opposite for two signals with low similarity. In addition, we applied this approach to measure the influence on the similarity between different tissues (Figure S1.4d).

### S2. Supp. Content for Main Text Section "Large-scale Determination of AS SNVs & Construction of the AS Catalog"

#### S2.1. Calling AS events

##### S2.1.1. Allele-specific Expression (ASE), Binding (ASB), and Chromatin Accessibility (ASCA)

**Overview**

AS phenomena arise from differential activity and/or modifications between the two haplotypes of the same individual, which can lead to allele-specific RNA and protein expression in a specific tissue or cell type. Most frequently, measurements of AS activity are performed using RNA-seq, ChIP-seq, ATAC-seq, or DNase-seq data (Baran et al., 2015; Castel et al., 2020; Chen et al., 2016; Maurano et al., 2015; Onuchic et al., 2018; Pirinen et al., 2015). The EN-TEx project simultaneously and uniformly characterizes almost all the functional genomics activity (i.e., expression, binding, methylation, etc.) at every heterozygous locus of each individual.

ASE and ASB were measured with an extended version of the AlleleSeq pipeline, now dubbed AlleleSeq2 (see website for github with c(Rozowsky *et al.*, 2011, 2011)ode). AlleleSeq (Rozowsky *et al.*, 2011) was originally developed for the 2012 ENCODE rollout (Gerstein et al., 2012; Rozowsky *et al.*, 2011). It was subsequently refined for a number of applications, including the 1,000 Genomes functional interpretation group and other projects (Chen *et al.*, 2016; Khurana et al., 2013; Onuchic *et al.*, 2018). Broadly, the pipeline incorporates personal variation, including large SVs, in order to account for reference bias (Chen *et al.*, 2016; Degner et al., 2009; Rozowsky *et al.*, 2011) in a straightforward way. We have included additional filters to mitigate ambiguous mapping biases (Chen *et al.*, 2016; van de Geijn et al., 2015). In order to account for the overdispersed nature of the functional genomics readcount data, the significance of the allelic imbalance is assessed with the beta-binomial test (Chen *et al.*, 2016) (Figure S2.1a).

**Mapping**

For each available replicate of the EN-TEx experiments, functional genomics reads were mapped to both personal haplotypes simultaneously using STAR-2.6.0c (Dobin et al., 2013). We required stringent mapping criteria, allowing the maximum number of mismatches to be 3% of the read length. For ChIP-seq and ATAC-seq datasets, mapping was performed forbidding spliced alignments. Adapters were also removed from the ATAC-seq reads with cutadapt (Martin, 2011). For RNA-seq data, we used GENCODE v24 (Frankish *et al.*, 2019) annotation converted to personal coordinates. RNA-seq mapping was performed in the two-pass mode to identify and incorporate novel junctions. Read duplicates were identified and removed from all alignments using picard (http://broadinstitute.github.io/picard/). The fraction of assay reads that were preferentially aligned to either haplotype and overlapped heterozygous SNVs (hetSNVs) across all samples ranged from 1.1-7.3%.

To visualize functional genomics reads on individual haplotypes (Figure 3C), we used SAMtools (1.9) (Danecek et al., 2021) to extract haplotype-specific reads from the BAM files generated by STAR from the last step. If an assay had multiple replicates, we merged all the BAM files. The number of reads mapped to a given region in the personal genome was calculated by bedtools (2.29.2) (Quinlan and Hall, 2010) and stored in bedgraphs, lifted over to the reference genome with UCSC liftOver (Kuhn et al., 2013), and converted to bigwigs with bedgraphToBigWig (2.8) (Kent *et al.*, 2010). Figure S2.1b summarizes the pipeline used to generate the haplotype-specific bigwigs. The bigwigs are displayed with the Integrative Genomics Viewer (IGV) (Robinson et al., 2011). See Figure S2.1c for accession numbers of the data used to generate the signal tracks in Figure 3-4. A script that generates the haplotype-specific read coverage from the BAM files is provided in https://github.com/gersteinlab/AlleleSeq2. See File: sample_signal_track.tar.gz and File: AlleleSeq2_workflow_examples.tar.gz for an example of the intermediate files (except for BAM files) in generating haplotype-specific signal tracks.

**Filtering and Assessment of Read Imbalance at Heterozygous SNVs (hetSNVs)**

The number of reads overlapping each hetSNV and carrying the corresponding alleles was calculated after filtering. The filtering included:

- Potentially misphased loci;
- Reads bearing an incorrect allele;
- HetSNVs located in potential copy number variation (CNV) sites through assessment of the surrounding read depth (+/- 1 kb);
- Sites with potential ambiguous mapping (Chen *et al.*, 2016; van de Geijn *et al.*, 2015);
- Non-autosomal chromosomes; for most downstream analyses we used call sets that only include loci from autosomes.

We aggregated read counts from all replicates available for each experiment (sample). We then calculated the significance of the imbalance at each heterozygous locus as described previously (Chen *et al.*, 2016) and called ASE and ASB sites at an FDR of 10% (Figure S2.1d).

We provide the read counts and p-values for all the ASE and ASB sites that are either significantly imbalanced or accessible (SNVs that have at least the minimum number of reads needed to be statistically detectable for allele specificity). See File: hetSNVs_default_AS.tsv for the full list of accessible heterozygous SNV loci. Columns in the hetSNV files are:

| | |
|---|---|
| 1) chr | : chromosome |
| 2-3) ref_start, ref_end | : GRCh38 locus positions (0-based, half-open) |
| 4) ref_allele | : reference allele |
| 5-6) hap1_allele/hap2_allele | : haplotype 1/2 allele |
| 7) experiment_accession | : ENCODE experiment ID |
| 8) donor | : ENTEx individual |
| 9) tissue | : tissue |

10) assay : assay
11-14) cA/cC/cG/cT : number of reads with A/C/G/T
15) ref_allele_ratio : number of reads with reference / total number of reads
16) p_betabinom : p values calculated from the beta-binomial test
17) imbalance significance : '1' passes the FDR10% threshold, '0' not a significantly imbalanced site.

S2.1.2. Allele-specific Methylation (ASM)

We used WGS variant calls to determine the positions of hetSNVs and identify all homozygous CpG positions in the genome of each donor (Figure S2.1e-f). With such information, and with the fully processed tissue-specific WGBS-aligned reads, an in-house script was then used to identify positions exhibiting significant allelic differences in CpG methylation. Our script counted the number of times a methylated or unmethylated homozygous CpG occurred in the same read as each of the two possible alleles at the hetSNV position for autosomal chromosomes. If the same read overlapped multiple CpGs, they were each considered as independent observations. Reads that overlapped with indels, had a low-quality score (Phred < 20) on the SNP position, or had a base call that did not match either of the two alleles expected in that position based on the WGBS calls were discarded. Due to the nature of bisulfite sequencing data, where cytosines may be observed as thymines during bisulfite conversion, it was not possible to determine which allele the read came from in several cases. In such cases, the read was also discarded. If a low-quality score, or an unexpected base call, was observed on a CpG position for a particular read, that observation did not contribute to the final counts. The significance of the association between the allele at the hetSNV position and the methylation state of the CpGs in the 300 bp surrounding region was assessed using Fisher's exact test. The 300 bp windows surrounding the hetSNV position were chosen as the WGBS dataset was composed of paired-end 150 bp reads. The test was only performed for hetSNV positions that showed a minimum of 6 observations of either a methylated or unmethylated CpG position for both alleles, and the p-values were subsequently corrected with Benjamini-Hochberg method for FDR control. The difference in the level of methylation between alleles was also computed for each hetSNV. Finally, ASM calls were made by identifying the heterozygous SNP positions with FDR values below a specified threshold (10%), and absolute differences in methylation between alleles above a minimum threshold of 10%. See File: ENTEx.TissueStacked.phased.final.txt for ASMs that met the accessible criteria and appeared in at least one tissue. Coordinates are 0-based in hg38. Explanation of the columns are the following:
Allele1(2).(Un)Methylated (int): Number of (un)methylated CpG in the 300 bp surrounding region of Allele1(2)
Number.of.good.reads (int): Number of reads that used to count methylated and unmethylated CpGs
Is.on.heterozygous.CpG (binary): 0 indicates that the variant is not on CpG while 1 indicates that the variant is on CpG
P.Values (float): p-value of Fisher's exact test based on Allele1.Methylated, Allele1.Unmethylated, Allele2.Methylated, and Allele2.Unmethylated
FDR (float): Adjusted p-value based on Benjamini-Hochberg method

Methylation.Allele1(2) (float): Fraction of methylated CpG in the 300 bp window of allele1(2)
Methylation.Difference (float): Methylation.Allele1 - Methylation.Allele2
Phasing.Set (string): Phasing set designated by individual vcf
Tissue (string): Tissue from which the sequenced sample originated
Individual (string): ENTEx individual ID

S2.1.3. Hi-C - Allele-Specific Interactions

**Creating Haplotype-specific Interaction Matrices**

Each pair of the paired-end reads are aligned separately to both of the parental haplotypes using BWA-MEM (Li and Durbin, 2010). Sequencing reads are then paired based on their read names. Each paired-end read is then assigned to either one or both of the parental haplotypes as follows: for each paired-end read, a score is assigned to each parental haplotype based on the number of mismatches of the mapping to that haplotype. Paired-end reads are then either assigned to haplotype 1 or haplotype 2 based on their corresponding score. In brief, pairs of reads are assigned to a haplotype if they map exclusively or with a better score to that haplotype. Additionally, pairs of reads that exclusively map to one of the haplotypes are also assigned to that haplotype. After every paired-end read is assigned to a parental haplotype, chimeric reads and PCR duplicates are removed and we generate an interaction matrix for each haplotype of each tissue of each individual (see Figure S2.1g for the pipeline and Figure S2.1h for the matrices).

**AS Interactions**

For each significant interaction captured by Fit-Hi-C, we found the number of reads that map to haplotype 1 and haplotype 2 using the haplotype-specific interaction matrices. If there was a difference in the number of reads that mapped to one haplotype vs. the other, we then calculated the p-value for the significance of the allelic imbalance using a binomial test. File: hic_files.tar.gz contains two folders: “ref” and “pgenome”. The “ref” folder contains .hic files for each individual and tissue (each individual and tissue combination is a separate folder, totaling up to eight folders); these files contain information on the genome-wide interaction matrices. The information can be extracted using Juicer tools and the contact matrices can be visualized using Juicebox (see Figure S2.1h for an example). The “Pgenome” folder contains two subfolders: “hap1” and “hap2”. Each of these folders contain two .hic files for each chromosome of each individual and tissue. Chr*.hap*.hic files contain the Hi-C data for that chromosome in personal genome coordinates and Chr*.hap*2ref.hic files contain the Hi-C data for that chromosome in a reference genome coordinate (lifted over using personal genome chain files). Figure S2.1i shows the total number of raw AS interactions and significant allelic imbalances per sample (calculated using the binomial test described above).

S2.1.4. Proteomics - Allele specific Peptide (ASP) Analysis

The proteomics data were mapped at the gene level and filtered to a set containing one or more AS peptides in any donor. These fell into two categories: genes with AS peptides for one allele only or those with peptides specific to both alleles. Both groups were considered for ASP ratios.

The ASP ratios were calculated for each tissue and donor in which allelic peptides were quantified, based on the ratio of the summed peptide intensities of peptides specific to the two alleles. Individual ASPs were filtered to require a minimum of three distinct peptides unambiguously identifying a gene, an expression level for the tissue of not less than five-fold lower than the highest expressed tissue and an ASP ratio of greater than 0.75. Figure S5.2b summarizes key numbers of genes with allelic peptides. See File: Supp_data_proteomics.xlsx for a full list of allelic peptides.

#### S2.2. Allele-specific Functional Elements

##### S2.2.1. Genes and cCREs

We extended our pipeline to measure allelic imbalance at genomic regions and elements of interest. To do so, we aggregated read counts from all hetSNVs within the relevant region and assessed the significance of imbalances between personal haplotypes for individual hetSNVs as described above. We provide a large catalog of genomic elements measured for AS activity (e.g., ASE genes and cCREs) with corresponding haplotype-specific assay read counts and significance scores of the imbalance (Figure S2.2a). See File: genes_default_AS.tsv and File: cCREs_default_AS.tsv for lists of accessible genes and cCREs with allelic imbalance calculations. Columns are similar to those described in S2.1.1 with the following differences:
4) region_id : gene name (GENCODE v24) or cCRE id
5-6) hap1_count/hap2_count : number of reads mapped to haplotype 1/2

##### S2.2.2. Correlation Between AS Genes and Diseases

We compared the set of AS genes to a set of genes associated with certain diseases. The list of disease genes are those known to be affected by disease-associated mutations and expressed in disease-related tissues (Jiang et al., 2020). For every tissue and individual, we noted the genes that were present in both the set of AS genes and the set of disease genes. The list of the overlapping genes and their associated diseases can be found in File: Associated_AS_Disease_Genes.xlsx. Many of the correlations were sensible. For example, TSHR, TG, and PAX8, which are associated with hyperthyroidism, showed AS behavior in the thyroid, and TNNT2, LDB3, and SCN5A, associated with cardiomyopathy, showed AS expression in the heart.

##### S2.2.3. Gene Ontology Enrichment Analysis of AS Genes

To determine the characteristics of active AS genes, we performed gene ontology enrichment analysis of protein-coding genes that showed AS activity in different assays (Figure S2.2b). DAVID Bioinformatics Resources 6.8 (Huang da et al., 2009a; 2009b) was used to perform the functional annotation clustering. For ASB, the background list for each assay includes all protein-coding genes with accessible promoters in that assay; for ASE, the background list includes all protein-coding genes with an accessible expression level from RNA-seq. The AS gene list for each assay includes genes showing AS activity in any EN-TEx individual and tissue, and genes were ranked by p-value to be AS genes. For ASE analysis, since DAVID has a 3,000 gene limit, the top 3,000 mostly ASE+ protein-coding genes were selected for the enrichment analysis, and the top 20 enriched terms are shown. We found that protein-coding genes showing AS activity in assays are mostly enriched in phosphoprotein, and their sequences are featured with polymorphism and variants.

#### S2.3. Aggregation Across Tissues and Assays

##### S2.3.1. AS Expression and Binding

We observed a large increase in detection power when we pooled reads for each hetSNV across all tissues in each individual. We calculated the significance of the imbalance at each hetSNV for the pooled call set in the same manner as for individual tissues and called ASE and ASB sites at an FDR of 10%. Figure S2.3 provides a summary of the AS catalog, including the number of AS hetSNV and AS elements, with different aggregation methods and levels. See File: hetSNVs_pooled_AS.tsv for a list of accessible hetSNVs with allelic imbalance calculations.

##### S2.3.2. Methylation

We aggregated the counts of methylated and unmethylated homozygous CpG positions surrounding both alleles of each heterozygous SNV across tissues for each individual to assess the cross-tissue association between the allele at the hetSNV position and the methylation state of the homozygous CpGs. The significance of association was computed using Fisher's exact test; Benjamini-Hochberg method was used to control the FDR. For aggregated observation, the test was only performed for accessible hetSNV positions that showed a minimum observation of methylated or unmethylated homozygous CpG positions for both alleles. The number of positions n (around 12) was determined by maximizing the sum of p-values. ASM were called at FDR values under 10% and absolute methylation differences larger than 10%.
We then generated a combined ASM call set that includes cross-tissue counts of methylated and unmethylated homozygous CpG observations surrounding accessible hetSNVs for all four individuals. Identical hetSNVs across individuals were included as separate records of CpG counts. All accessible hetSNVs, their associated gene, distance to gene, and genomic region were annotated based on the refGene database. Alternative allele frequency was annotated based on the Genome Aggregation Database (gnomAD) 3.0 database using ANNOVAR (Wang et al., 2010). cCREs were annotated based on the ENCODE. See File: ENTEx.TissueAggregated.final.txt for cross-tissue ASMs in autosomal chromosomes. Columns are similar to those described in S2.1.2.

#### S2.4. Generalizability of the AS catalog

We discovered over one million SNVs that show AS activity in gene expression, DNA methylation, histone modification, and/or TF binding. This catalog should cover a large fraction of AS activity of common SNVs. To estimate the coverage, we started by using the 1,000 Genomes project high-coverage data (1000 Genomes Project Consortium et al., 2015) to assign allele frequencies to the EN-TEx SNVs. We found that 76% (i.e., 5,276K) of the common SNVs in 503 European individuals (specifically, individuals of GBR, FIN, IBS, TSI, and CEU) were discovered in EN-TEx, 4,414K of which were heterozygous and unambiguously genotyped in at least one of the four EN-TEx individuals. Among these 4,414K SNVs, 946K (21.4%) show AS activity in at least one assay. If the EN-TEx project was conducted on all 503 European individuals from the 1,000 Genomes project, which contains 6,946K common SNVs with AF<1, then the number of AS SNVs would be 6,946K * 21.4% = 1,486K (assuming each of the 6,946K SNVs is heterozygous in at least one individual). This number is only a 540K increase from the 946K that are currently in the AS catalog, indicating that our catalog includes a majority of AS events at common SNV loci in the European population.

While previous studies also compiled AS histone modifications and/or DNA methylation, our catalog is larger. For example, while Onuchic et al. (Onuchic *et al.*, 2018) reported 125K ASM loci, 36K loci with ASB H3K27ac, and 0.5K loci with ASB H3K27me3 (Table S1 of Onuchic et al.), our catalog includes 469K, 79K, and 96K loci, respectively. Similarly, Chen et al. (Chen *et al.*, 2016) used SNVs discovered by the 1,000 Genomes project (2,504 individuals) to construct a diploid genome for each of the 384 individuals compiled by Geuvadis. They mapped RNA-seq and ChIP-seq to these diploid genomes and identified 63K ASE hetSNVs and 6.1K ASB (ChIP-seq) hetSNVs, the latter of which is a much smaller number than the 361K in our catalog. In terms of AS activity in the regulatory regions, we also found more (28K vs. 11.7K) AS cCREs than a similar study using the Roadmap data (Leung et al., 2015). While we note that the EN-TEx resource does not have more ASE events than GTEx (Castel *et al.*, 2020) or AlleleDB (Chen *et al.*, 2016), we found that ASE is only a small fraction of all the AS events in the genome (Figure S2.4). Most AS events are related to the chromatin states of the regulatory regions.

##### S2.5. Calling AS Events in External Datasets Used for Validation (Roadmap and NA12878)

In addition to the EN-TEx AS catalog, we have also generated ASE and ASB call sets for the CEPH individual NA12878 and Roadmap (Roadmap Epigenomics *et al.*, 2015) individuals STL002 and STL003. We used this for external validation of our predictive models.

Personal genome sequences for STL002 and STL003 were constructed using variant calls generated previously (Onuchic *et al.*, 2018). For NA12878, we used SNVs and indels available from the Illumina Platinum Genomes project (Eberle et al., 2017) (2016-1.0) and large deletions generated by the 1,000 Genomes Phase 3 SV Analysis Group(Sudmant et al., 2015). All datasets with matching assays (and all tissues for STL002 and STL003) that are available from the ENCODE portal were utilized to generate the AS call sets. These personal genome files are available as File: pgenome_NA12878.tar.gz (NA12878), File: pgenome_STL002.tar.gz (STL002), and File: pgenome_STL003.tar.gz (STL003).

See Figure S2.5 for the number of AS hetSNVs detected in each sample. The STL002, STL003, and NA12878 AS hetSNV call sets are available as File: hetSNVs_default_AS_validation.tsv and the file is formatted as described in S2.1.1. These call sets were used for validation of predictive models described in S4.

#### S2.6. High-confidence and High-power Call Sets

We also developed a "high-confidence" call set requiring that at least one read from both alleles was detected in the functional genomics assay, thus accounting for potential false-positive genotype calls. See File: hetSNVs_high-confidence_AS.tsv for a list of hetSNVs with high-confidence allelic imbalance calculations.

In addition, we generated a "high-power" tissue-specific call set by allowing a more relaxed threshold (FDR 20%) for loci that were detected as significantly imbalanced after read pooling-based joint calling across all tissues (Figure S2.6a). See File: hetSNVs_high-power_AS.tsv for a list of hetSNVs with the high-power allelic imbalance calculation.

In addition, we tested two methods for increasing detection power of AS hetSNVs in datasets with low read counts. Both methods impose a less strict test for allele specificity on hetSNVs that have been determined to be AS in prior experiments. This prior knowledge can be taken from other experiments on the same individual, or from experiments on different individuals entirely. The first “high-power” method relaxes the FDR threshold from 10% to 20% for all hetSNVs that have prior evidence of allele specificity. All other hetSNVs are evaluated at the usual 10% FDR threshold. The second method uses a one-sided beta-binomial test, instead of the default two-sided test, to determine whether the direction of imbalance is consistent with prior data. With both high-power methods, new AS hetSNVs are identified that did not meet the threshold using the default calling method.

For the purpose of validating these high-power methods, we tested them on a deep RNA-seq experiment of the cell line GM12878 (Figure S2.6b). We first identified ASE hetSNVs in the dataset using our default calling method. This was our “gold standard” list of 24,685 ASE hetSNVs. Then, we simulated a shallower sequencing experiment by downsampling by a factor of 4. Using the default ASE calling method on the downsampled dataset, we identified 6,928 ASE hetSNVs. Approximately 80% of these hetSNVs were in common with the gold-standard list. We expect that the error rate is a product of the randomness of downsampling. Then, both high-power calling methods were performed on the downsampled dataset.

We generated priors for the high-power methods using the pooled reads across all tissues from the four EN-TEx individuals. If a hetSNV was ASE in at least one individual, it was included in the high-power test. If two or more individuals had ASE of the same hetSNVs, the direction of imbalance should agree (e.g., both favor the reference allele over the alternative allele), otherwise that hetSNV was excluded. We did not take into account the identity of the alternative allele for any hetSNV.

For the relaxed FDR method, 122 new ASE hetSNVs were identified that did not meet the threshold for allele specificity using the default method. For the one-sided method, 275 new

ASE hetSNVs were identified. For both, approximately 60% of the new ASE hetSNVs were in common with the “gold standard” list. The validation shows that both methods can be used to identify a modest number of new ASE hetSNVs, at the cost of somewhat reduced specificity.

It should be noted that the results of the high-power methods are dependent on the nature of the prior. Both methods can only be used to evaluate hetSNVs for which there is prior information. Using data from the EN-TEx individuals as a prior for non-EN-TEx cell lines such as GM12878 captures most common hetSNVs, but excludes most rare variants. If more of the individual’s hetSNVs are in common with the prior, it is likely that the high-power methods will identify more AS hetSNVs. In the case of the EN-TEx individuals, we circumvent this problem by using AS hetSNVs identified from the all-tissues pooled reads as the prior for high-power analysis of individual tissue datasets. Because both sets come from the same individual, they share both common and rare variants.

#### S2.7. Integration with the ClinGen Allele Registry

The variants identified in all four EN-TEx individuals are registered in the ClinGen Allele Registry (Pawliczek et al., 2018), which provides unique variant identifiers for canonical alleles defined at the level of nucleic acid sequences or at the level of proteins. The unique identifier integrates different types of labels and definitions of the same allele across multiple databases including dbSNP (Sherry et al., 2001), gnomAD (Karczewski et al., 2020), ClinVar (Landrum et al., 2018), and ExAC (Lek et al., 2016); approximately 24K variants were previously recorded in ClinVar (Ngatchou et al., 2013). 58 variants are classified as 'pathogenic' or 'likely pathogenic' and 14 of them show AS behavior (including ASM) in at least one of the EN-TEx samples. All variants are bulk registered in VCF format using API specified by Allele Registry documentation (http://reg.clinicalgenome.org/doc/AlleleRegistry_1.01.xx_api_v1.pdf). Variants can be queried either programmatically via APIs or via search interface using any type of ID associated with the variant. Metadata for allele(s) are available in machine readable form (JSON) on ClinGen and can be queried in bulk as well.

#### S2.8. Examples of Coordinated AS Activity Across Assays

We detected the inactive copy of the X chromosome in the majority of tissues from female individuals. We found that both gene expression and active chromatin signal is significantly skewed toward one haplotype, while repressive chromatin signal is significantly skewed toward the opposite haplotype in many tissues.

We then investigated the specific allelic coordination at the chromosome level using the X chromosome. X chromosome inactivation ensures that females have only one functional copy of the X chromosome, and occurs by random selection of the inactivated copy early in embryonic development. For the two female individuals, the EN-TEx data enabled comprehensive analysis of the AS activity in 24 tissues. We identified the active copy of the X chromosome by examining the overall gene expression levels in 24 tissues of individual 3. The gene expression, active histone marks, and repressive histone marks were coordinated in terms of their haplotype-specific activity. As shown in Figure 3C, the gene expression values of all genes in the X chromosome were higher in haplotype 2 than those in haplotype 1 (see Figure S2.8a and Figure S2.8b). In accordance with this finding, enrichment of the active histone mark H3K27ac was also higher in haplotype 2 than in haplotype 1. Moreover, enrichment of the repressive histone mark H3K27me3 was imbalanced in the opposite direction (i.e., higher in haplotype 2). See Figure S2.8d for a similar analysis performed on individual 4. A similar coordination was observed using other AS activities such as POL2R and CTCF binding (Figure S2.8a).

When focusing on specific genes, we found that the *DHRSX* gene located on the pseudo-autosomal region had balanced expression in both haplotypes, whereas the *SLC25A5* gene located on the inactivated region showed a significant skew in gene expression towards the active haplotype in accordance with the chromosomal-level imbalance. We also identified a known “escaper” gene, *KDM6A* (Itoh et al., 2019), that demonstrated balanced expression in both haplotypes while being located on the inactivated region of ChrX. The expressed haplotype of the AS *SLC25A5* gene in tibial nerve tissue showed significant AS activity for the activating histone mark H3K27ac, while the “inactive” haplotype had significant AS activity for the repressive histone mark H3K27me3 and for DNA methylation. In addition, our analysis of haplotype-specific Hi-C data revealed an AS skew in Hi-C interactions between another gene, *XACT,* and its potential distal regulatory element on the active haplotype of Chr X (see Figure S2.8c).

We found an example of AS activity for a less-characterized locus. We detected AS Hi-C interactions in the *XACT* locus (Figure S2.8c) on the active copy of chromosome X. We first determined the active copy of chromosome X by looking at the gene expression distribution on both haplotypes and found that haplotype 2 has more gene expression than haplotype 1. We then looked at the differential interaction of chromosome X by subtracting the Hi-C matrices of the haplotypes. We found that an interaction between the *XACT* locus and a region upstream of it is significantly elevated in the active haplotype. We also found that both the *XACT* locus and the upstream region are bound to CTCF, which might be mediating the interaction. *XACT* is a long non-coding RNA (lncRNA) found to be active in the active copy of chromosome X early in cell development. This CTCF-mediated haplotype-specific interaction could play a role in activating the *XACT* locus established at early stages of cell development. While such observations are interesting, they are provisional on additional supportive data.

#### S3. Supp. Content for Main Text Section “Interrelating SVs & Chromatin Modifications”

##### S3.1. Analysis of Structural Variants

We focused our analysis on SVs that are larger than or equal to 50 bp, while the VCF files of each individual (see S1.1.2) also contain smaller “SVs”. We noted that SVs identified from Oxford Nanopore data (individuals 1 and 4) had fewer large insertions than those identified from

Pacbio data (individuals 2 and 3) (Figure S3.1). This likely resulted from differences between the two sequencing technologies (Aganezov *et al.*, 2020; Kirsche et al., 2021).

To analyze the sequence composition of the SVs, we used RepeatMasker (ver. 4.0.7, slow search mode) (http://www.repeatmasker.org) to classify the sequences that are inserted, deleted, or inverted.

We also estimated the allele frequencies of the SVs. For this purpose, we checked for overlaps in the location between the EN-TEx SVs and those reported by Audano et al. (2019) (Audano et al., 2019). To increase the chance of finding an overlap between these two datasets, we used the confidence intervals (CIs) of an EN-TEx SV's coordinates as the location of the SV. Specifically, the CIs of the breakpoints were denoted by CI_POS and CI_END in VCFs of individuals 2 and 3. The SVs of individuals 1 and 4 were called by different tools, therefore the corresponding VCFs did not have CI_POS and CI_END. Instead, we used +/- 2*STD_quant_start as the CI of POS and +/- 2*STD_quant_stop as that of END. For DEL and INV, we extended POS upstream by its CI and END downstream by its CI. For INS, we extended the POS upstream and downstream but did not extend the END. When an overlap was found, we further checked whether the two SVs were of the same type (e.g., both are deletions). If the two SVs were not the same type, we considered the two SVs to be different. Through this analysis, we matched SVs in Audano et al. (2019) with 68.3%, 65.9%, 63.4%, and 65.3% of the SVs in the four individuals, respectively. We assigned these EN-TEx SVs an allele frequency in European populations estimated by Audano et al. (2019) (Audano *et al.*, 2019). We performed a similar analysis by using more recent SVs called from long-read DNA sequencing data (Ebert et al., 2021) and gnomAD SVs (Karczewski *et al.*, 2020). A total of 71.4%, 68.9%, 66.5%, and 68.0% of the SVs in the four individuals, respectively, overlap with the former dataset. Because gnomAD annotates SVs differently, we allowed EN-TEx "INS" to match "INS", "DUP", "BND", and "MCNV" in gnomAD, EN-TEx "DEL" to match gnomAD "DEL", "BND", and "MCNV", and EN-TEx "INV" to match gnomAD "INV" and "BND". In this way, we found a match for 63.4%, 61.4%, 60.3%, and 63.1% of the SVs in the four individuals, respectively.

To understand how SVs distribute in the genome, we generated a null expectation of SV distribution by shuffling the locations of SVs, using a method similar to that used in the 1,000 Genomes SV study (Sudmant *et al.*, 2015). Specifically, we put the SVs in random locations on the same chromosome while avoiding gaps in the assembly. We calculated the ratio of the number of unshuffled SVs intersecting a given genomic region over the number of shuffled SVs. We repeated the shuffling 1,000 times.

#### S3.2. Associating SVs with Expression Quantitative Trait Loci (eQTLs)

We aimed to identify heterozygous SVs that potentially cause AS gene expression and underlie the action of known eQTLs. To do this, we first identified eQTLs (GTEx Consortium, 2020) that are compatible with the AS expression of the associated genes. For each ASE gene, we checked if the two alleles at each associated eQTL locus had the expected regulatory effect. The numbers of heterozygous SNPs and indels that were identified as compatible eQTLs in at least one tissue were 219K, 190K, 184K, and 137K in the four individuals, respectively (Figure S3.2a). We used the compatible eQTLs associated with a given ASE gene to define a window spanning from –10 kb of the compatible eQTL on the far 5' end to +10 kb of the compatible eQTL on the far 3' end. For a heterozygous SV that intersects with this window, we determined whether the SV and the compatible eQTLs may locate on the same linkage block by comparing their allele frequency and haplotype. Specifically, for each SV identified in the last step, we identified all compatible eQTLs (with respect to the given ASE gene) that fell within +/- 10 kb of the SV. Suppose the SV is on haplotype 1, then we calculated the allele frequencies of the alleles of the compatible eQTLs on haplotype 1. Here, we used the allele frequency in the European population reported by the 1,000 Genomes project (1000 Genomes Project Consortium *et al.*, 2015) for the alleles at each compatible eQTL. For each individual, about 500 ~ 800 compatible eQTLs carried an alternative allele that could be found in the 1,000 Genomes project. These compatible eQTLs were excluded from the next steps. If at least 30% of the hap1 alleles of the compatible eQTLs within +/- 10 kb of the SV had similar allele frequencies as the SV's allele frequency (defined between 80% and 120% of the SV's allele frequency), then we considered the SV to be potentially linked to the compatible eQTLs and that it may contribute to the AS expression of the given gene. We listed SVs that meet this criteria, the associated ASE gene, and the compatible eQTLs +/- 10 kb from the SVs in File: Supp_Data_SVs_associated_with_eQTL.xlsx.

We identified known eQTL-associated SVs (including SV-eQTLs) (Chiang et al., 2017; Sudmant *et al.*, 2015) in our list of potential eQTL-associated SVs. We considered that an SV was a match if a reported eQTL-associated SV was found within +/-100 bp of this SV and both SVs were associated with the same gene. We searched for matches in tissue-specific and non-tissue-specific ways. For individual 1, our list includes 337 SVs that are associated with eQTLs in at least one tissue, of which 67 match known eQTL-associated SVs. The fractions are 84/317, 70/304, and 46/215 for individuals 2 to 4, respectively. Details of these results are listed in Supp_Data_SVs_associated_with_eQTL.xlsx. For comparison, we also calculated the fraction of known eQTL-associated SVs in our SVs that are close to genes with AS expression (Figure S3.2b). We pooled genes that have AS expression in at least one tissue. Because GTEx eQTLs fall within +/- 1 Mb of the TSS of genes (GTEx Consortium, 2020), we used the same window to look for SVs near the genes with AS expression, requiring the SVs to at least partially overlap with the windows. We further required SVs to be heterozygous, clearly phased, and relatively common (i.e., present in Audano et al. (2019) (Audano *et al.*, 2019)). We found 4,912 SVs in individual 1 that meet these criteria, of which 596 match known eQTL-associated SVs. This fraction is significantly lower than the observed fraction of 67/337 ($p = 3.4e-5$, Chi-square test). We observed similar enrichment in the other three individuals ($p = 4.2e-12$, $3.1e-4$, $4.9e-6$).

See Figure S3.2c-j for examples of indels potentially changing the gene expression and examples of splicing variants.

#### S3.3. Aggregating the Impact of SVs on Neighboring Chromatin

Our goal is to calculate potential changes in the chromatin state in the neighborhood of SVs. Intuitively, this can be done by comparing the chromatin state near heterozygous SVs between the haplotypes of an individual. We excluded heterozygous SVs that are closer than 5 kb from other SVs in the same individual.

In the remaining heterozygous SVs, we focused on those that have relatively precise breakpoints. Specifically, we kept SVs where the total length of the start position's confidence interval and the end position's confidence interval is at most 50 bp. To minimize the influence of SVs on mapping sequence reads, we further excluded SVs for which the average mappability of a window +/- 500 bp of the SV is below 0.9. Because EN-TEx requires the length of a ChIP-seq read to be at least 50 bp, we used 50-mer multi-reads Umap mappability (Karimzadeh et al., 2018) to filter SVs when calculating potential disruption to chromatin openness (measured by ATAC-seq) and histone modifications (measured by ChIP-seq). We used 100-mer multi-reads Bismap mappability (Karimzadeh *et al.*, 2018) to filter SVs when working with WGBS data. We also excluded SVs that fall in blacklist regions that are known to give problematic ChIP-seq reads (Amemiya et al., 2019). When Umap mappability was used, the numbers of SVs that passed the filters were 3,931, 3,636, 3,006, and 3,258 for the four individuals, respectively. When Bismap mappability was used, the numbers were 4,522, 4,246, 3,497, and 3,777.

For each SV that passed the above filters, we calculated the average chromatin state in the SV's flanking regions. We define flanking regions of an SV as the -500 bp ~ -100 bp region and the 100 bp ~ 500 bp region (Figure S3.3a) – the extra 100 bp upstream and downstream of the SV are extra buffer regions that should reduce the influence of SVs on mapping sequencing reads. Because the chromatin state can be tissue specific and individual specific, we treated the SV-sample combinations as independent data points. We summed the allelic ATAC-seq and ChIP-seq reads on hetSNVs (see section S2.1.1) that fall into the flanking regions of a given SV in a given sample. For ATAC-seq and each ChIP-seq assay, we excluded SV-sample combinations in which the total reads from both haplotypes were less than 15. This step left about 4.4 K ~ 7.2 K SV-sample combinations (each SV has 2.9 ~ 3.8 samples on average) for the ChIP-seq assay. If the haploid that carries the SV has 70% or less reads (e.g., ATAC-seq reads) than the other haploid, we consider that the SV reduces the given chromatin state (e.g., chromatin openness). For CpG methylation measured by WGBS, we averaged the ASM levels around hetSNVs (section S2.1.2) that fall into the flanking regions of SVs. About 58% of the SV-sample combinations lacked suitable hetSNVs in the SV flanking regions. We similarly required at least 70% reduction in the methylation levels near SVs, but we included all 45.6 K SV-sample combinations (6.2 samples per SV), since the general methylation levels of CpGs were high (in 98% of SV-sample combinations the methylation levels of the SV flanking regions averaged over the two haplotypes were above 50%). For each chromatin state, we reported the fraction of SV-sample combinations where the chromatin state was reduced near the SVs (Figure S3.3b).

We also repeated the analysis by comparing one individual that carries the SV with one who does not. In this case, we excluded SVs that are closer than 5 kb from any other SVs in individual 2 or individual 3, and SVs that fall on the sex chromosomes. We similarly filtered out

SVs that have imprecise breakpoints and/or low mappability in the neighborhoods. A total of 1,974 SVs in individual 2 and 2,154 in individual 3 passed the filters when Umap mappability was used; the numbers were 2,294 and 2,478 when Bismap mappability was used.

We calculated the average fold-change over control of ATAC-seq and histone ChIP-seq, and the average methylation levels of CpG sites, in the flanking regions of SVs. For ATAC-seq and histone ChIP-seq, we also excluded SV-assay combinations in which the sum of the fold-change over the two individuals is less than 1.0, leaving 43 ~ 93% of the SV-assay combinations to determine reduction in chromatin state. For DNA methylation, we included all SV-assay combinations. To qualify a reduction in chromatin state, the above signal in the individual who carries the given SV must be 70% or lower than in the individual who does not. The right panels of Figure S3.3b show the fraction of SV-assay combinations where a given chromatin state is reduced near the SVs. Again, TE-related SVs tended to reduce chromatin openness and H3K27ac levels in the neighboring regions.

#### S4. Supp. Content for Main Text Section “Generalized Application #1: Predicting AS ChIP-seq Activity from Nucleotide Sequence”

##### S4.1. Relation between AS SNPs and Transcription Factor (TF) Motifs

We collected 660 human TF motifs from the Cis-BP database (Weirauch et al., 2014). Specifically, we required the motifs to be from protein-binding microarray (PBM) and SELEX-based experiments. Position weight matrices (PWMs) from multiple motifs are combined into a single PWM file for each TF. We used FIMO (Grant et al., 2011) with p<10-e4 to scan the motif occurrence in the human genome. We then intersected the AS SNP file with each motif occurrence file with bedtools and retrieved the contingency table of whether a SNP was AS and whether a SNP was in a motif. An odds ratio (OR) was used as the measurement of AS enrichment and Fisher’s exact test was used for statistical significance. The motifs were then ranked based on the OR. For H3K4me3 ChIP-seq-based or any other assay-based ranking, only the SNPs that were accessible in that assay were used to intersect with the motifs. For the list of motifs as well as the OR and p-values in all assay or specific assay, see File: motif_ranking.tsv.

For the conservation score of the motifs, we downloaded the genome-wide phyloP score from UCSC Genome Browser. For each occurrence of the motif, the conservation score of the region was determined by the mean of each base (from UCSC Kent_tool bigWigAverageOverBed). The conservation of the TF was the mean of the scores from all occurrences of its motifs. The entropy of a motif was calculated by sum(-plog(p)) where p is the relative frequency of each base in each position. The fraction of CG of a motif was calculated as the number of positions where C or G was the most frequent base, divided by the length of the motif. Spearman’s correlation was used for all pairs of the rank and each motif property (Figure S4.1a).

Each motif's family information was obtained from Cis-BP as well. We noticed that the top-ranked motifs were more likely to be in the C2H2 zinc finger family. A C2H2-ZF domain typically contains 3-4 base-contacting residues, and zinc finger proteins usually contain multiple tandem C2H2-ZF domains. The individual DNA motifs of these tandem domains often overlap with each other and assemble into the full-length motifs we observed in SELEX (e.g., 4-mer and another 4-mer overlapping by one base results in a 7-mer) (Najafabadi et al., 2017). Thus, mutations in the overlapping base in the middle of the motif might be more likely to affect the binding affinity of the TF. Consistent with this reasoning, we observed that AS SNVs happened more frequently in the "conjunction" base while non-AS SNVs happened relatively randomly across all positions of the motif (see FOXO3 in Figure 5A and Figure S4.1b).

#### S4.2. Allele-Specific Effect Prediction with the BERT Model

BERT is a natural language model based on the Transformer neural network architecture. It has been widely applied to natural language processing due to its ability to incorporate long-range contextual information (Devlin et al., 2018). Thus, it can also be applied to extract meaningful sequential patterns from genomic sequences, such as to predict AS effects of SNPs.

We extracted the 128 bp sequence upstream and downstream of the SNP in question as the input. The sequences were labeled as positive or negative based on their AS effects. For balancing considerations, the negative set was randomly downsampled to the same size as the positive set. The dataset was then split into training, cross-validation, and testing sets at a 8:1:1 ratio.

We initialized the BERT model with the weights of the pre-trained DNABERT model (Ji et al., 2021). A single-layer classifier was added on top of the output of DNABERT and the model was fine-tuned on the AS datasets. For fine tuning, we selected from a range of hyperparameters (learning rate=1e-5, 5e-5; training epoch = 5, 10, 20). As the pre-trained DNABERT model has different versions with k-mer sizes of 3-6, we report the model with the highest performance.

The model was first trained with only SNPs from donor individual 3 to predict the "pooled" AS SNVs (i.e., SNVs that were active in at least one tissue). For many of the prediction tasks, the model achieved performance of AUROC > 0.7 on the validation set, significantly higher than logistic regression and random forest on sequence embeddings (Figure S4.2a-d; see below for more details). We then tested the model performance on validation sets composed of SNPs exclusive to the other three donors. The validation sets for these three individuals have been randomly downsampled to the same size as the validation set for individual 3. The sampling was repeated ten times and average results are reported. As expected, the performance was lower compared to individual 3. Specifically, the model showed exceptional performance on the prediction for CTCF (AUROC = 0.7936) and could be generalized well to the other three donors (average AUROC = 0.6876). The model for H3K27ac AS SNVs also showed high validation performance on the test set (AUROC = 0.8001), other individuals (average AUROC = 0.7286), and an external validation set from Roadmap individuals (average AUROC = 0.7426.

For model interpretation, we used the method implemented by (Ji *et al.*, 2021), where the attention scores of the last layer for the first token are averaged over all 12 attention heads, and then regularized by k-mer coverage. As a comparison, we used the dna2vec model released by (Ng, 2017) to transform k-mers to continuous-valued vectors, preserving their contextual preference. Using the same training, test, and validation data as above, we represented each input sequence as an average over the embedding of all its k-mers. We then trained a logistic regression classifier based on the average embedding vector. We performed embedding with k-mer sizes of 3-8 and reported the one with the highest performance.

We also implemented a much simpler model based on motif information only. We overlapped the hetSNVs from the same training set as above with identified CTCF motifs from the Cis-BP database. The following features were used to build a random forest classifier:

| Feature | Feature frequency |
|---|---|
| A: Whether the SNP overlaps with CTCF motif | 287/28,891 (positive)<br>45/28,891 (negative) |
| B: Whether the SNP overlaps with conserved positions in CTCF motif | 161/28,891 (positive)<br>27/28,891 (negative) |
| C: Whether there is CTCF motif within 256 bp window | 1489/28,891 (positive)<br>689/28,891 (negative) |
| D: # of TF motifs within the 256 bp window | Average 56.8 (positive)<br>Average 62.2 (negative) |

A logistic regression model using feature A and B only has almost no predictive performance (AUROC=0.504). By adding features C and D, which include some contextual information, the performance increases to AUROC=0.5618, which is still much lower than other models in comparison. This is expected because the frequency of CTCF motifs is quite low and only accounts for a very small portion of the dataset.

#### S5. Supp. Content for Main Text Section “Generalized Application #2: Models Interrelating AS Activity at Promoters and Genes”

##### S5.1. Prediction of Promoter AS Activity With a Random Forest Model

We trained a random forest model that could predict the AS binding state of gene promoters in an assay and in individual tissue-specific manner. We call this the "reverse" model as we are going from gene to promoter. Models achieved good performance on both internal EN-TEx and Roadmap STL002/3 data (Figure S5.1a). After testing different combinations of features, models were built using four features (Figure S5.1b), which are numbers of ranked TFs’ motifs intersecting, nearby and distal to the hetSNV in the promoter and the imbalance ratio of haplotypic expression. In addition to the feature importance score from the random forest model, we investigated the association of each feature with ASB promoters, indicated by an R2 score (Figure S5.1c). Other features (including gene expression level, eQTL, all 660 non-ranked TF features in the promoter) were tested but proven to not be informative (Figure S5.1d). We applied our model on a large scale to the entire GTEx cohort (>800 people) to predict AS promoters from the available genotypes and ASE data. The results of the model are in File: ASB-predictions-on-GTEx-cohort.tsv, which lists GTEx individuals, gene names, assay types, and predictions of the associated promoter.

We also constructed a "forward" model to predict ASE from ASB. Specifically, to interrelate AS activities of genes and promoters (Figure S5.1e), a random forest model was trained using assay-based annotations of the promoters. The assay-based ASB of the promoters was informative for the prediction of ASE genes (Figure S5.1f), but we did not have enough validation data for a full evaluation.

#### S5.2. Compatibility Between Assays

##### S5.2.1. Compatible and Incompatible: Single Chromatin Mark vs. Gene Expression

Using the methods described in previous sections, we identified promoters (± 2 kb from the TSS) with allelic imbalance in the chromatin states measured by H3K27ac, H3K27me3, etc. We determined the compatibility between AS promoter chromatin states and AS gene expression in a straightforward way. The allele with more active promoter chromatin should have higher expression levels, otherwise the promoter and the gene are incompatible. Similarly, alleles with more repressed promoter chromatin are compatible with lower expression levels. We treated histone marks H3K27ac, H3K4me3, and H3K4me1, chromatin openness indicated by ATAC-seq or by DNase-seq, and the binding of EP300, POLR2A, POLR2AP, and CTCF, as marks of active chromatin. Histone marks H3K27me3 and H3K9me3, and CpG methylation were considered as marks of repressed chromatin.

Because AS gene expression and/or AS chromatin state can be tissue specific and/or individual specific, we did not merge compatible (or incompatible) promoter-gene pairs that appeared in multiple samples. Overall, the average number of ASE genes compatible with at least one of the 13 marks was 35 per tissue per individual, while the average number of ASE genes suitable for the compatibility analysis (i.e., the promoters of these genes were accessible for measuring the potential AS chromatin state indicated by any of the 13 marks) was 226 per tissue per individual. See Figure S5.2a for more compatibility results and File: AS_allhets_alt_allele_ratio_eqtl_intersect.tsv for all the eQTL-ASB events in the plots.

We note that some assays were performed twice for a given tissue of a given individual. For example, the RNA-seq of individual 3’s liver includes two experiments (ENCSR226KML and ENCSR504QMK), while there is only one H3K27ac ChIP-seq experiment for the same sample. In another example, there are two CTCF ChIP-seq experiments for individual 3’s spleen (ENCSR756URL and ENCSR773JBP), while there is only one RNA-seq experiment for the same sample. In these cases, we combined ASE genes or ASB promoters that were called from either of the duplicated experiments, excluding those where the directions of the allelic imbalance were the opposite in the two experiments. The combined ASE genes and ASB promoters were analyzed for compatibility. File: Supp_data_compatibility.xlsx lists the compatibility of genes with ASE in each tissue and individual.

To test the numbers of compatible vs. incompatible promoter-gene pairs, we identified genes that have ASE in at least one tissue of at least one individual. We shuffled the gene-promoter relation for these genes and calculated the ratio of N_compatible vs. N_incompatible. We repeated this process 1,000 times for each chromatin mark (after excluding replicates where N_incompatible is zero) to calculate a Z-score of the ratios shown in Figure 7D.

##### S5.2.2. Compatibility with AS Proteomics

Of the high-stringency ASP set, 114 were overlapped with ASE events calculated from RNA-seq data, 58 showed compatibility, and 56 showed incompatibility (Figure S5.2b). The z-score 0.26 of the ratio of the compatible to the incompatible (based on ASP/ASE pairs being randomized 1,000 times) was not significant, indicating that the compatibility between the RNA-seq and protein-level allele expression is near random. Although some of this incompatibility is likely due to technical issues, manual examination of the most biased ASPs-overlapping ASEs showed compelling evidence for AS protein expression, implicating post-translational regulation (Figure S5.2c) (Ghaemmaghami et al., 2003; Greenbaum et al., 2003). For some of the incompatible cases, there are clear biological reasons for the difference between ASP and ASE ratios such as frameshift variants. File: Supp_data_compatibility.xlsx shows details of the significant 114 ASPs overlapping with ASEs.

##### S5.2.3. Enrichment of ASE Genes Near ASM Promoters

We checked the association between the ASM of the promoters and the ASE of the corresponding genes while ignoring the compatibility between the two. We annotated all hetSNVs showing ASM with the closest associated gene based on the refGene database (O'Leary et al., 2016) using ANNOVAR (Wang *et al.*, 2010). Chi-squared tests were used to determine whether ASE is significantly enriched among genes associated with ASM hetSNVs in promoter-like sequences (PLSs) identified by ENCODE (Encode Project Consortium et al., 2020) with or without AS of TF binding compared to that among genes only associated with ASM hetSNVs in non-cCREs (Encode Project Consortium *et al.*, 2020). Significant enrichment was called at p-values < 0.05 and error in enrichment was estimated based on binomial distribution. Figure S5.2d shows that genes with ASE are more highly enriched near PLSs with ASM and/or TF binding than near non-cCREs with ASM.

### S6. Supp. Content for Main Text Section "Generalized Application #3: Using the EN-TEx Resource to Extend eQTL Annotations to Hard-to-obtain Tissues"

#### S6.1. Correlation Between Chromatin Features and eQTL Activity

We overlapped chromatin (histone marks and TFs ChIP-seq, DNase-seq, ATAC-seq) peaks with GTEx catalogs of eQTLs and fine-mapped eQTLs for every EN-TEx sample, and observed a higher overlapping proportion in the case of fine-mapped eQTLs (Figure S6.1a). We obtained fine-mapped eQTLs after intersecting eQTLs with posterior probability ≥ 0.8 from the three GTEx fine-mapping eQTL catalogs (CAVIAR, CaVEMaN, and DAP-G; see https://gtexportal.org/home/datasets#filesetFilesDiv15).

Next, we identified 1,353,101 SNVs that show tissue-specific eQTL activity: these SNVs are GTEx eQTLs in ≥ 5 EN-TEx tissues, and are not GTEx eQTLs in ≥ 5 other EN-TEx tissues. Thus, for every SNV we defined two groups of tissues: i) tissues in which the SNV is an eQTL, and ii) tissues in which the SNV is not an eQTL. In this way, we could compute at which frequency the SNVs are marked by a particular histone modification when they do or do not show eQTL activity. We observed that SNVs are more likely to be marked by a given histone modification in the tissues in which they are eQTLs, compared to the tissues in which they are not eQTLs (Figure S6.1b). For each histone mark, we excluded lowly marked SNVs, i.e., SNVs overlapping chromatin peaks in < 10% of all EN-TEx ChIP-seq samples for that particular histone mark.

#### S6.2. Building a Predictive Model That Transfers eQTLs From a Donor Tissue to a Target Tissue

In this section, we explain how we trained a machine learning model that can predict the tissue-specific activity of a set of eQTLs. Specifically, one such model takes as input a set of SNV-eQTLs previously identified in a given tissue (i.e., donor tissue) and predicts whether each of these SNVs is an eQTL also in another tissue (i.e., target tissue). Practically, the goal is to transfer eQTLs from a donor tissue to a target tissue (Figure 8A and Figure S6.2a).

For a given donor-target tissue pair, we first retrieved the set of GTEx donor-tissue SNV-eQTLs associated with one single eGene (see Figure S6.2b, "n. of eQTLs"), and randomly partitioned them into training and test sets (containing 70% and 30% of SNV-eQTLs, respectively). Next, we trained a random forest model by providing, for every SNV-eQTL, a number of features related to either the donor or target tissue (see Figure S6.2c). The response class was defined as "yes" if the SNV-eQTL was annotated as GTEx eQTL in the target tissue, and otherwise as "no". We trained the random forest model using the R package caret (Kuhn, 2008) and by implementing a 5-fold cross-validation schema.

Only chromatin data from EN-TEx assays (histone marks and TFs ChIP-seq, DNase-seq, ATAC-seq) were employed. For this reason, the number of features employed in the model for a given target tissue depends on the type of EN-TEx experiments available for that particular tissue (i.e., if no ATAC-seq experiments were performed for lung tissue, then features “ATAC” and “ATAC_k” would not be employed to predict eQTLs in lung tissue). We downloaded a BED file containing repeated regions annotated in GRCh38 from http://genome.ucsc.edu/cgi-bin/hgTables, after setting “group” = “repeats” and “track” = “Repeatmasker”. We downloaded a BED file containing ENCODE candidate cCREs from https://api.wenglab.org/screen_v13/fdownloads/GRCh38-ccREs.bed.

Because GTEx eQTLs catalogs are available for matched EN-TEx tissues, we considered all possible pairs of donor-target tissues among 28 deeply sampled EN-TEx tissues (see Figure S6.2b), leading to a total of 756 (28*27) predictive models. For simplicity, we can consider these as different tissue-specific parameterizations of the general predictive model. We hereafter refer to the 756 donor-target models as “submodels”. Thus, a submodel is defined as the model for a particular donor-target tissue pair. For each target tissue, we have 27 submodels, each using a different donor tissue.
In the case of artery aorta, we combined data from experiments performed on both ascending aorta (Individuals 1 and 2) and thoracic aorta (Individuals 3 and 4).

#### S6.3. Model Performance, Validation, and Novel Application

We used several metrics (see Figure S6.3a) to evaluate the performance of each submodel on either the 5 cross-validation folds (Figure S6.3b) or the test set (Figure 8B and Figure S6.3c).

We further decomposed the submodels’ performance by considering different sets of SNVs (Figure 8C). Specifically, within a given submodel’s test set, we identified sets of true positives (TP: SNVs classified as eQTLs in a given target tissue that are also GTEx eQTLs in the same tissue), false negatives (FN: SNVs not classified as eQTLs but that are GTEx eQTLs in the target tissue), false positives (FP: SNVs classified as eQTLs but that are not GTEx eQTLs), and true negatives (TN: SNVs not classified as eQTLs that are not GTEx eQTLs) SNVs. The violin plots in Figure 8C show distributions of GTEx nominal p-values in the corresponding target tissue for these four sets of SNVs (each point of the distribution corresponds to the median p-value of an SNV set in one of the 756 submodels). Of note, the significance of the TP and FP sets is stronger compared to the FN and TN sets, respectively. This suggests that our model i) predicts the strongest among all GTEx eQTLs in a target tissue and ii) could help prioritize some of the SNVs with marginally significant p-values discarded by GTEx.

For 4 of the 28 tissues we focused on additional eQTL catalogs other than GTEx, available from (Kerimov et al., 2021): i) pancreatic islets eQTLs (van_de_Bunt_2015 dataset) matched to pancreas (PNCREAS); ii) muscle eQTLs (FUSION dataset), matched to skeletal muscle (GASMED); iii) skin eQTLs (TwinsUK dataset), matched to both suprapubic (SNINNS) and lower-leg (SKINS) skin. Thus, for these 4 tissues we evaluated the proportion of eQTLs identified by these studies (SNVs with p-value < $10^{-5}$) that were also classified as “eQTLs” in the test set of the relevant target tissue (pancreas, muscle, or skin) by our submodels (see Figure

8D). eQTL catalogs used in this analysis were downloaded from ftp://ftp.ebi.ac.uk/pub/databases/spot/eQTL/csv/ (files ".all.tsv.gz").

Currently, large-cohort eQTL studies are restricted to tissues such as blood that can be easily obtained from donors (e.g., over 30,000 donors in (Vosa et al., 2021)), while most other tissues are difficult to profile in a large number of individuals. For this reason, and to showcase the utility of our predictions, we have directly applied our model to a set of >1.5 M blood eQTLs from (Vosa *et al.*, 2021). In this way, we could predict which of these blood eQTLs are active in every EN-TEx tissue (Figure 8E and Figure S6.3d-e). We downloaded the eQTL catalog from https://molgenis26.gcc.rug.nl/downloads/eqtlgen/cis-eqtl/ (2019-12-11-cis-eQTLsFDR0.05-ProbeLevel-CohortInfoRemoved-BonferroniAdded.txt.gz). Specifically, we selected 1,547,430 blood eQTLs found to be associated with only one eGene in the original study, and lifted them over to assembly GRCh38 using LiftOver (https:
278 //genome.ucsc.edu/cgi-bin/hgLiftOver). The metric "GTEx v8 eQTL-eGene regression slope", which was one of the predictive features employed in the training step (Figure S6.2c), was not available in this second eQTL catalog; thus, for these predictions we instead used the metric $\log_2$(Z-score). Since blood tissue is not included in the EN-TEx collection, and given that we do not currently have any submodel using blood as donor tissue, we applied each of the 756 submodels trained on GTEx data to this blood eQTL set. These results are available on the EN-TEx webpage (see File: "predictions.blood.eQTLs.tar.gz"). As an example, when using artery aorta as donor tissue, we transferred up to 60% of the blood eQTLs to some of the EN-TEx tissues, such as thyroid and tibial artery (Figure 8E and Figure S6.3d-e). These results were computed after excluding those eQTLs contained in the original training sets.

#### S6.4. Model Interpretation

To facilitate the interpretation of the model, we computed, across the test set of each submodel, the correlation between the level of a particular feature at a donor-tissue eQTL and the probability of classifying the donor-tissue eQTL as an eQTL in the target tissue. In Figure 8F we show, for the first 15 features in Figure S6.2c, the strongest correlation coefficient (i.e., the coefficient with the largest absolute value) obtained across all 756 submodels. This analysis highlights how most chromatin features, with the exception of H3K27me3, are positively correlated with predicting eQTLs in a given tissue, while other features are negatively correlated (e.g., tissue specificity of the eGene and distance from the eGene's TSS).

Figure S6.4a shows a comprehensive representation of these correlation patterns across all predictive features and submodels. In this specific case we discarded 4 of the 39 features ("POLR2A", "POLR2Aphospho5", "EP300", and "POLR2A_k") since they were not used in a large proportion of the submodels (because ChIP-seq assays for these 3 TFs were performed on a limited number of tissues; for more details on these features see Figure S6.2c). We thus focused on 432 donor-target submodels that did not have missing data for the remaining 35 features. The heatmap in Figure S6.4a shows, for each of these 432 submodels (rows), Pearson's correlation coefficients between the level of predictive features (columns) at donor-tissue eQTLs in the target tissue, and the probability of donor-tissue eQTLs being classified as eQTLs also in the target tissue (clustering method: “Ward.D2”, clustering distance: “manhattan”). The vast majority of the chromatin features show stronger correlations in a specific set of submodels, as highlighted by hierarchical clustering (cluster at the bottom). We have found that these submodels use donor tissues with larger GTEx sample sizes (Figure S6.4a, right side). Thus, chromatin features have a stronger impact when transferring eQTLs from donor tissues with larger sample sizes, which tend to detect more eQTLs albeit with lower effects (GTEx Consortium, 2017; 2020). By contrast, certain features appear to be systematically either negatively (“tissue_specificity”, “tss_distance”, “H3K27me3_k”) or positively (“sum”, “is_proximal”, “H3K36me3”, “H3K36me3_p”) correlated with the SNV’s probability of being an eQTL, independent of the donor tissue. These results suggest that donor-tissue eQTLs with a higher number of chromatin peaks and/or marked by H3K36me3 in the target tissue are more likely to be eQTLs in the target tissue as well. Conversely, donor-tissue eQTLs associated with tissue-specific genes, or that are located far from the eGene’s TSS, are less likely to be transferable from one tissue to another.

Because of this, and with the goal of simplifying the interpretation of these models, we evaluated whether two simple rules could help transfer eQTLs from one tissue to another. In Figure 8G we show that, on one hand, donor-tissue eQTLs either characterized by strong chromatin activity (feature “sum” ≥ 3) in the target tissue, or whose eGene is constitutively expressed across EN-TEx samples (feature “tissue_specificity” < 0.8), tend to be eQTLs also in the target tissue (rule #1: 67% eQTLs, 33% not eQTLs). On the other hand, donor-tissue eQTLs that have low chromatin activity (feature “sum” = 0) in the target tissue and whose eGene shows tissue-specific expression (feature “tissue_specificity” > 5) are less likely to be eQTLs in the target tissue (rule #2: 23% eQTLs, 77% not eQTLs). Figure 8G refers to the specific case employing testis as the donor tissue, and thyroid as the target tissue. Figure S6.4b shows that these findings are generalizable across all the 756 donor-target tissue pairs.

### S7. Supp. Content for Main Text Section “Generalized Application #4: Decorating ENCODE Regulatory Elements with EN-TEx Tissue & AS Information”

#### S7.1. Decoration of Candidate Cis Regulatory Elements (cCREs) from ENCODE Encyclopedia

##### S7.1.1. Signal Normalization Method

In order to overcome batch effects, matrices of gene expression and histone marks’ values were quantile-normalized across samples (tissues and donors). The choice of the quantile normalization method was made after performing benchmarking of a number of normalization methods. The methods selected for the benchmarking are among the ones analyzed in a recent publication (Valikangas et al., 2018): quantile normalization, smooth quantile normalization, upper-quartile normalization, variance stabilization normalization, and local regression normalization (two variants: LoessF and LoessCyc). These normalization techniques are widely applied in other bioinformatics fields, such as microarray and proteomics analyses. The pilot analysis was performed independently for two cell lines, K562 and GM12878, for which different polyA+ RNA-seq evaluation datasets were produced by the Wold, Gingeras, and Graveley labs during ENCODE Phase II. The benchmarking consisted of three steps: i) for each method, we computed the distribution of Pearson's and Spearman's correlation coefficients across all genes between each pair of samples; ii) we then ranked the methods based on the mean of the distribution of all genes' variance across samples (Chen et al., 2018), and iii) we then calculated the relative log expression distribution (distribution of $\log_2$ ratio for a given gene between one particular sample and the median across all samples), which should be close to 0 (Berghoff et al., 2017). Overall, the quantile and smooth quantile normalization techniques performed similarly between each other and better than the other methods. We thus opted for quantile normalization. In particular, for each of the histone modifications used in the decoration procedure below, we provide the quantile normalized fold-change signals of cCREs across all the available tissues and individuals. The data file for each of the histone modifications is a data matrix, in which each row corresponds to a cCRE and each column corresponds to a tissue from an individual. As a result, the element in the matrix is the quantile normalized signal of the histone modification observed in the cCRE from a tissue. These files are available as File: cCRE_histoneSignals_qnorm.tar.gz.

##### S7.1.2. Decoration of Regulatory Annotations

We used the ChIP-seq datasets of both active and repressed marks to decorate (i.e., re-annotate) the cCREs from the Encyclopedia, which are based on a set of high-quality DNase hypersensitive sites (DHS) (Encode Project Consortium *et al.*, 2020). The ENCODE regulatory elements consist of 0.9 million cCREs averaging ~400 bp. For each type of functional genomic data, we normalized the activity signals of the cCREs from all tissues and focused on the cCREs with relatively strong signals (Figure S7.1a). In the decoration, we considered three active marks (H3K27ac, H3K4me1, and H3K4me3) and three repressed marks (H3K27me3, H3K9me3, and DNA methylation). ChIP-seq datasets were uniformly processed using the ENCODE standard pipeline, including alignment, quality control, and peak calling. With the uniformly processed ChIP-seq datasets, the average epigenomic signals were calculated and normalized for a registry of cCREs from the Encyclopedia (Figure S7.1a). Namely, we first calculated the average fold-change against the control, typically input DNA, for each cCRE. The average fold-change was quantile normalized independently across experiments but jointly between individuals and tissues. Finally, the scores for each experiment were scaled from 1 to 10. For a particular tissue type, we defined a set of cCREs for each epigenomic mark that are considered as "active" (i.e., thresholding the normalized and scaled quantile values of the cCREs). The thresholding value was calculated for each assay by maximizing the similarity – the fraction of shared active cCREs – among the four individuals across tissues. We used the average threshold score across the transverse colon, spleen, and esophagus since those were the most commonly comprehensive assays across individuals.

For each tissue, we then defined a set of active, repressed, and bivalent cCREs based on their active and repressed epigenomic signals, respectively (Figure S7.1b; Figure S7.1c as an example from spleen). Briefly, the active cCREs show high activity for only the active marks (i.e., H3K27ac, H3K4me1, and H3K4me3); the repressed cCREs show high activity for only the repressed marks (i.e., H3K27me3, H3K9me3, and DNA methylation); and the bivalent cCREs show high activity for both the active and the repressed marks. The cCREs were then separated into distal and proximal groups according to their distance to TSSs (proximal as those within 2 kb of annotated TSSs). We also intersected these cCREs with the CTCF binding sites from the matched tissue type to define CTCF+ and CTCF- cCREs. Finally, the active and repressed cCREs were further annotated using their allelic signature to identify a set of AS and non-AS ones, respectively. In the AS decoration, we used the allelic signature from the matched epigenomic marks to define the active/repressed AS and non-AS cCREs. Any active/repressed cCREs intersecting with the AS cCREs were considered to be active/repressed AS. The active/repressed AS cCREs from different individuals were pooled together to generate a set of active/repressed AS cCREs in the corresponding tissue. We found that the numbers of repressed cCREs are comparable to those of active cCREs in many tissue types, highlighting the necessity of decoration using the repressed markers (Figure S7.1d). Finally, we provided the cCRE decoration results in all the tissue types (see File: cCRE_Decoration.matrix or separately File: active.combined_set.txt.zip, File: bivalent.combined_set.txt.zip, and File: repressed.combined_set.txt.zip) (Figure S7.1e).

In order to further subset the cCREs, we created an annotation set that focuses on regions with high H3K27ac signals. We call this set the “stringent” annotation set. To create this stringent annotation, we intersected the cCRE regions with the top 1% of scored regions as prioritized by the H3K27ac feature from the Matched Filter (Sethi et al., 2020). This stringent annotation was further used in other analyses and labeled as “stringent” in the main manuscript and figures. A file containing these stringent regions (bed file) can be found in File: stringent.regions.MF.hg38.bed. These bed regions represent the highest-scoring Matched Filter regions.

##### S7.1.3. Repressed Regions

Gene activation and repression can be mediated through the combination of different histone marks. Historically, much effort has been devoted to elucidating how genes are activated; however, evidence is emerging to demonstrate the requirement of appropriate heterochromatin formation for the preservation of genome stability and the cell type-specific silencing of genes (Becker et al., 2016). In the mammalian genome, H3K9me3 and H3K27me3 are well-documented histone marks enriched for “constitutive” and “facultative” heterochromatin, respectively. For genomic regions not containing any active regulatory elements (cCREs), we have identified a set of elements that are marked by either H3K9me3 or H3K27me3 and do not have any active marks (H3K27ac, H3K36me3, H3K4me1, and H3K4me3) or transcriptional activities as fully repressed in the EN-TEx tissues. Regions within the ENCODE4 GRCh38 blacklist (ENCSR636HFF) and GENCODE gene list (GRCh38_v24) were removed. In summary, 45,207 non-overlapping elements of size no less than 200 bp (roughly approximate nucleosome size) are uniquely marked by H3K9me3, spanning 12,655,795 bp (less than 0.4%) of the reference genome, and 24,006 elements by H3K27me3, spanning 7,474,178 bp (less than 0.3%). Identified elements can be found in File: ENTEx_fully_repressed_regions_independent_of_cCREs.bed. As shown in Figure S7.1f, nearly 75% of these elements are specifically repressed in a certain tissue, and the rest show some degree of tissue specificity. It was previously known that H3K27me3-enriched facultative heterochromatin contains repressed genes in a cell-type-specific manner, whereas H3K9me3-enriched constitutive heterochromatin mainly occur at the same gene-lacking regions in every cell type (Gerlitz, 2020; Saksouk et al., 2015). Observations also suggest that large domains of H3K9me2/3 form in a cell-type-specific manner and can influence cell identity by silencing lineage-inappropriate genes and impeding the conversion of terminally differentiated cells into a different cell type, highlighting a role for H3K9me3 in cell-type-specific gene regulation (Becker et al., 2017; Becker *et al.*, 2016; Nicetto and Zaret, 2019; Ninova et al., 2019; Pace et al., 2018).

DNA methylation is one major contributor to gene repression, and it has been reported to interact with H3K9me3 in chromatin repressive pathways (Du et al., 2015). We further analyzed the methylation rate of CpG sites within these repressed elements. WGBS of CpG sites were available for 11 EN-TEx tissues that also have H3K9me3 and H3K27me3 ChIP-seq data. For the same tissue from different donors, we aggregated the CpG reads by taking the sum of reads from all donors, and considered a CpG site to be methylated (meCpG) when the site is covered by at least 5x reads and the ratio of meCpG reads is at least 50%. The overall meCpG rate in each tissue was calculated and used as a control to evaluate the meCpG rate in H3K9me3- and H3K27me3-marked elements. As shown in Figure S7.1f, H3K9me3 uniquely marked elements show significantly (t-test, p-values $< 0.05$) higher meCpG rates than elements uniquely marked by H3K27me3. Compared with the control, H3K9me3-marked regions seem to be hypermethylated, whereas H3K27me3-marked regions are hypomethylated. This is consistent with the current understanding of constitutive heterochromatin and facultative heterochromatin, of which the former is defined by high levels of DNA methylation and H3K9me3 and the later displays DNA hypomethylation and high H3K27me3 (Saksouk et al., 2014).

##### S7.1.4. Validating Annotations Using 3D Genome Organization

Chromosome compartments that are observed from principal component analysis (PCA) analysis on a Hi-C correlation matrix give insight into the activity level of the chromatin. Chromosomes are divided into two distinct compartments, A and B, at a megabase scale (Lieberman-Aiden et al., 2009). The A compartment (positive values) corresponds to the active regions on the chromosome and the B compartment (negative values) corresponds to the inactive regions. Chromatin interactions are constrained within the compartment types, e.g., the loci in A compartment interact with the loci in the same compartment. Since A/B compartment assignments are proxies for the activity level of different loci, our tissue-specific regulatory element annotations can be validated by looking at their corresponding compartment in the tissue-level Hi-C data. We showed that our annotated tissue-specific active regulatory elements are dominantly located in the active compartment of the chromosomes of corresponding tissues, with a significantly higher number of regulatory elements per megabase observed in the positive compartment values when layered onto the first principal component of the Hi-C data.

We have assessed where the cCREs are located with respect to the chromatin compartments. For that, we first binned the genome into 1 MB consecutive bins. We then counted the total number of cCREs in each bin and divided that number by the total number of cCREs in the genome. This gave us the cCRE density per 1 MB. We then plotted this density against the A/B compartment score obtained by the first eigenvector of the correlation matrix calculated from the Hi-C contact matrix. We performed this analysis for the master cCRE list from ENCODE3, tissue-specific active cCRE list derived in this study, more restrictive tissue-specific active cCRE list derived in this study, and tissue-specific repressed cCRE list derived in this study. Below are the scatter plots for two tissues and four individuals (Figure S7.1g).

#### S7.2. Tissue Specificity

In this section, we compared the tissue specificity of genes, cCREs, and epigenomic peaks in a systematic manner. We included protein-coding genes, non-coding genes, different catalogs of decorated cCREs, and diverse types of epigenetic marks.

##### S7.2.1. Tissue-specific Results

There are many methods for determining tissue specificity, most of which are based on continuous positive values (Kryuchkova-Mostacci and Robinson-Rechavi, 2017). Here, we chose the simple method of tissue count to determine the tissue specificity of genes/cCREs based on thresholds (Kryuchkova-Mostacci and Robinson-Rechavi, 2017). We did this because we can consistently apply this method across different annotations including cCREs, genes, TSSs, and epigenomic peaks. Most of the other methods that are based on continuous positive values can be only applied on one annotation category (e.g., genes). Briefly, all genes and cCREs (and also equally applicable to peaks and TSSs) were defined as active or inactive by thresholding their expression/activity level in a particular tissue type. The numbers of tissue types in which these genes/cCREs are active were then summarized. For each gene/cCRE annotation category, we then calculated a tissue-specificity score using the number genes/cCREs that are active in only one tissue type divided by the total number of genes/cCREs in the category. The uniqueness scores range from 0 to 1, with higher scores indicating stronger tissue specificity.

For the genes, we included three gene types: protein-coding genes (from MS and RNA-seq), lncRNAs, and pseudogenes. To better estimate the expression level of pseudogenes, we applied our previously developed pipeline to quantify the expression level of pseudogenes, which can minimize the effects of multiple mapping bias in RNA-seq data (Sisu et al., 2020) (Figure S7.2c). We then applied this pipeline to all the three gene types, and defined a set of active genes in the tissues by thresholding the FPKM (fragments per kilobase of transcript per million mapped reads) values (FPKM > 1 for protein-coding genes; FPKM > 0.5 for lncRNAs and pseudogenes) (Figure S7.2a). Over 40% and 35% of the detected pseudogenes and lncRNAs, respectively, were actively transcribed in a single tissue, confirming that non-coding RNAs exhibit higher tissue specificity than protein-coding genes (Necsulea et al., 2014; Ransohoff et al., 2018). Of the pseudogenes demonstrating tissue specificity, a large fraction showed transcriptional activity only in testis (Figure S7.2b). For the cCREs, we used the decorated annotations in the tissues to calculate the tissue-specificity scores as described above (Figure S7.2d and Figure S7.2e). We also explored the tissue specificity of regulatory elements and epigenomic peaks (Figure 9B-C). The epigenetic profiles analyzed, including H3K27ac and DNase-seq, demonstrated tissue specificity, with the exception of DNA methylation, which exhibited ubiquity. An example of the tissue specificity of RAMPAGE data (for TSSs) is shown in Figure S7.2f.

The tissue specificity of the genes, cCREs, and epigenomic peaks are in File: Tissue_Specificity.zip.

##### S7.2.2. Tissue Specificity of ASB and ASE

Similar to H3K27ac-ASB cCREs (Figure 9), most ASE genes were detected in a single tissue (Figure S7.2g). For the ~20 genes that were detected ASE across all tissues, the allelic imbalance is in the same direction (Figure S7.2h). We further compared our pan-tissue H3K27ac-ASB and ASE genes with housekeeping genes. Annotation results are shown in Figure S7.2i-j.

##### S7.2.3. The Effect of Tissue Specificity on Conservation

The tissue-specificity influence on conservation is shown in Figure S7.2k. Candidates are separated into categories of active, bivalent, and repressive. The number of candidates, rare derived allele frequency (DAF) values, and corresponding total SNP counts (from gnomAD) are given as a function of increasing tissue specificity (shared tissue count). In order to select rare variants, a minor allele frequency (MAF) of 0.05 was used.

Various decorations further subset categories and affect the conservation level, based specifically on whether elements are distal or proximal as well as if they are CTCF bound or not. Conservation is shown for both phastCons (cross-species) and rare DAFs (cross-population) in Figure S7.2l.

We show the conservation across active and repressed cCREs in both ubiquitous and tissue specific cases in Figure S7.2m. We include the results across the 1,000 Genomes, Pan Cancer Analysis Working Group (PCAWG), and gnomAD projects. Additionally, we show an increase in conservation when filtering for high H3K27ac signals (using stringent definitions for active elements with Matched Filter (Sethi *et al.*, 2020)), which is supported by all three datasets. The overall conservation calculation is described in section S6.1.4.

##### S7.2.4. The Relationship Between Purifying Selection and Regions Exhibiting Allele Specificity

The fraction of causative variants may be estimated by purifying selection. The analysis by the NIH Roadmap Epigenome project of epigenomes from 36 distinct cell and tissue types from 13

donors suggests that the donor genomes harbor on average at least 200 regulatory variants that are under purifying selection and therefore detrimental (Onuchic *et al.*, 2018).
In order to calculate the purifying selection on AS events, population-scale variants from three cohorts were used. We used two measures of purifying selection and conservation for this analysis. The first is the fraction of rare variants, which is calculated as #rare/(#rare + #common) for variants falling in a given AS region. In order to categorize variants as rare or common, ancestral alleles were used (i.e., the measurement was a DAF) and a MAF threshold of 0.05 was used. MAF is a commonly used metric for calculating selection in populations (Chen *et al.*, 2016; Khurana *et al.*, 2013; Onuchic *et al.*, 2018). When considering the number of variants in each category, we found that across all tissues and individuals, 8,294 out of 128,448 ASB variants were rare. Of the 2,711,078 total non-ASB variants, we found a total of 274,287 rare variants. In total there were 40,123 ASE+ variants, of which 2,961 were rare. Finally, of the 624,210 ASE- variants, 70,370 were rare. The second is phastCons, which measures the cross-species conservation (Siepel et al., 2005). All purifying selection analyses (rare DAF and phastCons) were performed for AS/non-AS cCREs, ASB/non-ASB H3K27ac regions, and ASE/non-ASE genes. The results are shown in Figure S7.2n. Furthermore, we found that proximal AS events in the promoter were under stronger selection as compared to distal AS events.

#### S7.3. Relating Encyclopedia Decorations to QTLs & GWAS Loci

We utilized various methods to evaluate the regulatory impact of our cCRE decorations. QTL and GWAS SNPs are important functional genomic variants and are useful for interpreting the function of our decorations. We performed GWAS enrichment analysis using eQTL and GWAS SNPs to assess the disease relevance of our cCRE decorations.

##### S7.3.1. QTL Enrichment Analysis

We estimated the QTL (eQTL and sQTL) enrichment in the cCREs by calculating an odds ratio (OR) score using the numbers of real QTL SNPs and control SNPs located in the cCREs compared to those in the baseline regions (Figure S7.3a).

$$OR = \frac{a / b}{c / d}$$

in which a is the number of QTL SNPs in the cCREs; b is the number of control SNPs in the cCREs; c is the number of QTL SNPs in the baseline region; and d is the number of control SNPs in the baseline region.

The eQTL and sQTL SNPs were downloaded from GTEx v8 (GTEx Consortium, 2017). The baseline regions are the union of all the functional and putative functional regions in the human genome, including coding regions (CDS), untranslated regions (UTRs), noncoding RNA genes, open chromatin regions, TF binding sites, active and repressed histone peaks from multiple tissue and cell types, and evolutionary conserved regions (Finucane et al., 2015). The set of control SNPs was generated with the same number and same MAF distribution as the real QTLs, and this procedure was repeated 30 times to calculate a standard deviation for the SNP enrichment. The results of the QTL enrichment are in File: QTL_enrichment.zip.

We also compared the eQTL/sQTL enrichment in the regulatory elements from EN-TEx with those from Roadmap (Figure S7.3b and Figure S7.3c). First, we found that the distal regulatory elements from EN-TEx show stronger enrichment than the enhancer annotation from Roadmap. In addition, the active proximal regulatory elements from EN-TEx show stronger eQTL/sQTL enrichment than the TSS-associated annotations from Roadmap.

S7.3.2. GWAS Enrichment Analysis

We downloaded the GWAS tag SNPs from the GWAS Catalog (Buniello et al., 2019). We performed several steps of quality control to generate a set of high-quality GWAS tag SNPs by removing some insignificant SNPs (p-values > 5*10-8), low-confidence SNPs, and SNPs from non-European studies. We also removed all SNPs in the HLA locus (for hg38: chr6:29,723,339-33,087,199). Next, we extended the set of tag SNPs by including the SNPs in high linkage disequilibrium (LD scores > 0.6) with the tag SNPs, which can generate more SNPs to increase the statistical power in the enrichment analysis. Some GWAS with very few LD-extended SNPs were removed. Finally, this resulted in a clean dataset with ~70K unique tag SNPs from 1,140 GWAS covering 717 unique traits.

We then applied the hypergeometric test to estimate the enrichment of the GWAS tag SNPs in the cCREs from a particular tissue type (Figure S7.3d).

$$P(X=k)=\frac{\binom{K}{k}\binom{N-K}{n-k}}{\binom{N}{n}}$$

in which N is the total number of cCREs in the genome; K is the total number of cCREs that carry GWAS tag SNPs; n is the number of cCREs in a particular tissue type; and k is the number of cCREs in a particular tissue type that also carry GWAS tag SNPs. Notably, we extended the cCREs 500 bp on both sides in the calculation (Figure S7.3f). The results of the GWAS tag SNP enrichment are in File: GWAS_enrichment.zip.

For the active distal cCREs, we identified 141 GWAS that are enriched in at least one tissue type (Figure S7.3g). However, for the active proximal cCREs, we did not find any enriched GWAS in any tissue type. These results are consistent with previous studies showing that the causal GWAS SNPs are enriched in the enhancers instead of the near-gene promoters (Whalen and Pollard, 2019; Yao et al., 2014) and also suggest that the active distal cCREs from our decoration are indeed significantly enriched in enhancers as we observed in the original Roadmap annotations (Figure S7.3h).

Stratified linkage disequilibrium score regression (LDSC) values were also calculated for each tissue using 1,000 Genomes LD scores and GWAS summary statistics provided by Bulik-Sullivan, et al. This approach regresses chi-square statistics from the GWAS summary statistics with LD scores to estimate partitioned heritability in a disease-specific manner. The p-value indicates enrichment for a particular trait within an annotation.

In Figure S7.3g, we show the p-value enrichment of each tissue with respect to various GWAS traits. Notably, distal active AS regions experienced higher enrichment compared to distal active non-AS regions (Figure S7.3i), and both types of regions experienced higher enrichment compared to the original Roadmap annotations (Figure S7.3c). For LDSC enrichment analysis of distal active elements in coronary artery (Figure S7.3e), we found stronger associations between AS elements with respect to celiac disease, neuroticism, and type II diabetes, which were elucidated in previous clinical studies (Almas et al., 2017; Gajulapalli and Pattanshetty, 2017; Naito and Kasai, 2015). These results demonstrate that AS elements can significantly improve GWAS trait enrichment compared with the total set of elements across different traits as well as diverse tissue types, indicating that AS elements are valuable for the interpretation of GWAS data and that they potentially help pinpoint small subsets of regulatory elements driving a trait in specific tissues.

#### S8. Supp. Content for Main Text Section “Discussion”

##### S8.1. The EN-TEx Supplemental Data Repository

All processed data files are described in detail in their corresponding sections of this document. When mentioned, these files are referred to as "File: file_name". All of these files are hosted on the EN-TEx data portal website ENTEx.encodeproject.org, with the same file names as in "File: file_name". Additionally, on the website, each file is followed with the supplement section number that contains the description of that file. Links to all the raw data of this project can be found on the EN-TEx data portal website as well.

##### S8.2. Open-consent of Data

In concert with the GTEx project, an Institutional Review Board-approved consent form was written and given to the next-of-kin of each donor. The consent form allows for unrestricted access to data collected as part of the GTEx and EN-TEx project, including unrestricted use of the primary data and metadata collected from each donor. It was made clear that although no identification of the donor or family constituted part of these data, it is within the realm of possibly that individual identification could be made. Specific details of the consent document are contained here:
https://www.genome.gov/Pages/Research/ENCODE/GTEx_Consent_ENCODE_addendum_10-9-14.pdf

##### S8.3. Explorer Tool

The EN-TEx explorer tool, which can be run in R, installed as an offline executable, or hosted on a website through integration with Amazon Web Services, allows for the interactive exploration of low-dimensional visualizations created by an in-house data analysis pipeline (Figure S8.3). This pipeline performs dimensionality reduction on cCRE signals, genomic data, and proteomic data. Methods include PCA, variational autoencoder (VAE), uniform manifold approximation and projection (UMAP) (McInnes et al., 2018), potential of heat diffusion for affinity-based transition embedding (PHATE) (Moon et al., 2019), set intersection plots generated by user-specified thresholds (Sets), and t-distributed stochastic neighbor embedding (tSNE) (Van der Maaten and Hinton, 2008). The pipeline then generates the tool programmatically in R Shiny in one of the three forms above.

The visualizations generally cluster samples from common tissues together. Through extensive precomputation, the tool allows users to interactively adjust analysis parameters, including scaling, normalization, feature subsetting, method-specific hyperparameters, the type of visualization used (ggplot2, plotly 2D, plotly 3D, boxplot, heatmap, UpSetR, Venn diagram), and the appearance of the resulting figures. Users are able to save figures as images, download analysis results as Excel spreadsheets, or bookmark their sessions as short URLs that can be easily shared (Figure S8.3). To install the tool, please consult the Github README. Instructions and documentation regarding tool usage can be found by pressing the “Instructions” button on the tool. The input files for the explorer tool are available on the EN-TEx website. File: ENTEx.Explorer.cCRE.Combined.zip contains the cCREs. File: ENTEx.Explorer.Expression.Combined.zip contains the expressed genes and File: ENTEx.Proteomics.cCRE.Combined.zip contains the mass spectrometry proteomic data.

#### S8.4. The EN-TEx Chromosome-Level Data Visualization Tool

Because the EN-TEx data span a wide range of the human genome, it may be useful to visualize the distribution over each chromosome. Accordingly, we present the EN-TEx Chromosome-Level Data Visualization Tool, which generates heatmaps for datasets for all assays, individuals, and tissues present in the EN-TEx data catalog. The data, which were initially in BED format, were preprocessed with in-house Bash and Python scripts and converted to GRCh38 coordinates using liftOver (Hinrichs et al., 2006) prior to the generation of the plots using the R package chromoMap (Anand and Rodriguez Lopez, 2020). The EN-TEx Chromosome-Level Data Visualization Tool was also used to generate the plots in Figure 4B of the main text.

The EN-TEx Chromosome-Level Data Visualization Tool can be accessed at ENTEx.gersteinlab.org. Users can specify any combination of parameters (individual, assay, ploidy, and color) for a track and subsequently generate interactive plots containing one to four tracks each by pressing the “Submit” button (Figure S8.4a). By default, the tool generates heatmaps for the data of each chromosome at a fixed resolution of 2.5 Mb. The user can get information about the data displayed in a specific bin by hovering over the bin with a mouse cursor.

The “Advanced” tab contains tools for custom chromosome and region selections. To view the data in only one chromosome, one can select the chromosome of interest in the

“Chromosomes” dropdown menu. To view a subset region of the chromosome, the user can input the region in the format initial_position:final_position in the “Region” text box (e.g., if the user wishes to visualize data between 1 Mb and 2 Mb, the user would input 1000000:2000000). The tool automatically sets the resolution of the data for subset regions of the chromosome to the length of the inputted interval divided by a factor of one hundred (e.g., for the 1000000:2000000 interval, the resolution will be equal to 10 kb). Users can also visualize the data as heatmaps accompanied by either histograms or scatterplots by selecting the desired option in the “Plot Type” dropdown. A series of plots generated with this tool is shown in Figure S8.4b.

#### S8.5. Additional Data Exploration with SCREEN

The SCREEN website (https://screen.encodeproject.org/) is a center for the ENCODE cCRE registry and annotation. This website is routinely used by researchers all over the world. The annotations of cCREs using our EN-TEx data are also available at the SCREEN website (Figure S8.5). The EN-TEx data provide many unique annotations. For example, different from the annotation from other datasets, the EN-TEx data indicate the repressive states of cCREs. In addition, the EN-TEx data specify whether each cCRE is AS in terms of functional genomic signals. Figure S8.5 is a step-by-step guide from the main webpage of SCREEN to an cCRE with repressive states in multiple human tissues. In line with the repressive states, this cCRE has no CTCF binding and is not AS.

#### S8.6. Providing Evidence for the Buffering Hypothesis Using AS cCREs and Housekeeping Genes

Genetic variants in cCREs can change functional signal and gene expression. For these changes to occur, the variants need to escape from buffering effects (Onuchic *et al.*, 2018). Such effects are strong in important genomic regions. We used allele specificity as a proxy for escaping buffering. Based on our allelic decoration, we evaluated the allele specificity of housekeeping genes expressed in EN-TEx tissues, as shown in Figure S8.6a. For each tissue, expressed protein-coding genes were split into housekeeping genes and non-housekeeping genes according to Housekeeping and Reference Transcript Atlas (http://www.housekeeping.unicamp.br) (Hounkpe et al., 2021). The two-sided Fisher exact test was performed to measure the enrichment of AS housekeeping genes. We found that, compared with non-housekeeping genes, the expression of housekeeping genes shows less allele specificity, supporting the buffering hypothesis. We further examined the allele specificity of proximal active (pAct) cCREs in a ± 10 kb window centered on the TSS (defined by the gene starting site) of each housekeeping and non-housekeeping gene. The cCREs flanking housekeeping genes are significantly (Figure S8.6a, paired-tissue two-sided t-test, p-value < 2.2e-16) longer than cCREs flanking non-housekeeping genes. To control for this factor, we split genes into 20 bins based on the total length of flanking cCREs. Within each bin, cCRE length remains similar (paired-tissue two-sided t-test, p-value > 0.05) between housekeeping and non-housekeeping genes. The bins with less than 30 housekeeping or non-housekeeping genes were removed from further analysis. The pAct cCREs flanking housekeeping genes are less likely AS than the ones flanking non-housekeeping genes (two-sided t-test).

The buffering effect is likely due to redundant TFs. To test this, we counted the number of TF motifs that intersect with each CTCF+ and CTCF- cCRE in each tissue. For this calculation, we used the motifs of 206 TFs (CTCF excluded) from Cis-BP (Weirauch *et al.*, 2014). The total count of all TF motifs was compared between CTCF+ cCREs and CTCF- cCREs using a two-sided t-test. As shown in Figure S8.6b, for both distal and proximal cCREs, CTCF+ cCREs have significantly ($p$-value $< 0.05$) more TF motifs than CTCF- cCREs. In addition, we tested whether the genetic variants in large motif clusters tend to be associated with ASB. To this end, we identify the locations of the motifs of 660 TFs in the human genome. For each motif, we counted the number of all motifs within 500 bp of the motif. A larger number suggests that the motif likely has many functionally redundant motifs. According to the proxy of redundancy, we divided all the motifs evenly into two groups: likely redundant or not. The genetic variants are considered ASB if significantly imbalanced reads are observed in any of the tissues of the four individuals with any assays. As a result, we found that the genetic variants in the motifs that are likely redundant tend not to be AS, consistent with the buffering effect.

#### Supplementary Figures

#### Figure S1. Supp. Figures for Main Text Section “Improvements in Genome Analysis from Uniform Multi-tissue Data Collection & Diploid Genome Mapping”

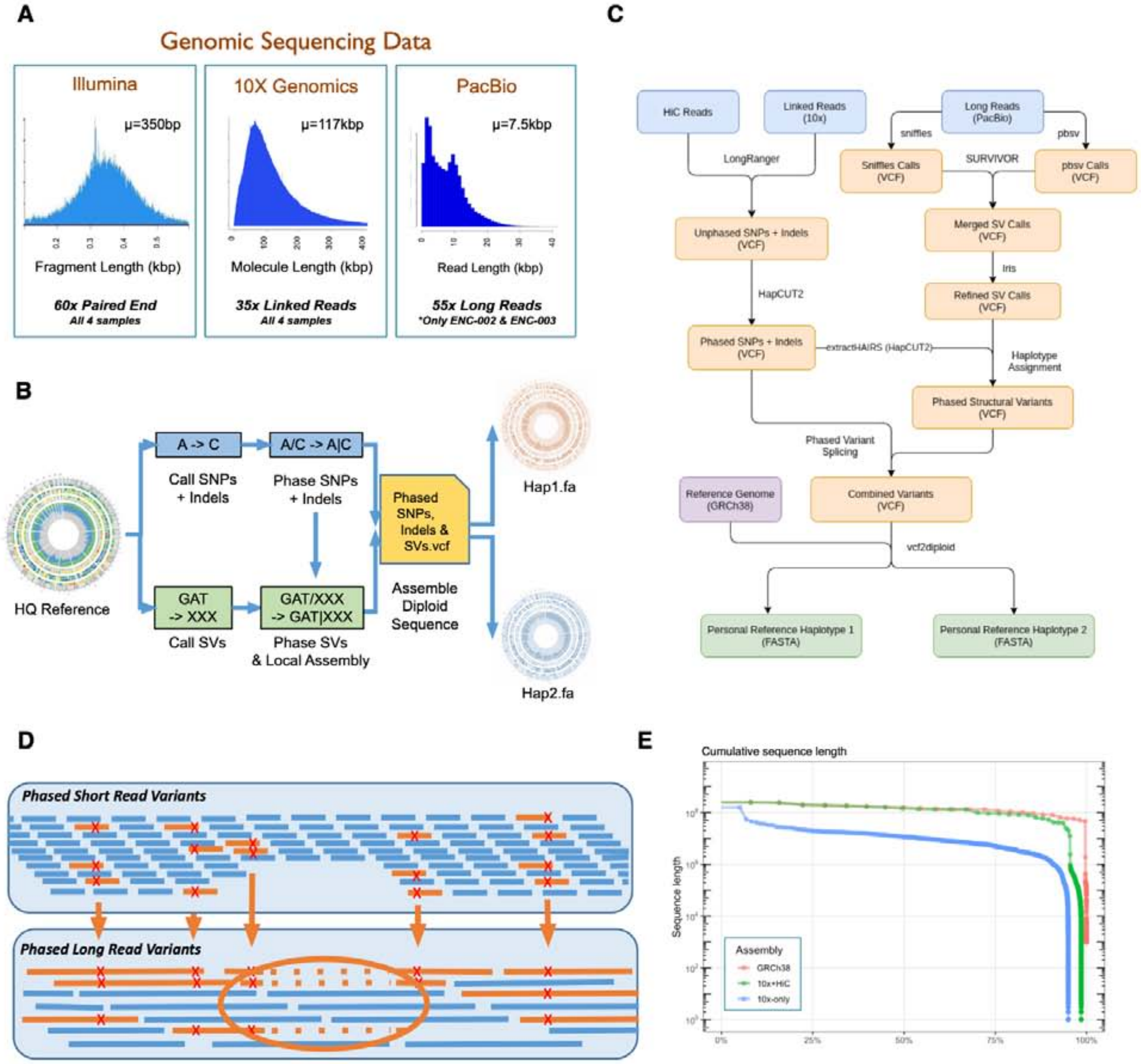

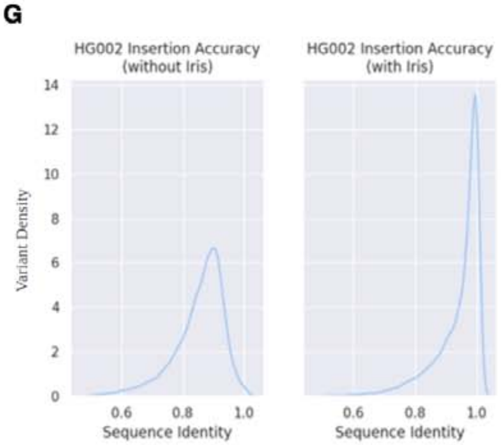

**F**

| Individual | Illumina | 10x linked-read | long-read |
|---|---|---|---|
| 1 | ENCSR246MMZ | ENCSR410ALS (ENCFF854ZQU) | individual1_ONT_WGS.fastq (individual1_SV.vcf) |
| 2 | ENCSR549QWF | ENCSR613XEH (ENCFF065JXC) | ENCSR723PGJ (ENCFF867GGB) |
| 3 | ENCSR961ZRM | ENCSR997HAI (ENCFF166PQX) | ENCSR664TEU (ENCFF840IWS) |
| 4 | ENCSR420NDH | ENCSR456SNK (ENCFF541HLI) | individual4_ONT_WGS.fastq (individual4_SV.vcf) |

**Figure S1.1a. Personal genome construction**

**(A)** Summary of whole-genome sequencing. All four individuals were sequenced with regular Illumina short reads and 10x linked reads. Individuals 2 and 3 were additionally sequenced with PacBio long reads. The figure shows the sequencing depth and read-length distribution under each platform. **(B)** An overview of the CrossStitch workflow. SNPs and small indels are called and phased, while unphased SV calls are obtained independently. Then, the phase blocks from the small variants are used to assign haplotypes to heterozygous SVs, and the phased variants are used to construct a phased personal genome assembly based on a high-quality reference sequence. **(C)** Detailed summary of the CrossStitch method. This diagram shows an overview of the CrossStitch methods with the specific software and data types used. **(D)** SV phasing with CrossStitch. Phased small variants are used to assign a haplotype to each long read, and SVs are phased by observing the haplotypes of the long reads, which indicate the presence of that variant. In this example, a deletion is phased when all three of the long reads including that deletion have small variants that are unique to the orange haplotype. **(E)** Phase block length. This figure shows the size of phase blocks in individual 2 obtained with HapCUT2 when performing small variant phasing with 10x reads only, as well as with a combination of 10x and Hi-C reads. When both data types are used, the contiguity of the phase blocks obtained is very similar to that of GRCh38. **(F)** Accession numbers of WGS data. Accession numbers in parentheses are for VCF files. The long-read sequencing data and VCF files of individuals 1 and 4 are available at ENTEx.encodeproject.org. The other data can be downloaded from the ENCODE portal. **(G)** Refining novel insertion sequences with Iris. This figure shows the sequence similarity of ONT calls to CCS calls in the Genome-in-a-Bottle sample HG002, used to benchmark the performance of Iris. The sequence similarity between two sequences S and T is calculated as edit_distance(S, T) / [max(length(S), length(T))].

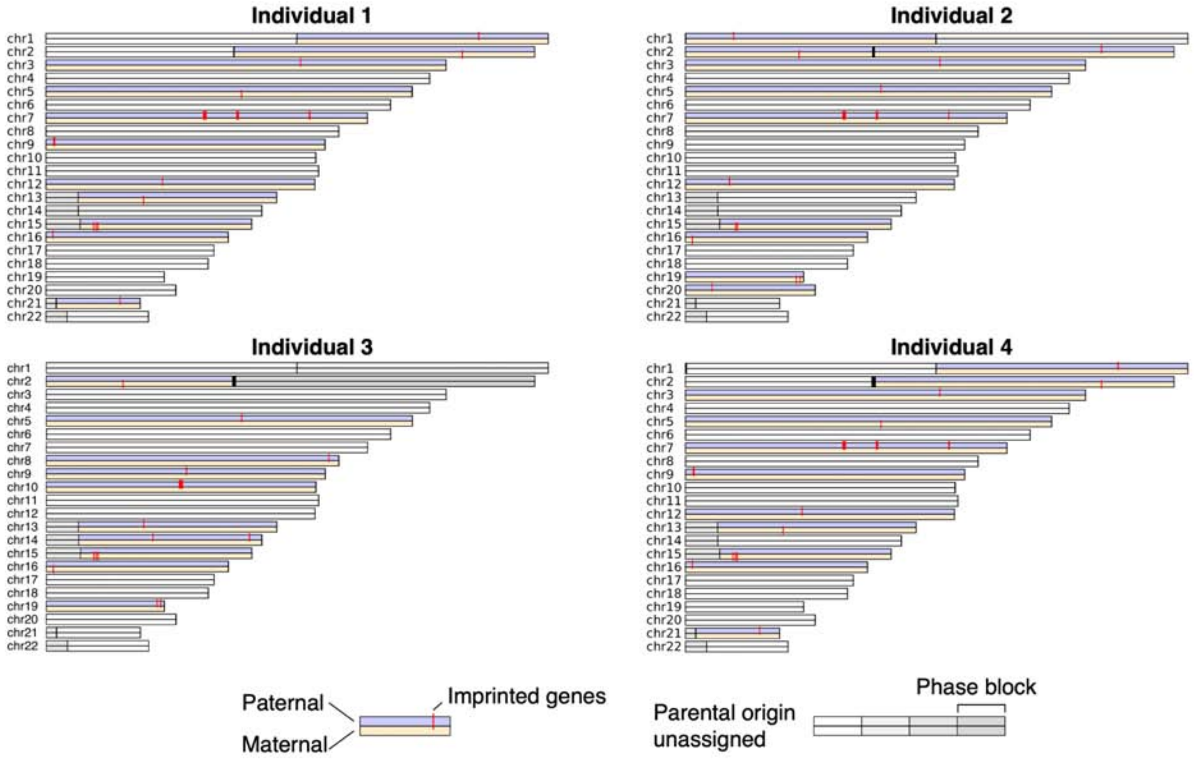

**Figure S1.1b. Phase blocks of personal genomes with parental origins**
The parental origin of each phase block was determined based on the consistency between the direction of allele-specific gene expression and the direction of known imprinted genes. See Figure S1.1c for comparison details and S2.2 for the calculation of allele specific genes.

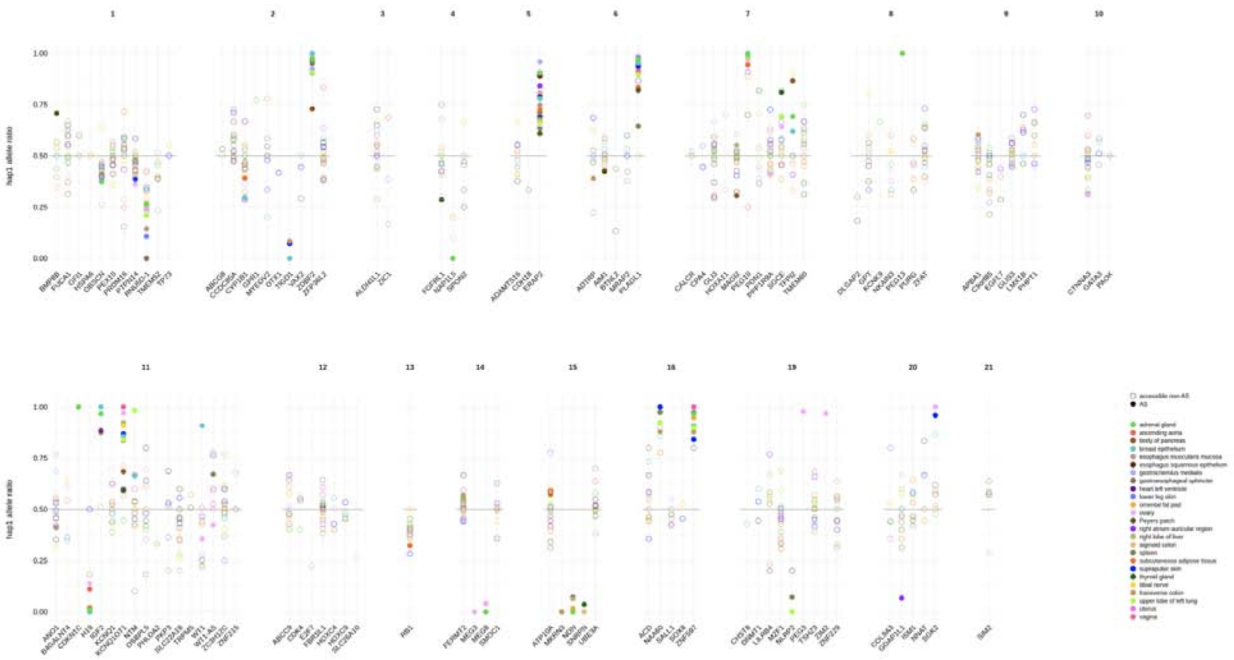

**Figure S1.1c. Consistency of ASE imbalance direction in known imprinted genes across tissue samples (individual 3)**

Fraction of reads preferentially mapping to haplotype 1 in known imprinted genes. For most genes, the direction of the significantly imbalanced genes (filled circles) is consistent across samples from different tissues (colors).

A

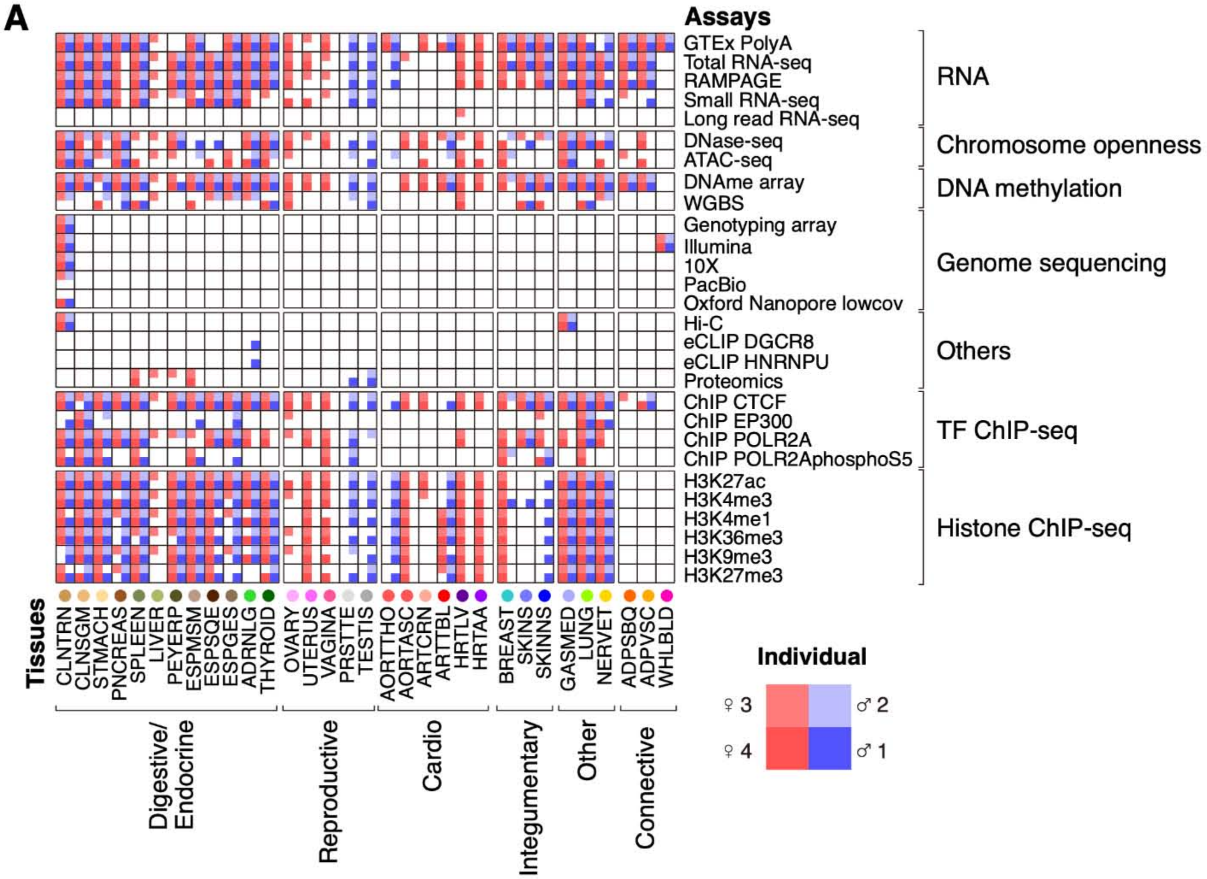

B

| ENTEx Tissue Names | Abbrev | Color Hex | GTEx Tissue Names |
|---|---|---|---|
| transverse colon | CLNTRN | #CC9955 | Colon_Transverse |
| sigmoid colon | CLNSGM | #EEBB77 | Colon_Sigmoid |
| upper lobe of left lung | LUNG | #99FF00 | Lung |
| stomach | STMACH | #FFDD99 | Stomach |
| spleen | SPLEEN | #778855 | Spleen |
| gastrocnemius medialis | GASMED | #AAAAFF | Muscle_Skeletal |
| adrenal gland | ADRNLG | #33DD33 | Adrenal_Gland |
| esophagus muscularis mucosa | ESPMSM | #BB9988 | Esophagus_Muscularis |
| thyroid gland | THYROID | #006600 | Thyroid |
| gastroesophageal sphincter | ESPGES | #8B7355 | Esophagus_Gastroesophageal_Junction |
| tibial nerve | NERVET | #FFD700 | Nerve_Tibial |
| body of pancreas | PNCREAS | #995522 | Pancreas |
| esophagus squamous epithelium | ESPSQE | #552200 | Esophagus_Mucosa |
| Peyer's patch | PEYERP | #555522 | |
| breast epithelium | BREAST | #33CCCC | Breast_Mammary_Tissue |
| suprapubic skin | SKINNS | #0000FF | Skin_Not_Sun_Exposed_Suprapubic |
| prostate gland | PRSTTE | #DDDDDD | Prostate |
| heart left ventricle | HRTLV | #660099 | Heart_Left_Ventricle |
| testis | TESTIS | #AAAAAA | Testis |
| vagina | VAGINA | #FF5599 | Vagina |
| lower leg skin | SKINS | #7777FF | Skin_Sun_Exposed_Lower_leg |
| tibial artery | ARTTBL | #FF0000 | Artery_Tibial |
| uterus | UTERUS | #FF66FF | Uterus |
| right atrium auricular region | HRTAA | #9900FF | Heart_Atrial_Appendage |
| ovary | OVARY | #FFAAFF | Ovary |
| omental fat pad | ADPVSC | #FFAA00 | Adipose_Visceral_Omentum |
| subcutaneous adipose tissue | ADPSBQ | #FF6600 | Adipose_Subcutaneous |
| ascending aorta | AORTASC | #FF5555 | Artery_Aorta |
| right lobe of liver | LIVER | #AABB66 | Liver |
| thoracic aorta | AORTTHO | #FF5555 | Artery_Aorta |
| coronary artery | ARTCRN | #FFAA99 | Artery_Coronary |

**Figure S1.2a. Functional genomics data**
**(A)** Data matrix of the EN-TEx resource showing tissues vs. assays. Each square is partitioned into four regions, representing the four individuals. **(B)** Information for EN-TEx tissues.
The table shows the full name, abbreviation, and color code of the EN-TEx tissues, as well as their matching relationship with GTEx tissues. This tissue color scheme is also used in other main and supplementary figures. Note there are 1,635 total experiments in EN-TEx, which includes control experiments and replicates (both of which are not explicitly shown in the data matrix). If we remove replicates and controls, the number of experiments is 1,275, which is equivalent to the number of cells in the data matrix.

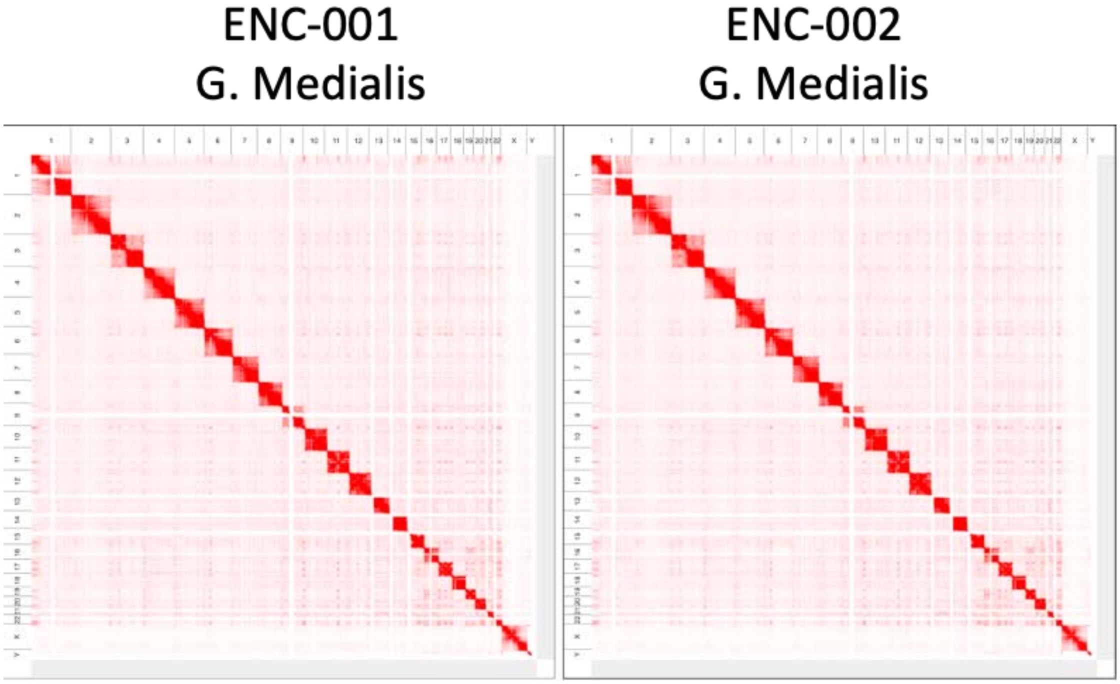

**Figure S1.2b. Example of reference-aligned genome-wide Hi-C maps for the skeletal muscle tissue of two individuals**

Number of paired reads (billions)

| | ENC-004 | ENC-003 | ENC-001 | ENC-002 |
|---|---|---|---|---|
| Gastrocnemius medialis | 1.53 | 1.41 | 1.60 | 1.38 |
| Transverse colon | 1.44 | 1.51 | 1.50 | 2.07 |

Number of contacts (billions)

| | ENC-004 | ENC-003 | ENC-001 | ENC-002 |
|---|---|---|---|---|
| Gastrocnemius medialis | 0.964 | 0.997 | 1.02 | 0.992 |
| Transverse colon | 1.06 | 1.10 | 1.08 | 0.958 |

**Figure S1.2c. Number of paired reads and number of contacts from reference-aligned genome-wide Hi-C contact maps**

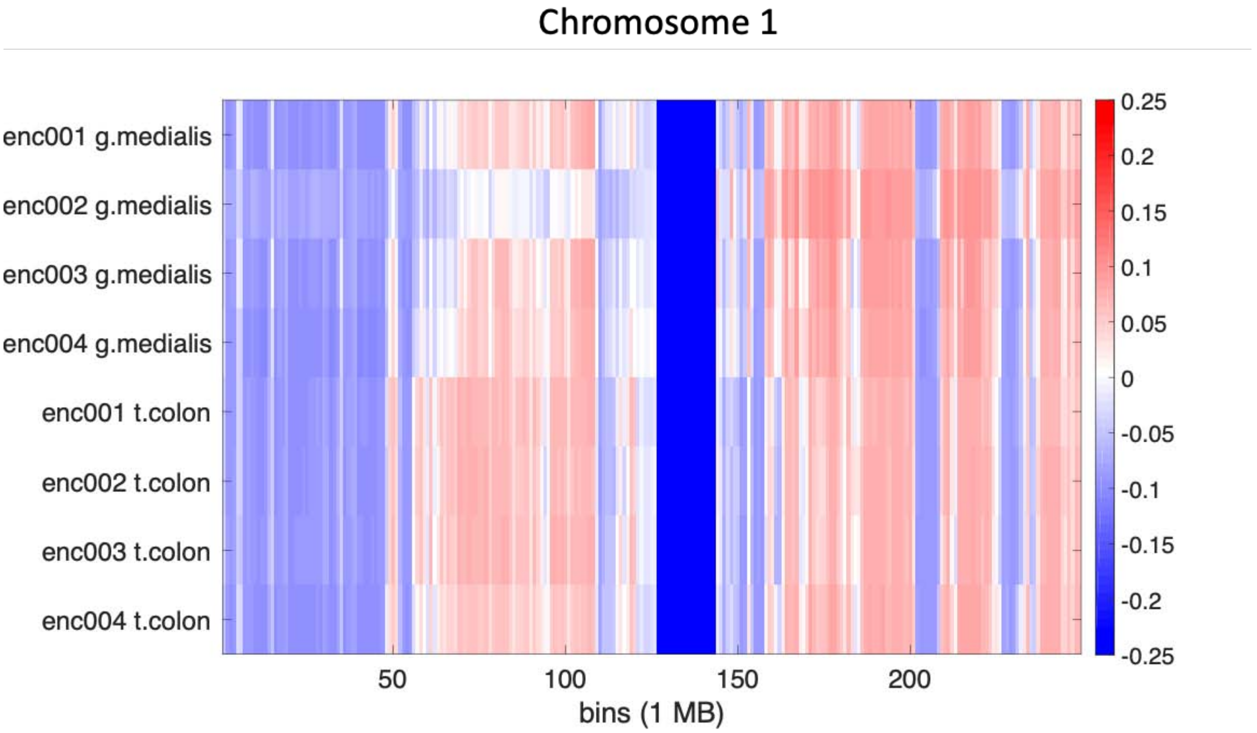

**Figure S1.2d. A/B compartment annotation of four individuals and two tissues for chromosome 1**

Red indicates that the 1 MB region is in the A compartment, whereas blue indicates that the region is in the B compartment. The dark blue band corresponds to the centromere.

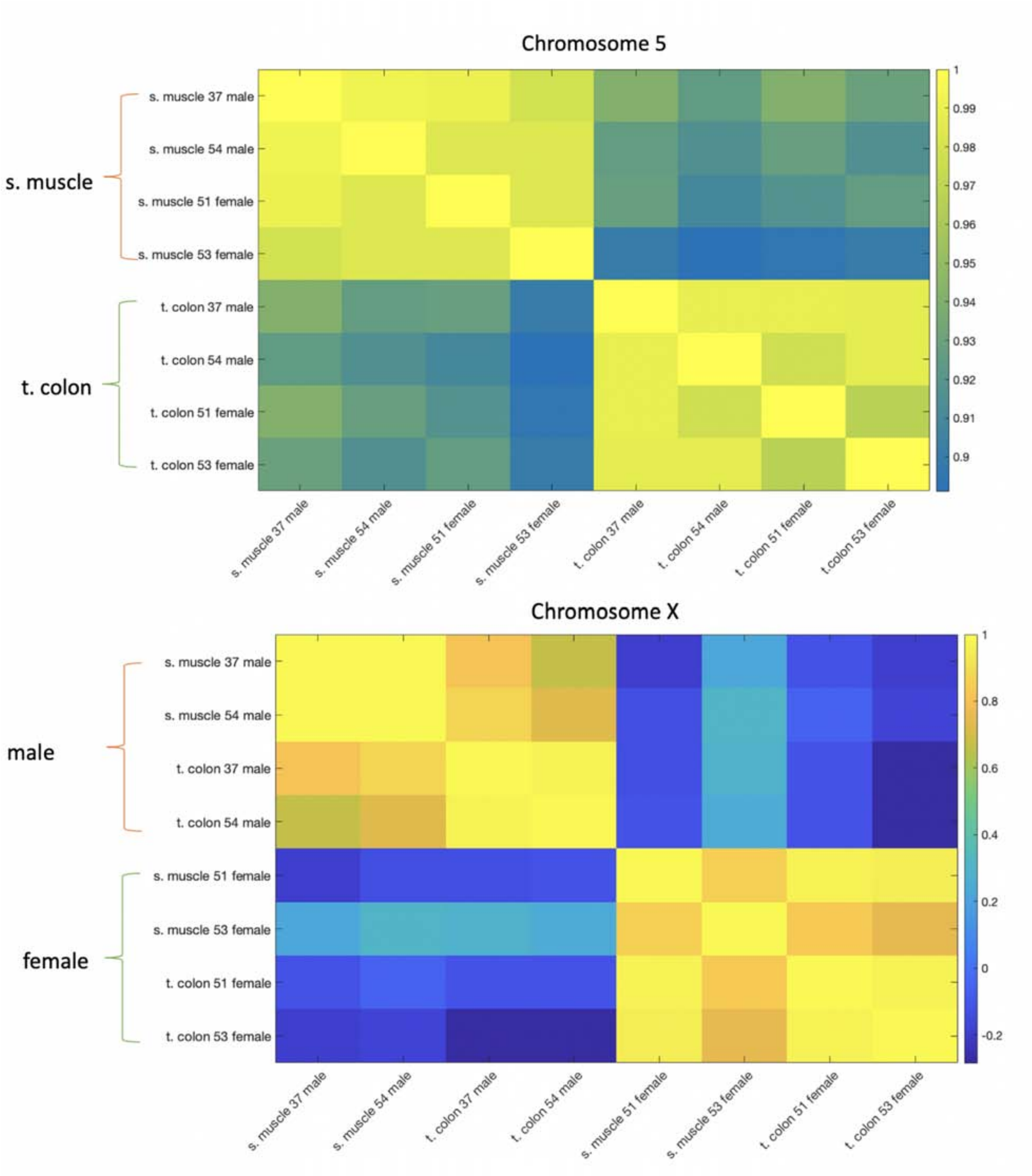

**Figure S1.2e. A/B compartments cluster based on tissue in autosomes or sex in chromosome X**

37 male, 54 male, 51 female, and 53 female correspond to individuals 1, 2, 3, and 4, respectively.

| | | CLNTRN | | | | GASMED | | | |
|---|---|---|---|---|---|---|---|---|---|
| Chrom | Total | Indiv 1 | Indiv 2 | Indiv 3 | Indiv 4 | Indiv 1 | Indiv 2 | Indiv 3 | Indiv 4 |
| chr1 | 12397710 | 6788490 | 6857968 | 6640741 | 6263058 | 6128125 | 6218242 | 6188268 | 6282354 |
| chr10 | 3579150 | 2953439 | 2990590 | 2916989 | 2831756 | 2915114 | 2923222 | 2929788 | 2996873 |
| chr11 | 3649051 | 2869655 | 2948299 | 2816640 | 2778830 | 2860206 | 2887171 | 2860242 | 2961438 |
| chr12 | 3552445 | 2969628 | 3041869 | 2919601 | 2856463 | 2939877 | 2980095 | 2940746 | 3026456 |
| chr13 | 2616328 | 1704423 | 1724468 | 1682424 | 1668618 | 1628849 | 1620200 | 1607366 | 1674749 |
| chr14 | 2290870 | 1410898 | 1444375 | 1384357 | 1381694 | 1413111 | 1418356 | 1402899 | 1433712 |
| chr15 | 2079780 | 1119023 | 1143407 | 1119432 | 1094365 | 1107323 | 1136572 | 1128846 | 1138837 |
| chr16 | 1631721 | 945521 | 979481 | 945231 | 922589 | 908303 | 956847 | 932352 | 922697 |
| chr17 | 1386945 | 1028955 | 1055611 | 1029448 | 1009073 | 1014825 | 1056832 | 1042864 | 1050488 |
| chr18 | 1292028 | 1076987 | 1086267 | 1064598 | 1062482 | 1067970 | 1061511 | 1060048 | 1081935 |
| chr19 | 687378 | 560395 | 570534 | 560516 | 559704 | 564088 | 575239 | 571257 | 578818 |
| chr2 | 11729746 | 8394990 | 8525226 | 8309864 | 7938906 | 7840021 | 7789126 | 7833348 | 8259552 |
| chr20 | 830116 | 679685 | 692916 | 663099 | 670959 | 674590 | 680375 | 673769 | 691748 |
| chr21 | 436645 | 207828 | 213035 | 203748 | 205200 | 206996 | 207260 | 206634 | 212110 |
| chr22 | 516636 | 217507 | 220358 | 218031 | 217149 | 216516 | 220675 | 219461 | 220895 |
| chr3 | 7862595 | 6015900 | 6134904 | 5902231 | 5774444 | 5434165 | 5464917 | 5391741 | 5857473 |
| chr4 | 7237110 | 5562209 | 5681942 | 5600166 | 5271711 | 5254951 | 5245142 | 5313566 | 5522909 |
| chr5 | 6590265 | 5129851 | 5233827 | 5049672 | 4852197 | 4903566 | 4859322 | 4876589 | 5108741 |
| chr6 | 5836236 | 4649176 | 4768735 | 4545579 | 4416778 | 4436206 | 4469419 | 4422407 | 4647679 |
| chr7 | 5076891 | 3880470 | 3956534 | 3827833 | 3644663 | 3778164 | 3740318 | 3791757 | 3882221 |
| chr8 | 4212253 | 3506901 | 3567642 | 3465779 | 3373636 | 3384990 | 3372726 | 3383295 | 3507176 |
| chr9 | 3829528 | 1910384 | 1963561 | 1897205 | 1867856 | 1748547 | 1802543 | 1761180 | 1909601 |
| chrX | 4868760 | 2923158 | 4102701 | 4010558 | 2739157 | 2745686 | 3901268 | 3961831 | 2903581 |
| chrY | 654940 | 44238 | 224 | 319 | 42376 | 42181 | 636 | 338 | 43792 |

**Figure S1.2f. Summary of significant interactions determined by FitHiC2**
The “Total” column presents the total number of intrachromosomal interactions for a given chromosome (e.g., chr1, chr2, …, chrY).

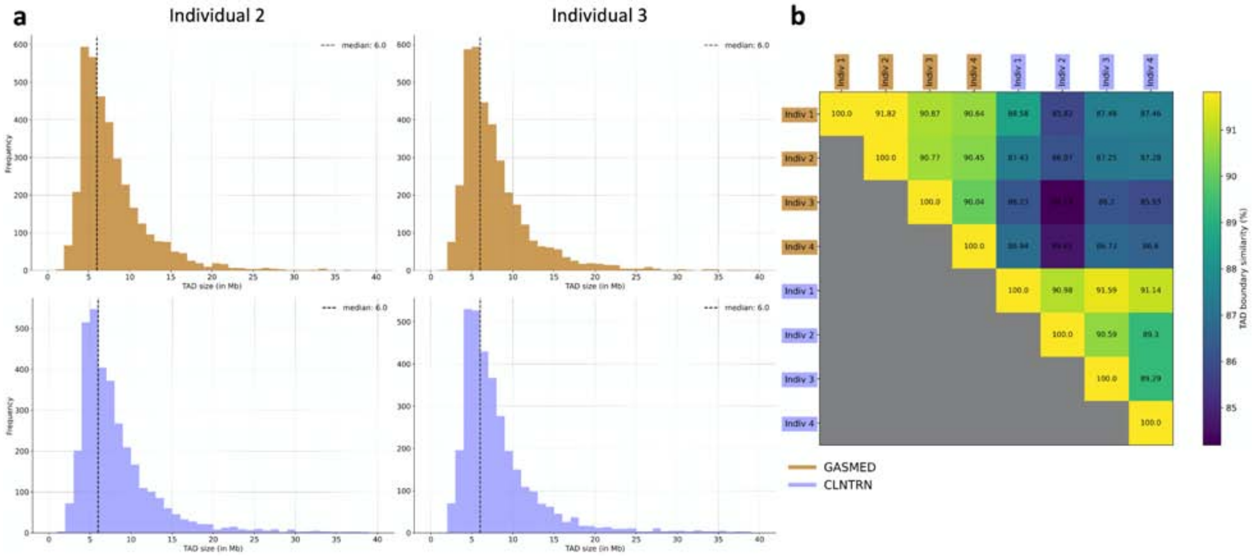

**Figure S1.2g. Comparison of TopDom TAD calls for EN-TEx individuals and available Hi-C tissues**

**(A)** TADs were shown to have a similar size distribution and median TAD size across individuals and tissues. The TAD size distribution of individuals 2 (left) and 3 (right) for available Hi-C tissue types gastrocnemius medialis (GASMED - top) and transverse colon (CLNTRN - bottom) are shown. **(B)** Pair-wise comparison of TAD calls across all four individuals and available Hi-C tissue types. TAD calls were shown to be more similar (i.e., located at the same position along a chromosome) for the same tissues from different individuals than between different tissues.

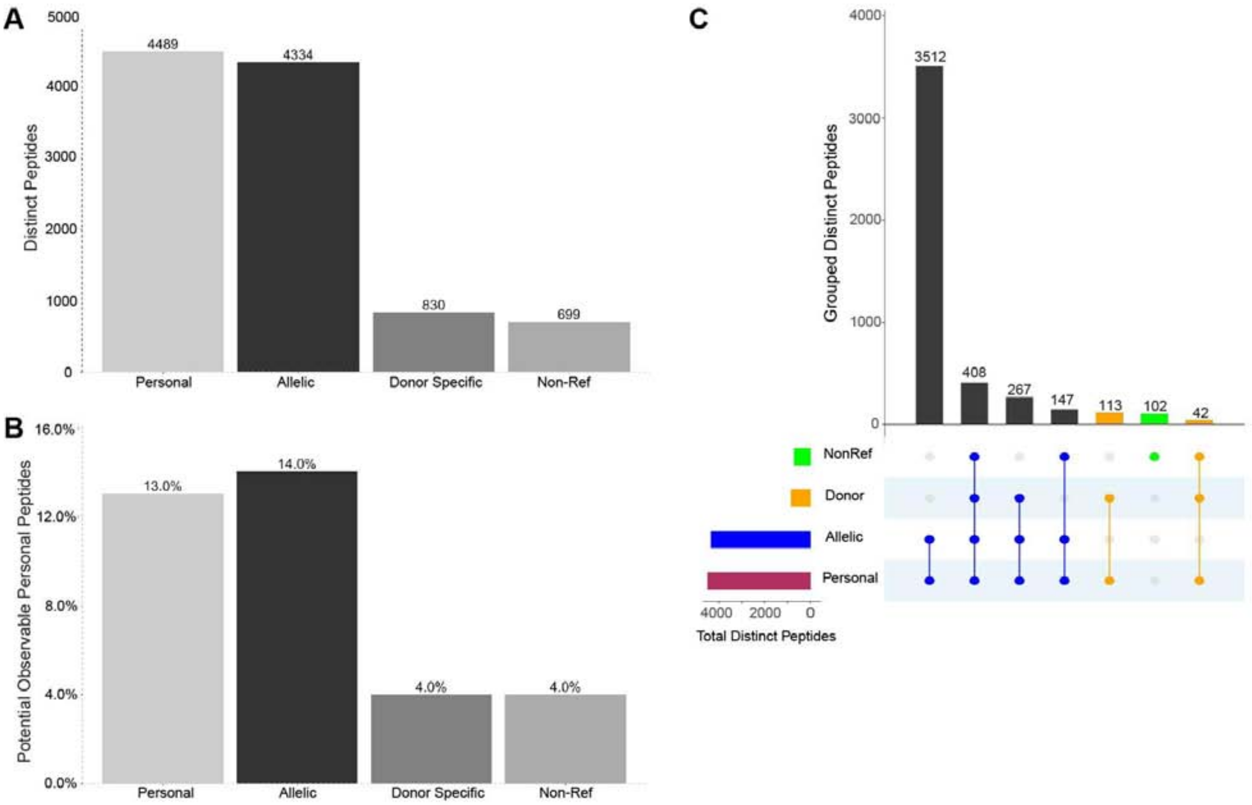

**Figure S1.2h. Observed personal peptide summary**

**(A)** Total number of significantly identified unambiguous personal peptides (filtered for 0.01 posterior error probability and unambiguous gene mapping). The personal category includes all types of personal peptides, allele-specific peptides are those that are specific to only one allele in at least one individual, donor-specific peptides are those that are completely absent in at least one of the four donors, and non-reference peptides are those that do not match the reference genome. Due to the use of the TMT method, there is a bias towards the most common peptide from among the four donors (usually the reference peptide), as TMT boosts the signal for common peptides. **(B)** Coverage of all potentially observable personal peptides calculated by an *in silico* tryptic digest. Although *in silico* peptides are filtered for unambiguity and are limited to amino acid lengths of 6-60, there will be a vast number of unobservable peptides due to MS-incompatible charge states and chemical properties. **(C)** UpSet plot showing the overlap in distinct peptides identified in the MS experiments among the four personal peptide categories. The majority of the peptides are considered personal because they are either donor-specific, AS, or both. There are a small number of non-reference peptides that are neither AS nor donor-specific; these are variant peptides that are common across all donors and alleles.

| Peptide | Type | Gene | Ensemble id |
|---|---|---|---|
| VETAGSEPGDTEPJEJGGPGAEPEQK | NewModel | HYOU1 | ENSG00000149428 |
| RPESPGDAEAAAAAAPGAPGGR | NewModel | SNX25 | ENSG00000109762 |
| SHMMDVQQGSTQDSAJK | NewModel | PDIA4 | ENSG00000155660 |
| SQGVQPJPSQGGK | NewModel | FAM120A | ENSG00000048828 |
| ASAAEGVGEPGASAGR | NewModel (nonATG) | WDR26 | ENSG00000162923 |
| HPKPEVJGSSADGAJJVSJDGJR | AddedModel | TNXB | ENSG00000168477 |
| DSNQGJYGJSPEGVDR | AddedModel | TNXB | ENSG00000168477 |
| SSJDTGSSJSTDR | AddedModel | IQSEC1 | ENSG00000144711 |
| SGASGASAAPAASAAAAJAPSATR | REFerror | CENPV | ENSG00000166582 |
| QTFENQVNR | REFerror | POLR2A | ENSG00000181222 |
| GGGSCVJCCGDJEATAJGR | REFerror | ZNF598 | ENSG00000167962 |
| VJWJDEJQQAVDEANVDEDR | REFerror | IQGAP2 | ENSG00000145703 |
| GPGGVWAAEAJSDAR | Multiple-Variants | SAA1 | ENSG00000173432 |
| JPQEQSQJPNPSEASTTFPESHJR | Multiple-Variants | IFI16 | ENSG00000163565 |
| GTJVTVSSASTK | IG Allelism | | |
| VTVSSASTK | IG Allelism | | |
| GTTVTVSSASTK | IG Allelism | | |
| VDEYJAWQHTTJR | AltAssembly | GSTT1 | ENSG00000277656 |
| GQHJSDAFAQVNPJK | AltAssembly | GSTT1 | ENSG00000277656 |
| VEAAVGEDJFQEAHEVJJK | AltAssembly | GSTT1 | ENSG00000277656 |
| AJEMENSQJCK | Some Evidence | HNRNPA0 | ENSG00000177733 |
| AEATESAMER | Some Evidence | HNRNPA2B1 | ENSG00000122566 |
| GAGSMATGJGEPVYGJSEDEGESR | Weak Evidence | NEDD4L | ENSG00000049759 |
| GSSPEAGAAAMAESJJJR | Weak Evidence | NPLOC4 | ENSG00000182446 |
| AJPGSSMADQAPFDTDVNTJTR | No Evidence | FBP1 | ENSG00000165140 |
| DTEQTJYQER | No Evidence | LAMB2 | ENSG00000172037 |

**Figure S1.2i. Novel peptides**

These are non-variant novel peptides identified with very high confidence that do not match known protein annotations in the GENCODE reference at the time of the experiment. These peptides were manually curated by GENCODE and annotated in combination with orthogonal evidence. Column 1 is the sequence of each novel peptide. Column 2 is the type of annotational outcome: NewModel = a new protein was annotated in GENCODE reference (nonATG, means the new model did not have a canonical ATG TSS); AddedModel = an existing protein annotation was adjusted or peptide provided additional support for an annotational change already in progress; REFerror = Although peptides matched genuine unannotated proteins an underlying error in the reference genome GRCh38 sequence assembly meant that they cannot be added to current GENCODE annotation; Multiple-Variants = The peptide could be potentially explained by complex variants; IG Allelism = Peptides matched in highly variable genomic regions; AltAssembly = Peptides matching alternative genome assemblies; Some Evidence = Peptides had some orthogonal evidence but fell just below current criteria for annotation of new protein model; Weak evidence = Very little orthogonal evidence to support change in annotation; No Evidence = No support for a change in annotation The affected genes and their Ensembl IDs are listed in columns 3 and 4. Additional information, e.g., the number of spectral reads matching the peptide, are provided in File: Supp_data_proteomics.xlsx on the EN-TEx resource website.

**A**

Precision Mapping Table

| | RNAseq | DNAseq | CHIPseq | Hi-C |
|---|---|---|---|---|
| Unique to Reference | 0.001 | 0.002 | 0.001 | 0.001 |
| Unique to Haplotypes | 0.020 | 0.045 | 0.016 | 0.024 |
| Gain= ((Hap1 U Hap2)-Ref)/Ref | 0.019 | 0.045 | 0.015 | 0.023 |

**B**

| DNAseq | ENC-001 37yo male | ENC-002 54 yo male | ENC-003 53 yo female | ENC-004 51 yo female | Average |
|---|---|---|---|---|---|
| Hap1&Hap2 (reads) | 224,383,498 | 260,499,463 | 243,467,335 | 266,766,596 | 244,872,476 |
| Ref (reads) | 214,959,471 | 249,767,410 | 233,252,112 | 254,366,883 | 234,192,822 |
| Ref only (%) | 0.2 | 0.2 | 0.3 | 0.2 | 0.2 |
| Hap1&&Hap2¬Ref (reads) | 9,842,890 | 11,359,688 | 10,864,254 | 12,883,160 | 11,196,768 |
| Hap1&&Hap2¬Ref ( %) | 4.4 | 4.4 | 4.5 | 4.8 | 4.5 |
| Improvement | 0.044 | 0.043 | 0.044 | 0.049 | 0.045 |

| HiC | ENC-001 37yo male | ENC-002 54 yo male | ENC-003 53 yo female | ENC-004 51 yo female | Average |
|---|---|---|---|---|---|
| Hap1&Hap2(reads) | 185,631,684 | 663,535,209 | 217,727,831 | 181,462,412 | 312,089,284 |
| Ref (reads) | 181,352,202 | 649,127,067 | 212,497,875 | 177,218,164 | 305,048,827 |
| Ref only (%) | 0.1 | 0.2 | 0.2 | 0.1 | 0.1 |
| Hap1&&Hap2¬Ref (reads) | 4,479,970 | 15,707,837 | 5,583,723 | 4,436,038 | 7,551,892 |
| Hap1&&Hap2¬Ref ( %) | 2.4 | 2.4 | 2.6 | 2.4 | 2.4 |
| Improvement | 0.024 | 0.022 | 0.025 | 0.024 | 0.024 |

| CHIPseq | ENC-001 37yo male | ENC-002 54 yo male | ENC-003 53 yo female | ENC-004 51 yo female | Average |
|---|---|---|---|---|---|
| Hap1&Hap2(reads) | 16,523,306 | 30,068,339 | 12,834,649 | 7,886,359 | 16,828,163.250 |
| Ref (reads) | 16,271,254 | 29,616,292 | 12,637,708 | 7,770,250 | 16,573,876 |
| Refonly (%) | 0.1 | 0.1 | 0.1 | 0.1 | 0.1 |
| Hap1&&Hap2¬Ref (reads) | 267,944 | 491,961 | 212,830 | 122,841 | 273,894 |
| Hap1&&Hap2¬Ref ( %) | 1.6 | 1.6 | 1.7 | 1.6 | 1.6 |
| Improvement | 0.015 | 0.015 | 0.016 | 0.015 | 0.015 |

| RNAseq | ENC-001 37yo male | ENC-002 54 yo male | ENC-003 53 yo female | ENC-004 51 yo female | Average |
|---|---|---|---|---|---|
| Hap.1(reads) | 14,512,594 | 38,983,875 | 39,631,595 | 38,983,875 | 33,027,984.750 |
| Hap.2(reads) | 14,508,357 | 39,004,820 | 39,660,757 | 39,004,820 | 33,044,688.500 |
| Reference (reads) | 14,252,554 | 38,639,784 | 39,294,659 | 38,639,784 | 32,706,695.250 |
| Ref (reads) | 14,053 | 40,126 | 58,062 | 40,126 | 38,091.750 |
| Ref only(%) | 0.1 | 0.1 | 0.1 | 0.1 | 0.1 |
| Hap1&&Hap2¬Ref (reads) | 383,723 | 721,245 | 724,982 | 721,245 | 637,798.750 |
| Hap1&&Hap2¬Ref (%) | 2.6 | 1.8 | 1.8 | 1.8 | 2.0 |
| Improvement | 0.025 | 0.018 | 0.017 | 0.018 | 0.019 |

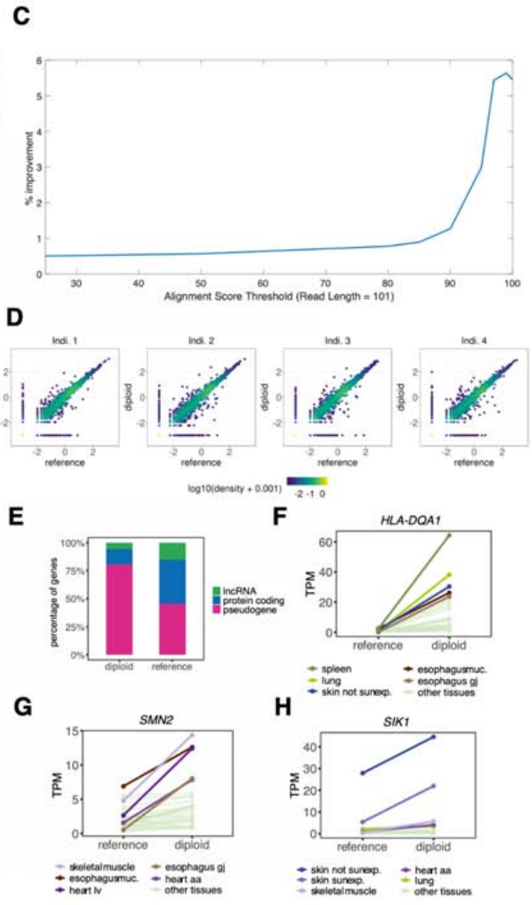

**Figure S1.3a. Mapping to personal genomes**

**(A)** Summary of percentage with precision mapping. **(B)** Using DNA from transverse colon tissues, we constructed both haplotype sequences for each individual. Here, we show the summary statistics when comparing the mapping efficiency across different assays (DNA-seq, Hi-C, ChIP-seq, and RNA-seq) between mapping to haplotypes and the reference genome for each individual. For the mapping of raw reads, stringent filtering criteria were applied (2 mismatches, Qm = 255 and Q > 30). Improvement was calculated as ((Haplotype1 U Haplotype 2)-Reference)/Reference. **(C)** The change in improvement as a function of alignment score threshold for Hi-C data. More stringent criteria yields more improvement. However, in the alignment score thresholds that are commonly used in pipelines (e.g., 80-100 for a 101 bp read length), we observe improvements from 1.5% to 4%. **(D)**–**(G)** Comparison of gene expression mapped to a personal genome vs. the reference genome. **(D)** Scatterplots reporting gene expression quantifications obtained after mapping to the reference genome (x axis) and the diploid genomes (y axis) for each of the four individuals. Expression values are reported as log10(TPM + 0.001), and correspond to the median value across tissues. The gene density is color-coded. **(E)** Differential gene expression analysis between quantifications obtained after mapping to the reference genome and to the diploid genomes. Differential expression analysis was performed with DESeq2 (Love et al., 2014) for each donor on RNA-seq read counts across multiple tissues. Genes with an adjusted p-value (Benjamini–Hochberg) < 0.1 and |log2 FC| > 1 were considered to be differentially expressed. See File: Supp_DE_genes.tsv for a full list of differentially expressed genes. The barplot depicts the gene type for genes upregulated when mapping to the diploid genome (left bar, n genes = 107) or to the reference genome (right bar, n genes = 112). **(F)–(G)** Examples of genes upregulated when mapping to the diploid genome (ind 3). HLA-DQA1 belongs to the HLA class II alpha-chain paralogues. SMN2 belongs to the SMN complex and plays a role in pre-mRNA splicing. This gene is part of an inverted duplication on chromosome 5q13, a region prone to rearrangements and deletions. Mutations in this gene have been associated with spinal muscular atrophy (Muller-Felber et al., 2020; Son et al., 2019). SIK1 encodes a member of the salt-inducible kinase family, which has been associated with pigment gene expression (Mujahid et al., 2017). Mutations in this gene have been associated with neurodevelopmental impairments (Hansen et al., 2015; Proschel et al., 2017).

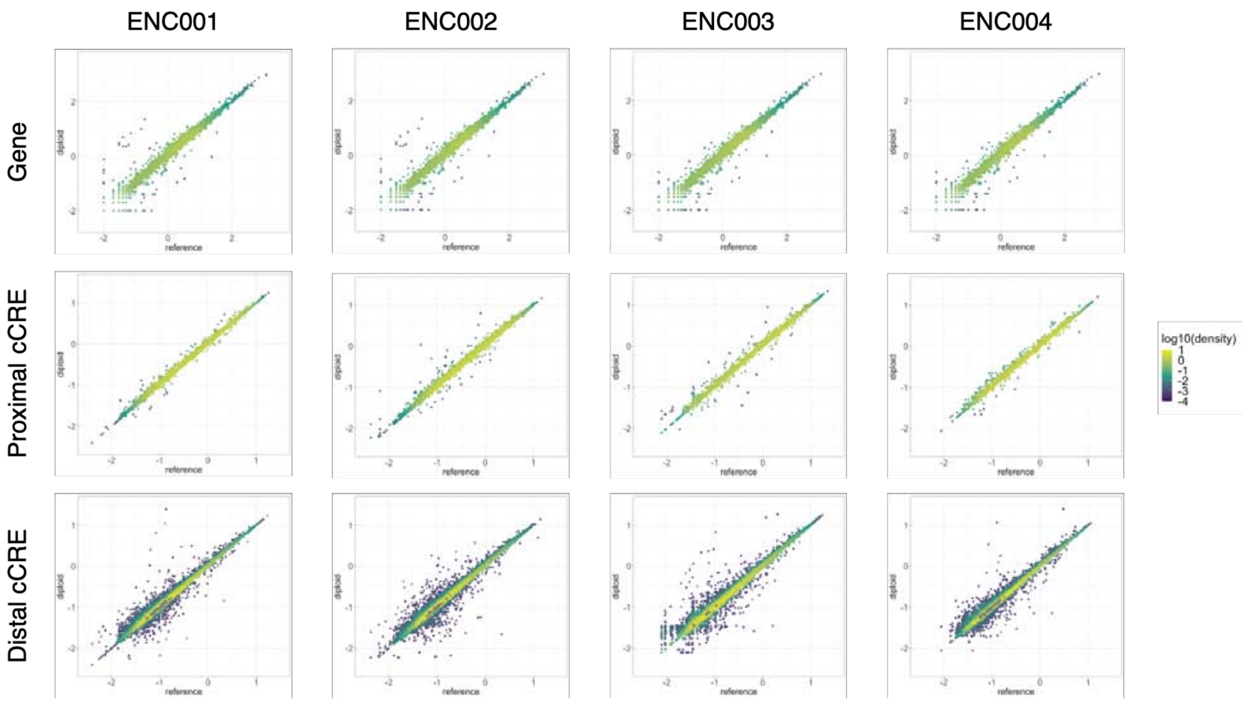

**Figure S1.3b. Gene expression and cCRE activity obtained from mapping to personal genomes**

Scatterplots reporting gene expression, proximal cCRE H3K27ac activity, and distal cCRE H3K27ac activity quantifications obtained after mapping to the reference genome (x axis) and the diploid genomes (y axis) for each of the four individuals. Gene expression and H3K27ac activity values are reported as log10(TPM + 0.001), and correspond to the median value across tissues. The gene and cCRE densities are color-coded. Note that these plots only include protein-coding genes (i.e., no ncRNAs).

#### Autochromosomes + Sex chromosomes

| Gene | up_in_reference | up_in_diploid | total |
|---|---|---|---|
| protein_coding | 44 | 15 | 59 |
| lncRNA | 17 | 6 | 23 |
| pseudogene | 51 | 86 | 137 |
| total | 112 | 107 | **219** |

| cCRE enc001 | up_in_reference | up_in_diploid | total |
|---|---|---|---|
| proximal cCRE | 25 | 110 | 135 |
| distal cCRE | 156 | 846 | 1002 |
| total | 181 | 956 | **1137** |

| cCRE enc002 | up_in_reference | up_in_diploid | total |
|---|---|---|---|
| proximal cCRE | 68 | 116 | 184 |
| distal cCRE | 470 | 878 | 1348 |
| total | 538 | 994 | **1532** |

| cCRE enc003 | up_in_reference | up_in_diploid | total |
|---|---|---|---|
| proximal cCRE | 76 | 9 | 85 |
| distal cCRE | 559 | 90 | 649 |
| total | 635 | 99 | **734** |

| cCRE enc004 | up_in_reference | up_in_diploid | total |
|---|---|---|---|
| proximal cCRE | 37 | 3 | 40 |
| distal cCRE | 219 | 59 | 278 |
| total | 256 | 62 | **318** |

#### Autochromosomes

| Gene | up_in_reference | up_in_diploid | total |
|---|---|---|---|
| protein_coding | 39 | 15 | 54 |
| lncRNA | 17 | 6 | 23 |
| pseudogene | 46 | 81 | 127 |
| total | 102 | 102 | **204** |

| cCRE enc001 | up_in_reference | up_in_diploid | total |
|---|---|---|---|
| proximal cCRE | 25 | 3 | 28 |
| distal cCRE | 156 | 66 | 222 |
| total | 181 | 69 | **250** |

| cCRE enc002 | up_in_reference | up_in_diploid | total |
|---|---|---|---|
| proximal cCRE | 68 | 10 | 78 |
| distal cCRE | 468 | 106 | 574 |
| total | 536 | 116 | **652** |

| cCRE enc003 | up_in_reference | up_in_diploid | total |
|---|---|---|---|
| proximal cCRE | 68 | 9 | 77 |
| distal cCRE | 543 | 89 | 632 |
| total | 611 | 98 | **709** |

| cCRE enc004 | up_in_reference | up_in_diploid | total |
|---|---|---|---|
| proximal cCRE | 37 | 3 | 78 |
| distal cCRE | 216 | 56 | 574 |
| total | 536 | 116 | **312** |

**Figure S1.3c. Differential gene expression and cCRE activity between mapping to reference and personal genomes**

Differential gene expression and cCRE H3K27ac activity analysis between quantifications obtained after mapping to the reference genome and to the diploid genomes. Differential expression analysis was performed with DESeq2 (Love *et al.*, 2014) for each donor on RNA-seq/H3K27ac ChIP-seq read counts across multiple tissues. Genes/cCREs with an adjusted p-value (Benjamini–Hochberg) $< 0.1$ and $|\log2 FC| > 1$ were considered to be differentially expressed. The left and right panel shows the numbers of differentially expressed genes and cCREs on the all chromosomes and autochromosomes, respectively.

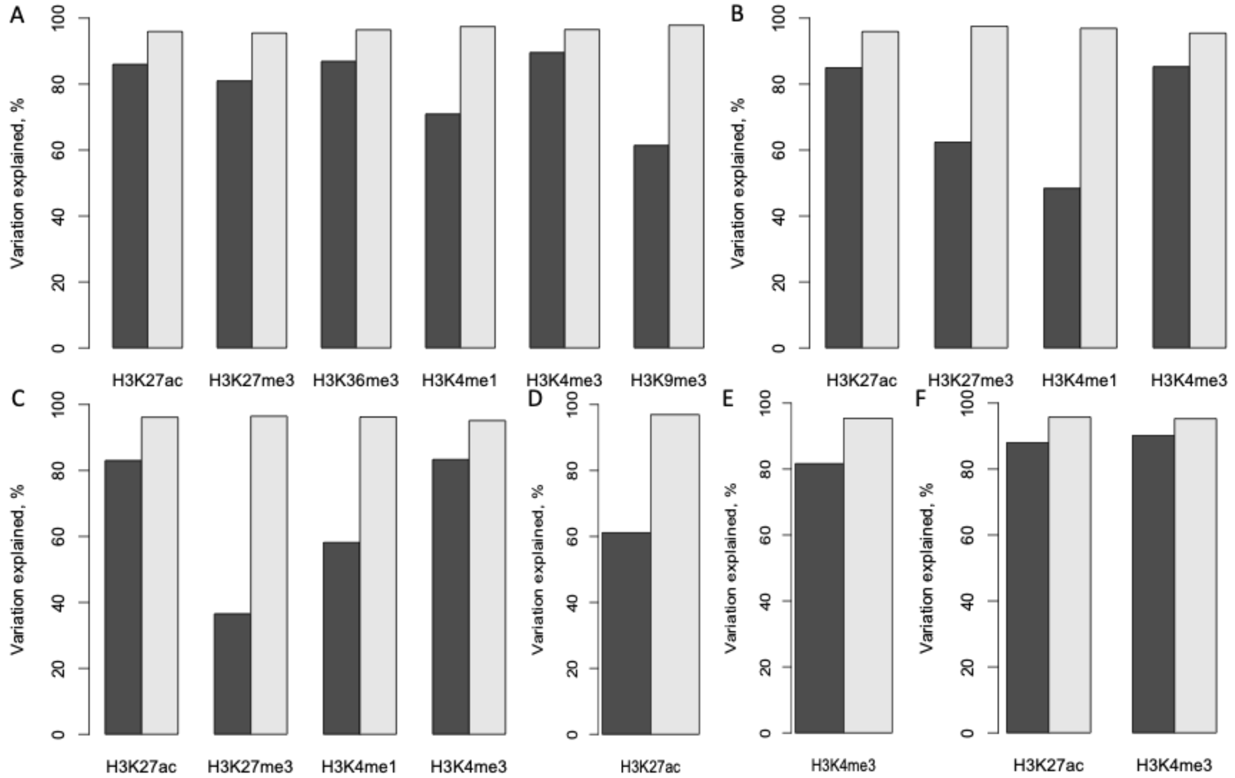

**Figure S1.4a. Variation explained between two experiments corrected by replicates**
To calculate the variation explained between experiments (e.g., the two H3K27ac ChIP-seq experiments of the spleens from two individuals), for each experiment, we identified the cCREs that drive high variation explained (> 95%) in the replicates of the experiment. The intersecting set of cCREs from the two experiments was used to calculate the variation explained between the two experiments (black bars; e.g., 87% for the two H3K27ac experiments). The average variation explained between the replicates from the two experiments is indicated by the white bars (e.g., 96% for H3K27ac). The results in spleen, transverse colon, gastrocnemius medialis, thyroid gland, pancreas, and prostate gland are shown in **(A)–(F)**.

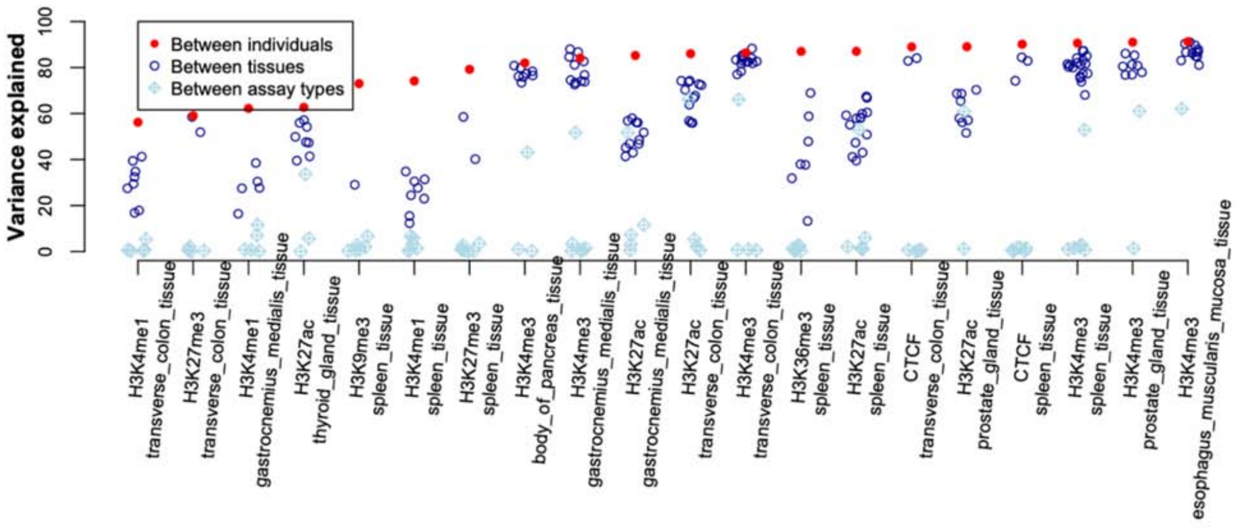

| Similarity of functional genomic activities of cCREs | | | | |
|---|---|---|---|---|
| % of variation explained | H3K27ac: adrenal_gland_tissue: male_adult_54_years | H3K27ac: body_of_pancreas_tissue: male_adult_37_years | H3K27ac: esophagus_muscularis_mucosa_tissue: female_adult_51_years | H3K27ac: gastrocnemius_medialis_tissue: female_adult_53_years |
| H3K27ac: adrenal_gland_tissue: male_adult_54_years | 95.63 | 64.05 | 64.17 | 52.16 |
| H3K27ac: body_of_pancreas_tissue: male_adult_37_years | 64.05 | 95.1 | 56.51 | 44.09 |
| H3K27ac: esophagus_muscularis_mucosa_tissue: female_adult_51_years | 64.17 | 56.51 | 95.35 | 56.47 |

**Figure S1.4b. Similarity between the signals of two functional genomic experiments**
For each cCRE, the signal of a functional genomic experiment was measured by the average fold-change over control across the cCRE region. For two experiments, linear regression was used for the cCREs with low technical noise between replicates. The variance of one experiment explained by the other is used to indicate the similarity between the experiments across the cCREs. The similarity between all possible pairs of experiments is reported in the table shown beneath the figure.

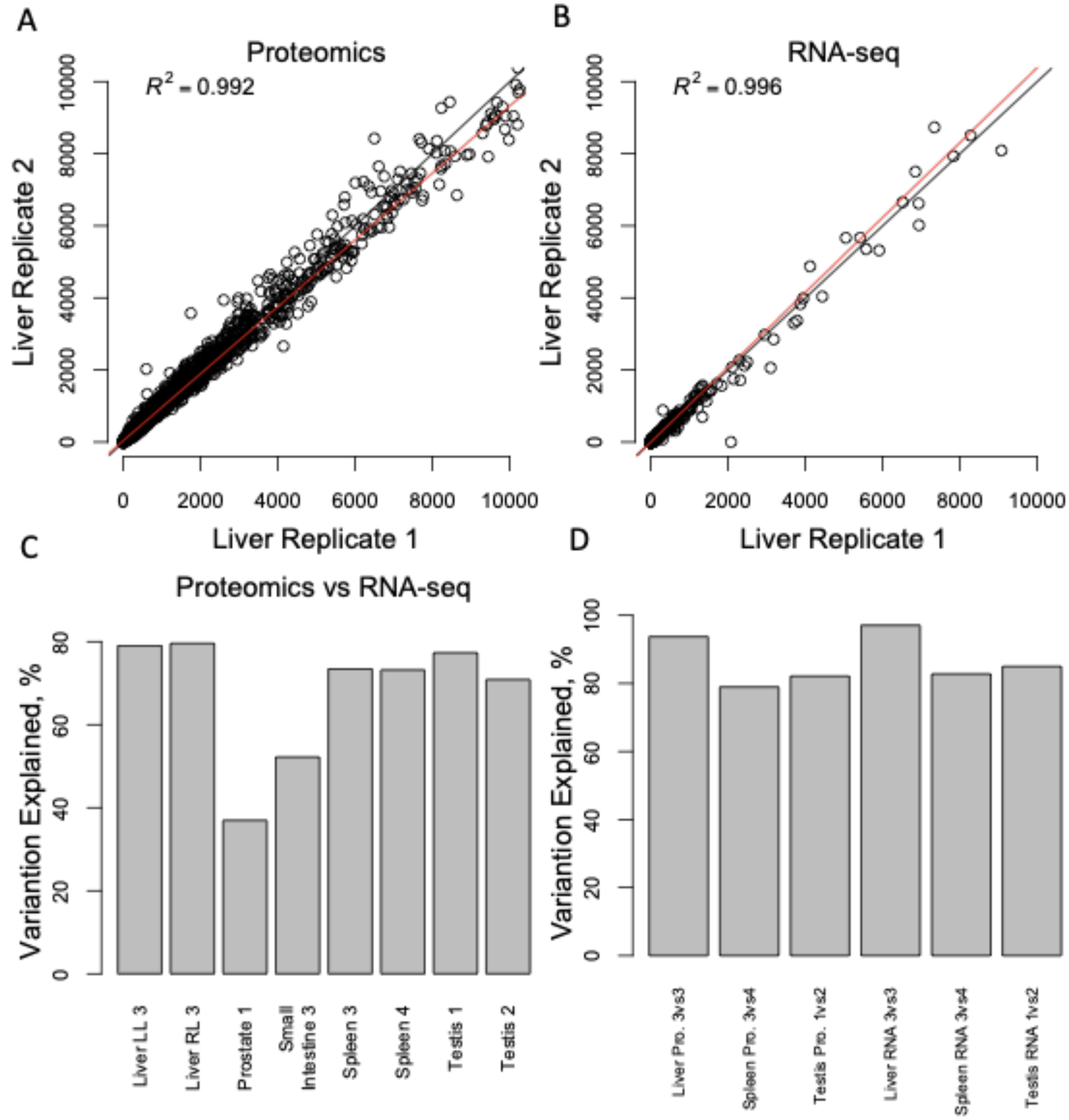

**Figure S1.4c. Variation explained between proteomics and RNA-seq data**
**(A)** The normalized protein abundances are highly consistent between replicates. **(B)** This is also true for the normalized RNA abundances. **(C)** The variation explained between the normalized protein abundances and the normalized RNA abundances varies across tissues, suggesting that for some tissues, protein abundances and RNA abundances have low consistency. LL indicates the left lobe of the liver, and RL indicates the right lobe. The numbers in the labels indicate the donors. However, respectively for protein abundances and RNA abundances, the variation explained between donors is higher **(D)** than the variation explained between the normalized protein abundances and RNA abundances **(C)**. The normalized proteomics and RNA-seq data matrix used for panels **(C)** & **(D)** is in File: normalized_proteomics_RNA-seq.dat.

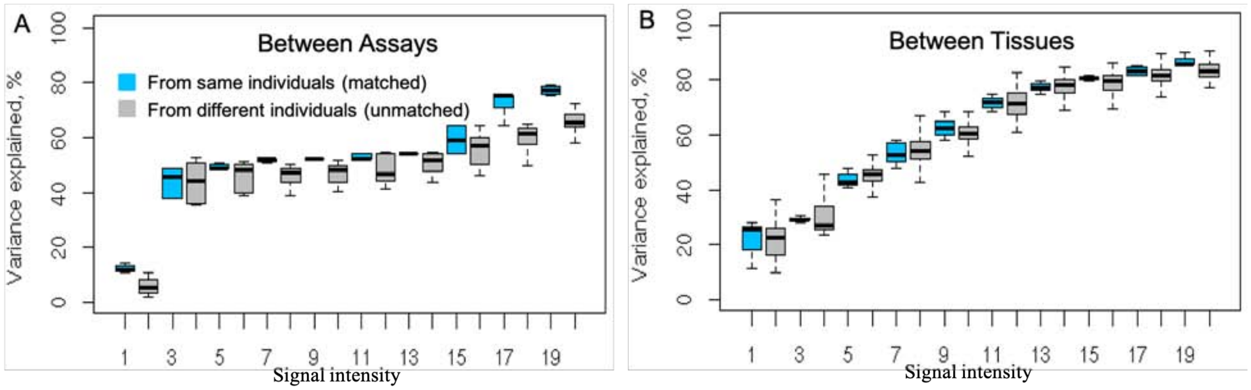

**Figure S1.4d. Comparing the explained variances calculated respectively using matched and unmatched data**

**(A)** The explained variance is calculated between two types of histone modifications (between assays) using matched data (blue), i.e., the two assays are generated from the same tissue of an individual. The unmatched data (grey) indicate the two assays are generated from the same tissues of different individuals. **(B)** The explained variance is calculated between the same histone modifications generated from two different tissues (between tissues). The tissues that are from the same individuals are referred to as matched, and are otherwise unmatched.

#### Figure S2. Supp. Figures for Main Text Section “Large-scale Determination of AS SNVs & Construction of the AS Catalog”

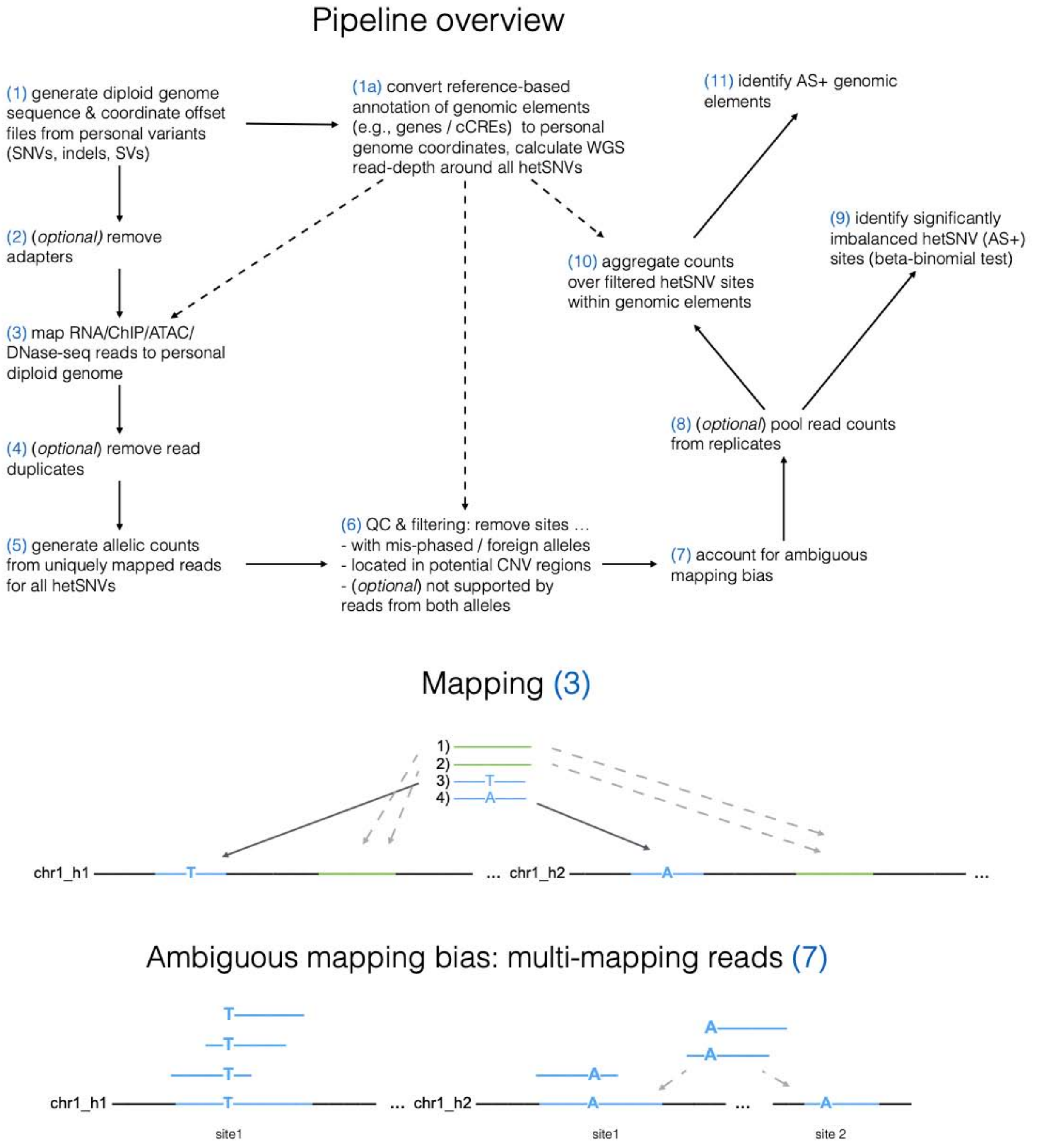

**Figure S2.1a. Workflow of the AlleleSeq2 pipeline**

We used genomic variants to construct the sequence and coordinates for each haplotype, and then mapped functional genomics assay reads to each haplotype by using the phased hetSNVs in a read (middle panel). This generated BAM files that contain both reads that are mapped uniquely to a region in a haplotype and reads that are mapped to multiple regions (within a haplotype or between two haplotypes, bottom panel). Since it is not possible to unambiguously identify the origin of the reads that multi-map within the haplotypes, we made the conservative assumption that all of these reads originate from the heterozygous locus and unless the direction of the bias changed towards the opposite allele, we adjusted the allele counts including the multi-mapping reads. We pooled all of the reads for assays with replicates. We identified hetSNVs with allelic imbalance by performing a beta-binomial test on the allelic reads. To determine whether a genomic region has allelic imbalance in RNA-seq, ChIP-seq, or ATAC-seq, we summed up allele-specific reads from all hetSNVs within the region and performed a beta-binomial test.

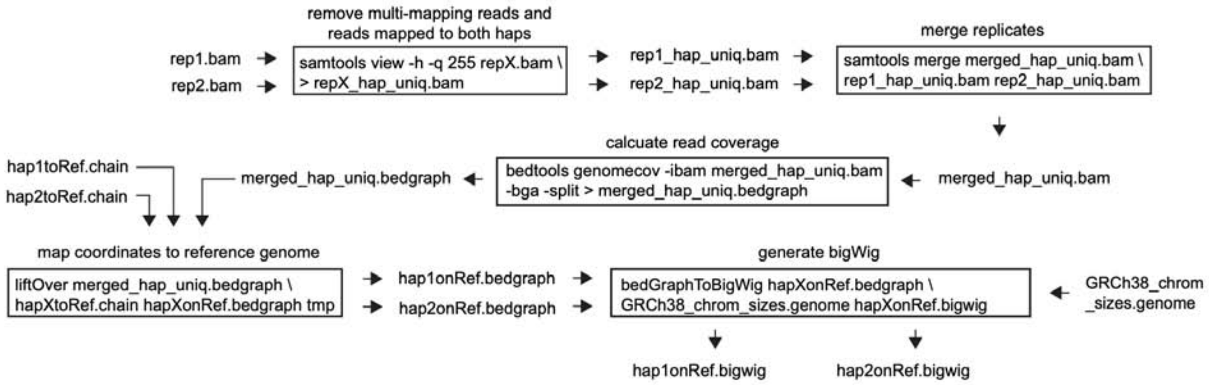

**Figure S2.1b. Generating haplotype-specific signal tracks**
Each box is a command to process files. The input BAM files are generated by step (3) in Figure S2.1a, containing reads that are uniquely mapped to each haplotype and reads with ambiguous mapping. The first box removes the reads with ambiguous mapping. In this example, the assay has two replicates so we merge the two BAM files of unique reads (box 2). If there are no replicates, then "merge replicates" is skipped. The read coverage of each chromosome in each haplotype is then calculated and stored in bedGraph files (box 3). Note that the coordinates of a given genomic region are in the personal genome, therefore the two haplotypes can give different coordinates even for the same gene. To compare the same region between the two haplotypes, we convert the coordinates in each haplotype from the personal genome to the reference genome (box 4). We also convert the read coverage from bedGraph to bigWig, which can be plotted in the IGV genome viewer (box 5). A script that generates the haplotype-specific read coverage from BAM files is provided in https://github.com/gersteinlab/AlleleSeq2. An example of the intermediate files (except for BAM files) in generating haplotype-specific signal tracks is available from File: sample_signal_track.tar.gz.

| | |
|---|---|
| Fig.3B | RNA: ENCFF660SLV, ENCFF751QEC<br>CTCF: ENCFF296YDQ, ENCFF255INZ<br>H3K27ac: ENCFF184LPK, ENCFF789APL, ENCFF707VEV, ENCFF298AKE |
| Fig.3C<br>Fig.S2.8d | ind3 RNA: ENCFF281PBY, ENCFF760KXM<br>ind3 H3K27ac: ENCFF699EFW, ENCFF075RQB<br>ind3 H3K27me3: ENCFF888EIC, ENCFF595OTK, ENCFF011LXD, ENCFF626DTV<br>ind4 RNA: ENCFF711JSM, ENCFF355UJC, ENCFF415QZI, ENCFF912BRJ<br>ind4 H3K27ac: ENCFF089KJG, ENCFF908MFI, ENCFF912LCB, ENCFF431PJB<br>ind4 H3K27me3: ENCFF417VAA, ENCFF876PUF, ENCFF992SRG, ENCFF463QBH |
| Fig.4D,<br>Fig.S3.2dA | RNA: ENCFF719MSG, ENCFF120MML, ENCFF337ZBN, ENCFF481IQE<br>H3K27ac: ENCFF339ODV, ENCFF870TZH<br>TF binding clusters: UCSC encRegTfbsClustered |
| Fig.4E,<br>Fig.S3.2eA | RNA: ENCFF038JEE, ENCFF897TAN<br>H3K27ac: ENCFF143SOY, ENCFF244ISL, ENCFF804MSF, ENCFF976BRQ<br>TF binding clusters: UCSC encRegTfbsClustered |
| Fig.4G,<br>Fig.S3.2fA | ind3 RNA: ENCFF534JLO<br>ind3 H3K27ac: ENCFF066DSD<br>ind3 CTCF: ENCFF417IMY<br>ind2 RNA: ENCFF232DNA<br>ind2 H3K27ac: ENCFF439NXI<br>ind2 CTCF: ENCFF178GEC<br>TF binding clusters: UCSC encRegTfbsClustered |
| Fig.4G.<br>Fig.S3.2g<br>Fig.S3.2h | ind3 RNA: ENCFF216VOH<br>ind3 H3K9me3: ENCFF423DVX<br>ind3 long-read RNA: ENCFF185VYD<br>ind2 RNA: ENCFF187KAR<br>ind2 H3K27ac: ENCFF095CZX<br>ind2 long-read RNA: ENCFF912HPY |
| Fig.S3.2c | RNA: ENCFF326CGI, ENCFF663VCC<br>H3K27ac: ENCFF935UTO, ENCFF653PKW, ENCFF235IVE, ENCFF226YFN<br>ATAC: ENCFF591BAY, ENCFF332SCG<br>CTCF: ENCFF800GHL, ENCFF100YUK, ENCFF861WPS, ENCFF056JNV, ENCFF608GCT, ENCFF682AOT<br>TF binding clusters: UCSC encRegTfbsClustered |
| Fig.S3.2eB | RNA: ENCFF122HNW, ENCFF069KBE, ENCFF483NBR, ENCFF226NNE<br>H3K27ac: ENCFF459LBY, ENCFF949SUD, ENCFF481TGO, ENCFF359AHW, ENCFF252NKY, ENCFF920PYS, ENCFF384MQH, ENCFF270YMP, ENCFF605JUU, ENCFF264CZV, ENCFF867PQG, ENCFF003LQT, ENCFF219DYV, ENCFF113KFQ<br>TF binding clusters: UCSC encRegTfbsClustered |
| Fig.S3.2eC | RNA: ENCFF411WXY, ENCFF543BVT, ENCFF072VKD, ENCFF484BLA, ENCFF086TFZ, ENCFF351OAS<br>H3K27ac: ENCFF214DHU, ENCFF209OKJ, ENCFF330KKH, ENCFF343NQH, ENCFF706KXN, ENCFF349JBL, ENCFF945XBP, ENCFF382QHO, ENCFF922CDY, ENCFF778KZF, ENCFF040XEO, ENCFF173QJF, ENCFF033YTT, ENCFF088QFN |
| Fig.S3.2eD | ind1 RNA: ENCFF751LMQ, ENCFF188LMP, ENCFF151GUG, ENCFF628TMU<br>ind1 H3K27ac: ENCFF259FFL, ENCFF570KZO, ENCFF176GZS<br>ind4 RNA: ENCFF711JSM, ENCFF355UJC, ENCFF415QZI, ENCFF912BRJ<br>ind4 H3K27ac: ENCFF930NCY, ENCFF089KJG, ENCFF908MFI, ENCFF912LCB, ENCFF431PJB |
| Fig.S3.2i | ind1 RNA: ENCFF150OQH, ENCFF460GHU, ENCFF661PVB, ENCFF152GUK<br>ind2 RNA: ENCFF900BSQ, ENCFF184XUR<br>ind3 RNA: ENCFF667RSP, ENCFF891NXQ |
| Fig.S3.2jA | ind1 RNA: ENCFF060XKP<br>ind1 H3K27ac: ENCFF665GNN<br>ind2 RNA: ENCFF251QRT<br>ind2 H3K27ac: ENCFF439ASM |

**Figure S2.1c. Data used to generate signal tracks**

Data in blue are given as the accession numbers in the ENCODE portal. TF binding clusters are available via the UCSC table browser.

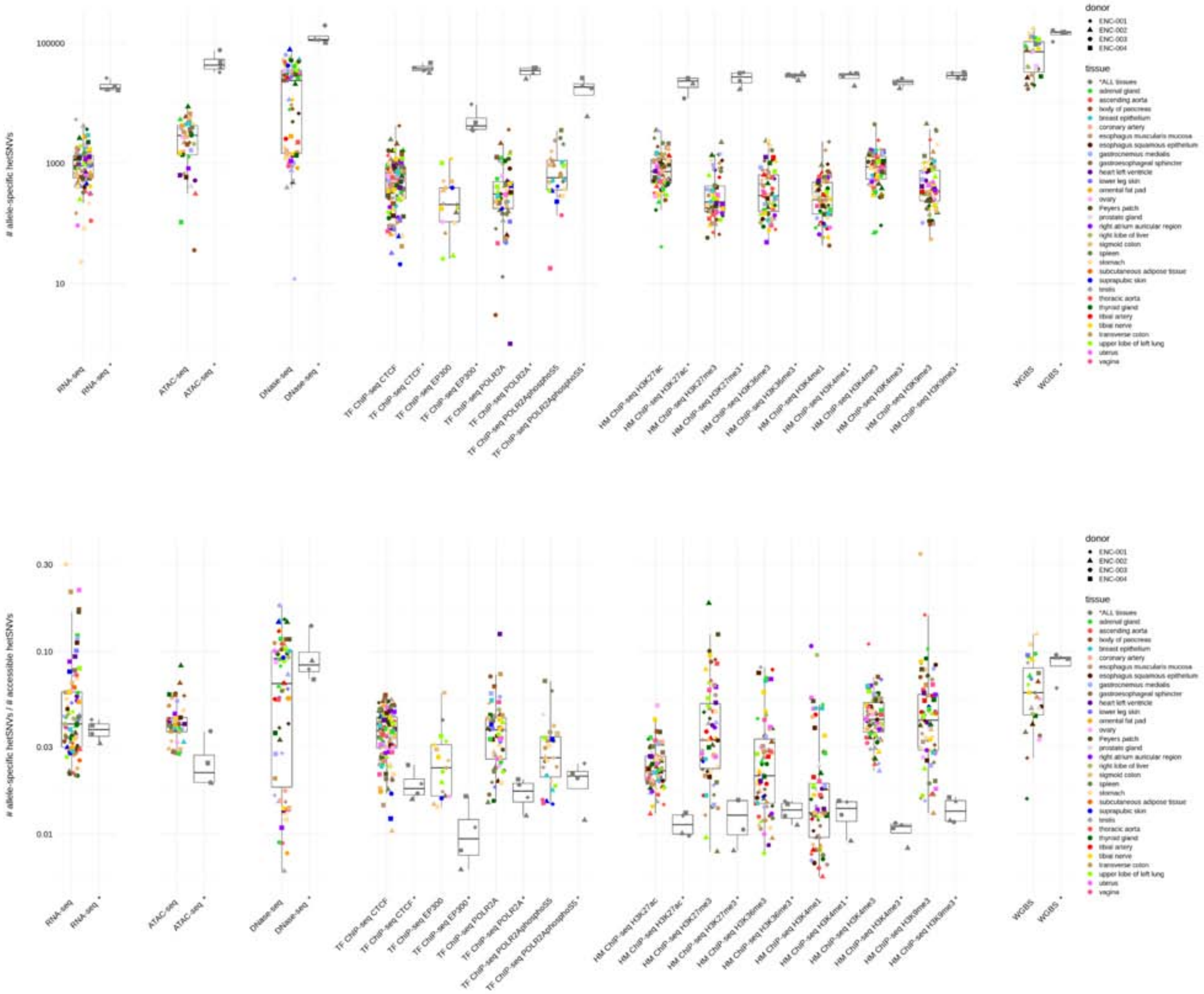

**Figure S2.1d. Distribution and fraction of the number of hetSNVs associated with AS behavior across different EN-TEx donors, tissues, and assays**

The fraction is the number of hetSNVs associated with AS behavior relative to the number of accessible hetSNVs. Call sets based on pooled reads from all tissues for each donor and assay are shown in gray. 820 AS events on average were detected in the RNA/ChIP/ATAC-seq samples (median 517, IQR 251-1030; ~3.8 % of the total number of accessible sites).

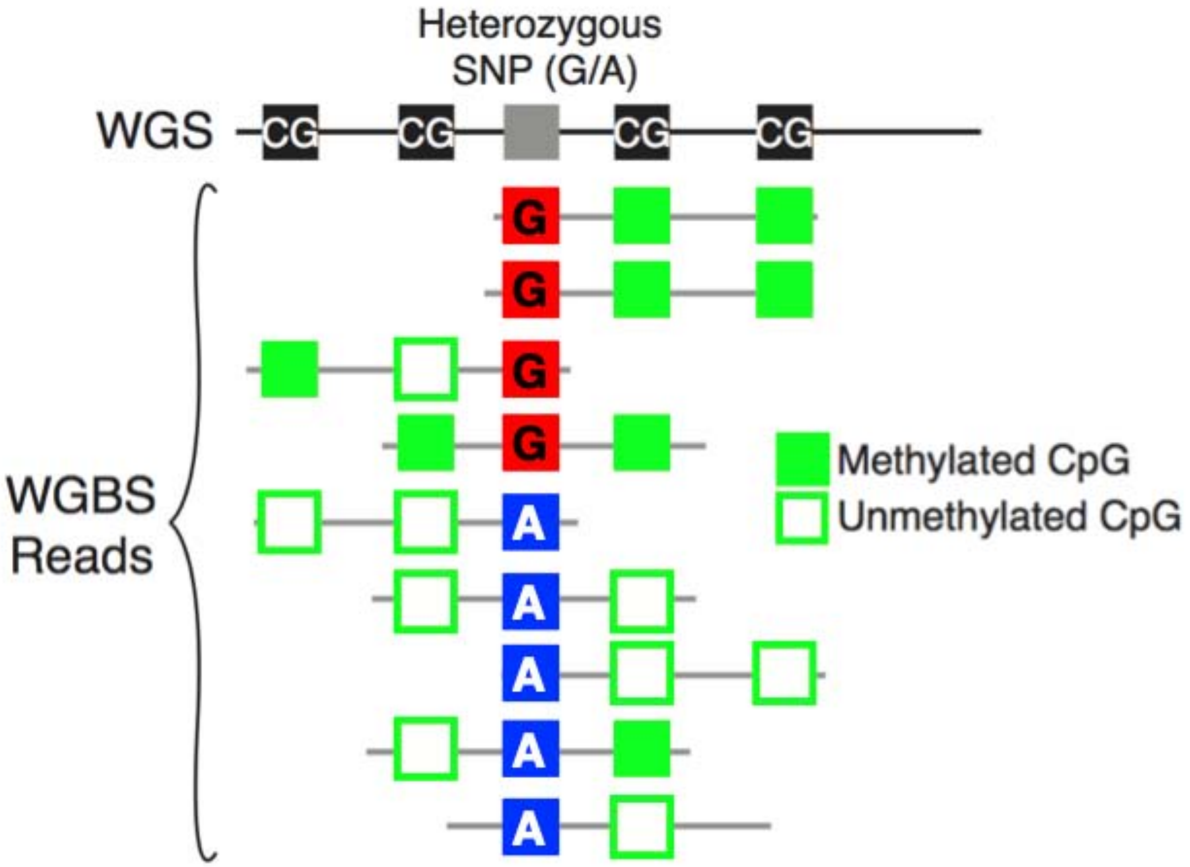

| | G | A |
|---|---|---|
| Methylated CpG | 7 | 1 |
| Unmethylated CpG | 1 | 8 |

→ Fisher's exact test

**Figure S2.1e. Schematic showing how AS methylation is calculated**
We can determine ASM by identifying AS methylated CpG sites near tag hetSNVs using the statistical test above. Since methylated C are sequenced differently from unmethylated C, we use a two-by-two contingency test (Fisher's exact test) in order to identify AS methylated CpGs in the vicinity of a tag hetSNV (Dianzani et al., 1986).

| Methylomes | PSIMCT-1,HM13 | BLCAP;NNAT | L3MBTL1 | GNAS | DDC;GRB10 | SGCE;PEG10 | MESTIT1;MEST | ZNF597;NAA60 | SNRPN;SNURF | NAP1L5 | INS;IGF2;H19 | KCNQ1OT1 | KCNK9 | DIRAS3 | ZDBF2 | ZIM2;PEG3 | DLK1;MEG3 | RB1 | FAM50B | PLAGL1; HYMAI | TCEB3C | INPP5F |
|---|---|---|---|---|---|---|---|---|---|---|---|---|---|---|---|---|---|---|---|---|---|---|
| ENC003 - AG | | 1/1 | | | | 0/1 | 1/1 | 4/4 | 4/4 | 1/1 | | | | | 4/4 | | | 0/1 | | | | |
| ENC003 - ESE | | 1/1 | | | | 0/1 | 2/2 | 1/2 | 3/4 | 0/2 | | | | | 1/3 | | | 0/1 | | | | |
| ENC003 - GS | | 1/1 | | | | 0/1 | | 1/2 | 2/2 | 0/1 | | | | | 0/1 | 0/1 | | 0 | | | | |
| ENC003 - U | | 1/1 | | | | 0/2 | 0/1 | 1/1 | 4/5 | 1/1 | | | 1/1 | | 4/4 | 1/1 | | 0/1 | 0/1 | | | |
| ENC003 - SLL | | 1/1 | | | | 0/1 | 0/1 | 1/1 | 2/3 | 0/1 | | | 0 | | 1/3 | 1/1 | | 0/1 | 0/1 | | | |
| ENC003 - TN | | 1/1 | 2/2 | | | 0/1 | 0/2 | 1/1 | 3/6 | 1/1 | | | 0/1 | | 3/3 | 1/1 | | 1/1 | 0/1 | | | |
| ENC004 - LLS | | | | | | | | | 3/3 | | | 2/2 | 1/2 | | | 0/1 | | 2/3 | | | | |
| ENC004 - O | | | | | | | | | 2/3 | | | 1/2 | | | | 1/3 | | 1/3 | | | | |
| ENC004 - Sp | | | | | | | | | 3/4 | | | 2/2 | 1/2 | | | | | 1/2 | | | | |
| ENC004 - SI | | | | | | | | | 2/3 | | | 1/1 | 1/2 | | | 0/1 | | 1/3 | | | | |
| ENC004 - TG | | | | | | | | | 3/3 | | | | | | | 2/2 | | 1/1 | | | | |
| ENC004 - ULLL | | | | | | | | | 2/2 | | | 1/1 | 1/2 | | | 2/3 | | 1/3 | | | | |
| ENC001 - LLS | | | | 4/4 | | | 1/1 | | | 1/2 | | 0/1 | 1/2 | 1/1 | 3/4 | 4/4 | 0/1 | 2/3 | | | | 0/1 |
| ENC001 - PR | | | | 3/4 | | | 1/1 | | | 1/1 | | 1/3 | 1/1 | 0/1 | 1/3 | 2/3 | | 0/2 | | | | 0/1 |
| ENC001 - Sp | | | | 4/4 | | | 0/1 | | | 1/2 | | 2/2 | 1/1 | 1/1 | 3/3 | 2/2 | | 1/1 | | | | 0/1 |
| ENC001 - T | | | | 4/4 | | | 1/1 | | | 1/2 | | 2/2 | 1/2 | 1/1 | 3/4 | 3/4 | 1/1 | 0/2 | | | | 1/1 |
| ENC001 - TG | | | | 3/4 | | | 1/1 | | | 1/2 | | 0/2 | 1/2 | 1/1 | 4/4 | 2/3 | | 0/3 | | | | 1/1 |
| ENC001 - ULLL | | | | 5/5 | | | 1/1 | | | 1/2 | | 1/3 | 1/2 | 1/1 | 3/5 | 2/3 | | 0/1 | | | | 1/1 |
| ENC002 - AG | | 1/3 | | | | | 3/4 | 4/4 | 2/4 | | | | | | 3/3 | 2/2 | | | 0/1 | 1/1 | | |
| ENC002 - ESE | | 1/1 | | | | | 1/2 | 0/1 | 2/4 | | | | | | 3/3 | 2/3 | | | 1/1 | 1/1 | | |
| ENC002 - GS | | 0/2 | | | | | 0/2 | 0/2 | 1/1 | | | | | | 2/2 | 1/1 | | | 1/1 | | | |
| ENC002 - PR | | 0/3 | | | | 1/1 | 2/3 | 4/4 | 2/3 | | | | | | 2/3 | 1/2 | | | 1/1 | 1/1 | | |
| ENC002 - PG | | 0/1 | | | | | | | 2/2 | | | | | | 3/3 | 3/5 | 1/3 | | | | | |
| ENC002 - SI | | 1/3 | | | | 1/1 | 2/4 | 2/3 | 1/5 | | | | | | 3/4 | 1/3 | 0/2 | | 1/1 | 1/1 | | |
| ENC002 - T | | 0/2 | | | | 0/1 | 2/3 | 2/3 | 3/4 | | | | | | 1/3 | 1/2 | | | 0/1 | 1/1 | | |
| ENC002 - TN | | 1/2 | | | | | 3/4 | 2/3 | 5/5 | | | | | | 3/3 | 5/5 | | | 1/1 | 1/1 | | |
| ENC002 - TC | | 0/1 | | | | | 0/3 | 0/1 | | | | | | | 0/2 | 0/1 | | | | 1/1 | | |

Region not covered | ASM in region | No ASM in region

**Figure S2.1f. ASM calls in known imprinting control regions**

Number of ASM calls made out of the total number of accessible hetSNVs that overlap an imprinting control region (ICR)(Fang et al., 2012) for each sample. Green cells represent ICRs overlapping at least one ASM call in that sample. Red cells represent ICRs that overlap with at least one accessible hetSNV but no hetSNVs with significant imbalance are observed. Yellow cells represent ICRs that did not overlap with any accessible hetSNVs in the sample.

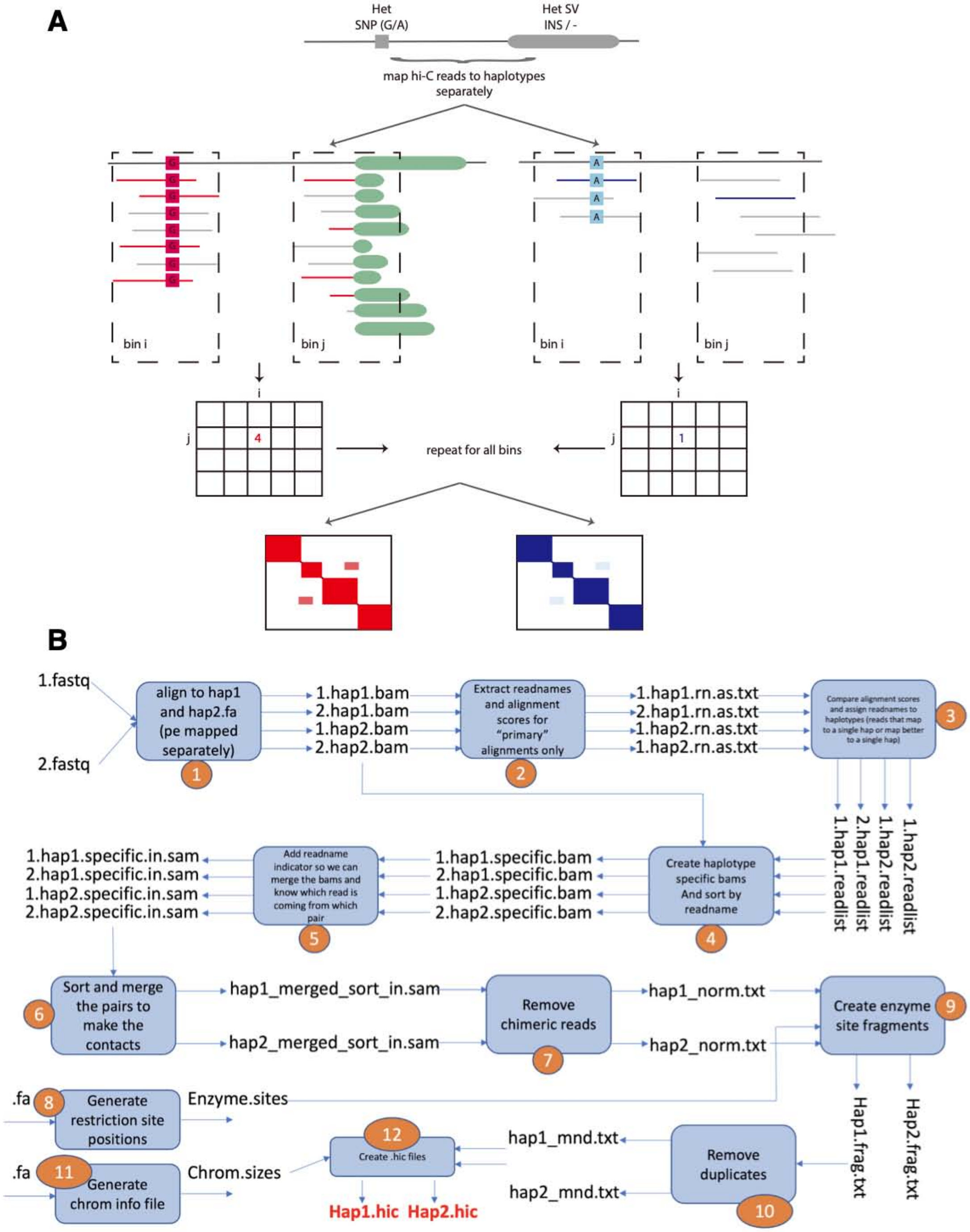

**Figure S2.1g. Generating haplotype-specific Hi-C**
**(A)** Schematic showing the overall methodology for determining haplotype-specific 3D contact interactions using Hi-C paired-end reads. **(B)** Workflow for the generation of haplotype-specific Hi-C contact maps.

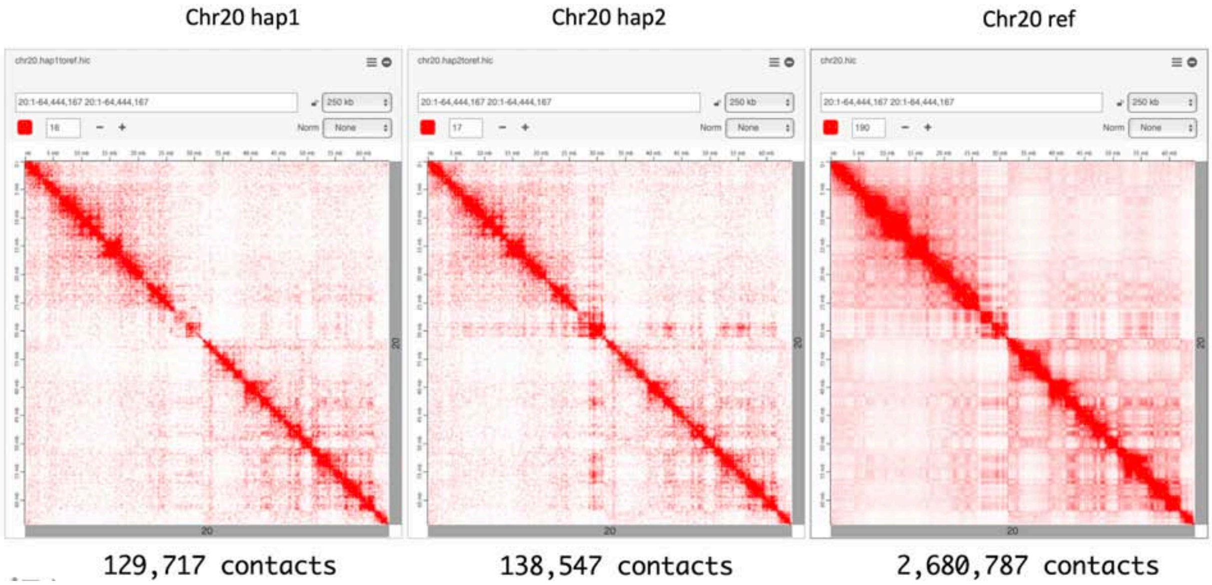

**Figure S2.1h. Haplotype-specific contact maps for Chr20 generated using the personal genome coordinates**

The third map is the bulk Hi-C contact map of Chr20 generated using the reference genome.

| Individual/Tissue | Intra-chromosomal Interactions | Hap1 | Hap2 | Hap1 or Hap2 | Significantly Imbalanced |
|---|---|---|---|---|---|
| ind1 skeletal muscle | 39,013,901 | 4,049,203 | 4,034,602 | 7,041,417 | 577,728 |
| ind2 skeletal muscle | 4,405,480 | 1,117,328 | 1,146,381 | 2,072,227 | 140,317 |
| ind3 skeletal muscle | 40,412,585 | 4,345,533 | 4,359,297 | 7,493,069 | 574,836 |
| ind4 skeletal muscle | 41,569,344 | 4,028,293 | 4,021,800 | 6,983,660 | 523,931 |
| ind1 transverse colon | 45,534,793 | 4,942,660 | 4,924,000 | 8,574,917 | 702,953 |
| ind2 transverse colon | 25,548,308 | 2,148,267 | 2,151,803 | 3,842,621 | 261,752 |
| ind3 transverse colon | 43,917,995 | 4,716,549 | 4,722,227 | 8,118,858 | 609,973 |
| ind4 transverse colon | 43,406,680 | 4,343,051 | 4,334,617 | 7,506,125 | 583,468 |

**Figure S2.1i. Number of Hi-C contacts obtained from haplotype specific Hi-C contact maps**

Of the average 6,454,111 interactions per sample, 496,859 showed significant AS behavior.

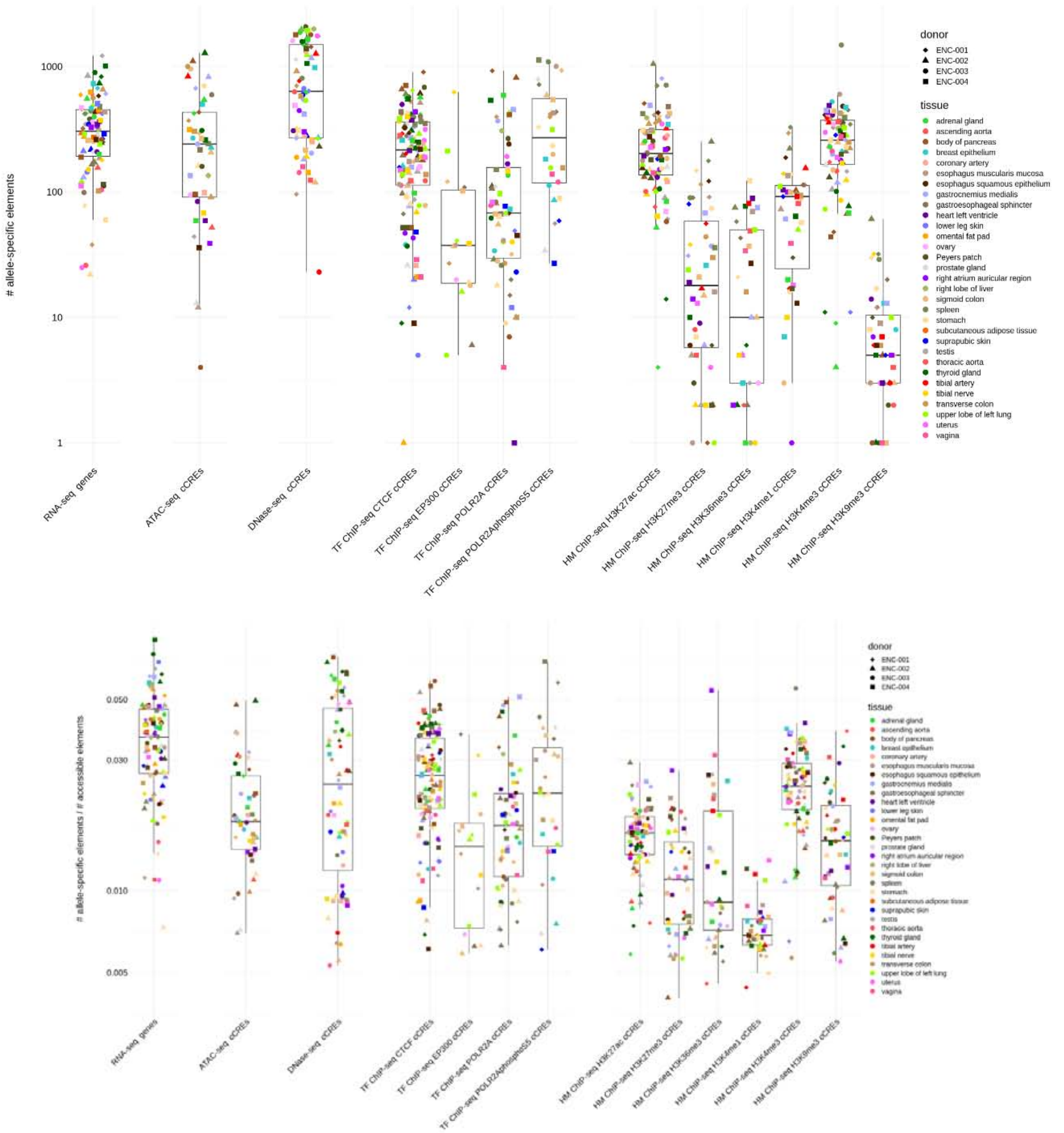

**Figure S2.2a. Distribution and fractions of the number of genomic elements (genes and cCREs) associated with AS behavior across different EN-TEx donors, tissues, and assays**

On average, for each individual and tissue in the RNA/ChIP/ATAC-seq samples, 193 cCREs (median 123, IQR 32-284) and 351 genes (median 205, IQR 193-452) showed a significant AS imbalance per assay.

| Type | Assay | Category | Term | Total AS+protein-coding genes | Category gene count | Percentage (%) | P-Value | FDR |
|---|---|---|---|---|---|---|---|---|
| ASB+ | H3K27ac | UP_KEYWORDS | **Phosphoprotein** | 2,115 | **1,011** | **47.8** | 7.0E-07 | 3.9E-04 |
| | | UP_KEYWORDS | Acetylation | | 433 | 20.5 | 1.9E-04 | 5.3E-02 |
| | H3K4me3 | UP_KEYWORDS | **Phosphoprotein** | 2,312 | **1,080** | **46.7** | 1.0E-06 | 5.5E-04 |
| | | GOTERM_CC_DIRECT | Golgi membrane | | 106 | 4.6 | 4.5E-05 | 3.7E-02 |
| | H3K27me3 | INTERPRO | Immunoglobulin-like fold | 335 | 33 | 9.9 | 2.3E-05 | 1.5E-02 |
| | | UP_KEYWORDS | Developmental protein | | 38 | 11.3 | 7.7E-05 | 2.4E-02 |
| | | KEGG_PATHWAY | Systemic lupus erythematosus | | 10 | 3 | 3.2E-04 | 4.7E-02 |
| | | KEGG_PATHWAY | Alcoholism | | 11 | 3.3 | 5.2E-04 | 4.7E-02 |
| | CTCF | UP_KEYWORDS | Ubl conjugation | 1,227 | 144 | 11.7 | 1.5E-05 | 7.4E-03 |
| | | UP_KEYWORDS | Isopeptide bond | | 97 | 7.9 | 1.3E-04 | 3.1E-02 |
| | | UP_KEYWORDS | **Phosphoprotein** | | **574** | **46.8** | 1.9E-04 | 3.1E-02 |
| | | UP_KEYWORDS | Acetylation | | 253 | 20.6 | 4.2E-04 | 5.1E-02 |
| | H3K36me3 | No | | | | | | |
| | H3K9me3 | | | | | | | |
| | H3K4me1 | | | | | | | |
| | ATAC | | | | | | | |
| | DNase | | | | | | | |
| ASE+ (top 3,000) | RNA-seq | UP_KEYWORDS | Glycoprotein | 2,966 | 902 | 30.4 | 3.8E-35 | 2.0E-32 |
| | | UP_SEQ_FEATURE | glycosylation site:N-linked (GlcNAc...) | | 835 | 28.2 | 6.0E-31 | 4.2E-27 |
| | | UP_SEQ_FEATURE | signal peptide | | 699 | 23.6 | 1.9E-29 | 6.6E-26 |
| | | UP_KEYWORDS | Disulfide bond | | 638 | 21.5 | 2.7E-26 | 7.2E-24 |
| | | UP_KEYWORDS | Signal | | 782 | 26.4 | 5.7E-26 | 1.0E-23 |
| | | UP_KEYWORDS | Secreted | | 405 | 13.7 | 6.1E-22 | 8.2E-20 |
| | | GOTERM_CC_DIRECT | extracellular exosome | | 629 | 21.2 | 1.1E-22 | 9.0E-20 |
| | | UP_KEYWORDS | **Polymorphism** | | **2,071** | **69.8** | 2.3E-21 | 2.5E-19 |
| | | UP_SEQ_FEATURE | disulfide bond | | 533 | 18.0 | 6.9E-21 | 1.6E-17 |
| | | UP_SEQ_FEATURE | **sequence variant** | | **2,089** | **70.4** | 3.6E-20 | 6.1E-17 |
| | | GOTERM_CC_DIRECT | plasma membrane | | 760 | 25.6 | 1.0E-17 | 4.3E-15 |
| | | UP_KEYWORDS | Membrane | | 1,289 | 43.5 | 3.3E-16 | 3.0E-14 |
| | | INTERPRO | Epidermal growth factor-like domain | | 94 | 3.2 | 1.3E-17 | 4.1E-14 |
| | | GOTERM_CC_DIRECT | extracellular region | | 321 | 10.8 | 2.2E-16 | 6.0E-14 |
| | | SMART | EGF | | 82 | 2.8 | 8.5E-16 | 4.5E-13 |
| | | UP_KEYWORDS | EGF-like domain | | 89 | 3.0 | 1.0E-14 | 7.6E-13 |
| | | GOTERM_CC_DIRECT | extracellular matrix | | 103 | 3.5 | 4.0E-15 | 8.3E-13 |
| | | UP_KEYWORDS | Cell membrane | | 566 | 19.1 | 1.7E-14 | 1.1E-12 |
| | | UP_KEYWORDS | Extracellular matrix | | 87 | 2.9 | 6.0E-13 | 3.6E-11 |
| | | UP_KEYWORDS | Calcium | | 222 | 7.5 | 8.7E-13 | 4.6E-11 |

**Figure S2.2b. Gene ontology enrichment analysis of AS genes**
Functional annotation of EN-TEx AS protein-coding genes detected from different assays. Analysis was performed using DAVID Bioinformatics Resources 6.8. For each assay, the background list includes all protein-coding genes with accessible promoters for ASB or with accessible expressions for ASE. For ASE analysis, since DAVID has a 3,000 gene limit, the top 3,000 mostly ASE protein-coding genes were selected for the enrichment analysis, and the top 20 enriched terms are shown in the table. Terms with the largest number of genes are highlighted in bold.

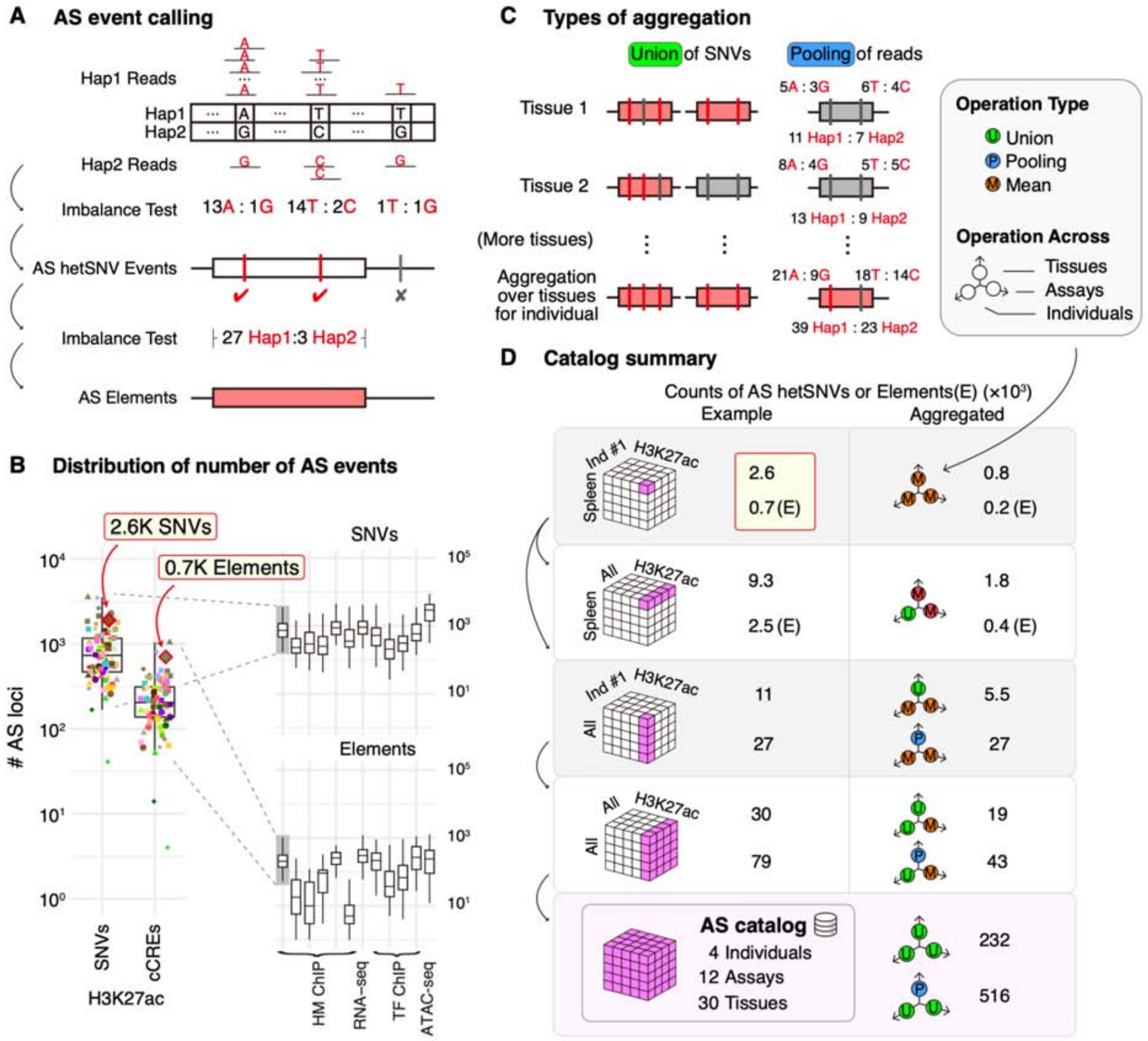

**Figure S2.3. Summary of AS catalog**

**(A)** Schematic of calling AS SNVs and AS elements. Examples of the number of reads are shown for each SNV and element. See S2.1 for details. **(B)** Distribution of the number of AS events in different assays. This is an abstract of Figure S2.2a. H3K27ac AS events in spleen from Individual 1 are shown as an example here and in panel (D). **(C)** Details of union and pooling of reads aggregation. For the union method, the AS SNV set is simply the union of AS SNV sets in all tissues. For the pooling method, reads from different tissues were summed first. The statistical test was done on the summary number, thus increasing the statistical power. **(D)** Summary of AS catalog. The colored circle at the right shows the aggregation method along that axis. Take the third row as an example. The upper one indicated that the operation is union across tissues, average across assays and individuals, while the bottom one is pooling reads across tissues, average across assays and individuals. Data obtained from merging the tissue-specific AS call sets resulted in ~5.5K unique AS hetSNVs (for either AS binding or expression) and ~1K AS genomic elements (not shown in figure) for each individual per assay. Pooling the reads from each assay across all tissues dramatically increased by ~5X fold the detection power, making it possible to identify ~27K AS hetSNVs per assay for each individual. Finally, merging across all assays provided a catalog of all loci where AS activity could be assessed in any of the tissues of the four individuals.

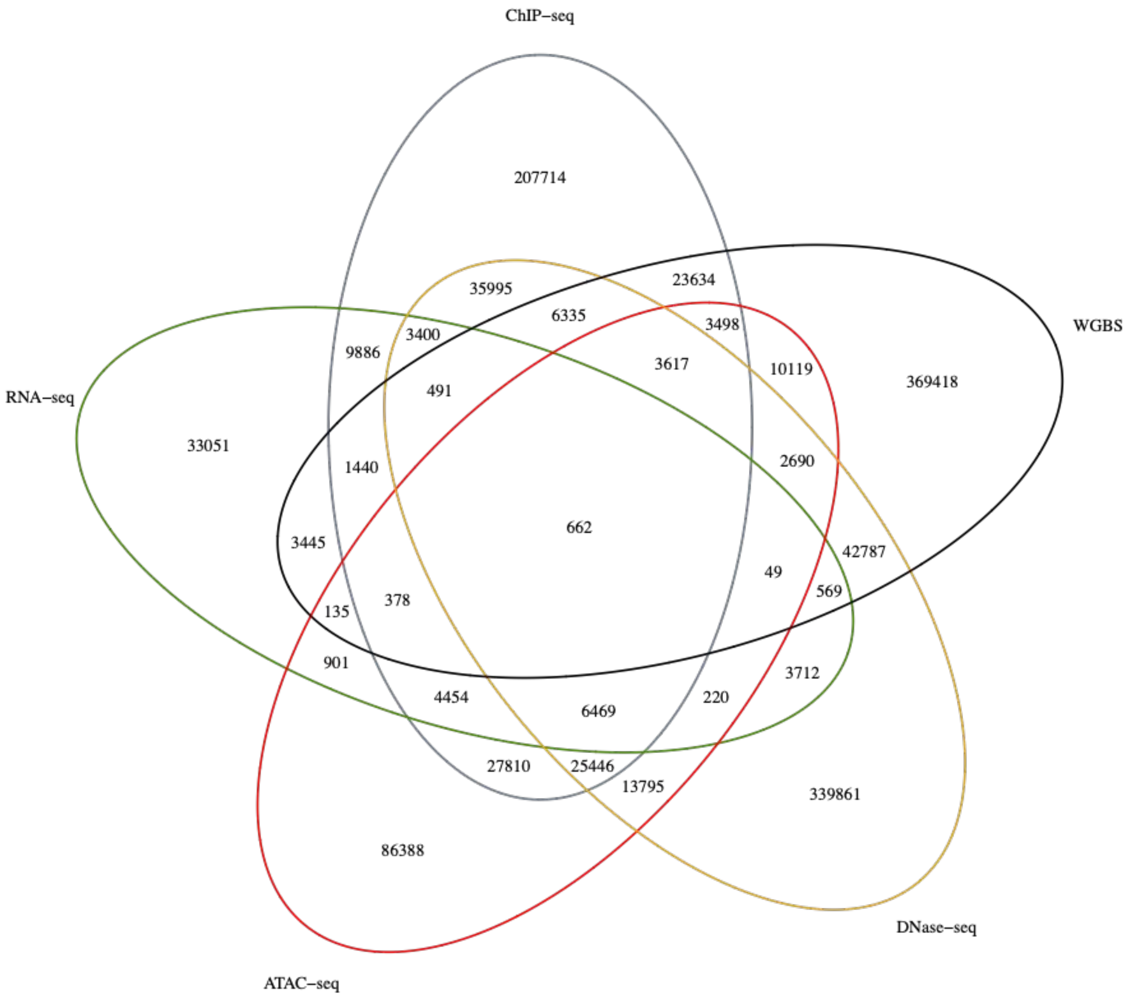

| | all (avg.) variants in 4 individuals | unique variants in 4 individuals | unique/all |
|---|---|---|---|
| **hetSNVs** | 9483K (2371K) | 6023K | 0.64 |
| **AS SNVs** | 1459K (365K) | 1268K | 0.87 (0.93) |
| **ASM SNVs** | 516K (129K) | 469K | 0.91 (0.97) |
| **ASE SNVs** | 79K (19.8K) | 69K | 0.87 (1.0) |
| **ASB SNVs** | 412K (103) | 361K | 0.88 (0.98) |
| **AS accessibility SNVs** | 682K (170.5K) | 620K | 0.91 (0.97) |

**Figure S2.4. Numbers of AS hetSNVs detected in different EN-TEx assays**
**(A)** Venn diagram of hetSNVs showing AS activity in different assays. The numbers are pooled from all four individuals. Heterozygous SNVs that appear in multiple individuals but have the same coordinates are collapsed into one unique heterozygous SNV. **(B)** The total numbers of AS hetSNVs (i.e., no collapsing; numbers in parentheses are the average in an individual), the numbers of unique ones, and the unique-to-all ratios. For a comparison with the unique-to-all

ratio of the AS SNVs, we randomly sampled the same number of heterozygous SNVs from the four individuals and calculated the unique-to-all ratio (numbers in parentheses). The smaller unique-to-all ratios of the AS SNVs suggest that these SNVs tend to be more common than random heterozygous SNVs. AS SNVs are those that show AS activity in any of the assays in **(A)**. ASM SNVs show AS activity in WGBS data, ASE SNVs in RNAs-seq data, ASB SNVs in ChIP-seq data, and AS accessibility SNVs in ATAC-seq or DNase-seq data.

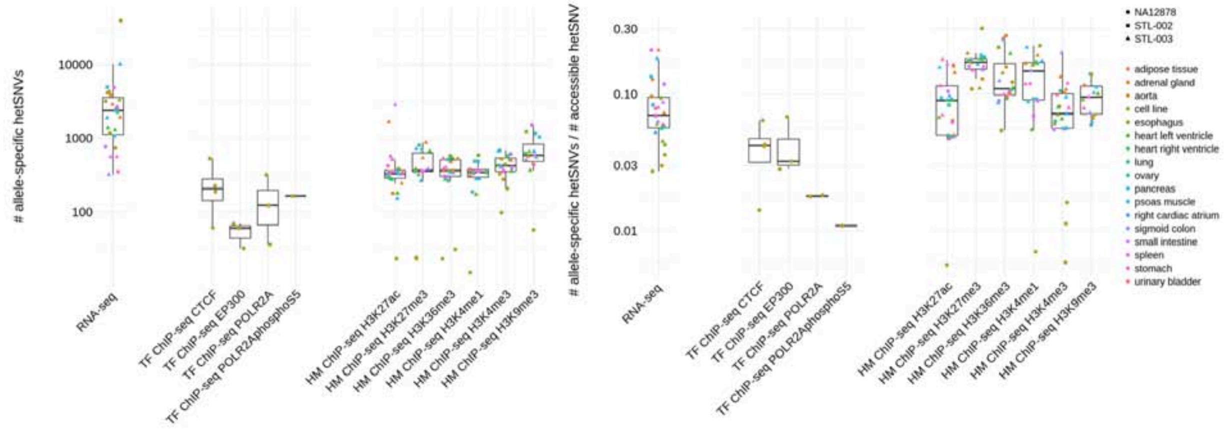

| Donor | Tissue | Assay | Experiment ID | # AS hetSNVs |
|---|---|---|---|---|
| STL-002 | adrenal gland | HM-ChIP-seq H3K27ac | ENCSR642HHF | 178 |
| STL-002 | aorta | HM-ChIP-seq H3K27ac | ENCSR322TJD | 350 |
| STL-002 | esophagus | HM-ChIP-seq H3K27ac | ENCSR645SYH | 386 |
| STL-002 | lung | HM-ChIP-seq H3K27ac | ENCSR540ADS | 312 |
| STL-002 | ovary | HM-ChIP-seq H3K27ac | ENCSR268JQE | 286 |
| STL-002 | pancreas | HM-ChIP-seq H3K27ac | ENCSR402HFW | 329 |
| STL-002 | psoas muscle | HM-ChIP-seq H3K27ac | ENCSR250NHD | 365 |
| STL-002 | small intestine | HM-ChIP-seq H3K27ac | ENCSR655XLM | 512 |
| STL-002 | spleen | HM-ChIP-seq H3K27ac | ENCSR086XCT | 353 |
| STL-002 | stomach | HM-ChIP-seq H3K27ac | ENCSR582UTE | 419 |
| STL-002 | aorta | HM-ChIP-seq H3K27me3 | ENCSR128VHV | 373 |
| STL-002 | esophagus | HM-ChIP-seq H3K27me3 | ENCSR641RQV | 353 |
| STL-002 | lung | HM-ChIP-seq H3K27me3 | ENCSR204NFO | 803 |
| STL-002 | ovary | HM-ChIP-seq H3K27me3 | ENCSR037SNV | 701 |
| STL-002 | small intestine | HM-ChIP-seq H3K27me3 | ENCSR877PAS | 357 |
| STL-002 | adrenal gland | HM-ChIP-seq H3K36me3 | ENCSR899MFS | 478 |
| STL-002 | aorta | HM-ChIP-seq H3K36me3 | ENCSR989AMI | 363 |
| STL-002 | esophagus | HM-ChIP-seq H3K36me3 | ENCSR279MCN | 580 |
| STL-002 | lung | HM-ChIP-seq H3K36me3 | ENCSR671NXL | 264 |
| STL-002 | ovary | HM-ChIP-seq H3K36me3 | ENCSR659MYS | 518 |
| STL-002 | pancreas | HM-ChIP-seq H3K36me3 | ENCSR393HBQ | 507 |
| STL-002 | small intestine | HM-ChIP-seq H3K36me3 | ENCSR073YZL | 304 |
| STL-002 | spleen | HM-ChIP-seq H3K36me3 | ENCSR078BHK | 300 |
| STL-002 | aorta | HM-ChIP-seq H3K4me1 | ENCSR848TLB | 376 |
| STL-002 | esophagus | HM-ChIP-seq H3K4me1 | ENCSR478BKA | 589 |
| STL-002 | lung | HM-ChIP-seq H3K4me1 | ENCSR356ANC | 187 |
| STL-002 | ovary | HM-ChIP-seq H3K4me1 | ENCSR113AFY | 488 |
| STL-002 | pancreas | HM-ChIP-seq H3K4me1 | ENCSR984UHU | 488 |
| STL-002 | small intestine | HM-ChIP-seq H3K4me1 | ENCSR538JMW | 355 |
| STL-002 | spleen | HM-ChIP-seq H3K4me1 | ENCSR115TSA | 332 |
| STL-002 | adrenal gland | HM-ChIP-seq H3K4me3 | ENCSR425NQT | 571 |
| STL-002 | aorta | HM-ChIP-seq H3K4me3 | ENCSR960EVO | 327 |
| STL-002 | esophagus | HM-ChIP-seq H3K4me3 | ENCSR697GPO | 443 |
| STL-002 | lung | HM-ChIP-seq H3K4me3 | ENCSR466DZW | 683 |
| STL-002 | ovary | HM-ChIP-seq H3K4me3 | ENCSR139TLA | 510 |
| STL-002 | pancreas | HM-ChIP-seq H3K4me3 | ENCSR315LPR | 423 |
| STL-002 | small intestine | HM-ChIP-seq H3K4me3 | ENCSR944QSH | 397 |
| STL-002 | esophagus | HM-ChIP-seq H3K9me3 | ENCSR200WDD | 1225 |
| STL-002 | lung | HM-ChIP-seq H3K9me3 | ENCSR728FLA | 625 |
| STL-002 | ovary | HM-ChIP-seq H3K9me3 | ENCSR956UFV | 1157 |
| STL-002 | pancreas | HM-ChIP-seq H3K9me3 | ENCSR533HDU | 1044 |
| STL-002 | small intestine | HM-ChIP-seq H3K9me3 | ENCSR270VNK | 1506 |
| STL-002 | adipose tissue | RNA-seq | ENCSR686JJB | 2440 |
| STL-002 | adrenal gland | RNA-seq | ENCSR146ZKR | 3838 |
| STL-002 | esophagus | RNA-seq | ENCSR993QGR | 3179 |
| STL-002 | lung | RNA-seq | ENCSR917YHC | 2278 |
| STL-002 | ovary | RNA-seq | ENCSR725TPW | 2373 |
| STL-002 | pancreas | RNA-seq | ENCSR571BML | 4929 |
| STL-002 | psoas muscle | RNA-seq | ENCSR502OTI | 1079 |
| STL-002 | small intestine | RNA-seq | ENCSR039ICU | 767 |
| STL-002 | spleen | RNA-seq | ENCSR510PSL | 4863 |
| STL-002 | stomach | RNA-seq | ENCSR980UEY | 351 |
| NA12878 | cell line | ATAC-seq | ENCSR095QNB | 3789 |
| NA12878 | cell line | ATAC-seq | ENCSR637XSC | 19793 |
| NA12878 | cell line | HM-ChIP-seq H3K27ac | ENCSR000AKC | 23 |
| NA12878 | cell line | HM-ChIP-seq H3K27me3 | ENCSR000DRX | 24 |
| NA12878 | cell line | HM-ChIP-seq H3K27me3 | ENCSR000AKD | 23 |
| NA12878 | cell line | HM-ChIP-seq H3K36me3 | ENCSR000AKE | 23 |
| NA12878 | cell line | HM-ChIP-seq H3K36me3 | ENCSR000DRW | 31 |
| NA12878 | cell line | HM-ChIP-seq H3K4me1 | ENCSR000AKF | 15 |
| NA12878 | cell line | HM-ChIP-seq H3K4me3 | ENCSR057BWO | 98 |
| NA12878 | cell line | HM-ChIP-seq H3K4me3 | ENCSR000AKA | 207 |
| NA12878 | cell line | HM-ChIP-seq H3K4me3 | ENCSR000DRY | 277 |
| NA12878 | cell line | HM-ChIP-seq H3K9me3 | ENCSR000AOX | 57 |
| NA12878 | cell line | RNA-seq | ENCSR000AEC | 4267 |
| NA12878 | cell line | RNA-seq | ENCSR151NGC | 734 |
| NA12878 | cell line | RNA-seq | ENCSR820PHH | 1166 |
| NA12878 | cell line | RNA-seq | ENCSR000AEE | 4122 |
| NA12878 | cell line | TF-ChIP-seq CTCF | ENCSR000DRZ | 188 |
| NA12878 | cell line | TF-ChIP-seq CTCF | ENCSR000DZN | 528 |
| NA12878 | cell line | TF-ChIP-seq CTCF | ENCSR000AKB | 61 |
| NA12878 | cell line | TF-ChIP-seq CTCF | ENCSR000DKV | 226 |
| NA12878 | cell line | TF-ChIP-seq EP300 | ENCSR000BHB | 32 |
| NA12878 | cell line | TF-ChIP-seq EP300 | ENCSR000DZG | 69 |
| NA12878 | cell line | TF-ChIP-seq EP300 | ENCSR000DZD | 60 |
| NA12878 | cell line | TF-ChIP-seq POLR2A | ENCSR000DKT | 36 |
| NA12878 | cell line | TF-ChIP-seq POLR2A | ENCSR000EAD | 122 |
| NA12878 | cell line | TF-ChIP-seq POLR2A | ENCSR000BGD | 318 |
| NA12878 | cell line | TF-ChIP-seq POLR2Aphos | ENCSR000BIF | 164 |

| Donor | Tissue | Assay | Experiment ID | # AS hetSNVs |
|---|---|---|---|---|
| STL-003 | adipose tissue | HM-ChIP-seq H3K27ac | ENCSR082SHT | 340 |
| STL-003 | adrenal gland | HM-ChIP-seq H3K27ac | ENCSR405ESP | 1681 |
| STL-003 | aorta | HM-ChIP-seq H3K27ac | ENCSR519CFV | 243 |
| STL-003 | esophagus | HM-ChIP-seq H3K27ac | ENCSR679OVD | 297 |
| STL-003 | heart left ventricle | HM-ChIP-seq H3K27ac | ENCSR150QXE | 179 |
| STL-003 | heart right ventricle | HM-ChIP-seq H3K27ac | ENCSR928HSI | 276 |
| STL-003 | pancreas | HM-ChIP-seq H3K27ac | ENCSR612BWE | 312 |
| STL-003 | psoas muscle | HM-ChIP-seq H3K27ac | ENCSR791ISZ | 152 |
| STL-003 | right cardiac atrium | HM-ChIP-seq H3K27ac | ENCSR074ECR | 333 |
| STL-003 | sigmoid colon | HM-ChIP-seq H3K27ac | ENCSR561YSH | 324 |
| STL-003 | small intestine | HM-ChIP-seq H3K27ac | ENCSR543CPW | 2849 |
| STL-003 | spleen | HM-ChIP-seq H3K27ac | ENCSR235ZBF | 324 |
| STL-003 | stomach | HM-ChIP-seq H3K27ac | ENCSR001SHB | 338 |
| STL-003 | urinary bladder | HM-ChIP-seq H3K27ac | ENCSR054BKO | 568 |
| STL-003 | adrenal gland | HM-ChIP-seq H3K27me3 | ENCSR181JFC | 876 |
| STL-003 | aorta | HM-ChIP-seq H3K27me3 | ENCSR196PGM | 537 |
| STL-003 | esophagus | HM-ChIP-seq H3K27me3 | ENCSR088GXB | 382 |
| STL-003 | heart left ventricle | HM-ChIP-seq H3K27me3 | ENCSR503YOF | 664 |
| STL-003 | heart right ventricle | HM-ChIP-seq H3K27me3 | ENCSR068CQX | 317 |
| STL-003 | pancreas | HM-ChIP-seq H3K27me3 | ENCSR186QKH | 352 |
| STL-003 | psoas muscle | HM-ChIP-seq H3K27me3 | ENCSR720SAS | 263 |
| STL-003 | right cardiac atrium | HM-ChIP-seq H3K27me3 | ENCSR972RKX | 715 |
| STL-003 | sigmoid colon | HM-ChIP-seq H3K27me3 | ENCSR042RIW | 383 |
| STL-003 | spleen | HM-ChIP-seq H3K27me3 | ENCSR408ONP | 356 |
| STL-003 | stomach | HM-ChIP-seq H3K27me3 | ENCSR527BFF | 352 |
| STL-003 | adrenal gland | HM-ChIP-seq H3K36me3 | ENCSR942XCE | 359 |
| STL-003 | aorta | HM-ChIP-seq H3K36me3 | ENCSR673JYT | 530 |
| STL-003 | esophagus | HM-ChIP-seq H3K36me3 | ENCSR034ZHF | 363 |
| STL-003 | heart left ventricle | HM-ChIP-seq H3K36me3 | ENCSR434MDA | 541 |
| STL-003 | heart right ventricle | HM-ChIP-seq H3K36me3 | ENCSR142OBQ | 277 |
| STL-003 | pancreas | HM-ChIP-seq H3K36me3 | ENCSR943JOF | 345 |
| STL-003 | sigmoid colon | HM-ChIP-seq H3K36me3 | ENCSR445RFF | 339 |
| STL-003 | spleen | HM-ChIP-seq H3K36me3 | ENCSR466DUB | 401 |
| STL-003 | stomach | HM-ChIP-seq H3K36me3 | ENCSR552MZH | 386 |
| STL-003 | urinary bladder | HM-ChIP-seq H3K36me3 | ENCSR449TNC | 531 |
| STL-003 | adrenal gland | HM-ChIP-seq H3K4me1 | ENCSR511GOF | 344 |
| STL-003 | aorta | HM-ChIP-seq H3K4me1 | ENCSR325VOA | 309 |
| STL-003 | esophagus | HM-ChIP-seq H3K4me1 | ENCSR306ZBD | 358 |
| STL-003 | heart left ventricle | HM-ChIP-seq H3K4me1 | ENCSR111WGZ | 173 |
| STL-003 | heart right ventricle | HM-ChIP-seq H3K4me1 | ENCSR076CZA | 342 |
| STL-003 | pancreas | HM-ChIP-seq H3K4me1 | ENCSR449PYI | 342 |
| STL-003 | psoas muscle | HM-ChIP-seq H3K4me1 | ENCSR410UUH | 279 |
| STL-003 | right cardiac atrium | HM-ChIP-seq H3K4me1 | ENCSR671BOA | 283 |
| STL-003 | sigmoid colon | HM-ChIP-seq H3K4me1 | ENCSR782OZZ | 362 |
| STL-003 | spleen | HM-ChIP-seq H3K4me1 | ENCSR490YCL | 374 |
| STL-003 | stomach | HM-ChIP-seq H3K4me1 | ENCSR257BCD | 376 |
| STL-003 | adrenal gland | HM-ChIP-seq H3K4me3 | ENCSR234YIU | 501 |
| STL-003 | aorta | HM-ChIP-seq H3K4me3 | ENCSR957BPJ | 520 |
| STL-003 | esophagus | HM-ChIP-seq H3K4me3 | ENCSR577ILY | 353 |
| STL-003 | heart left ventricle | HM-ChIP-seq H3K4me3 | ENCSR487BEW | 422 |
| STL-003 | heart right ventricle | HM-ChIP-seq H3K4me3 | ENCSR791GCO | 556 |
| STL-003 | pancreas | HM-ChIP-seq H3K4me3 | ENCSR747VED | 354 |
| STL-003 | psoas muscle | HM-ChIP-seq H3K4me3 | ENCSR949OYZ | 668 |
| STL-003 | right cardiac atrium | HM-ChIP-seq H3K4me3 | ENCSR548LZS | 605 |
| STL-003 | sigmoid colon | HM-ChIP-seq H3K4me3 | ENCSR421HUB | 398 |
| STL-003 | small intestine | HM-ChIP-seq H3K4me3 | ENCSR792IJA | 401 |
| STL-003 | spleen | HM-ChIP-seq H3K4me3 | ENCSR432KIH | 371 |
| STL-003 | stomach | HM-ChIP-seq H3K4me3 | ENCSR129NCV | 351 |
| STL-003 | urinary bladder | HM-ChIP-seq H3K4me3 | ENCSR632OWD | 607 |
| STL-003 | adrenal gland | HM-ChIP-seq H3K9me3 | ENCSR992VZG | 459 |
| STL-003 | aorta | HM-ChIP-seq H3K9me3 | ENCSR065ZNA | 564 |
| STL-003 | esophagus | HM-ChIP-seq H3K9me3 | ENCSR150GLE | 654 |
| STL-003 | heart left ventricle | HM-ChIP-seq H3K9me3 | ENCSR176KNR | 366 |
| STL-003 | pancreas | HM-ChIP-seq H3K9me3 | ENCSR035QNZ | 549 |
| STL-003 | right cardiac atrium | HM-ChIP-seq H3K9me3 | ENCSR596BHN | 661 |
| STL-003 | sigmoid colon | HM-ChIP-seq H3K9me3 | ENCSR737NLJ | 507 |
| STL-003 | spleen | HM-ChIP-seq H3K9me3 | ENCSR421FPV | 587 |
| STL-003 | stomach | HM-ChIP-seq H3K9me3 | ENCSR639RKZ | 438 |
| STL-003 | adipose tissue | RNA-seq | ENCSR741QEH | 2894 |
| STL-003 | adrenal gland | RNA-seq | ENCSR598KJX | 1918 |
| STL-003 | esophagus | RNA-seq | ENCSR102TQN | 3320 |
| STL-003 | heart left ventricle | RNA-seq | ENCSR769LNJ | 1285 |
| STL-003 | heart right ventricle | RNA-seq | ENCSR433XCV | 1375 |
| STL-003 | pancreas | RNA-seq | ENCSR629VMZ | 1893 |
| STL-003 | psoas muscle | RNA-seq | ENCSR843HXR | 10137 |
| STL-003 | right cardiac atrium | RNA-seq | ENCSR675YAS | 2560 |
| STL-003 | sigmoid colon | RNA-seq | ENCSR999ZCI | 321 |
| STL-003 | small intestine | RNA-seq | ENCSR719HRO | 557 |
| STL-003 | spleen | RNA-seq | ENCSR910QOX | 2674 |
| STL-003 | stomach | RNA-seq | ENCSR721HDG | 556 |

**Figure S2.5. Construction of validation dataset from AS events in non-EN-TEx datasets**
**(A)** Distribution and fraction of the number of hetSNVs associated with AS behavior detected in NA12878 and Roadmap individuals STL002 and STL003. **(B)** Datasets used for calling AS events in the Roadmap individuals.

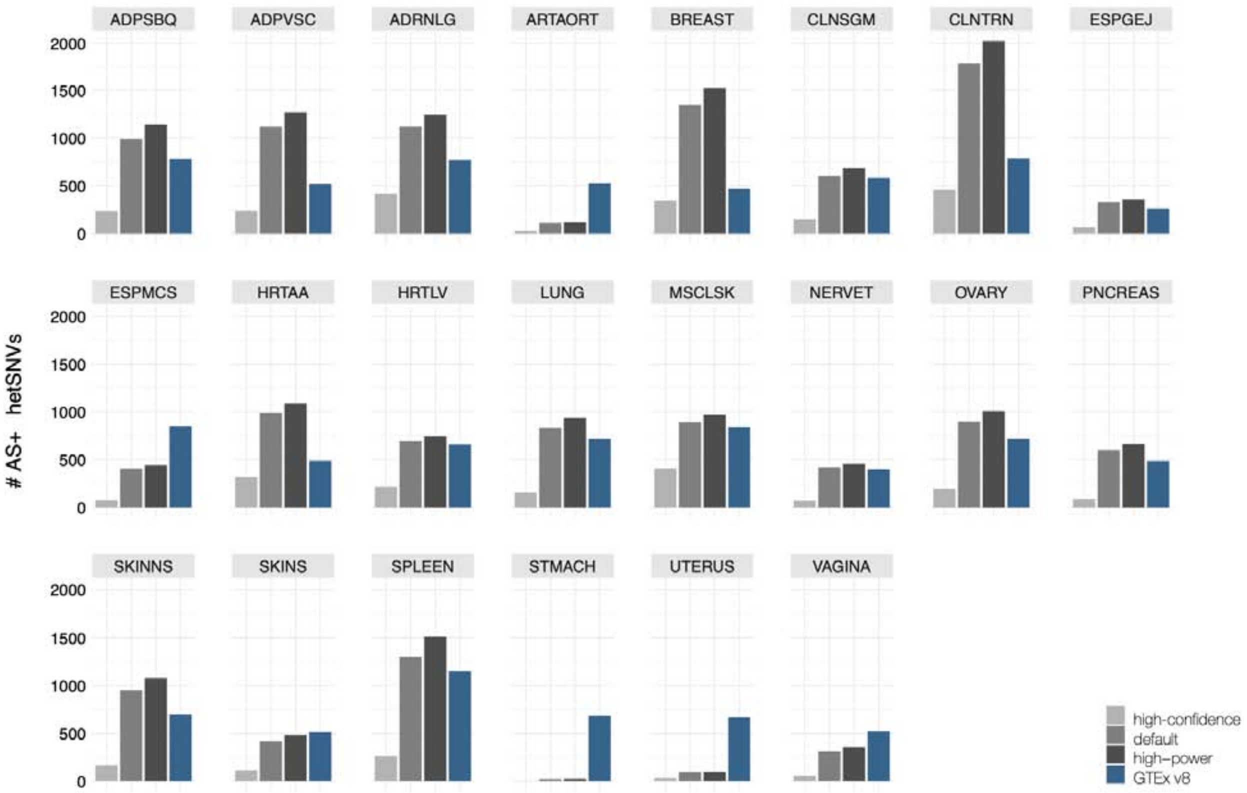

**Figure S2.6a. Numbers of AS hetSNVs detected from RNA-seq in different tissues from individual 3**

To produce high-power tissue-specific call sets, we called ASE and ASB sites for each tissue at a relaxed FDR threshold if the hetSNV was called AS in the pooled call set; otherwise, the sites were called at the usual 10% FDR. The “relaxed” FDR varied somewhat from tissue to tissue due to granularities in calculation but did not exceed 20%. Typically, high-power call sets produce 10%–20% more AS hetSNVs than typical call sets. See results in File: hetSNVs_high-power_AS.tsv.

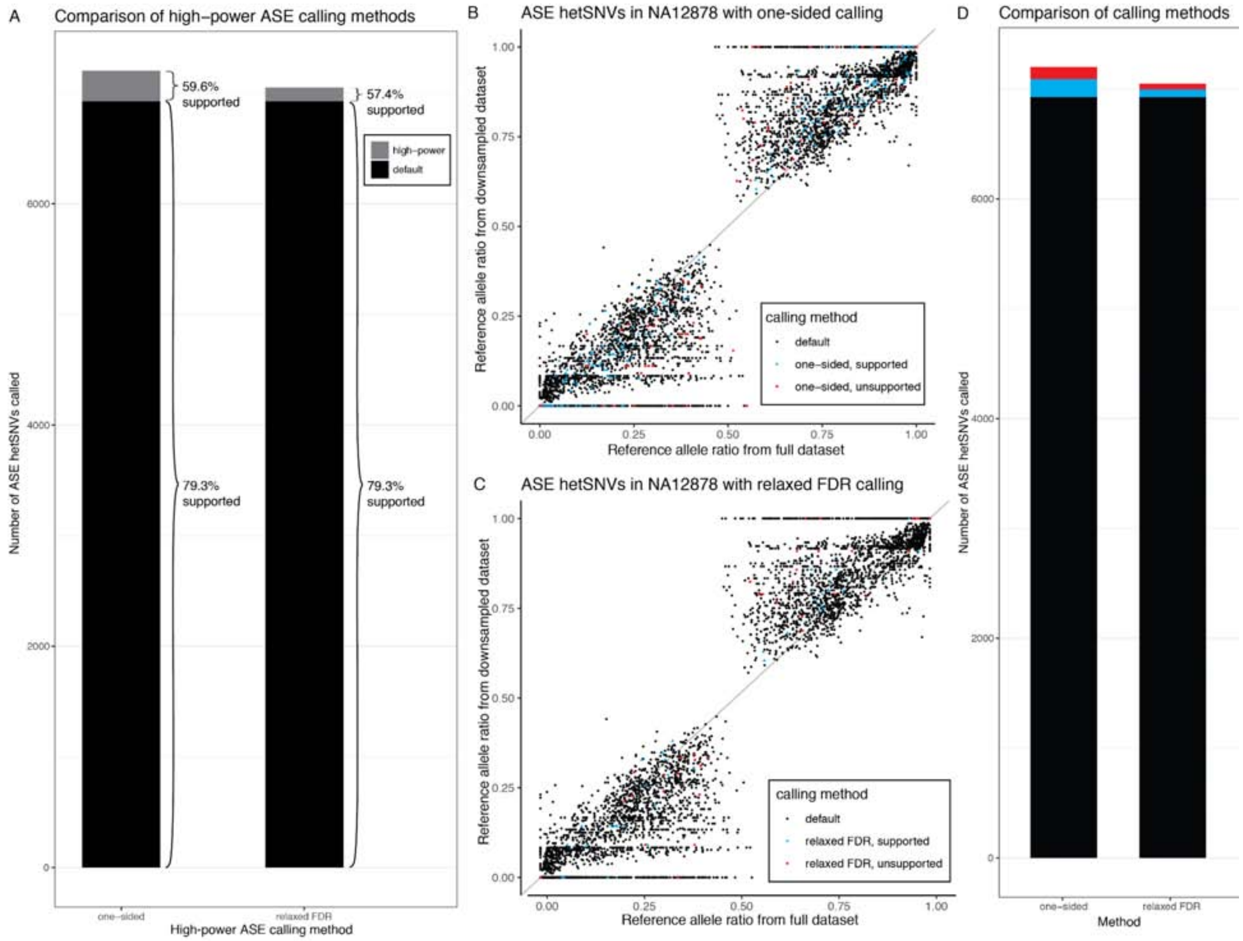

**Figure S2.6b. Validation of high-power AS calling methods**

To increase the detection power of ASE hetSNVs in datasets with fewer reads, we tested two high-power calling methods that selectively impose less-stringent tests on hetSNVs, which have been shown to have AS behavior in other experiments. The first method uses a one-sided beta-binomial test as its less-stringent test, while the second uses a two-sided beta-binomial test with a relaxed FDR of 20%. All hetSNVs that do not have prior evidence of being AS are evaluated with the standard two-sided beta-binomial test with an FDR of 10%. We validated both methods by testing on a deeply sequenced RNA-seq dataset from the GM12878 cell line, and simulating a shallower experiment by downsampling this dataset by a factor of 4. A, D. Using the default ASE calling method, 6,927 ASE hetSNVs were identified in the downsampled dataset. Of these, 79.3% were supported by the full RNA-seq dataset – that is, they were also called ASE in the full dataset. One-sided testing identified 275 additional ASE hetSNVs, and relaxed FDR testing identified 122 additional ASE hetSNVs. Both methods are enriched for supported hetSNVs as compared to the full pool of hetSNVs (59.6% and 57.4%, respectively), though they have a higher error rate than the default ASE calling method in this respect. B, C. We also show a comparison of reference allele ratios of ASE hetSNVs under different calling methods. Overall, the ASE hetSNVs added by both one-sided and relaxed FDR calling display similar reference allele ratios to ASE hetSNVs identified by default calling.

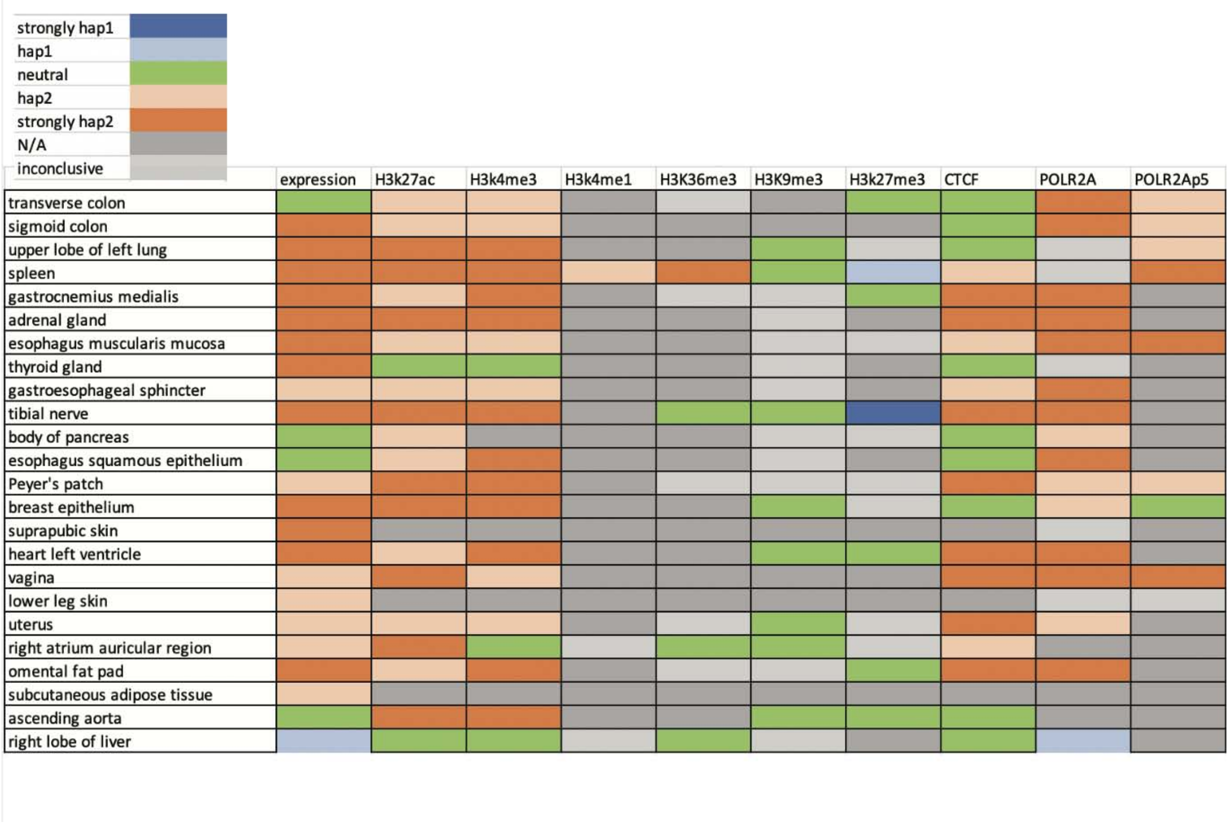

**Figure S2.8a. Heatmap to show haplotype specificity of Chr X for all assays and tissues from individual 3**

We detected that ~21% of the accessible X-chromosome genes have significantly imbalanced expression levels between the two haplotypes. Orange squares indicate more expression and binding peaks in haplotype 2, whereas blue squares indicate more expression and binding peaks in haplotype 1. Green squares indicate that the expression and binding are balanced between haplotypes. Light gray squares indicate that the number of data points is small and that, consequently, we cannot conclude which haplotype has more expression and binding. Dark gray squares indicate that data are not available for a given assay and tissue.

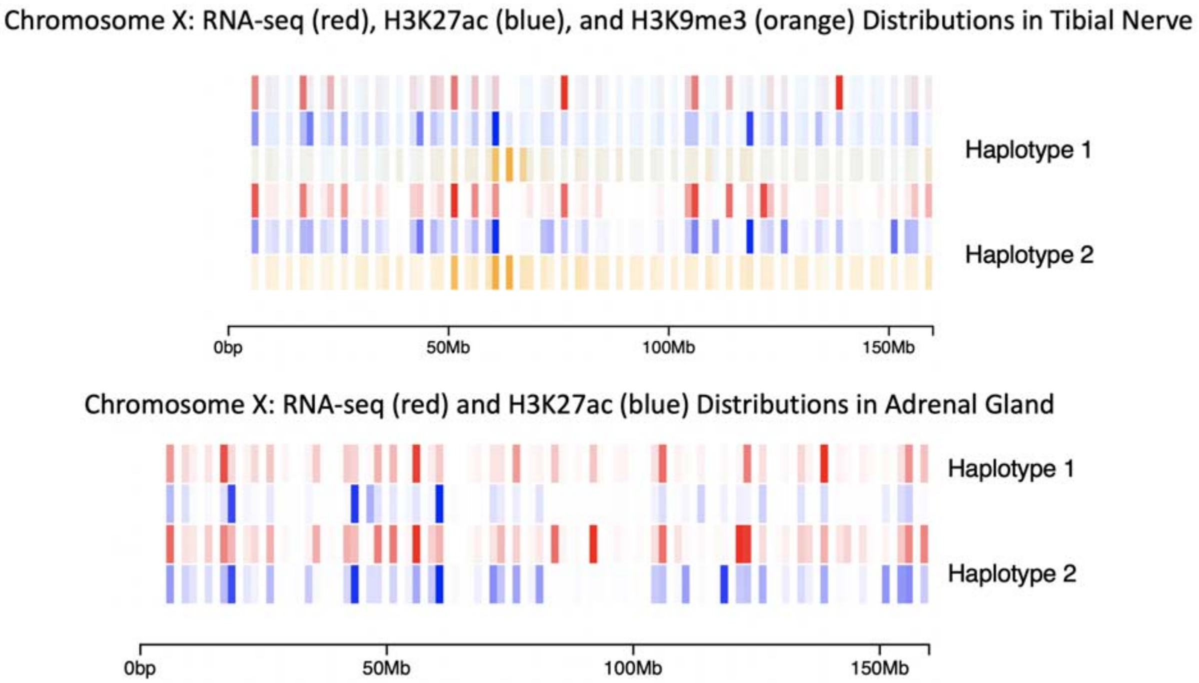

**Figure S2.8b. Chromosome painting of ChrX using RNA-Seq and ChIP-Seq in both haplotypes of individual 3 in two tissues**

This plot shows that the active haplotype is haplotype 2 in Chr X of individual 3, as there is more activity in haplotype 2.

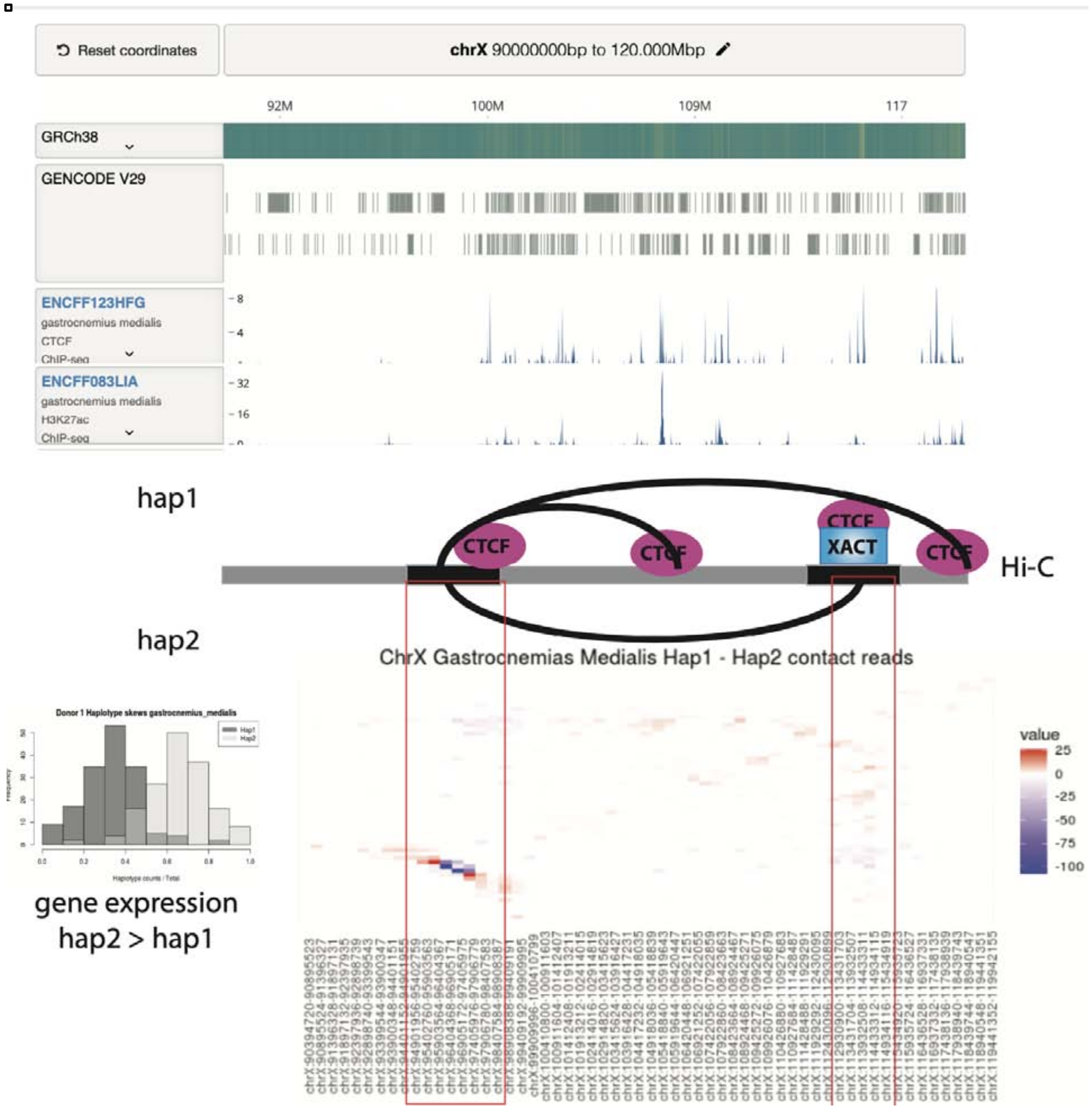

**Figure S2.8c. XACT locus on ChrX is shown to have haplotype-specific chromatin interactions with an upstream region.**

In the signal tracks, both XACT and upstream loci are shown to have CTCF bound, which is also associated with the H3K27ac signal. The heatmap shows differential chromatin interactions from haplotype-resolved Hi-C. The AS Hi-C interaction with the XACT locus and an upstream element occurs on the active haplotype, which was characterized by the difference in AS gene expression values (histogram).

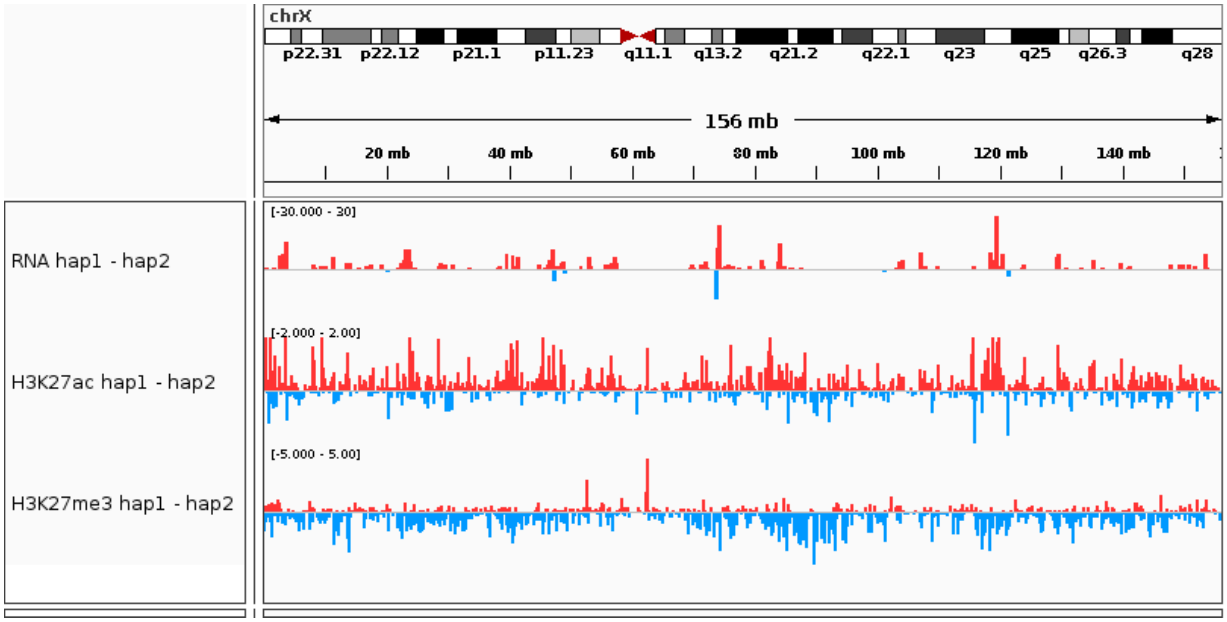

**Figure S2.8d. Coordinated AS activity in X chromosomes.**
Similar to Figure 3C, we show the differences in the levels of gene expression, H3K27ac, and H3K27me3 between the two X chromosomes in the thyroid gland of individual 4. The high RNA expression levels from haplotype 1 indicates that this X chromosome is active. Note the higher H3K27ac levels and lower H3K27me3 levels in this X chromosome.

#### Figure S3. Supp. Figures for Main Text Section “Interrelating SVs & Chromatin Modifications”

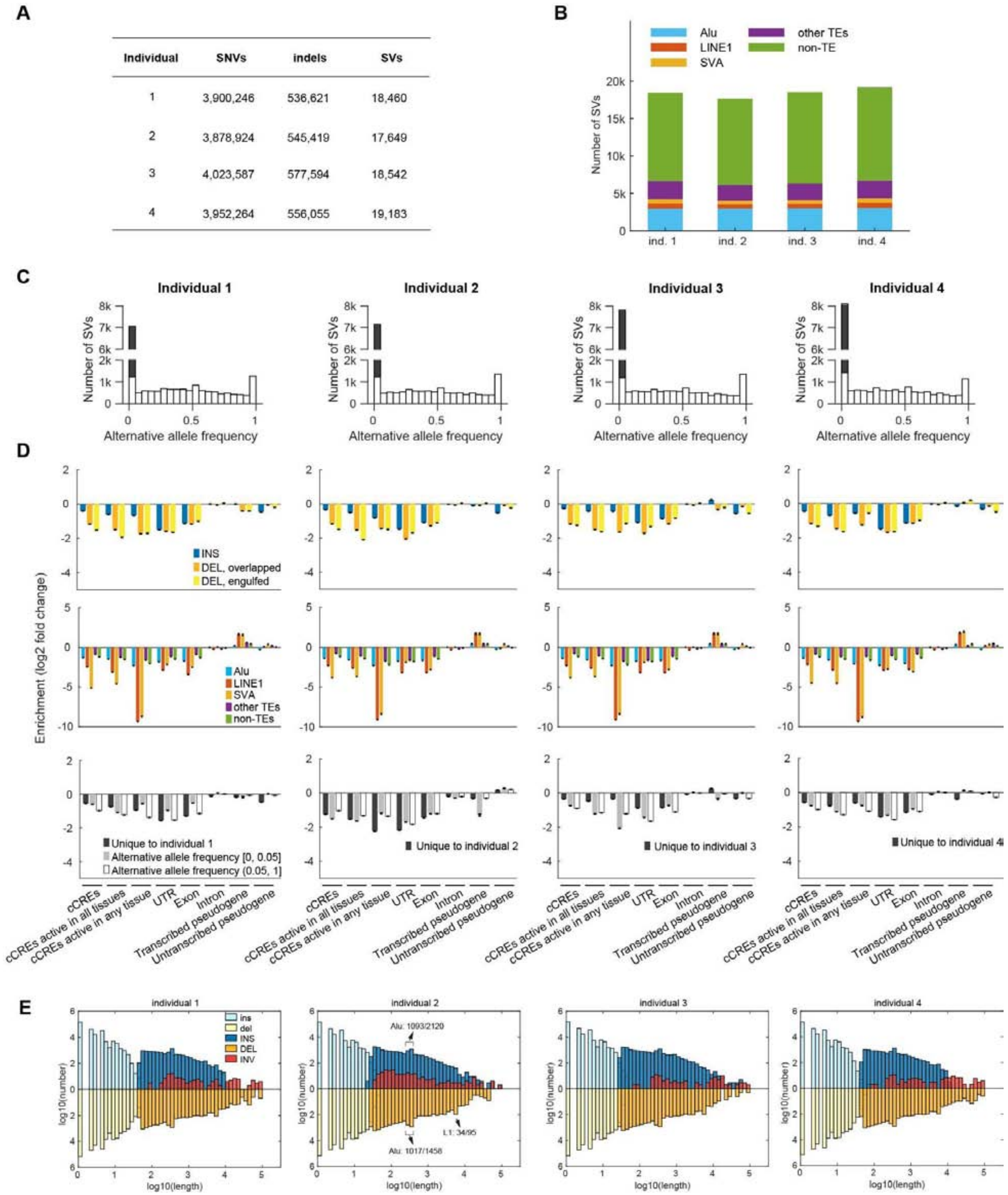

| Individual | SNVs | indels | SVs |
| --- | --- | --- | --- |
| 1 | 3,900,246 | 536,621 | 18,460 |
| 2 | 3,878,924 | 545,419 | 17,649 |
| 3 | 4,023,587 | 577,594 | 18,542 |
| 4 | 3,952,264 | 556,055 | 19,183 |

**Figure S3.1. Analysis of SVs**

**(A)** Number of genomic variants in the four individuals. **(B)** Numbers of SVs associated with transposable elements. **(C)** Allele frequencies of SVs in the European population calculated by overlapping with the results from Audano et al. (2019) (Audano *et al.*, 2019). SVs that have no overlap with the results from Audano et al. are shaded in the first bin. **(D)** Overlaps between SVs and functional genomic regions. We shuffle the locations of the SVs (see S3.1) to determine whether SVs are enriched or depleted in a given type of genomic region. For DELs, we consider cases in which a DEL partially overlaps with a given genomic region (DEL, partial) and cases in which a DEL is engulfed by a given genomic region (DEL, engulfed). **(E)** Lengths of indels and SVs in the four individuals. The peaks around 10^2.5 bp and 10^3.7 bp are due to Alu and LINE1. In individual 2, we show the fractions of SVs associated with Alu and LINE1 in the corresponding bins. Note that these fractions are much higher than those in **(B)**.

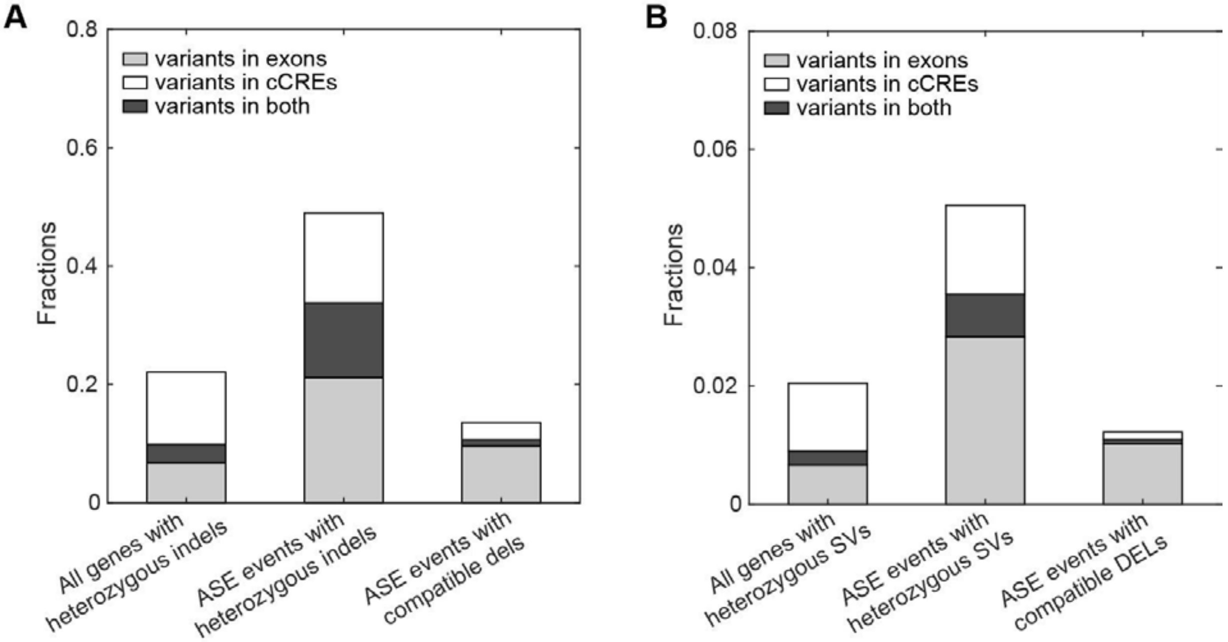

**Figure S3.2a. ASE events associated with indels and SVs.**
For an ASE event in a given tissue of a given individual, we looked for heterozygous indels and SVs that intersect with the exons of the ASE gene, and/or with cCREs within +/- 10 Kb of the gene's TSS. For comparison, we also show the fractions of genes (ASE or not) whose exons and/or nearby cCREs intersect with a heterozygous indel and SV. If the heterozygous small deletion and DEL had clear genotypes, we further evaluated whether they are compatible with the ASE event, i.e., the presence of the variants in exons and/or the tissue-specific active cCREs (see S7.2) should reduce gene expression. Since the exact breakpoints of SVs are often uncertain and SVs may disrupt nearby regions, we expanded the location of each SV by 100 bp upstream and 100 bp downstream when intersecting it with exons and cCREs. **(A)** The fractions of ASE events associated with indels and compatible deletions. **(B)** The fractions of ASE events associated with SVs and compatible DELs. In both panels, we pooled the ASE events and ASE events with associated variants from all tissues of all four individuals before calculating the fractions. Specifically in **(B)**, we found 42 ASE events that are associated with compatible DELs in the tissue-specific active cCREs, 323 associated compatible DELs in exons, and 22 associated with compatible DELs in both.

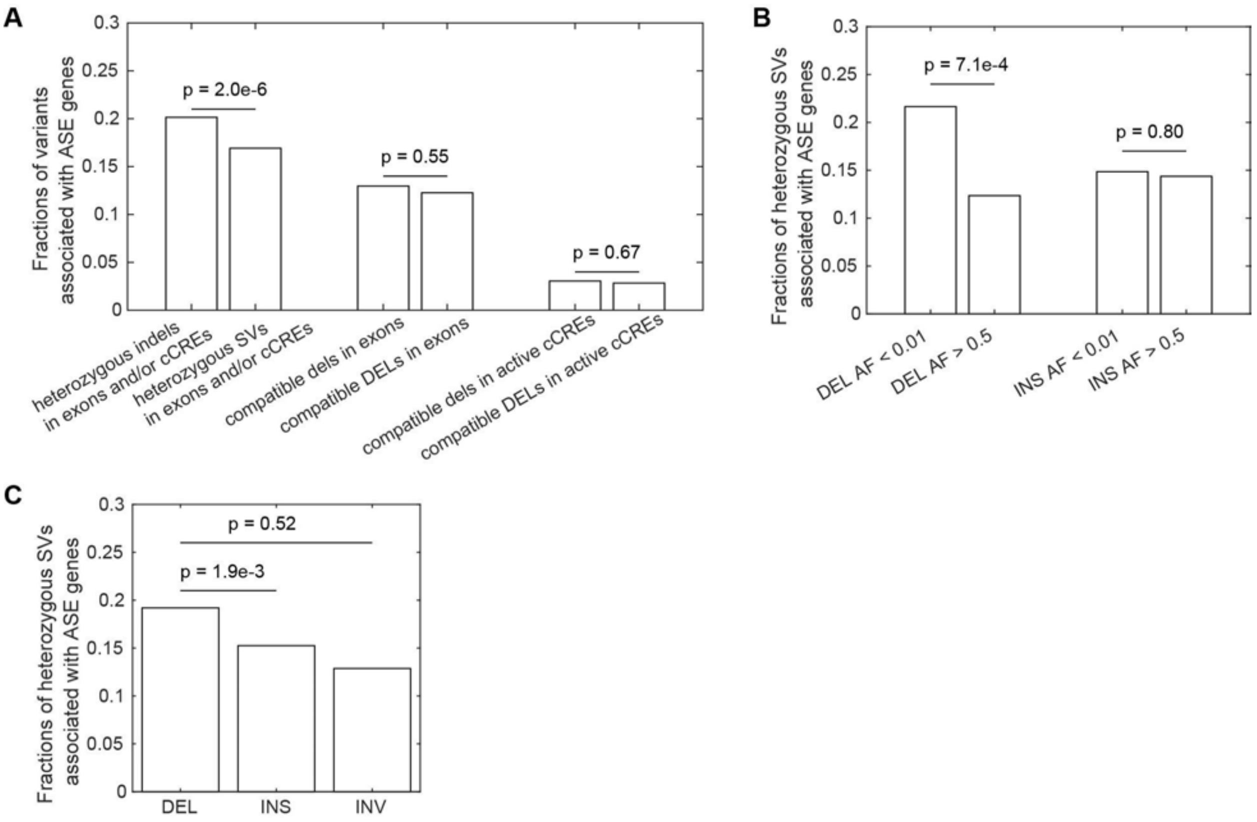

**Figure S3.2b. Indels and SVs associated with ASE**

Similar to Figure S3.2a, we looked for heterozygous indels and SVs that intersect with at least one of two genomic regions: an exon, and cCREs that are within +/- 10 Kb of a TSS. Among these variants, we calculated the fractions of those where the associated gene shows ASE in at least one tissue of the individual who carries the variants. We expanded the location of the SV by 100 bp upstream and 100 bp downstream before intersecting with exons and cCREs. (A) The fractions of indels and SVs associated with ASE. For heterozygous deletions and DEL that have clear genotypes, we further evaluated whether they are compatible with the associated ASE (Figure S3.2a). The fractions of compatible variants among those that intersect with an exon or any tissue-specific active cCREs are shown in separate groups. (B) The fractions of rare (AF < 0.01) and common (AF > 0.05) SVs that are associated with ASE among those that intersect with an exon and/or cCREs. Because the AF of INV are below 0.01 in each of the four individuals, we could not compare rare vs. common INV. (C) The fractions of DEL, INS, and INV that are associated with ASE among each type of SV that intersect with an exon and/or cCREs. In all panels, we pooled variants of interest from all tissues of all four individuals before calculating the fractions. Differences between fractions were tested via the $\chi 2$ test.

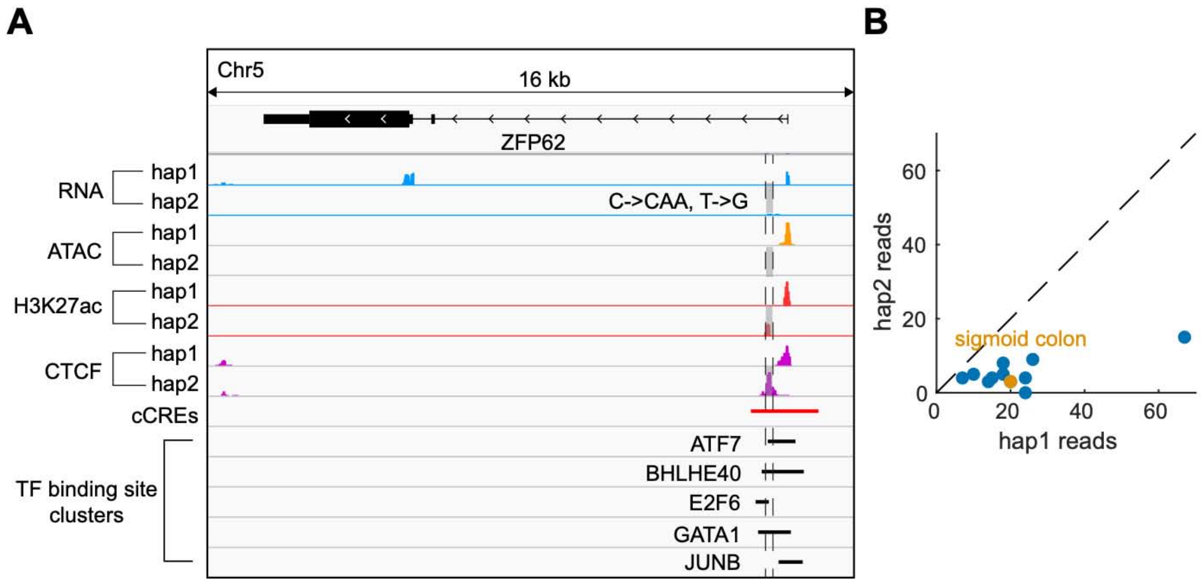

**Figure S3.2c. An indel that potentially changes gene expression**
**(A)** In the sigmoid colon of individual 2, the gene *ZFP62* has lower expression in haplotype 2. The TSS region of *ZFP62* in hap2 shows lower chromatin accessibility and changes in the positions of H3K27ac and CTCF binding peaks, compared with the same region in hap1. In hap2, a 2 bp insertion and an SNV were found in a cCRE near the TSS of the gene (the two variants are very close and are shown together by a single grey box). These variants and nearby variants that cannot be phased (not shown) might affect the function of the cCRE. **(B)** The gene has lower hap2 expression in multiple tissues, suggesting a universal factor changing the expression between haplotypes.

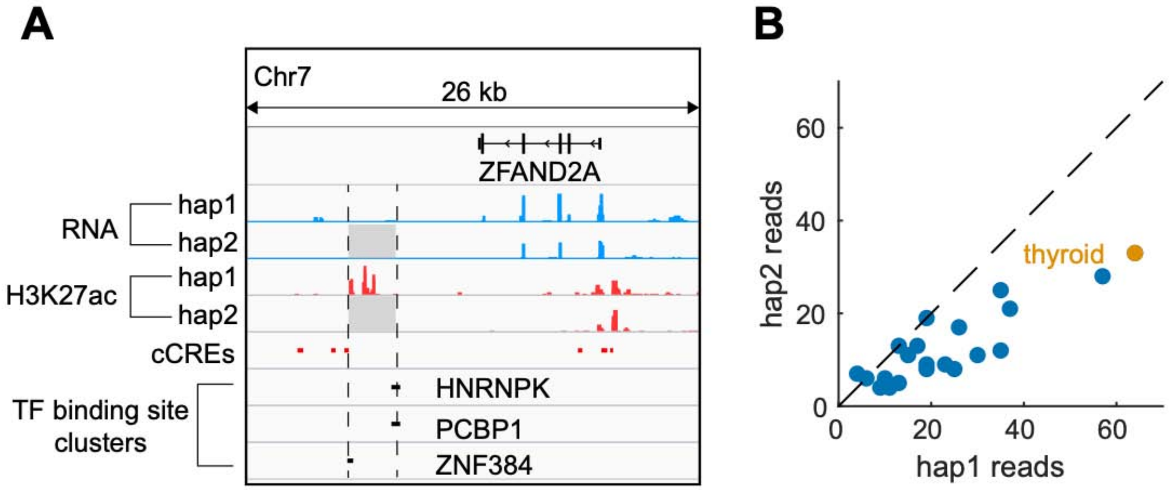

**Figure S3.2d. Shadow figure associated with Figure 4D**

**(A)** Similar to Figure 4D, the deletion in hap2 can disrupt cCREs identified in the thyroid and the binding of several TFs. **(B)** *ZFAND2A* has lower hap2 expression among multiple tissues, suggesting that the deletion may have a global effect on the expression of this gene.

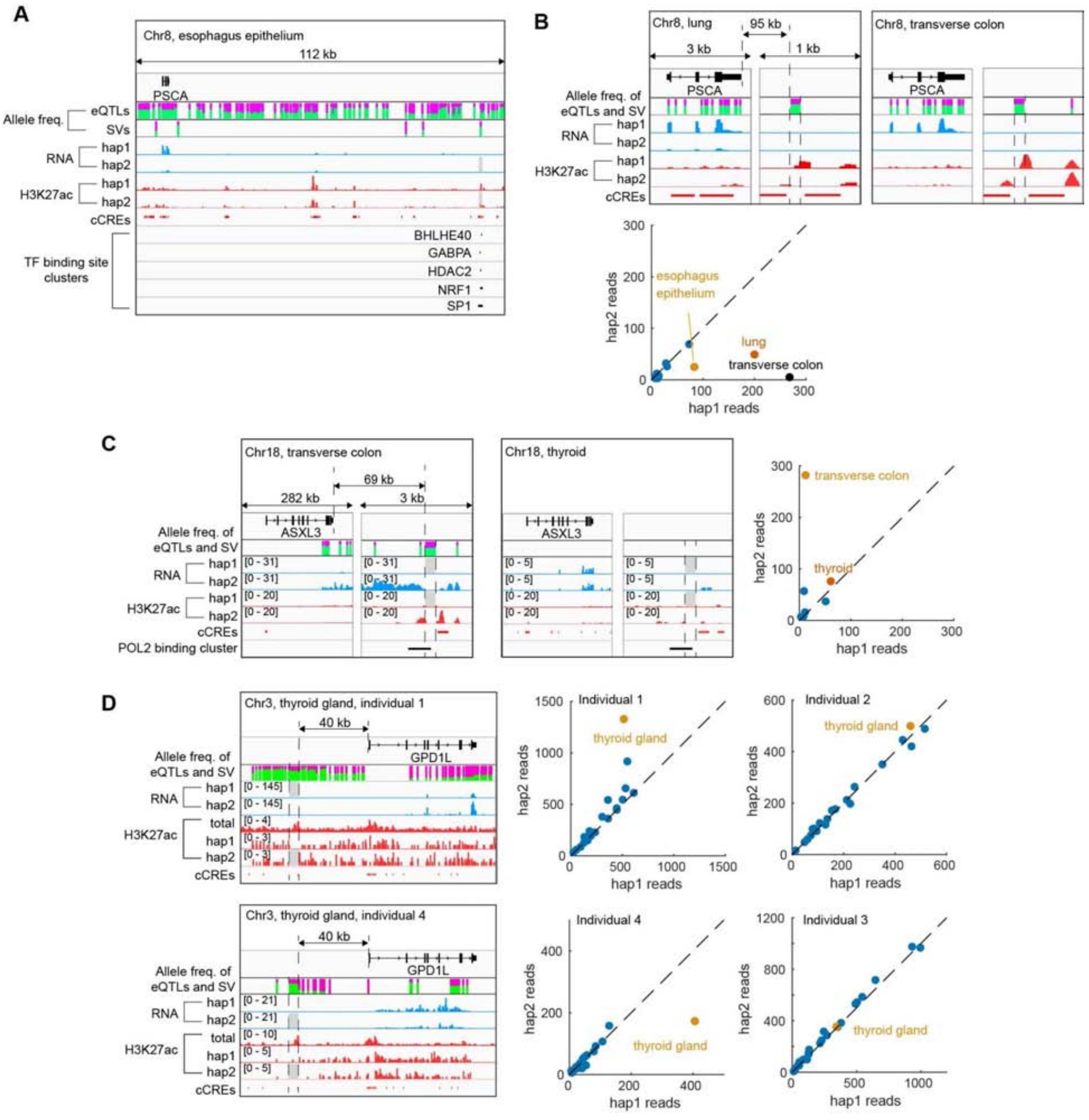

**Figure S3.2e. SVs potentially linked to eQTLs**

Panels **(A)** and **(B)** are shadow figures of Figure 4E. **(A)** This panel is the same as Figure 4E, but shows a panoramic view near the gene *PSCA*, including additional eQTLs that are compatible with the ASE of *PSCA*. The allele frequencies of the hap2 alleles at these eQTL sites are shown as the heights of the green bars. SVs near *PSCA* and their allele frequencies are also shown. The left four SVs are deletions in hap1, and the rightmost SV is the hap2 deletion shown in Figure 4E. cCREs and TF binding sites that can potentially be disrupted by the deletion of interest are shown. **(B)** *PSCA* also has a lower expression of hap2 in the lung and transverse colon of individual 3. In both tissues, the deletion has an allele frequency similar to that of some of the tissue-specific eQTLs compatible with the ASE of *PSCA*; moreover, this deletion appears to remove an H3K27ac peak in hap2, potentially causing the reduced

expression of *PSCA*. Imbalance in the ASE of *PSCA* appears to be restricted to three tissues shown in **(A)** and **(B)**. **(C)** Another example of a deletion that may be linked with compatible eQTLs of *ASXL3*. In the transverse colon of individual 2, *ASXL3* has lower expression in hap1. The relevant deletion occurs in hap1 and appears to disrupt H3K27ac and cCREs near the gene. Note that the H3K27ac levels at this cCRE and the expression levels of *PSCA* are both lower in the thyroid than in transverse colon, suggesting an association between the activity of this cCRE with *PSCA* expression. Imbalance in the ASE of *ASXL3* appears to be tissue specific. **(D)** A known SV-eQTL of *GPD1L* (Chiang *et al.*, 2017)
in the thyroid gland. Individuals 1 and 4 are heterozygous for this deletion, but the former has it on haplotype 1 and the latter has it on haplotype 2. As shown in the signal tracks, hap1 of individual 1 and hap2 of individual 2 show lower *GPD1L* expression than the other haplotype in the respective individual. There appears to be an active enhancer 40 kb upstream of *GPD1L*, as indicated by the total H3K27ac ChIP-seq signal (fold-change of the total reads from both haplotypes over the control) and by the locations of the active cCREs in the thyroid gland. This enhancer is removed by the deletion, potentially reducing the expression levels of *GPD1L* in the corresponding haplotype. The effect of the deletion is not obvious from the hap-specific H3K27ac ChIP-seq reads. This is likely because the region does not have enough SNVs, which are required to map ChIP-seq reads to both haplotypes. We note that individuals 2 and 3 are homozygous for this deletion, potentially explaining the lack of ASE of *GPD1L* in these two individuals.

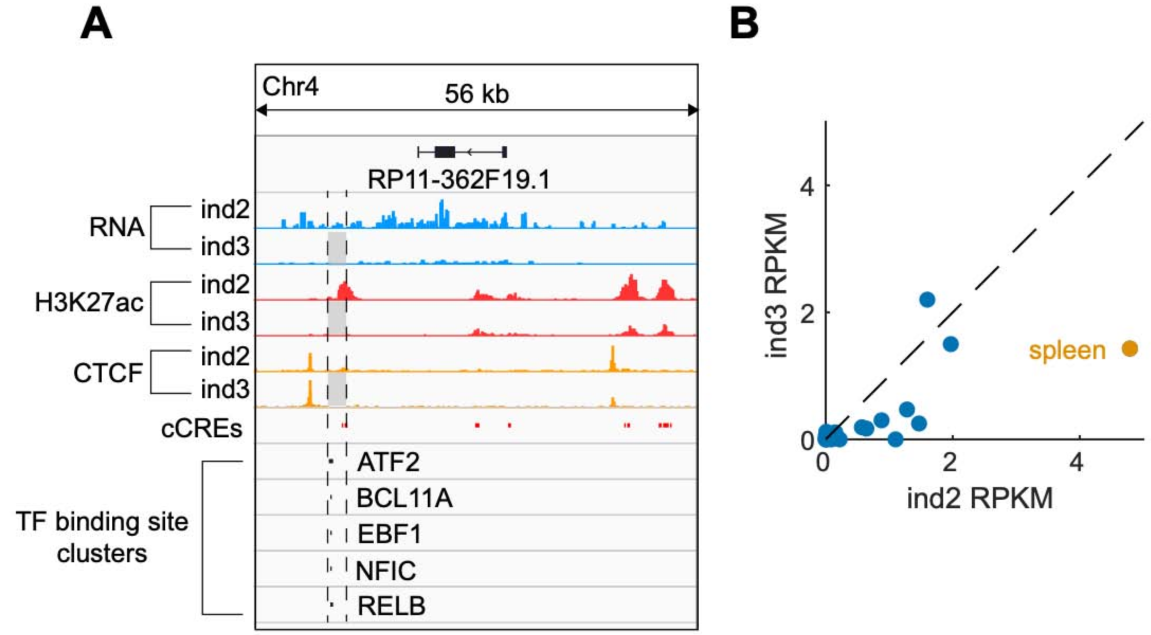

**Figure S3.2f. Shadow figure associated with Figure 4F**
**(A)** Similar to Figure 4F, the deletion in hap2 can disrupt spleen-specific cCREs and the binding of several TFs. **(B)** In multiple tissues, *RP11-362F19.1* has lower expression in individual 3 than in individual 2, suggesting that the deletion may have a global effect on the expression of this gene.

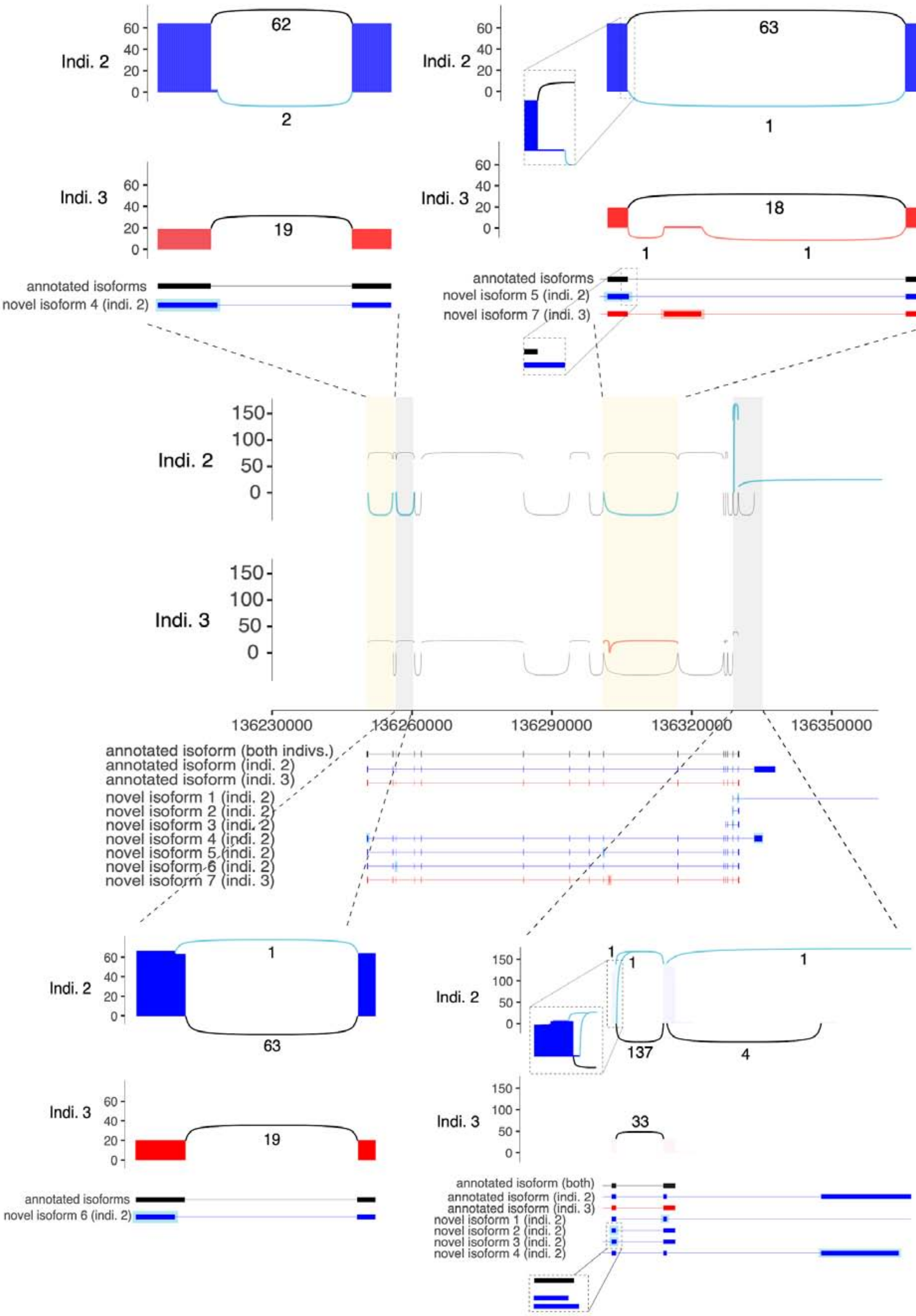

Indi. 2
62
2
Indi. 3
19
annotated isoforms
novel isoform 4 (indi. 2)
Indi. 2
63
1
Indi. 3
18
1
1
annotated isoforms
novel isoform 5 (indi. 2)
novel isoform 7 (indi. 3)
Indi. 2
Indi. 3
136230000
136260000
136290000
136320000
136350000
annotated isoform (both indivs.)
annotated isoform (indi. 2)
annotated isoform (indi. 3)
novel isoform 1 (indi. 2)
novel isoform 2 (indi. 2)
novel isoform 3 (indi. 2)
novel isoform 4 (indi. 2)
novel isoform 5 (indi. 2)
novel isoform 6 (indi. 2)
novel isoform 7 (indi. 3)
Indi. 2
1
63
Indi. 3
19
annotated isoforms
novel isoform 6 (indi. 2)
Indi. 2
1
1
1
137
4
Indi. 3
33
annotated isoform (both)
annotated isoform (indi. 2)
annotated isoform (indi. 3)
novel isoform 1 (indi. 2)
novel isoform 2 (indi. 2)
novel isoform 3 (indi. 2)
novel isoform 4 (indi. 2)

**Figure S3.2g. Novel splicing variants of *PCCB***

Shadow figure for Figure 4G. Sashimi plot and exonic structure representation of the *PCCB* isoforms expressed in individuals 2 (blue) and 3 (red) in adrenal gland and heart left ventricle tissues, respectively. The central panel provides a representation of the whole gene. In the sashimi plot, exons are represented by vertical lines either in blue (Ind. 2) or red (Ind. 3). Splicing connections of annotated isoforms are represented by black arcs, while novel connections observed in a specific individual are color-coded (magnifications of specific regions are provided, as well as the number of reads supporting each connection). The exonic structure of annotated and novel isoforms are reported at the bottom. The black isoform is expressed in both individuals, while those expressed in only one individual are color-coded. Annotated and novel isoforms were retrieved, for each individual, using Swan (Reese and Mortazavi, 2021). Specifically, a Swan gene report was generated for each individual by inputting transcriptome annotation and quantification files available, from long-read RNA-seq experiments, in the ENCODE portal (https://www.encodeproject.org/). These plots were obtained using ggashimi (Garrido-Martin et al., 2018).

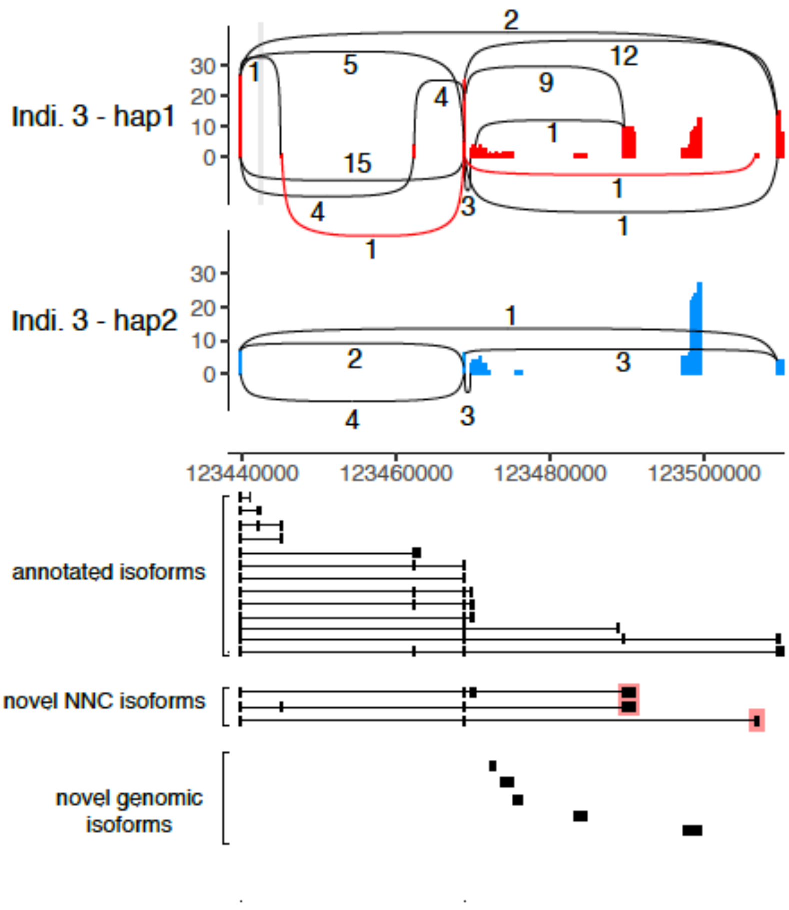

**Figure S3.2h. Novel splicing variants of *TRDN-AS1***
Sashimi plot and exonic structure representation of the lncRNA *TRDN-AS1* isoforms in individual 3 in the heart left ventricle. This gene carries a heterozygous deletion on haplotype 1 (highlighted in gray) and shows ASE in the right atrium auricular region (with hap1 being more highly expressed than hap2). For the sashimi representation, reads available from long-read RNA-seq experiments (see ENCODE portal) were phased to the two haplotypes using heterozygous SNVs that overlap the gene's exons. Read phasing was performed with ASCIIGenome (https://github.com/dariober/ASCIIGenome/) (Beraldi et al., 2017). Long-read RNA-seq reads show consistently higher expression of hap1 compared with hap2. Moreover, reads mapping to hap1 give rise to two novel splicing junctions (represented by red arcs) and two novel exons (highlighted in red in the exonic structure representation at the bottom). Annotated and novel isoforms were retrieved for each individual using SWAN (Reese and Mortazavi, 2021). Specifically, a SWAN gene report was generated for individual 3 by inputting transcriptome annotation and quantification files available from long-read RNA-seq experiments in the ENCODE portal. Only novel not in catalog (NNC) and genomic isoforms are shown. These plots were obtained using ggashimi (Garrido-Martin *et al.*, 2018).

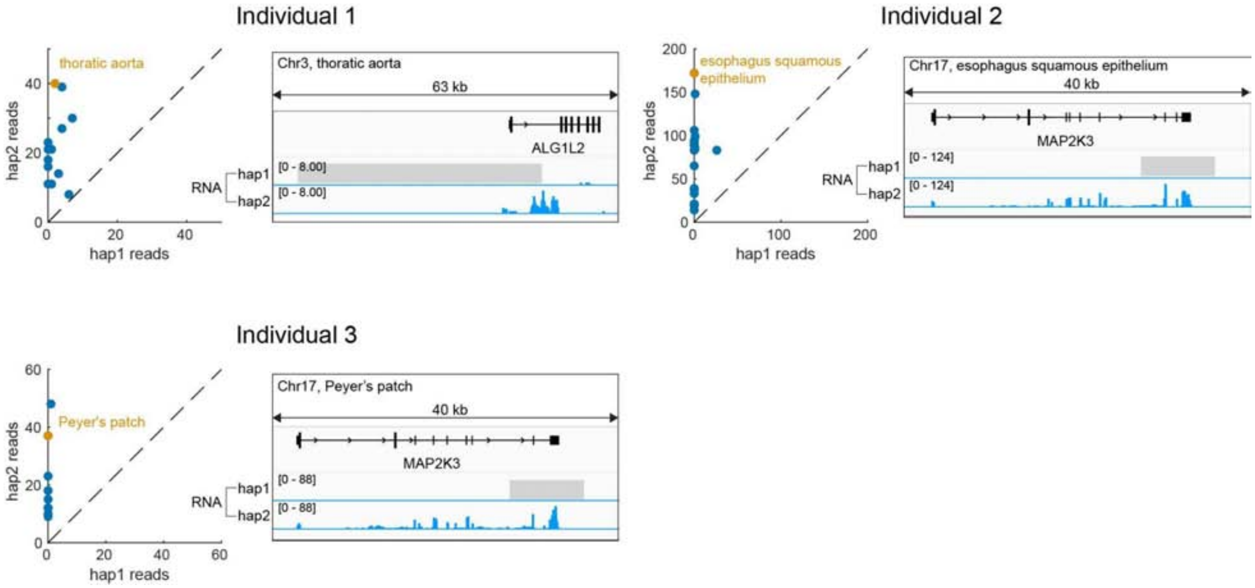

**Figure S3.2i. Null alleles potentially caused by deletion of entire exons.**
These examples show genes whose expression comes almost exclusively from haplotype 2 in multiple tissues of a given individual. Further analysis revealed that each example contains a deletion in haplotype 1 that removes multiple exons of the given gene in the given individual. We did not find null alleles associated with SV deletions in individual 4.

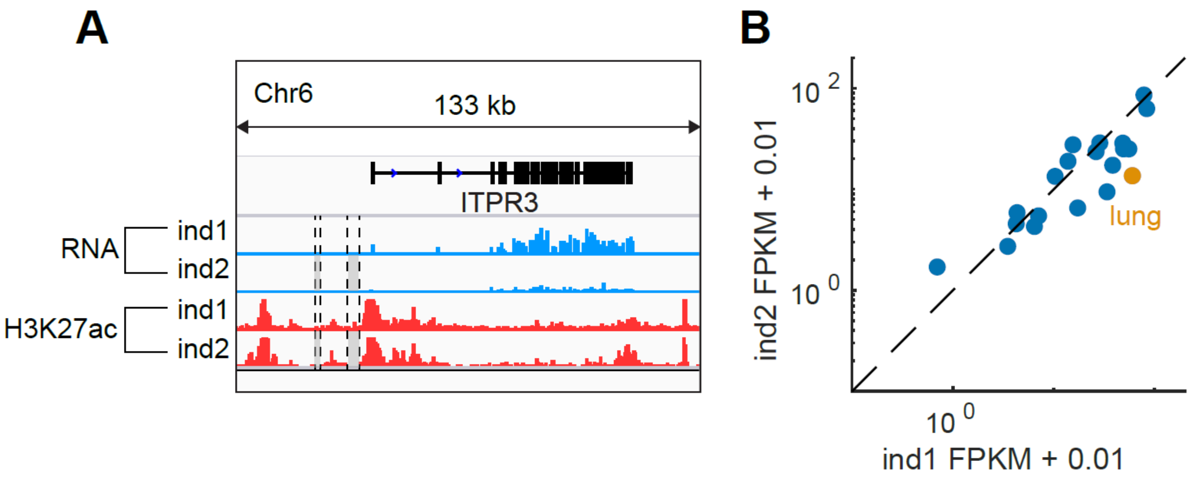

**Figure S3.2j. Homozygous deletions in individual 2 but not individual 1.**
**(A)** Two homozygous deletions upstream of *ITPR3* are found in individual 2 (the two variants are very close and are shown together by a single grey box), but are missing in individual 1. The deletions knock out part of the H3K27ac peak near the TSS of ITPR3, potentially reducing the gene's expression in the lung of individual 2 compared with individual 1. **(B)** Across multiple tissues, the expression levels of ITPR3 appear to be lower in individual 2 than individual 1.

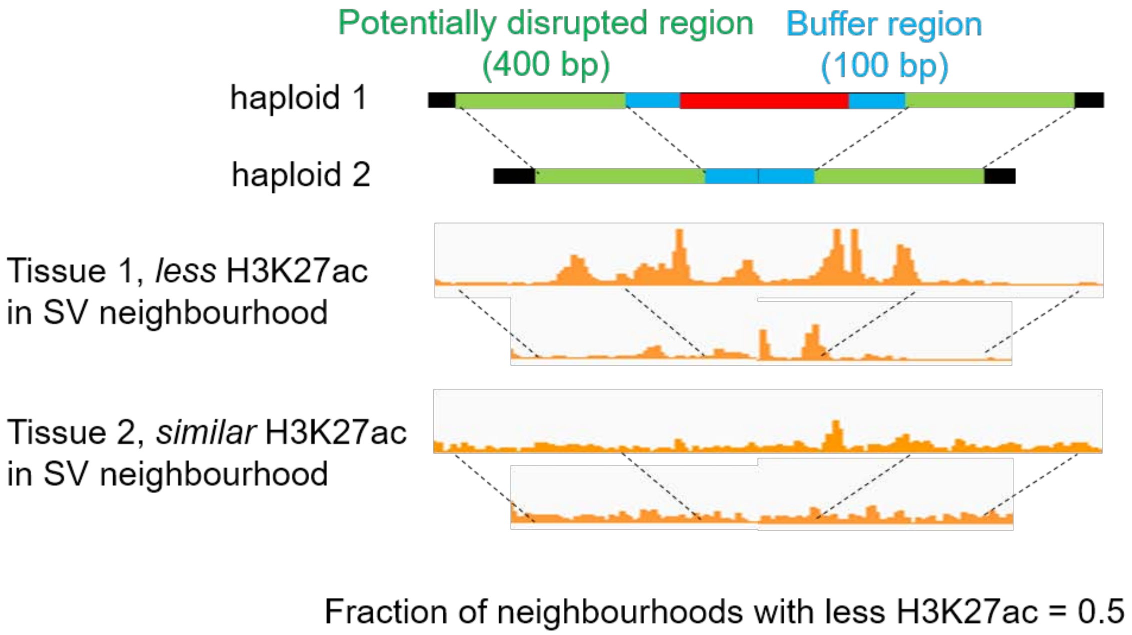

**Figure S3.3a. Calculating changes in the chromatin state in the SV neighborhood**
Using H3K27ac levels as an example, haploid 1 of individual 1 carries a deletion (red bar) while haploid 2 is wild type at the same locus; therefore, we compared the chromatin states in the two green regions between the two haploids. In tissue 1, the H3K27ac levels in the green region are lower in haploid 1, whereas the H3K27ac levels in tissue 2 are similar in both haploids. Therefore, only half of the neighborhoods of this deletion show a reduction in H3K27ac levels. Similar analyses can be performed between two individuals by substituting the two haploids with two individuals.

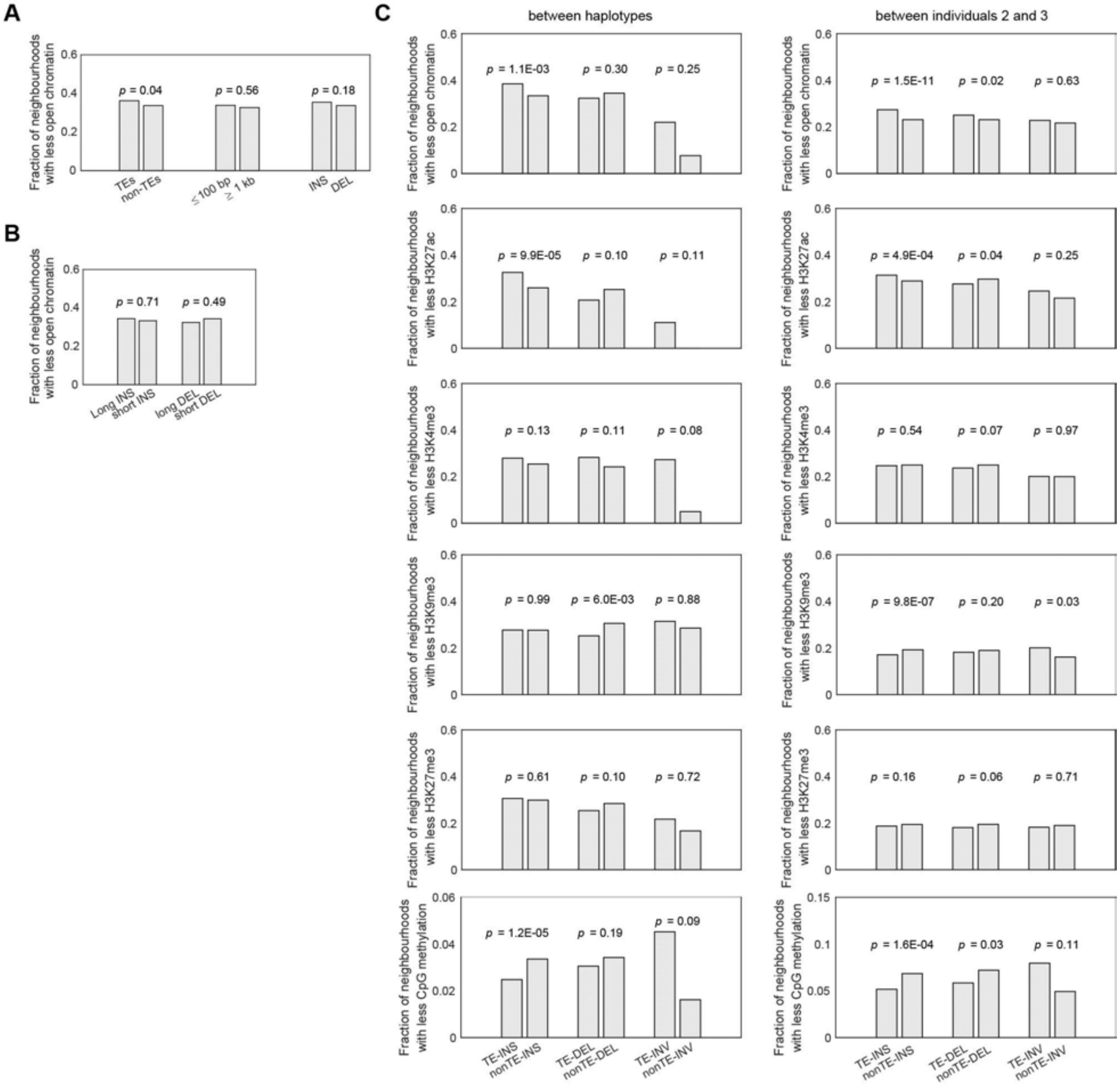

**Figure S3.3b. Changes in the chromatin state of SV neighborhoods**
Similar to Figure 4C, we investigated whether the presence of an SV changes the chromatin state of nearby regions and whether these changes are associated with different characteristics of the SVs. **(A)** The reduction in the chromatin openness near SVs does not differ by SV length or SV type, respectively. **(B)** Nor does it differ between long (> 1 kb) and short (< 100 bp) SV insertions. **(C)** Changes in other chromatin states near SVs. Left panels: changes in the chromatin states near heterozygous SVs in all four individuals. The changes were calculated by comparing the chromatin states between two haploids of the same individual. Right panels: changes in the chromatin states near SVs that are only present in either individual 2 or 3 (but not both). The changes were calculated by comparing the chromatin states between two individuals. p-values were calculated from the $\chi 2$ test.

#### Figure S4. Supp. Figures for Main Text Section “Generalized Application #1: Predicting AS ChIP-seq Activity from Nucleotide Sequence”

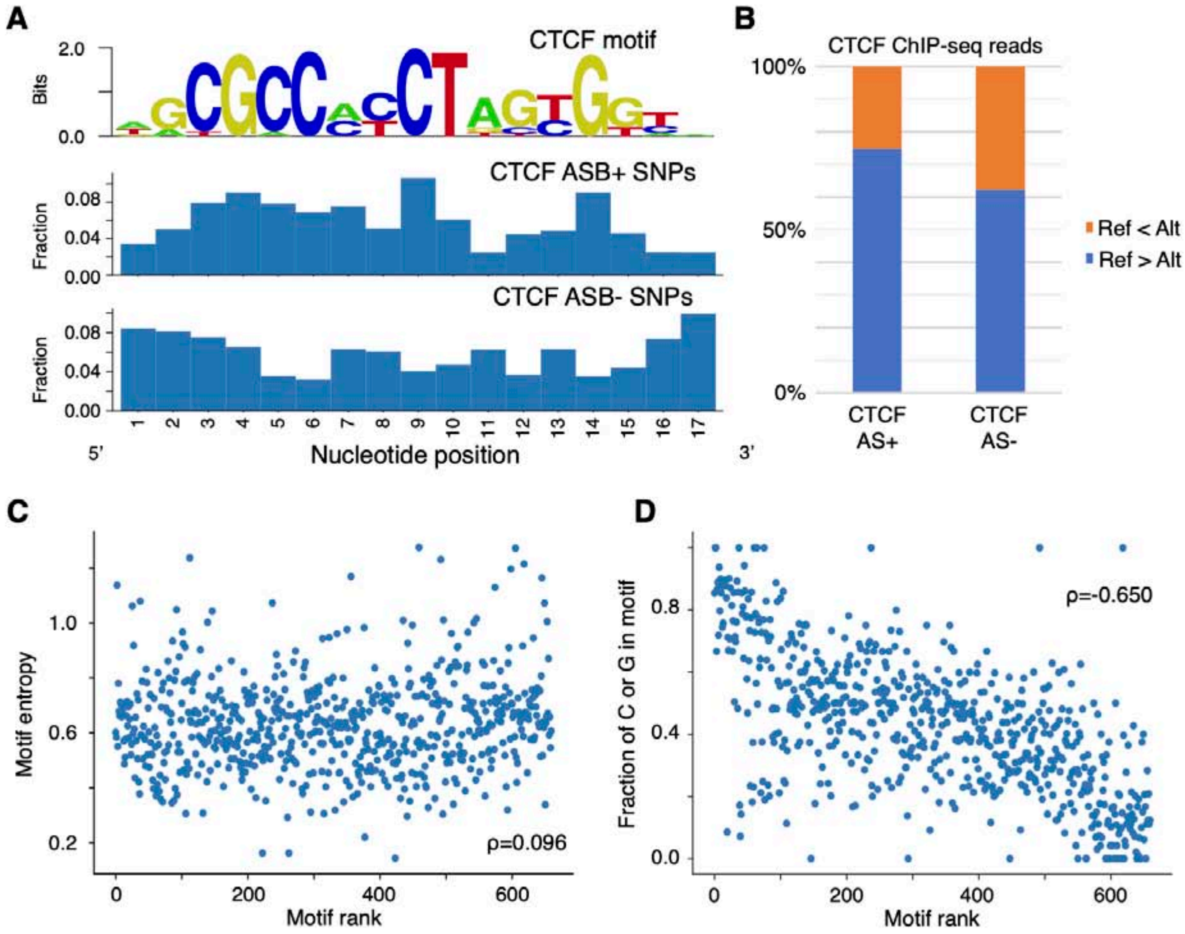

**Figure S4.1a. AS enriched motif ranking**

(A) Similar to Figure 5A, among the accessible SNPs from CTCF ChIP-seq, AS SNPs occur more frequently in the key positions of the CTCF motif. (B) For all CTCF accessible SNPs intersecting with the CTCF motif, >70% of the AS SNPs have more reads in the reference allele than the alternative allele. This number is ~60% for non-AS SNPs, indicating that the observation of AS is likely to be caused by the disruption of the motifs. (C) Motif ranking based on the enrichment of any ChIP-seq AS SNPs is not correlated with the entropy of the motifs. (D) The same ranking is negatively correlated with the GC content, determined by the number of positions where C or G is the most frequent base, divided by the motif length.

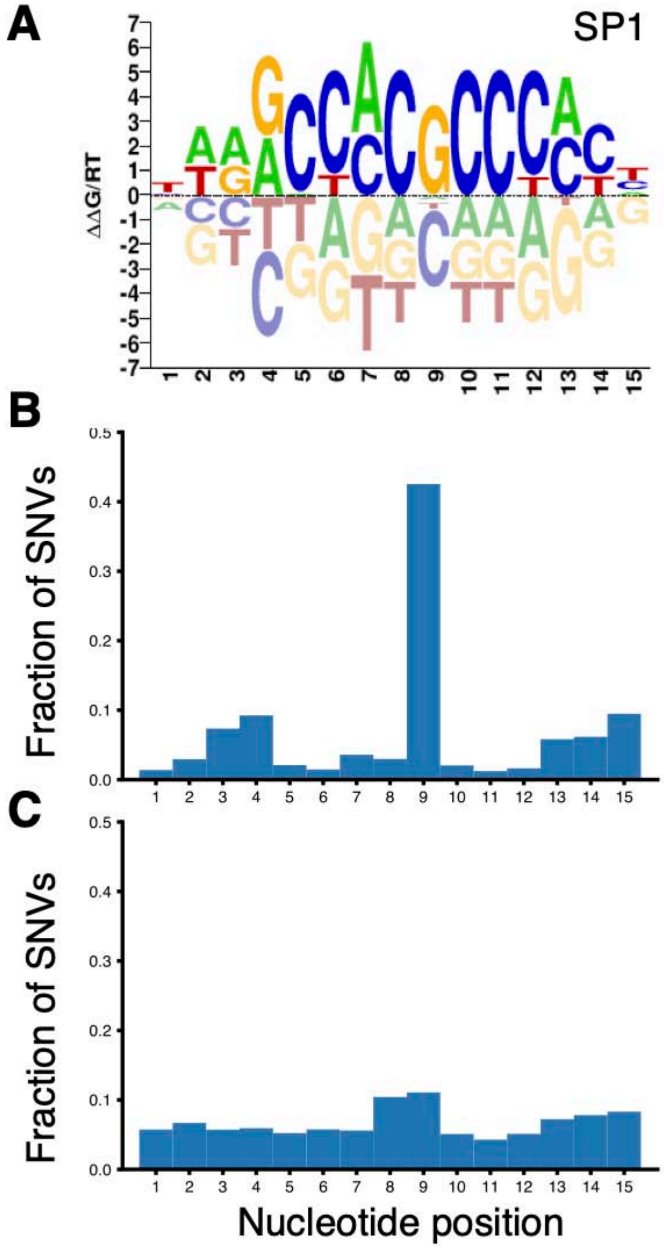

**Figure S4.1b. Fraction of AS/non-AS SNVs overlapping with the motif**

(A) The motif of transcription factor SP1. The logo is downloaded from the Cis-BP database. (B) Among all AS SNVs that overlap with the SP1 motif, >40% occur at position 9 of the motif. (C) For non-AS SNVs, they occur relatively randomly across all the positions of the motif.

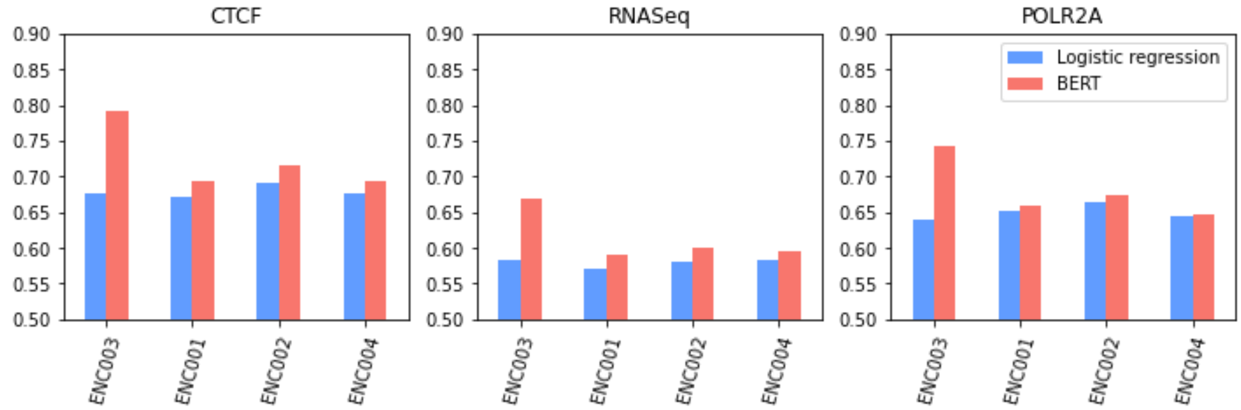

**Figure S4.2a. Performance of trained allelic effect prediction models**
"Logistic regression" results were derived from simple logistic regression on the dna2vec embedding of the input sequence; "BERT" results were derived from the fine-tuned DNABERT model. Both models were trained on SNPs from individual 3, and the results are reported for the validation sets from all four individuals.

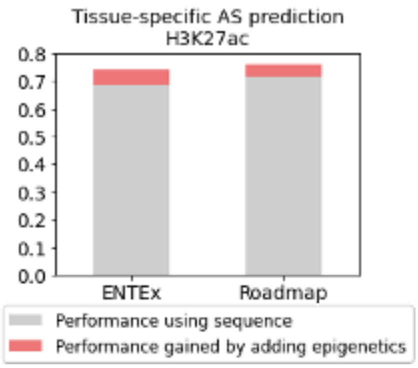

**Figure S4.2b. Tissue-specific performance in H3K27ac (with Roadmap external validation)**

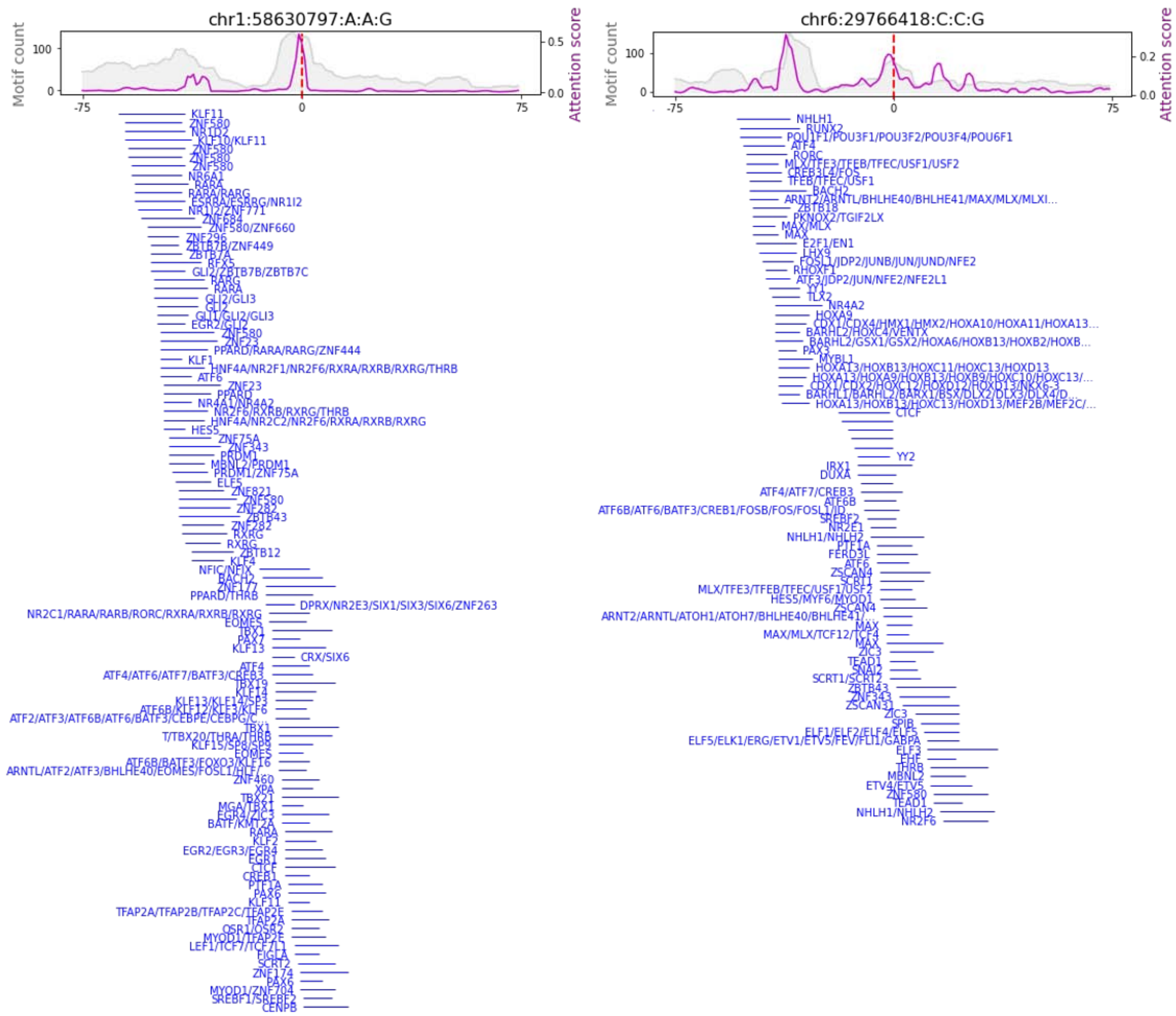

**Figure S4.2c. Additional examples of attention patterns learned by the model**

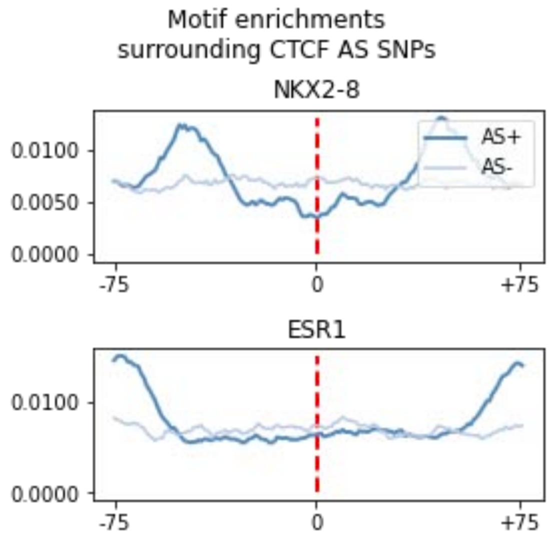

**Figure S4.2d. Motifs that peak in the proximity of the AS CTCF SNPs**

#### Figure S5. Supp. Figures for Main Text Section “Generalized Application #2: Models Interrelating AS Activity at Promoters and Genes”

| RF model on protein-coding genes | | | Positive Size | Negative Size | Accuracy | Sensitivity | Specificity | MCC | F1 | Precision | ROC_AUC | PR_AUC |
|---|---|---|---|---|---|---|---|---|---|---|---|---|
| H3K27ac | No cross-individual | Feature 1+2+3 | 1224 | 41005 | 0.8080 | 0.7895 | 0.8265 | 0.6164 | 0.8043 | 0.8198 | 0.8797 | 0.8967 |
| | | Feature 1+2+3+4 | 1323 | 45889 | 0.8249 | 0.7926 | 0.8571 | 0.6512 | 0.8191 | 0.8474 | 0.8922 | 0.9088 |
| | Cross-individual testing | Enc1 | 424 | 12489 | 0.8518 | 0.6527 | 0.8586 | 0.2498 | 0.2246 | 0.1357 | 0.8137 | 0.3335 |
| | | Enc2 | 277 | 10373 | 0.8756 | 0.5025 | 0.8855 | 0.1872 | 0.1737 | 0.1051 | 0.7574 | 0.1969 |
| | | Enc3 | 275 | 11629 | 0.8549 | 0.6041 | 0.8608 | 0.1958 | 0.1615 | 0.0932 | 0.8171 | 0.1888 |
| | | Enc4 | 351 | 11388 | 0.8404 | 0.7504 | 0.8432 | 0.2665 | 0.2197 | 0.1287 | 0.8479 | 0.3665 |
| | Independent testing | STL002_3 | 44 | 1070 | 0.7239 | 0.8726 | 0.7177 | 0.2504 | 0.2006 | 0.1135 | 0.8829 | 0.2842 |
| H3K4me3 | No cross-individual | Feature 1+2+3 | 1607 | 42579 | 0.7731 | 0.7449 | 0.8014 | 0.5472 | 0.7665 | 0.7896 | 0.8466 | 0.8721 |
| | | Feature 1+2+3+4 | 1738 | 48343 | 0.7995 | 0.7594 | 0.8397 | 0.6011 | 0.7912 | 0.8257 | 0.8686 | 0.8899 |
| | Cross-individual testing | Enc1 | 593 | 14424 | 0.8299 | 0.6428 | 0.8375 | 0.2429 | 0.2300 | 0.1401 | 0.7983 | 0.3510 |
| | | Enc2 | 330 | 10799 | 0.8563 | 0.5304 | 0.8662 | 0.1909 | 0.1797 | 0.1082 | 0.7703 | 0.1968 |
| | | Enc3 | 404 | 12503 | 0.8388 | 0.5878 | 0.8469 | 0.2031 | 0.1859 | 0.1104 | 0.8032 | 0.2126 |
| | | Enc4 | 416 | 10610 | 0.8216 | 0.7285 | 0.8252 | 0.2661 | 0.2357 | 0.1406 | 0.8285 | 0.3661 |
| | Independent testing | STL002_3 | 23 | 51 | 0.6854 | 0.8872 | 0.6771 | 0.2317 | 0.1827 | 0.1019 | 0.8799 | 0.3082 |
| CTCF | No cross-individual | Feature 1+2+3 | 384 | 25040 | 0.7058 | 0.7017 | 0.7098 | 0.4119 | 0.7044 | 0.7077 | 0.7759 | 0.7903 |
| | | Feature 1+2+3+4 | 430 | 27755 | 0.7236 | 0.6978 | 0.7494 | 0.4480 | 0.7162 | 0.7359 | 0.7897 | 0.8156 |
| | Cross-individual testing | Enc1 | 64 | 6509 | 0.7491 | 0.6367 | 0.7502 | 0.0875 | 0.0472 | 0.0245 | 0.7661 | 0.0552 |
| | | Enc2 | 78 | 5907 | 0.8007 | 0.2885 | 0.8075 | 0.0276 | 0.0364 | 0.0194 | 0.5996 | 0.0207 |
| | | Enc3 | 152 | 7298 | 0.7364 | 0.5741 | 0.7398 | 0.1005 | 0.0817 | 0.0440 | 0.7120 | 0.0775 |
| | | Enc4 | 136 | 8033 | 0.7343 | 0.5804 | 0.7369 | 0.0918 | 0.0680 | 0.0361 | 0.6878 | 0.0674 |
| POLR2A | No cross-individual | Feature 1+2+3 | 404 | 21092 | 0.7014 | 0.6884 | 0.7144 | 0.4032 | 0.6974 | 0.7070 | 0.7679 | 0.7948 |
| | | Feature 1+2+3+4 | 424 | 22436 | 0.7495 | 0.7192 | 0.7799 | 0.5002 | 0.7416 | 0.7659 | 0.8199 | 0.8445 |
| | Cross-individual testing | Enc1 | 54 | 4630 | 0.7878 | 0.5570 | 0.7905 | 0.0906 | 0.0571 | 0.0301 | 0.7435 | 0.0760 |
| | | Enc2 | 55 | 2914 | 0.8208 | 0.3738 | 0.8292 | 0.0722 | 0.0719 | 0.0398 | 0.6792 | 0.0664 |
| | | Enc3 | 145 | 8195 | 0.7831 | 0.6674 | 0.7852 | 0.1424 | 0.0969 | 0.0523 | 0.7945 | 0.1095 |
| | | Enc4 | 170 | 6699 | 0.7724 | 0.6288 | 0.7760 | 0.1489 | 0.1207 | 0.0668 | 0.7554 | 0.1402 |
| POLR2A phosphoS5 | No cross-individual | Feature 1+2+3 | 235 | 10153 | 0.6611 | 0.6445 | 0.6776 | 0.3226 | 0.6552 | 0.6669 | 0.7204 | 0.7537 |
| | | Feature 1+2+3+4 | 241 | 10538 | 0.7172 | 0.6840 | 0.7505 | 0.4357 | 0.7074 | 0.7330 | 0.7856 | 0.8107 |
| | Cross-individual testing | Enc1 | 76 | 3400 | 0.7412 | 0.6102 | 0.7441 | 0.1178 | 0.0937 | 0.0508 | 0.7208 | 0.1163 |
| | | Enc2 | 15 | 601 | 0.7818 | 0.5721 | 0.7871 | 0.1335 | 0.1136 | 0.0631 | 0.7483 | 0.1364 |
| | | Enc3 | 57 | 3334 | 0.7778 | 0.4835 | 0.7829 | 0.0827 | 0.0685 | 0.0369 | 0.6904 | 0.0819 |
| | | Enc4 | 93 | 3207 | 0.7552 | 0.6024 | 0.7597 | 0.1386 | 0.1222 | 0.0680 | 0.7307 | 0.1688 |

**Figure S5.1a. Performance validation of models predicting allele-specific bound promoters for each assay**

Models were trained separately for each assay with enough training data, including H3K27ac, H3K4me3, CTCF, POLR2A, and POLR2AphosphoS5. We targeted protein-coding genes with exactly one hetSNV in the promoter region ( ±1 kb of the TSS), and the ASB states of the hetSNV and the promoter were examined for consistency. For no cross-individual training/testing, sub-models were trained and tested with balanced data composed of the same set of positives and different subsets of negatives. For each sub-model, a five-fold cross validation strategy was used. Different sets of features (see Figure S5.1b for feature description) were tested for improved model performance. For cross-individual testing, for each of the four EN-TEx individuals, sub-models were trained with balanced data from the other three individuals and tested on the imbalanced data from the targeting individual. For independent testing, sub-models were trained with balanced EN-TEx data, including the four individuals, and tested on the imbalanced integrated data from Roadmap individuals STL002 and STL003. For all the metrics, the average performance of sub-models is shown in the table.

| Association of features with ASB event | | Positive_mean | Negative_mean | t.test p-value | R2 score | Assay | RF feature importance |
|---|---|---|---|---|---|---|---|
| Feature 1 | Top100TF_motifSum_w_hetSNV | 43 | 4 | < 2.2E-16 | 0.9421 | H3K27ac | 0.3298 |
| | | 42 | 4 | < 2.2E-16 | 0.9495 | H3K4me3 | 0.3127 |
| | | 16 | 4 | < 2.2E-16 | 0.8736 | CTCF | 0.1929 |
| | | 24 | 4 | < 2.2E-16 | 0.8895 | POLR2A | 0.2359 |
| | | 26 | 4 | 1.156E-15 | 0.8469 | POLR2AphosphoS5 | 0.2442 |
| Feature 2 | Top100TF_motifSum_nearby_hetSNV | 91 | 49 | < 2.2E-16 | 0.8642 | H3K27ac | 0.2263 |
| | | 89 | 48 | < 2.2E-16 | 0.9042 | H3K4me3 | 0.2285 |
| | | 63 | 44 | 1.092E-08 | 0.3008 | CTCF | 0.2500 |
| | | 81 | 52 | < 2.2E-16 | 0.4165 | POLR2A | 0.2371 |
| | | 73 | 49 | 2.757E-09 | 0.4254 | POLR2AphosphoS5 | 0.2347 |
| Feature 3 | motifSum_distal_to_hetSNV | 910 | 775 | < 2.2E-16 | 0.8035 | H3K27ac | 0.2129 |
| | | 898 | 780 | < 2.2E-16 | 0.8067 | H3K4me3 | 0.2303 |
| | | 817 | 769 | 0.0004748 | 0.1409 | CTCF | 0.2677 |
| | | 833 | 769 | 5.254E-06 | 0.3144 | POLR2A | 0.2201 |
| | | 854 | 778 | 0.0002361 | 0.3184 | POLR2AphosphoS5 | 0.2302 |
| Feature 4 | abs(RNAseq_hap1_allele_ratio -0.5) | 0.1403 | 0.0617 | < 2.2E-16 | 0.5884 | H3K27ac | 0.2311 |
| | | 0.1289 | 0.0633 | < 2.2E-16 | 0.5612 | H3K4me3 | 0.2285 |
| | | 0.1461 | 0.0625 | < 2.2E-16 | 0.5062 | CTCF | 0.2894 |
| | | 0.1638 | 0.0631 | < 2.2E-16 | 0.4754 | POLR2A | 0.3069 |
| | | 0.1423 | 0.0610 | 7.64E-14 | 0.5420 | POLR2AphosphoS5 | 0.2909 |

**Figure S5.1b. Association analysis between features and the promoter allele-specific binding events**

To predict the ASB state of a gene promoter, a random forest model was built using four features: three TF motif-based features of the promoter region and one AS expression feature of the gene. The three motif-based features are the total number of top 100 ranked TFs' motifs intersecting the hetSNV in the promoter, the total number of top 100 ranked TFs' motifs nearby (200 bp window centered on the hetSNV) but not intersecting the hetSNV in the promoter, and the total number of all 660 human TFs' motifs distal to the hetSNV in the promoter; the ASE feature is the imbalance ratio of gene expression between the haplotypes. Other features (including gene expression level, eQTL, all 660 non-ranked TF features in the promoter) were tested but proven not to be informative (Figure S5.1d). For each assay, we investigated the association of each feature with ASB promoters. Welch's two-sample t-test (two-tailed) was performed for each feature between ASB and non-ASB promoters. To test if each feature is positively correlated with ASB promoters, an R2 score was calculated (see Figure S5.1c). A larger R2 value indicates a stronger association between the targeted feature and the ASB event of the promoters. Datasets were shuffled 100 times and an averaged R2 score is shown in the table. For each feature, a random forest-based feature importance score is shown, which is the average of sub-random forest models trained from the balanced EN-TEx data composed of the same set of positives and different subsets of negatives. Protein-coding genes with one hetSNV in their promoters were used to train the model.

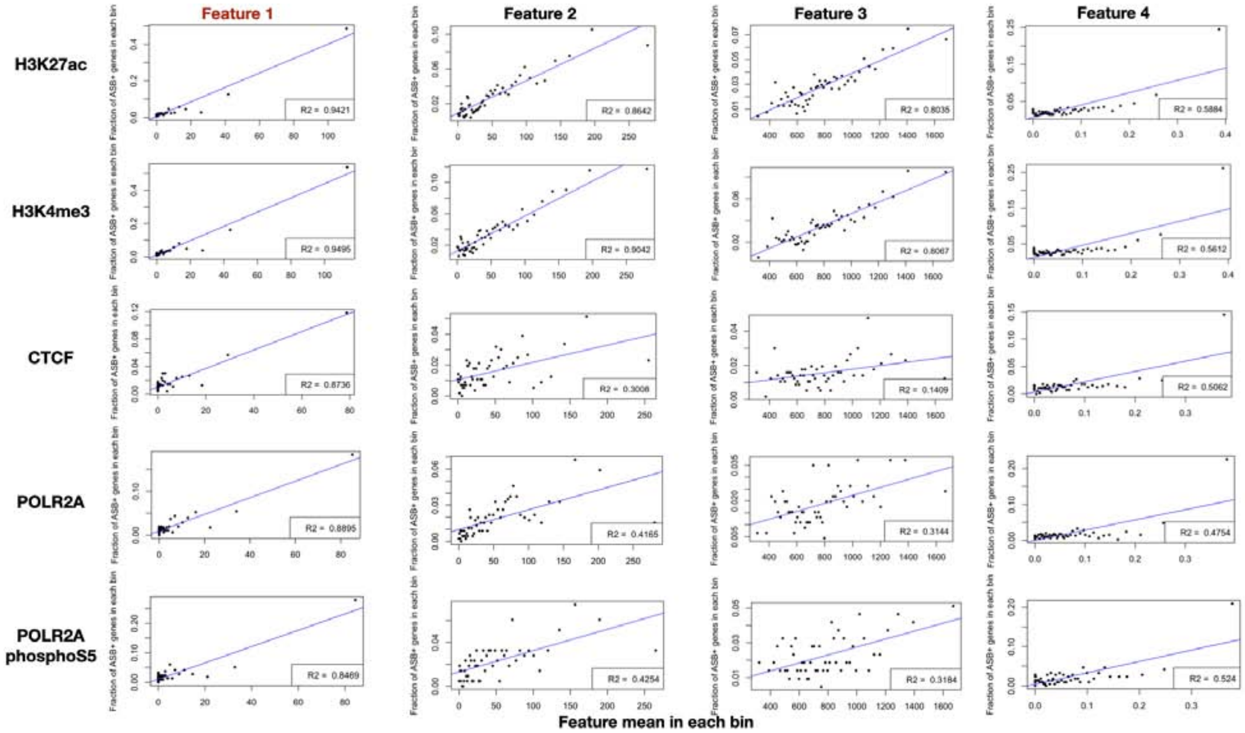

**Figure S5.1c. Scatter plots for the association analysis between features and the promoter ASB events**

First, for each assay, the whole dataset including all ASB and non-ASB promoters was ranked by the value of the targeting feature in an ascending order. Second, the ranked dataset was split into 50 bins. For each bin, we calculated the mean value of the targeting feature (x axis) and the ratio of ASB promoters in the bin (y axis), followed by a scatter plot shown above. For all assays, feature 1 (total number of top 100 ranked TFs' motifs intersecting the hetSNV in the promoter) showed the strongest association with ASB promoters.

| | |
|---|---|
| **ASE features** | P value to be ASE+ gene (Redundant to **Feature 4-** hap1 allelic ratio) |
| | Gene expression level in FPKM |
| **TF motif features** | All 660 TF motif sum in the promoter |
| | All 660 TF motif sum hit by the hetSNV in the promoter |
| | All 660 TF motif sum hit by accE hetSNV in the promoter |
| | Top 30 ranked TF motif sum hit by the hetSNV in the promoter |
| **hetSNV features** | If the hetSNV is GTEx eQTL |
| | eQTL effect (slope absolute value) |
| **Individual features** | The ratio of the promoter to be ASB+ in other EN-TEx individuals |

**Figure S5.1d. Features tested but not informative for the prediction of ASB promoters**
To build a machine learning model to predict ASB promoters, many other features were tested but not included in the final model since they did not improve the model performance.

**All gene types**

| | | | | | |
|---|---|---|---|---|---|
| H3K27ac | ASB+ | ASB- | No accB | Sum | Sum |
| ASE+ | 1402 | 8137 | 21384 | 30923 | 776187 |
| ASE- | 3855 | 225461 | 515948 | 745264 | |
| H3K27me3 | ASB+ | ASB- | No accB | Sum | Sum |
| ASE+ | 66 | 1697 | 29160 | 30923 | 776187 |
| ASE- | 133 | 29415 | 715716 | 745264 | |
| H3K36me3 | ASB+ | ASB- | No accB | Sum | Sum |
| ASE+ | 80 | 1147 | 29696 | 30923 | 776187 |
| ASE- | 73 | 19991 | 725200 | 745264 | |
| H3K4me1 | ASB+ | ASB- | No accB | Sum | Sum |
| ASE+ | 118 | 2085 | 28720 | 30923 | 776187 |
| ASE- | 278 | 64460 | 680526 | 745264 | |
| H3K4me3 | ASB+ | ASB- | No accB | Sum | Sum |
| ASE+ | 1400 | 8560 | 20963 | 30923 | 776187 |
| ASE- | 4527 | 229823 | 510914 | 745264 | |
| H3K9me3 | ASB+ | ASB- | No accB | Sum | Sum |
| ASE+ | 25 | 529 | 30369 | 30923 | 776187 |
| ASE- | 51 | 7994 | 737219 | 745264 | |
| CTCF | ASB+ | ASB- | No accB | Sum | Sum |
| ASE+ | 646 | 7505 | 22772 | 30923 | 776187 |
| ASE- | 1713 | 179835 | 563716 | 745264 | |
| EP300 | ASB+ | ASB- | No accB | Sum | Sum |
| ASE+ | 39 | 312 | 30572 | 30923 | 776187 |
| ASE- | 128 | 11016 | 734120 | 745264 | |
| POLR2A | ASB+ | ASB- | No accB | Sum | Sum |
| ASE+ | 872 | 4590 | 25461 | 30923 | 776187 |
| ASE- | 2450 | 133225 | 609589 | 745264 | |
| POLR2Apho | ASB+ | ASB- | No accB | Sum | Sum |
| ASE+ | 425 | 1878 | 28620 | 30923 | 776187 |
| ASE- | 1383 | 60965 | 682916 | 745264 | |
| ATAC | ASB+ | ASB- | No accB | Sum | Sum |
| ASE+ | 653 | 5048 | 25222 | 30923 | 776187 |
| ASE- | 2868 | 109180 | 633216 | 745264 | |
| DNase | ASB+ | ASB- | No accB | Sum | Sum |
| ASE+ | 1077 | 8071 | 21775 | 30923 | 776187 |
| ASE- | 4805 | 192779 | 547680 | 745264 | |

**Protein-coding genes**

| | | | | | |
|---|---|---|---|---|---|
| H3K27ac | ASB+ | ASB- | No accB | Sum | Sum |
| ASE+ | 1112 | 6032 | 15625 | 22769 | 631104 |
| ASE- | 3270 | 189499 | 415566 | 608335 | |
| H3K27me3 | ASB+ | ASB- | No accB | Sum | Sum |
| ASE+ | 35 | 1158 | 21576 | 22769 | 631104 |
| ASE- | 107 | 22842 | 585386 | 608335 | |
| H3K36me3 | ASB+ | ASB- | No accB | Sum | Sum |
| ASE+ | 18 | 695 | 22056 | 22769 | 631104 |
| ASE- | 40 | 13352 | 594943 | 608335 | |
| H3K4me1 | ASB+ | ASB- | No accB | Sum | Sum |
| ASE+ | 77 | 1556 | 21136 | 22769 | 631104 |
| ASE- | 219 | 53777 | 554339 | 608335 | |
| H3K4me3 | ASB+ | ASB- | No accB | Sum | Sum |
| ASE+ | 1125 | 6491 | 15153 | 22769 | 631104 |
| ASE- | 3977 | 197771 | 406587 | 608335 | |
| H3K9me3 | ASB+ | ASB- | No accB | Sum | Sum |
| ASE+ | 7 | 293 | 22469 | 22769 | 631104 |
| ASE- | 29 | 5433 | 602873 | 608335 | |
| CTCF | ASB+ | ASB- | No accB | Sum | Sum |
| ASE+ | 476 | 5373 | 16920 | 22769 | 631104 |
| ASE- | 1434 | 147190 | 459711 | 608335 | |
| EP300 | ASB+ | ASB- | No accB | Sum | Sum |
| ASE+ | 28 | 234 | 22507 | 22769 | 631104 |
| ASE- | 100 | 9218 | 599017 | 608335 | |
| POLR2A | ASB+ | ASB- | No accB | Sum | Sum |
| ASE+ | 634 | 3374 | 18761 | 22769 | 631104 |
| ASE- | 2024 | 111066 | 495245 | 608335 | |
| POLR2Apho | ASB+ | ASB- | No accB | Sum | Sum |
| ASE+ | 309 | 1376 | 21084 | 22769 | 631104 |
| ASE- | 1115 | 50458 | 556762 | 608335 | |
| ATAC | ASB+ | ASB- | No accB | Sum | Sum |
| ASE+ | 481 | 3696 | 18592 | 22769 | 631104 |
| ASE- | 2438 | 89692 | 516205 | 608335 | |
| DNase | ASB+ | ASB- | No accB | Sum | Sum |
| ASE+ | 787 | 5824 | 16158 | 22769 | 631104 |
| ASE- | 4035 | 157411 | 446889 | 608335 | |

**Figure S5.1e. Contingency table for ASE genes and ASB promoters**

For each assay, the numbers of ASE genes with ASB promoters are shown. No accB represents promoters that are not accessible for the assay.

| Feature Type | Feature Description | | Tested Feature Number | Positive | Negative | ENTEx Performance (ROC_AUC \| PR_AUC) | |
|---|---|---|---|---|---|---|---|
| ASB promoter annotation based features | ASB+ = (0,0,1)<br>ASB - = (0,1,0)<br>non-accB = (1,0,0) | Protein-coding genes | 10*3 | 117 | 707 | **0.87** | **0.88** |
| | | Protein-coding genes | 12*3 | 237 | 1807 | **0.88** | **0.88** |
| | | Protein-coding genes, FPKM >1 | 12*3 | 158 | 1608 | **0.87** | **0.88** |
| | | Protein-coding genes having 1 hetSNV in promoter | 12*3-1 (H3K9me3 no ASB+) | 51 (overfit) | 1011 | 0.73 | 0.72 |
| | | Protein-coding genes having 1 hetSNV in promoter, FPKM>1 | 12*3-1 | 37 (overfit) | 913 | 0.73 | 0.73 |
| | | Protein-coding genes having 1 hetSNV in promoter + eQTL | 12*3 -1 +1 | 51 (overfit) | 1011 | 0.78 | 0.76 |
| | | Protein-coding genes having 1 hetSNV in promoter + eQTL + housekeeping | 12*3 -1 +1 +1 | 51 (overfit) | 1011 | 0.74 | 0.73 |

| Validation Data | Feature Description | | Available Feature Number | |
|---|---|---|---|---|
| STL002 | ASB+ = (0,0,1)<br>ASB - = (0,1,0)<br>non-accB = (1,0,0) | Protein-coding genes | 3 | H3K27ac/ H3K36me3/ H3K4me1 |
| STL003 | | | 6 | H3K27ac/ H3K27me3/ H3K36me3/ H3K4me1/ H3K4me3/ H3K9me3 |
| NA12878 | | | 5 | H3K4me3/ CTCF/ EP300/ POLR2A/ POLR2AphosphoS5 |

**Figure S5.1f. Features and performance of a model to predict ASE for a gene from ASB on the associated promoter**

In addition to the ‘ASB from ASE’ model (Supplement S5.1), we constructed a model to predict ASE from ASB (going in the "forward" direction). The features of this model are summarized in this table; here, we used epigenetics as opposed to sequence features (as we did in the "reverse" model in Figure S5.1b). As shown in the feature description, different gene sets were tested to train the model. Other features that were tested but not informative included whether the hetSNV in the promoter was eQTL or whether the gene was a housekeeping gene. Overall, the model performed well on the EN-TEx samples but we do not have enough validation data on STL002, STL003, and NA12878 to properly evaluate the model.

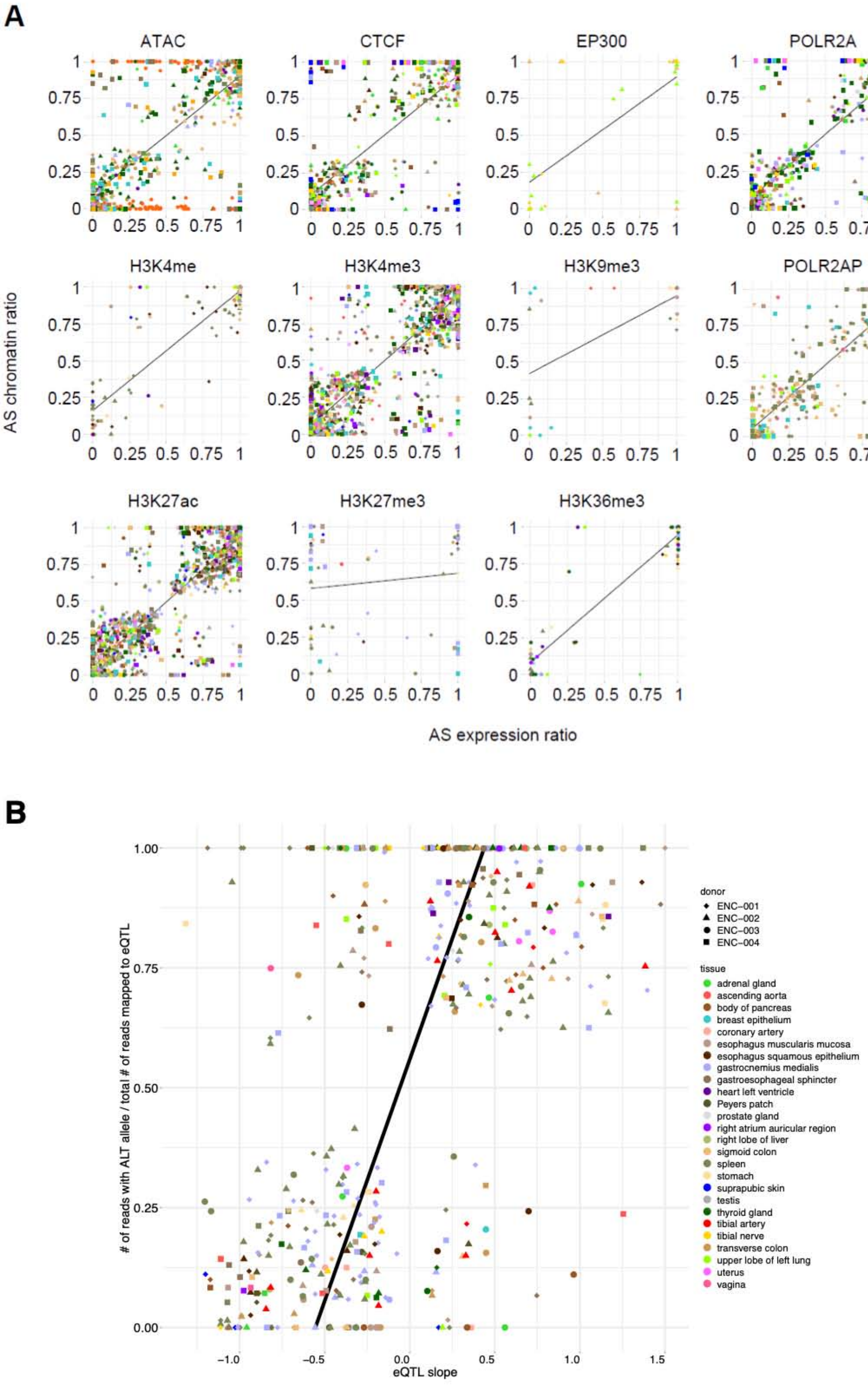
A
ATAC
CTCF
EP300
POLR2A
H3K4me
H3K4me3
H3K9me3
POLR2AP
H3K27ac
H3K27me3
H3K36me3
AS chromatin ratio
AS expression ratio
B
### of reads with ALT allele / total # of reads mapped to eQTL
eQTL slope
donor
ENC-001
ENC-002
ENC-003
ENC-004
tissue
adrenal gland
ascending aorta
body of pancreas
breast epithelium
coronary artery
esophagus muscularis mucosa
esophagus squamous epithelium
gastrocnemius medialis
gastroesophageal sphincter
heart left ventricle
Peyers patch
prostate gland
right atrium auricular region
right lobe of liver
sigmoid colon
spleen
stomach
suprapubic skin
testis
thyroid gland
tibial artery
tibial nerve
transverse colon
upper lobe of left lung
uterus
vagina

**C**

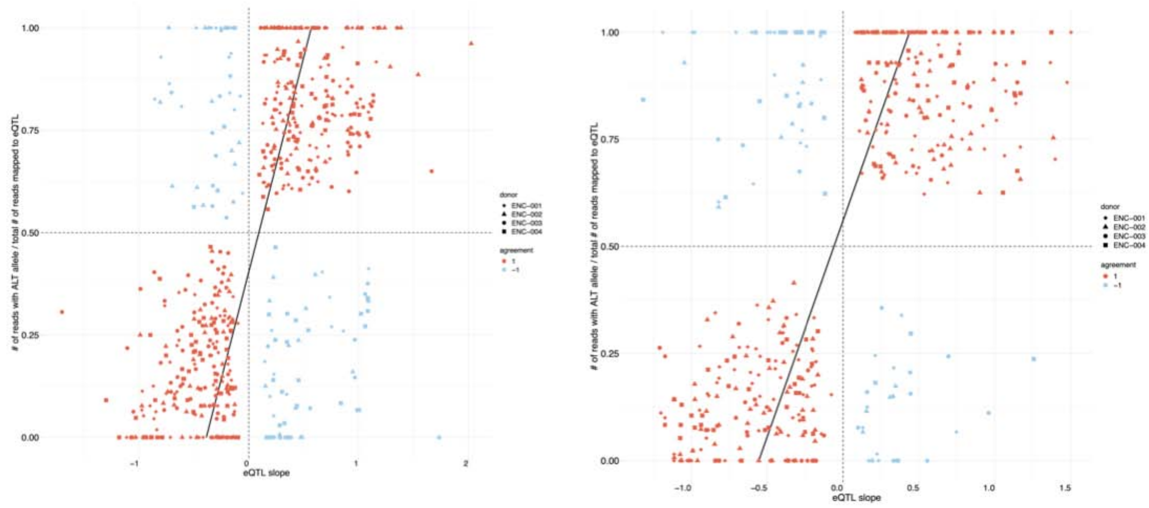

**Figure S5.2a. Compatibility between allelic events**

**(A)** Compatibility between AS chromatin state of the promoters (+/- 2 kb from the TSS) and the ASE of the corresponding genes. The AS chromatin ratio is the fraction of hap1 ChIP-seq reads among the total number of reads. The ASE ratio is the fraction of hap1 RNA-seq reads among the total number of reads. Each dot is a gene in a given tissue (marked by colors) and individual (marked by shape). See **(B)** for details regarding the colors and shapes. **(B)** AS H3K27ac at hetSNVs that are known GTEx eQTLs (GTEx Consortium, 2020). The y axis shows the fraction of H3K27ac ChIP-seq reads mapped to the alternative allele among the total reads mapped to either allele of an eQTL. **(C)** Same plots as Figure 7D (left) and **(B)** (right) but re-colored to show whether or not the eQTL effect (beta coefficient) and the ASE (left)/ASB (right) are compatible. A file with all the eQTL-AS binding events in the plots is available (File: AS_allhets_alt_allele_ratio_eqtl_intersect.tsv) with the compatibility represented by 1 or -1.

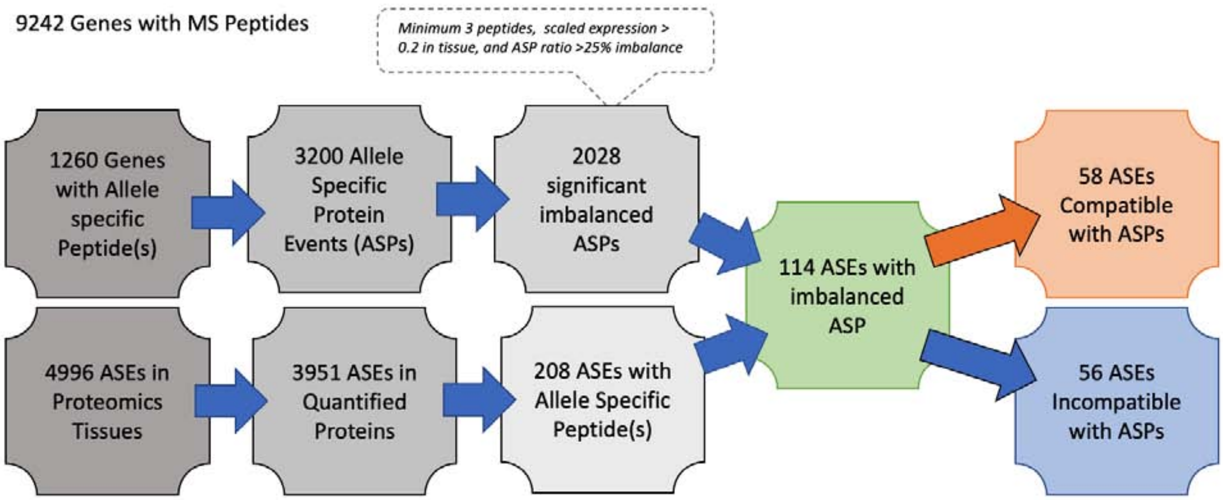

**Figure S5.2b. Flow chart of filtering and AS expression/AS Proteomics comparison**
Proteomics data were mapped at the gene level and filtered for proteins containing allele-specific peptides (ASPs). ASPs were calculated for each tissue in which allelic peptides were quantified. The ASP ratio was calculated as the summed peptide intensity of the first allele divided by the total specific to either allele. ASPs were filtered by the number of peptides, expression level, and ASP ratio. The p-value was calculated as 0.7 using z-scores.

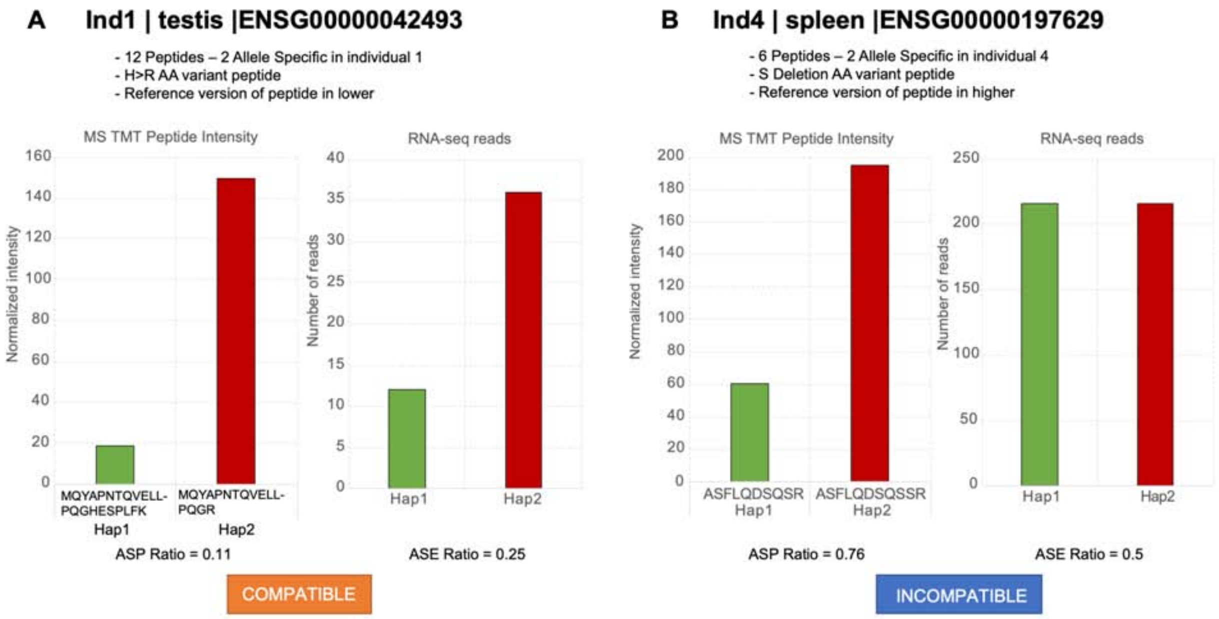

**Figure S5.2c. Compatibility between AS mRNA and AS peptide calculations**
**(A)** An example of a compatible ASP and ASE ratio. Both the proteomics and transcriptomics results indicate that the second allele is expressed at a higher level. **(B)** An example of an incompatible AS proteomics/ASE pairing. The transcriptomics result does not show any bias in the gene expression; however, the second allele is more highly expressed at the protein level.

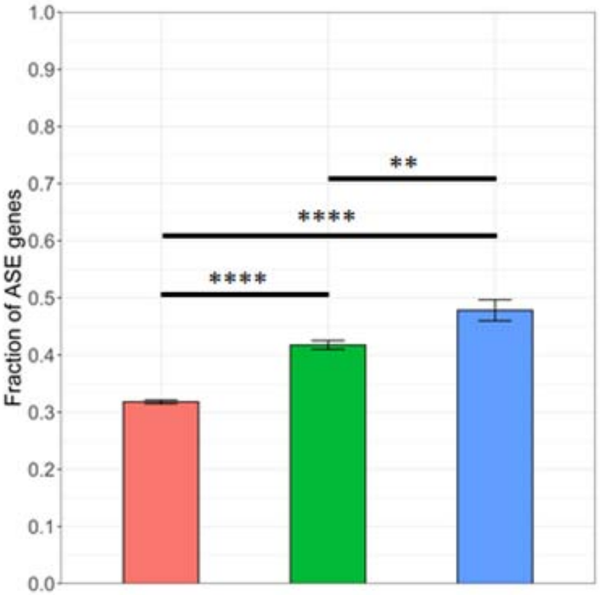

**Figure S5.2d. Enrichment of ASE genes near ASM promoters**
Enrichment of ASE genes near ASM promoters with (blue) ($\chi$2-test, OR = 1.96) or without (green) ($\chi$2-test, OR = 1.54) AS TF binding, relative to genes near ASM non-cCREs (red). ** $p < 0.01$, **** $p < 0.0001$.

#### Figure S6. Supp. Figures for Main Text Section “Generalized Application #3: Using the EN-TEx Resource to Extend eQTL Annotations to Hard-to-obtain Tissues”

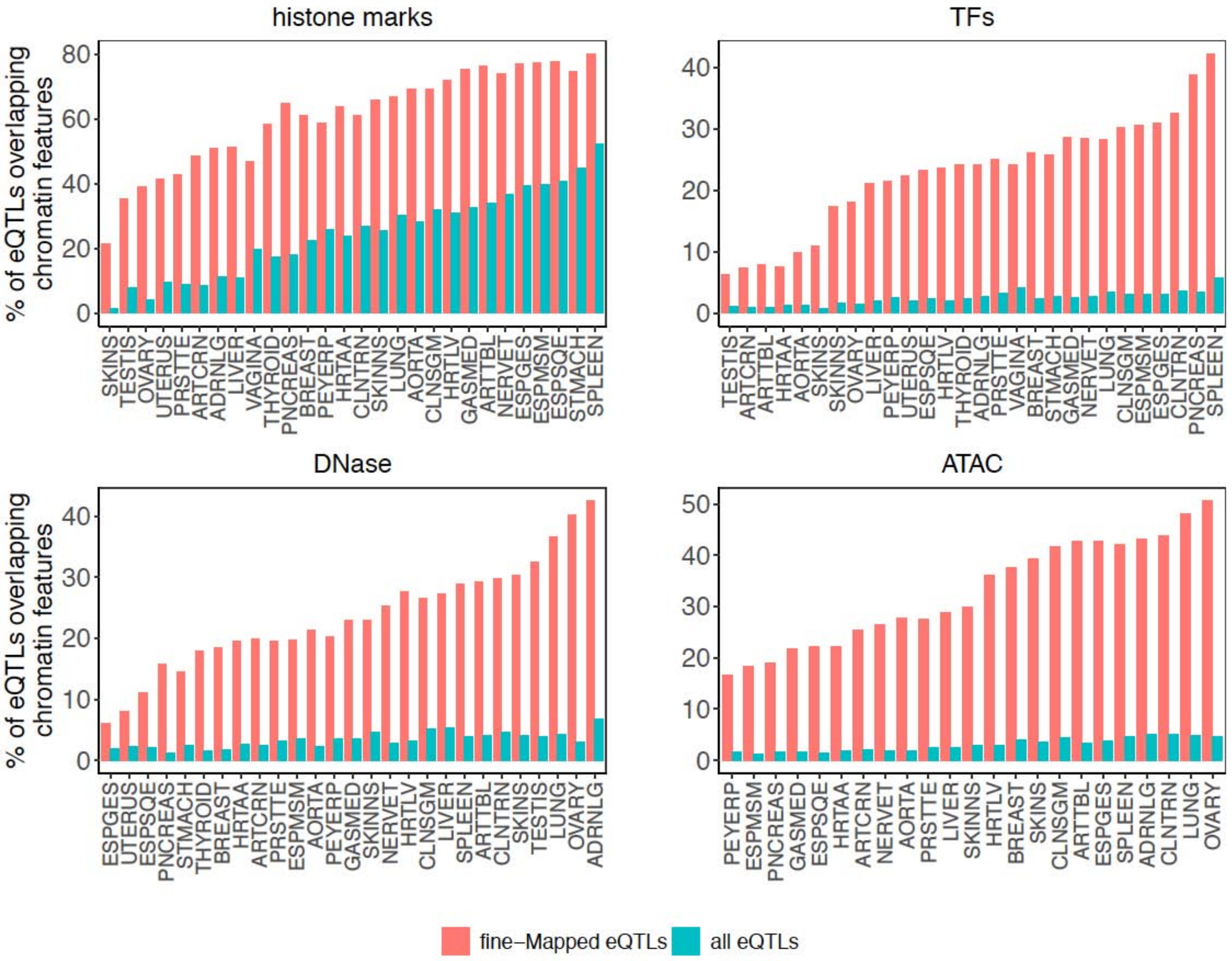

**Figure S6.1a. Chromatin features can help prioritize causal eQTLs**
Barplots showing the percentage of eQTLs overlapping a given feature. The fraction of fine-mapped (causal) eQTLs overlapping chromatin features is higher compared with the total set of GTEx eQTLs reported in a given tissue. In the case of histone marks and TFs, we report the proportion of eQTLs/fine-mapped eQTLs overlapping any of the six histone marks (H3K27ac, H3K4me1, H3K4me3, H3K27me3, H3K9me3, H3K36me3) and any of the four TFs (CTCF, EP300, POLR2A, POLR2APhosphoS5) assayed by the EN-TEx project, respectively.

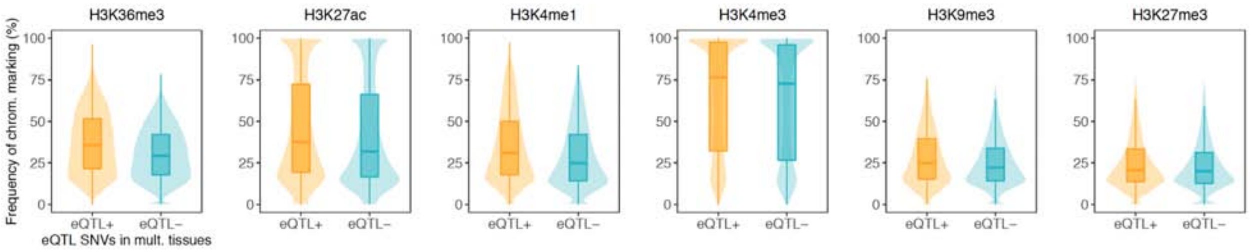

**Figure S6.1b. Chromatin-marked loci associated with eQTL activity**

We have identified 1,353,101 SNVs that show tissue-specific eQTL activity: these SNVs are GTEx eQTLs in ≥ 5 EN-TEx tissues and are not GTEx eQTLs in ≥ 5 other EN-TEx tissues. Thus, for every SNV we define two groups of tissues: i) tissues in which the SNV is an eQTL (eQTL+, orange) and ii) tissues in which the SNV is not an eQTL (eQTL-, cyan). We observed that SNVs are more likely to be marked by a given histone modification in the tissues in which they are eQTLs, compared with the tissues in which they are not eQTLs (p-value < 2.2e-16 for all histone marks, Wilcoxon paired test).

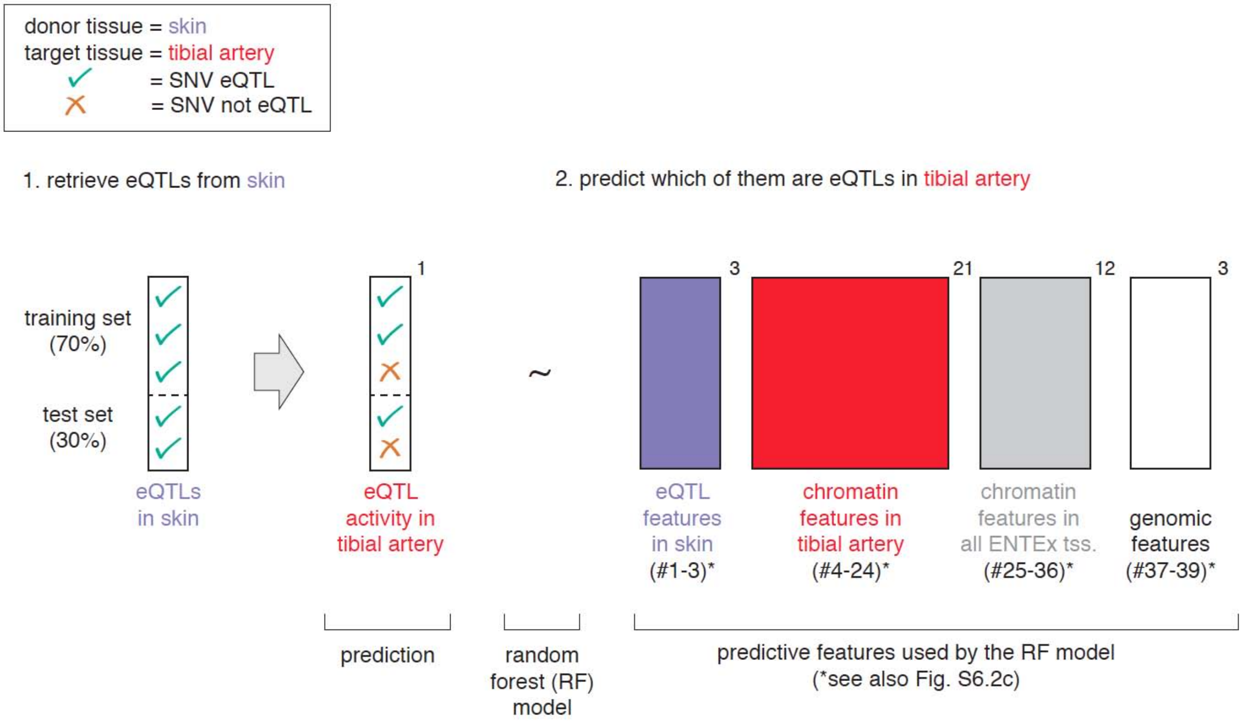

**Figure S6.2a. Schema of the predictive model using skin as the donor tissue and tibial artery as the target tissue**

We first retrieve GTEx eQTLs associated with one single eGene in sun-exposed skin (n = 967,288). This set of eQTLs is split into training and test sets (70% and 30%, respectively). Next, we train a random forest model in order to predict whether these eQTLs are eQTLs also in the tibial artery. The model uses a number of predictive features described in Figure S6.2c: i) eQTL-gene properties in the skin tissue (blue; features #1-3), ii) chromatin profiles at the eQTL loci in the tibial artery tissue (red; features #4-24), iii) chromatin profiles at the eQTL loci across all EN-TEx tissues (gray; features #25-36), iv) genomic features of the eQTL loci (white; features #37-39). The performance of the model is evaluated with a five-fold cross-validation schema.

| | Donor Tissue | Label | N. of samples | N. of eQTLs |
|---|---|---|---|---|
| 1 | Adrenal Gland | ADRNLG | 233 | 422,213 |
| 2 | Artery Aorta | AORTA | 387 | 724,353 |
| 3 | Artery Coronary | ARTCRN | 213 | 336,341 |
| 4 | Artery Tibial | ARTTBL | 584 | 911,849 |
| 5 | Breast - Mammary Tissue | BREAST | 396 | 586,535 |
| 6 | Colon - Sigmoid | CLNSGM | 318 | 573,063 |
| 7 | Colon - Transverse | CLNTRN | 368 | 607,436 |
| 8 | Esophagus - Gastroesophageal Junction | ESPGES | 330 | 579,297 |
| 9 | Esophagus - Mucosa | ESPSQE | 497 | 843,932 |
| 10 | Esophagus - Muscularis | ESPMSM | 465 | 829,886 |
| 11 | Heart - Atrial Appendage | HRTAA | 372 | 641,282 |
| 12 | Heart - Left Ventricle | HRTLV | 386 | 577,859 |
| 13 | Liver | LIVER | 208 | 293,783 |
| 14 | Lung | LUNG | 515 | 776,962 |
| 15 | Muscle - Skeletal | GASMED | 706 | 869,283 |
| 16 | Nerve - Tibial | NERVET | 532 | 1,007,246 |
| 17 | Ovary | OVARY | 167 | 272,146 |
| 18 | Pancreas | PNCREAS | 305 | 552,290 |
| 19 | Prostate | PRSTTE | 221 | 361,651 |
| 20 | Skin - Not Sun Exposed (Suprapubic) | SKINNS | 517 | 861,598 |
| 21 | Skin - Sun Exposed (Lower leg) | SKINS | 605 | 967,288 |
| 22 | Small Intestine - Terminal Ileum | PEYERP | 174 | 290,755 |
| 23 | Spleen | SPLEEN | 227 | 520,408 |
| 24 | Stomach | STMACH | 324 | 483,265 |
| 25 | Testis | TESTIS | 322 | 932,078 |
| 26 | Thyroid | THYROID | 574 | 998,513 |
| 27 | Uterus | UTERUS | 129 | 163,537 |
| 28 | Vagina | VAGINA | 141 | 171,342 |

**Figure S6.2b. Number of eQTLs available from each donor tissue**
For each of the 28 deeply sampled EN-TEx tissues, we list i) the corresponding number of samples with individuals' genotype used by GTEx to perform eQTL analyses (column "N. of samples") and ii) the number of eQTLs used by the model when setting the relevant tissue as "donor tissue" (column "N. of eQTLs"). More specifically, this number corresponds to the set of eQTLs associated with one single eGene in the donor tissue. We also require this single eGene to have non-missing (e.g., "not NA") coefficient of variation for gene expression across EN-TEx individuals and tissues.

| Feature # | Feature | Description | Feature type | Related to | Additional Info |
|---|---|---|---|---|---|
| 1 | tissue_ specificity | coefficient of variation of eGene's expression profile across EN-TEx samples (donors and tissues) | continuous | donor tissue | ratio between mean and standard deviation of gene expression profile |
| 2 | slope | GTEx v8 eQTL-eGene regression slope | continuous | donor tissue | - |
| 3 | tss_distance | GTEx v8 distance from eGene's TSS | continuous | donor tissue | - |
| 4 | ATAC | presence/absence of ATAC peak overlapping the SNV | binary | target tissue | set to 1 if peak present in $\geq$ 1 indiv. |
| 5 | CTCF | presence/absence of CTCF peak overlapping the SNV | binary | target tissue | set to 1 if peak present in $\geq$ 1 indiv. |
| 6 | DNase | presence/absence of DNase peak overlapping the SNV | binary | target tissue | set to 1 if peak present in $\geq$ 1 indiv. |
| 7 | H3K27ac | presence/absence of H3K27ac peak overlapping the SNV | binary | target tissue | set to 1 if peak present in $\geq$ 1 indiv. |
| 8 | H3K27me3 | presence/absence of H3K27me3 peak overlapping the SNV | binary | target tissue | set to 1 if peak present in $\geq$ 1 indiv. |
| 9 | H3K36me3 | presence/absence of H3K36me3 peak overlapping the SNV | binary | target tissue | set to 1 if peak present in $\geq$ 1 indiv. |
| 10 | H3K4me1 | presence/absence of H3K4me1 peak overlapping the SNV | binary | target tissue | set to 1 if peak present in $\geq$ 1 indiv. |
| 11 | H3K4me3 | presence/absence of H3K4me3 peak overlapping the SNV | binary | target tissue | set to 1 if peak present in $\geq$ 1 indiv. |
| 12 | H3K9me3 | presence/absence of H3K9me3 peak overlapping the SNV | binary | target tissue | set to 1 if peak present in $\geq$ 1 indiv. |
| 13 | POLR2A | presence/absence of POLR2A peak overlapping the SNV | binary | target tissue | set to 1 if peak present in $\geq$ 1 indiv. |
| 14 | POLR2A phospho5 | presence/absence of POLR2Aphospho5 peak overlapping the SNV | binary | target tissue | set to 1 if peak present in $\geq$ 1 indiv. |
| 15 | EP300 | presence/absence of EP300 peak overlapping the SNV | binary | target tissue | set to 1 if peak present in $\geq$ 1 indiv. |
| 16 | sum | sum of features 4-15 | discrete | target tissue | - |
| 17 | ATAC_k | fold-change ATAC signal ($\pm$ 5 bp window around the SNV) | continuous | target tissue | mean value across indivs. |
| 18 | CTCF_k | fold-change CTCF signal ($\pm$ 5 bp window around the SNV) | continuous | target tissue | mean value across indivs. |
| 19 | DNase_k | fold-change DNase signal ($\pm$ 5 bp window around the SNV) | continuous | target tissue | mean value across indivs. |
| 20 | H3K27ac_k | fold-change H3K27ac signal ($\pm$ 5 bp window around the SNV) | continuous | target tissue | mean value across indivs. |
| 21 | H3K27me3_k | fold-change H3K27me3 signal ($\pm$ 5 bp window around the SNV) | continuous | target tissue | mean value across indivs. |
| 22 | H3K4me1_k | fold-change H3K4me1 signal ($\pm$ 5 bp window around the SNV) | continuous | target tissue | mean value across indivs. |

| Feature # | Feature | Description | Feature type | Related to | Additional Info |
|---|---|---|---|---|---|
| 23 | H3K9me3_k | fold-change H3K9me3 signal ($\pm$ 5 bp window around the SNV) | continuous | target tissue | mean value across indivs. |
| 24 | POLR2A_k | fold-change POLR2A signal ($\pm$ 5 bp window around the SNV) | continuous | target tissue | mean value across indivs. |
| 25 | ATAC_p | fraction of tissues with ATAC peak over SNV | continuous | 28 EN-TEx tissues | - |
| 26 | CTCF_p | fraction of tissues with CTCF peak over SNV | continuous | 28 EN-TEx tissues | - |
| 27 | DNase_p | fraction of tissues with DNase peak over SNV | continuous | 28 EN-TEx tissues | - |
| 28 | H3K27ac_p | fraction of tissues with H3K27ac peak over SNV | continuous | 28 EN-TEx tissues | - |
| 29 | H3K27me3_p | fraction of tissues with H3K27me3 peak over SNV | continuous | 28 EN-TEx tissues | - |
| 30 | H3K36me3_p | fraction of tissues with H3K36me3 peak over SNV | continuous | 28 EN-TEx tissues | - |
| 31 | H3K4me1_p | fraction of tissues with H3K4me1 peak over SNV | continuous | 28 EN-TEx tissues | - |
| 32 | H3K4me3_p | fraction of tissues with H3K4me3 peak over SNV | continuous | 28 EN-TEx tissues | - |
| 33 | H3K9me3_p | fraction of tissues with H3K9me3 peak over SNV | continuous | 28 EN-TEx tissues | - |
| 34 | POLR2A_p | fraction of tissues with POLR2A peak over SNV | continuous | 28 EN-TEx tissues | - |
| 35 | POLR2A phosphoS5_p | fraction of tissues with POLR2AphosphoS5 peak over SNV | continuous | 28 EN-TEx tissues | - |
| 36 | EP300_p | fraction of tissues with EP300 peak over SNV | continuous | 28 EN-TEx tissues | - |
| 37 | is_proximal | whether the SNV overlaps any annotated TSS ($\pm$ 2 kb) | binary | - | set to 1 if SNV overlaps TSS wrt gencode v 24 |
| 38 | is_cCRE | whether the SNV overlaps any cCRE | binary | - | set to 1 if SNV overlaps cCRE wrt ENCODE3 |
| 39 | is_out_repeat | whether the SNV overlaps any repeated region | binary | - | set to 1 if SNV does not overlap repeated region |

**Figure S6.2c. List of predictive features employed by the random forest model to predict eQTL activity in a target tissue**

Features employed to predict which donor-tissue eQTLs can be transferred to a target tissue. For features 25-36, the fraction is computed over those of the 28 tissues with available data for the relevant experiment (e.g., if no ATAC-seq experiments were performed for lung tissue, then lung is not included in the calculation of ATAC_p).

| Metric | Description |
| --- | --- |
| Sensitivity | $\frac{TP}{TP + FN}$ |
| Specificity | $\frac{TN}{FP + TN}$ |
| Precision | $\frac{TP}{TP + FP}$ |
| Balanced Accuracy | $\frac{\text{sensitivity} + \text{specificity}}{2}$ |
| MCC | $\frac{TP \times TN - FP \times FN}{\sqrt{(TP + FP)(TP + FN)(TN + FP)(TN + FN)}}$ |
| Accuracy | $\frac{TP + TN}{TP + TN + FP + FN}$ |
| No-information Rate | the largest proportion of the observed classes |
| Accuracy Ratio | $\frac{\text{Accuracy}}{\text{No-information Rate}}$ |
| Prevalence | $\frac{TP + FN}{TP + TN + FP + FN}$ |

**Figure S6.3a. Description of the metrics used to evaluate the random forest model**
These metrics have been used in the submodels' evaluations shown in Figure 8B and Figure S6.3c.

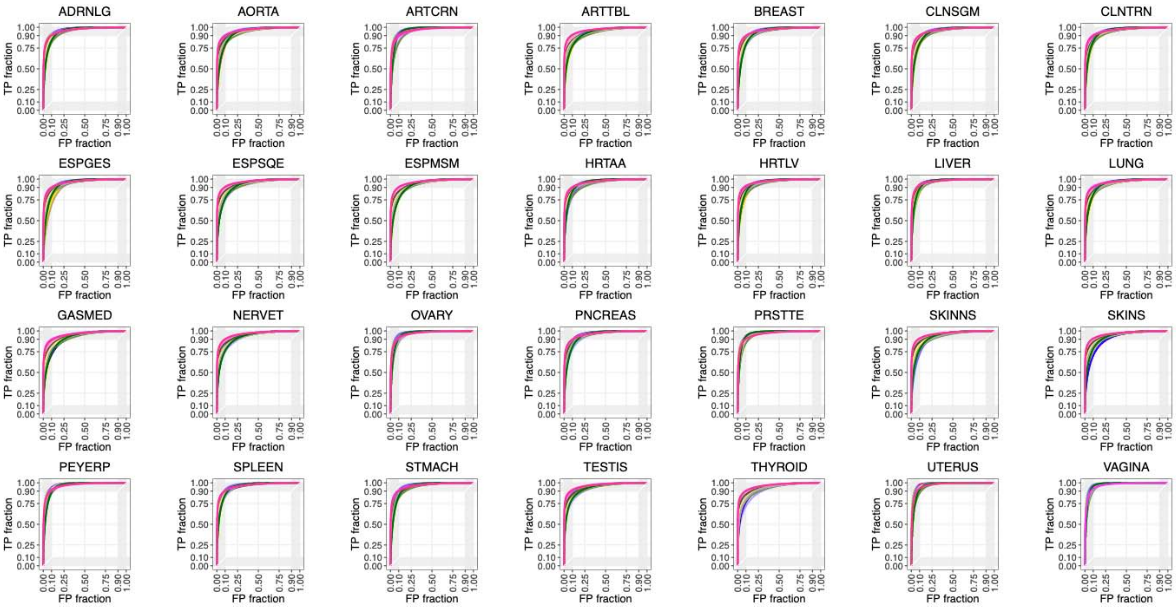

**Figure S6.3b. Performance of the random forest submodels by donor tissue**

Each plot shows receiver operating characteristic (ROC) curves from multiple target-tissue submodels obtained using the same donor tissue. For instance, the first plot shows ROC curves obtained from all submodels using adrenal gland (ADRNLG) as the donor tissue. ROC curves for each target tissue were computed on a five-fold cross-validation schema, and are color-coded in the figure (see Figure S6.2b for a correspondence between tissues and colors).

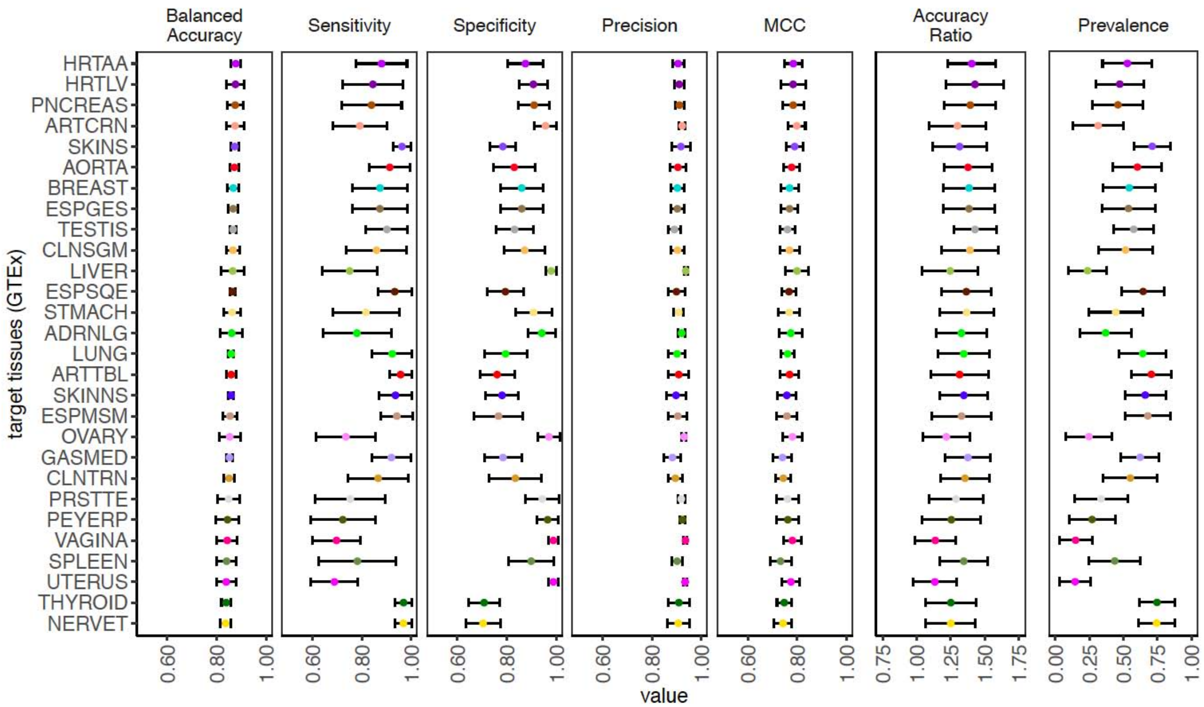

**Figure S6.3c. Performance of the random forest submodels by target tissue**
Dotplot reporting, for each target tissue (y axis), performance metrics (x axis) of the models obtained by using different donor tissues. For a particular target tissue, we report mean and standard deviation of the metric computed using different donor tissues. See Figure S6.3a for a detailed description of each performance metric.

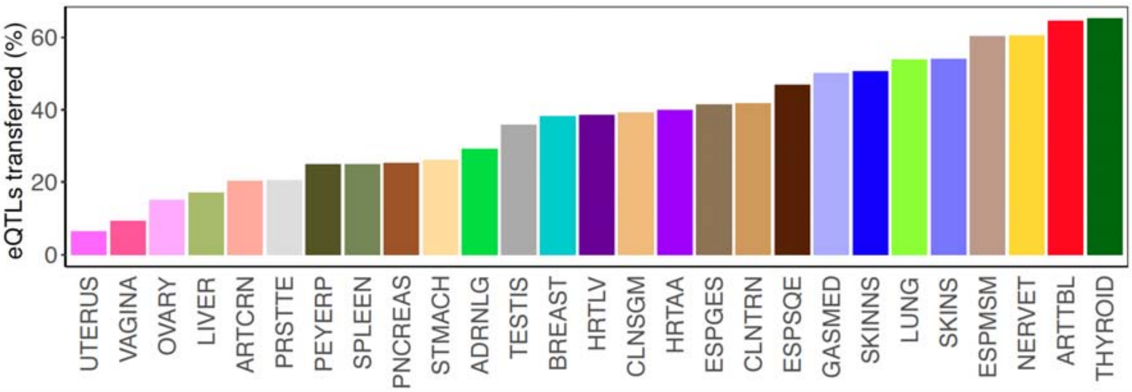

**Figure S6.3d. Proportion (%) of blood eQTLs from Vosa et al. 2021 that can be transferred to each EN-TEx tissue**

In this analysis, we aimed to predict the activity of 1,547,430 blood eQTLs from (Vosa *et al.*, 2021) in every EN-TEx tissue. To do so, we applied, for every target tissue, the submodel previously trained on GTEx data that uses artery aorta as the donor tissue (since blood is not among the EN-TEx tissues, we do not currently have a model using blood as the donor tissue). For each EN-TEx tissue (x axis) other than artery aorta, we report the proportion of blood eQTLs (y axis) predicted to be active by the model specific to each target tissue. Because of some overlap between this catalog of blood eQTLs and the GTEx catalogs, we computed these results after excluding the blood eQTLs that were also contained in the original training set used for every target tissue (see also Figure S6.3e).

| Target Tissue | Total | Predicted | Novel |
|---|---|---|---|
| THYROID | 1,467,572 (1,290,756) | 958,740 (841,058) | 816,420 (698,738) |
| ARTTBL | 1,467,916 (1,286,326) | 947,245 (828,900) | 837,659 (719,314) |
| NERVET | 1,467,940 (1,289,188) | 887,180 (777,348) | 758,986 (649,154) |
| ESPMSM | 1,468,001 (1,289,266) | 886,159 (776,500) | 799,913 (690,254) |
| SKINS | 1,468,004 (1,288,495) | 793,089 (695,178) | 683,870 (585,959) |
| LUNG | 1,467,841 (1,289,600) | 789,415 (692,658) | 720,074 (623,317) |
| SKINNS | 1,467,830 (1,288,516) | 744,235 (652,032) | 656,856 (564,653) |
| GASMED | 1,467,849 (1,284,844) | 734,548 (641,891) | 646,988 (554,331) |
| ESPSQE | 1,467,929 (1,285,876) | 688,868 (603,916) | 615,039 (530,087) |
| CLNTRN | 1,467,703 (1,288,973) | 612,148 (536,318) | 575,364 (499,534) |
| ESPGES | 1,467,849 (1,289,115) | 609,024 (534,229) | 575,021 (500,226) |
| HRTAA | 1,467,844 (1,287,852) | 586,424 (515,551) | 549,573 (478,700) |
| CLNSGM | 1,467,970 (1,289,237) | 574,549 (504,154) | 544,411 (474,016) |
| HRTLV | 1,467,925 (1,286,240) | 565,778 (496,803) | 533,022 (464,047) |
| BREAST | 1,467,981 (1,288,553) | 560,010 (490,672) | 526,559 (457,221) |
| TESTIS | 1,467,715 (1,293,072) | 526,338 (461,012) | 460,638 (395,312) |
| ADRNLG | 1,467,819 (1,289,468) | 427,103 (377,276) | 413,087 (363,260) |
| STMACH | 1,467,763 (1,289,089) | 382,833 (335,265) | 366,193 (318,625) |
| PNCREAS | 1,467,795 (1,287,656) | 369,129 (323,967) | 349,283 (304,121) |
| PEYERP | 1,467,623 (1,290,110) | 364,123 (322,879) | 357,375 (316,131) |
| SPLEEN | 1,467,822 (1,290,788) | 364,926 (319,817) | 347,251 (302,142) |
| PRSTTE | 1,467,921 (1,289,186) | 299,553 (263,773) | 290,817 (255,037) |
| ARTCRN | 1,467,752 (1,287,251) | 297,262 (260,061) | 291,965 (254,764) |
| LIVER | 1,467,913 (1,285,031) | 251,342 (220,727) | 245,376 (214,761) |
| OVARY | 1,467,879 (1,289,709) | 219,686 (193,123) | 215,232 (188,669) |
| VAGINA | 1,467,893 (1,290,434) | 136,579 (117,983) | 134,710 (116,114) |
| UTERUS | 1,467,947 (1,288,382) | 94,299 (83,539) | 93,199 (82,439) |

**Figure S6.3e. Number of potentially novel eQTLs predicted for each EN-TEx tissue that are not present in the GTEx catalog**

This table reports, for every target tissue, i) the total number of blood eQTLs (from (Vosa *et al.*, 2021)) analyzed after removing those contained in the training set (column “total”), ii) the number of blood eQTLs transferred, i.e., predicted to be active (column “predicted”), and iii) within the predicted eQTLs, the number of novel eQTLs that are not contained in the GTEx eQTL catalog for the relevant tissue. We identified 496,477 novel eQTLs on average across tissues. Because not all blood eQTLs from (Vosa *et al.*, 2021) might also have been tested by GTEx for gene associations in the relevant tissue, we also report (in parentheses) the numbers of total, predicted, and novel eQTLs out of those SNVs tested also by GTEx in a particular tissue.

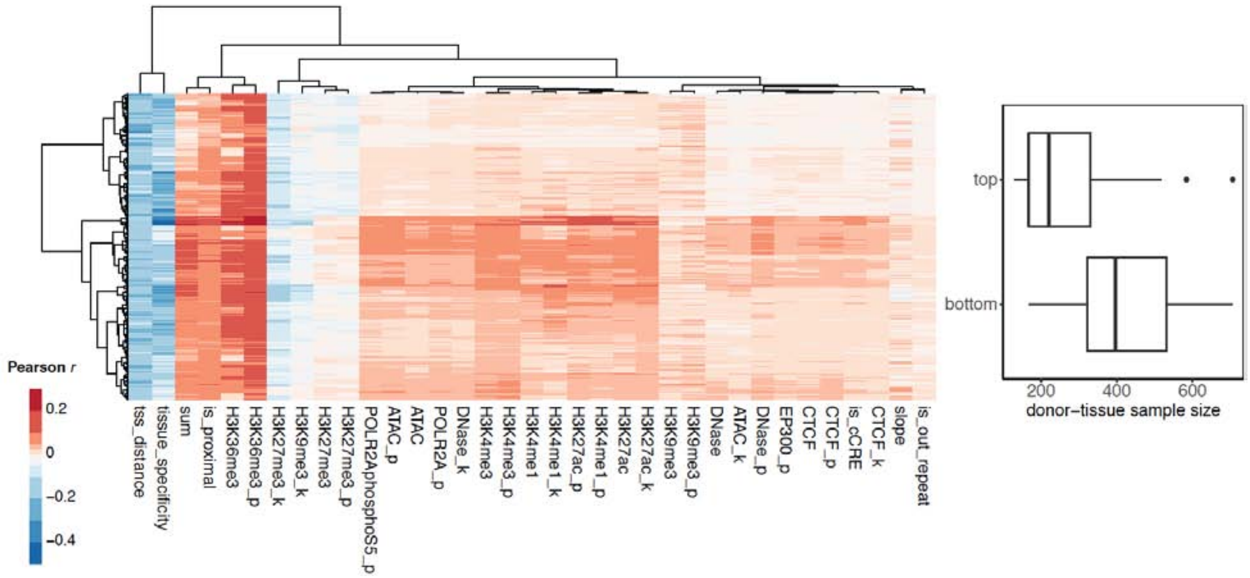

**Figure S6.4a. Dissecting the contribution of features to predicting tissue-specific eQTL activity**

[left] Heatmap showing, for each submodel (rows), Pearson's correlation coefficients between the level of predictive features (columns) at donor-tissue eQTLs and the probability of donor-tissue eQTLs of being classified as eQTLs in the target tissue (clustering method: "Ward.D2", clustering distance: "manhattan"). [right] Boxplot showing the sample size of donor tissues used for the submodels in the top and bottom (row) clusters of the heatmap. The sample size of each tissue is reported in Figure S6.2b (column "N. of samples").

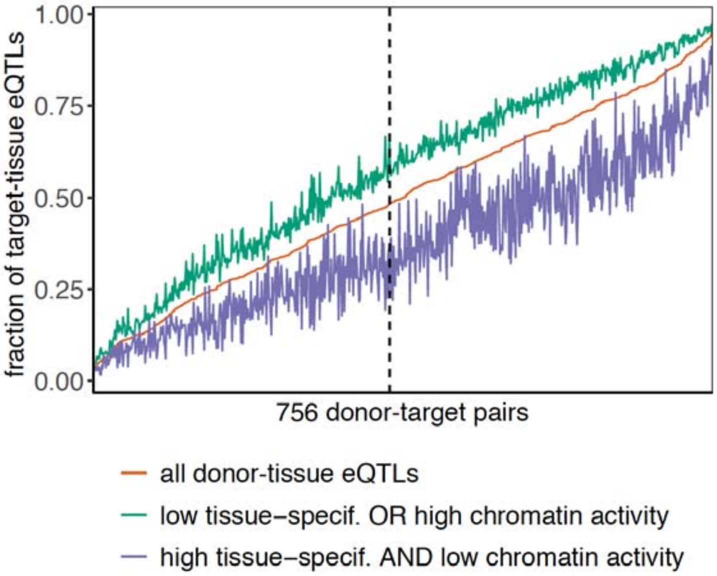

**Figure S6.4b. By analyzing the chromatin activity of donor-tissue eQTLs in the target tissue and the tissue specificity of their eGenes, we can identify eQTLs active in the target tissue**

Donor-tissue eQTLs either associated with housekeeping eGenes, or those that have high chromatin activity in the target tissue (green line), are more frequently also eQTLs in the target tissue, compared with donor-tissue eQTLs associated with tissue-specific eGenes and that have low chromatin activity in the target tissue (purple line). These enrichments are compared with the proportion of target-tissue eQTLs out of all donor-tissue eQTLs used across the 756 donor-target tissue pairs (orange line). Green line: donor-tissue eQTLs with “tissue specificity” < 0.8 or “sum” (chromatin marking) ≥ 3. Purple line: “tissue specificity” > 5 and “sum” = 0. See also Figure S6.2c for the definition of “sum” and “tissue specificity” features. The dashed vertical line corresponds to the results shown in Figure 8G, i.e., the case using testis as the donor tissue, and thyroid as the target tissue.

#### Figure S7. Supp. Figures for Main Text Section “Generalized Application #4: Decorating ENCODE Regulatory Elements with EN-TEx Tissue & AS Information”

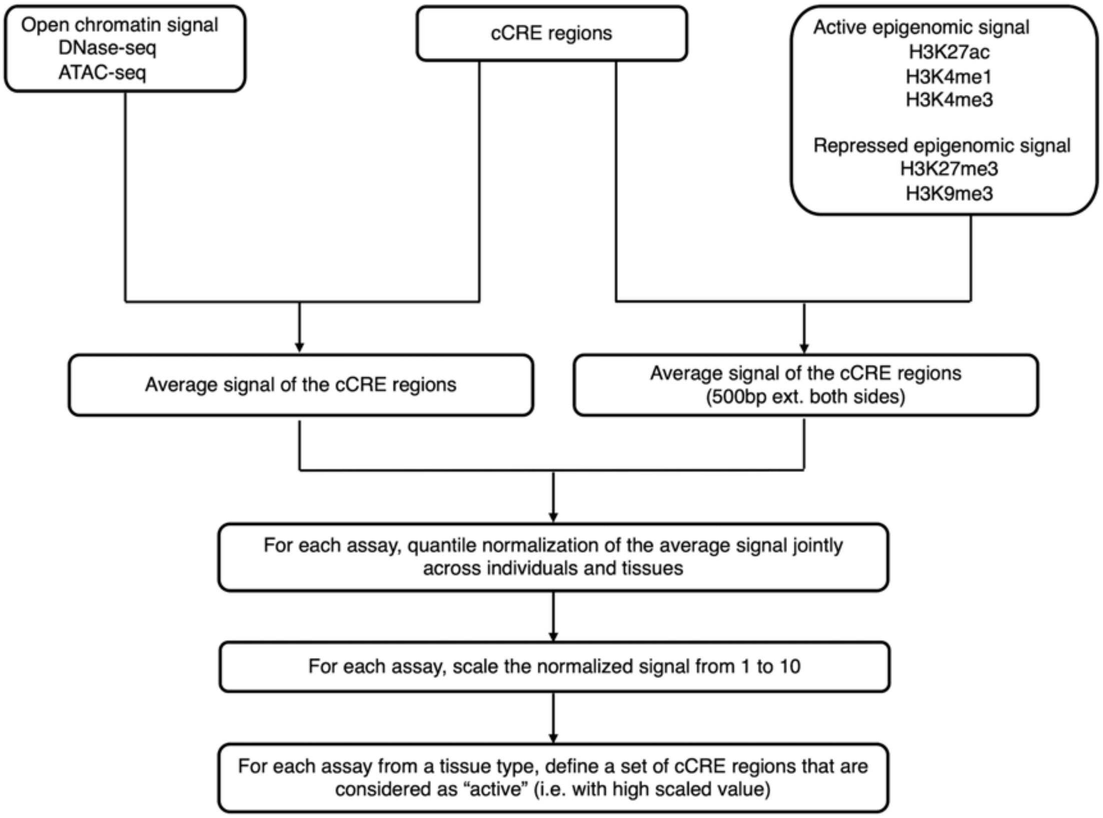

**Figure S7.1a. Data preprocessing**
We computed the average signal for each cCRE region using the datasets from DNase-seq, ATAC-seq, and five histone modifications (H3K27ac, H3K4me1, H3K4me3, H3K27me3, and H3K9me3). For DNase-seq and ATAC-seq, the signals were averaged across the genomic positions of the cCRE regions. The signals of histone modifications were averaged across the genomic positions of the cCRE regions with a 500 bp extended region on each side. For each assay, we performed quantile normalization on the average signal from the cCRE regions jointly across all of the biosamples. Then, we scaled the normalized signal from 1 to 10, and defined a set of "active" cCREs for each assay from each tissue type.

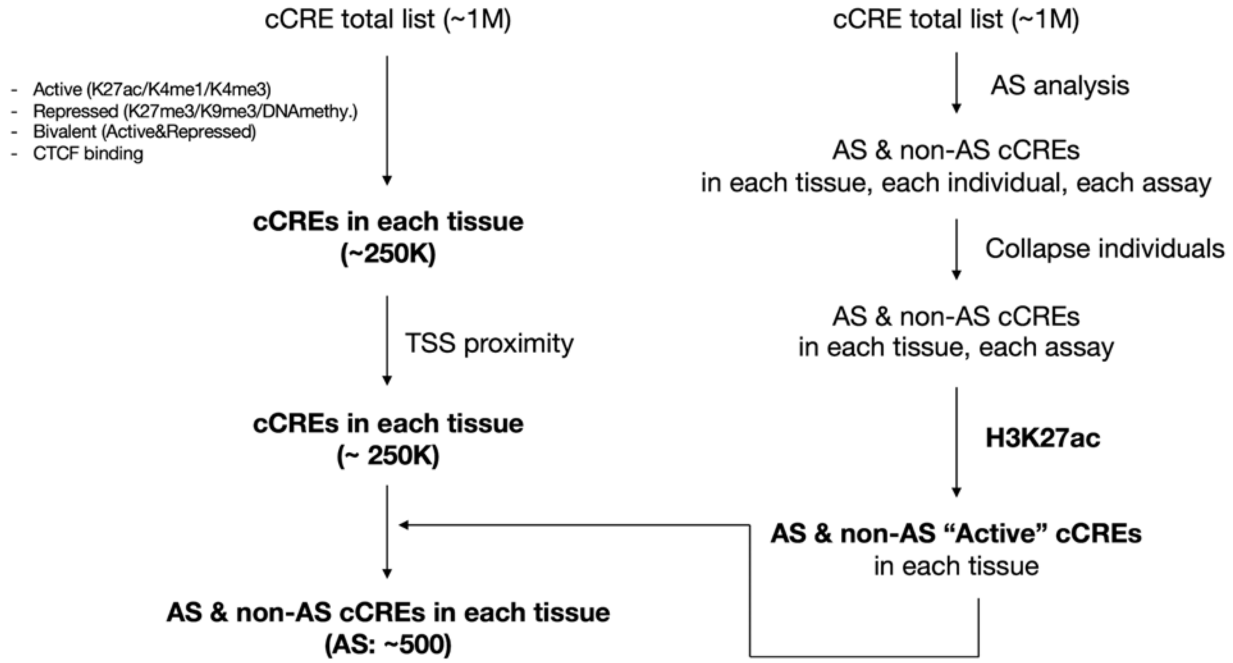

**Figure S7.1b. Framework of cCRE decoration**

We decorated the cCREs from the encyclopedia using the active and repressed histone modification signals and CTCF binding sites from tissues. The decorated cCREs were then separated into proximal and distal groups based on their proximity to the annotated TSSs. At another layer, these cCRE subgroups were further annotated as AS and non-AS based on their allelic signature.

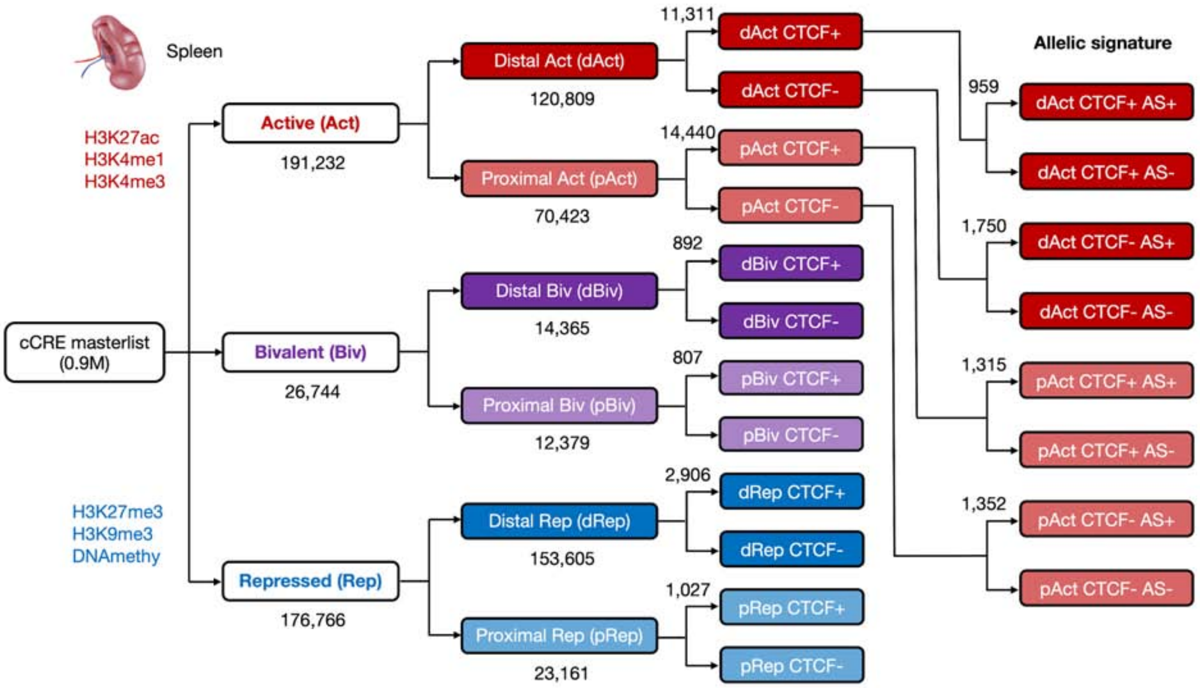

**Figure S7.1c. Framework of cCRE decoration in the spleen**

This figure shows the workflow of cCRE decoration and the numbers of different subgroups of cCREs in the spleen. Note that we define a number of abbreviations for the various decorations. dACT: distal active; pACT: proximal active; dBiv: distal bivalent; pBiv: proximal bivalent; dRep: distal repressed; pRep: proximal repressed; CTCT+ and CTCF- indicates with and without CTCF binding, respectively; AS+ and AS- indicates with and without allelic signature, respectively.

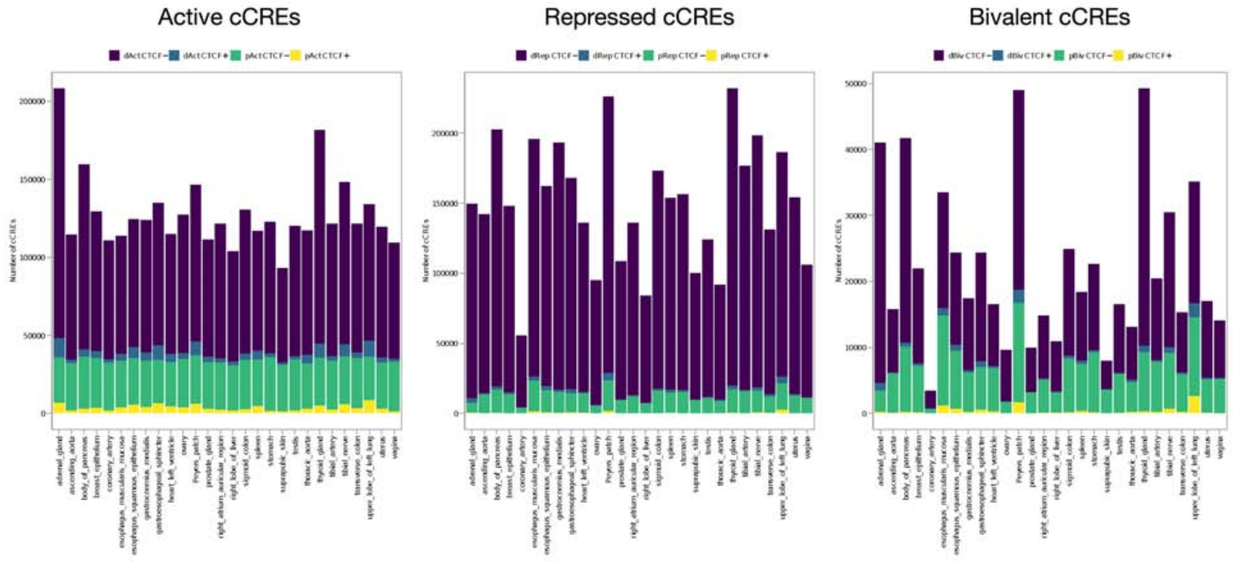

**Figure S7.1d. Number of cCREs in various tissues**
This figure shows the number of different subgroups of decorated cCREs in each tissue type. In each panel, the colors indicate the TSS proximity (proximal vs. distal) and CTCF binding state (CTCF+ vs. CTCF-). Note that the different decoration terms are defined in Figure S7.1c.

**Decoration of total cCREs**

[active/bivalent/repressed].[distal/proximal].[CTCF/nonCTCF]-[TissueType]

**Decoration of cCREs with allelic signature**

[active/repressed].[distal/proximal].[CTCF/nonCTCF].[AS/nonAS]-[TissueType]

890,906 cCREs

| cCRE_id | active.distal.CTCF-adrenal_gland | ... | ... | repressed.proximal.nonCTCF-vagina | active.distal.CTCF.AS-adrenal_gland | ... | ... | repressed.proximal.nonCTCF.nonAS-vagina |
|---|---|---|---|---|---|---|---|---|
| EH38E0001876 | 0 | ... | ... | 0 | 0 | ... | ... | 0 |
| EH38E0004911 | 0 | ... | ... | 0 | 0 | ... | ... | 0 |
| EH38E0005334 | 1 | ... | ... | 0 | 0 | ... | ... | 0 |
| EH38E0006178 | 0 | ... | ... | 1 | 0 | ... | ... | 1 |
| EH38E0006270 | 0 | ... | ... | 1 | 0 | ... | ... | 1 |
| ... | ... | ... | ... | ... | ... | ... | ... | ... |
| ... | ... | ... | ... | ... | ... | ... | ... | ... |
| ... | ... | ... | ... | ... | ... | ... | ... | ... |
| ... | ... | ... | ... | ... | ... | ... | ... | ... |
| EH38E2776491 | 0 | ... | ... | 0 | 0 | ... | ... | 0 |
| EH38E2776496 | 0 | ... | ... | 0 | 0 | ... | ... | 0 |
| EH38E2776512 | 1 | ... | ... | 0 | 0 | ... | ... | 0 |
| EH38E2776513 | 0 | ... | ... | 0 | 0 | ... | ... | 0 |
| EH38E2776514 | 0 | ... | ... | 0 | 0 | ... | ... | 0 |

**Figure S7.1e. cCRE decoration results matrix**

We generated an annotation matrix for all the decorated cCREs from tissue types. This corresponds to File: cCRE_decoration.matrix.

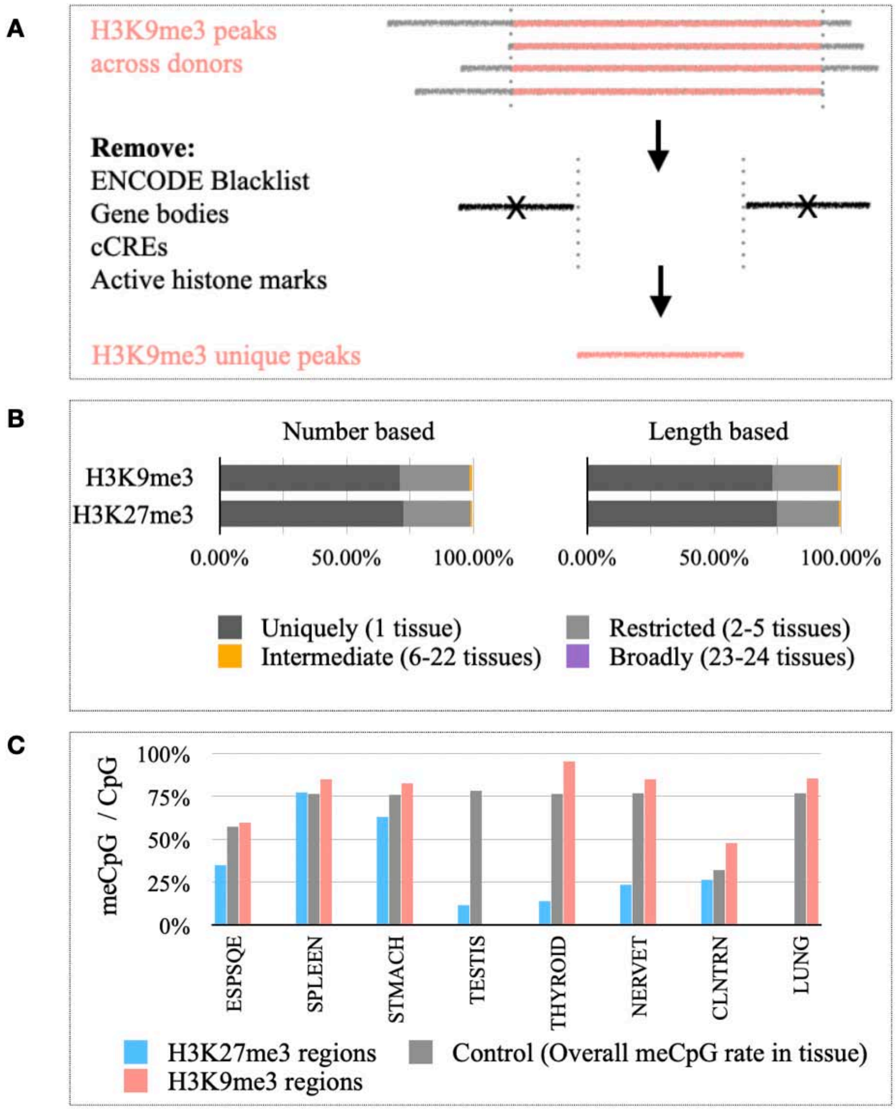

**Figure S7.1f. Identifying fully repressed elements independent of cCREs**
**(A)** For genomic regions outside of cCREs and annotated genes, elements longer than 200 bp that are uniquely marked by either H3K9me3 or H3K27me3 were defined as fully repressed. A total of 45,207 (covering 12,655,795 bp) and 24,006 (covering 7,474,178 bp) non-overlapping elements were identified based on H3K9me3 and H3K27me3, respectively. Identified elements can be found in File: ENTEx_fully_repressed_regions_independent_of_cCREs.bed. **(B)** The majority of these elements were repressed in a tissue-specific manner. **(C)** For tissues with available datasets, DNA methylation within these elements was evaluated, and H3K9me3-marked elements showed a significantly (t-test, p-value < 0.05) higher CpG methylation (meCpG) rate than elements marked uniquely by H3K27me3.

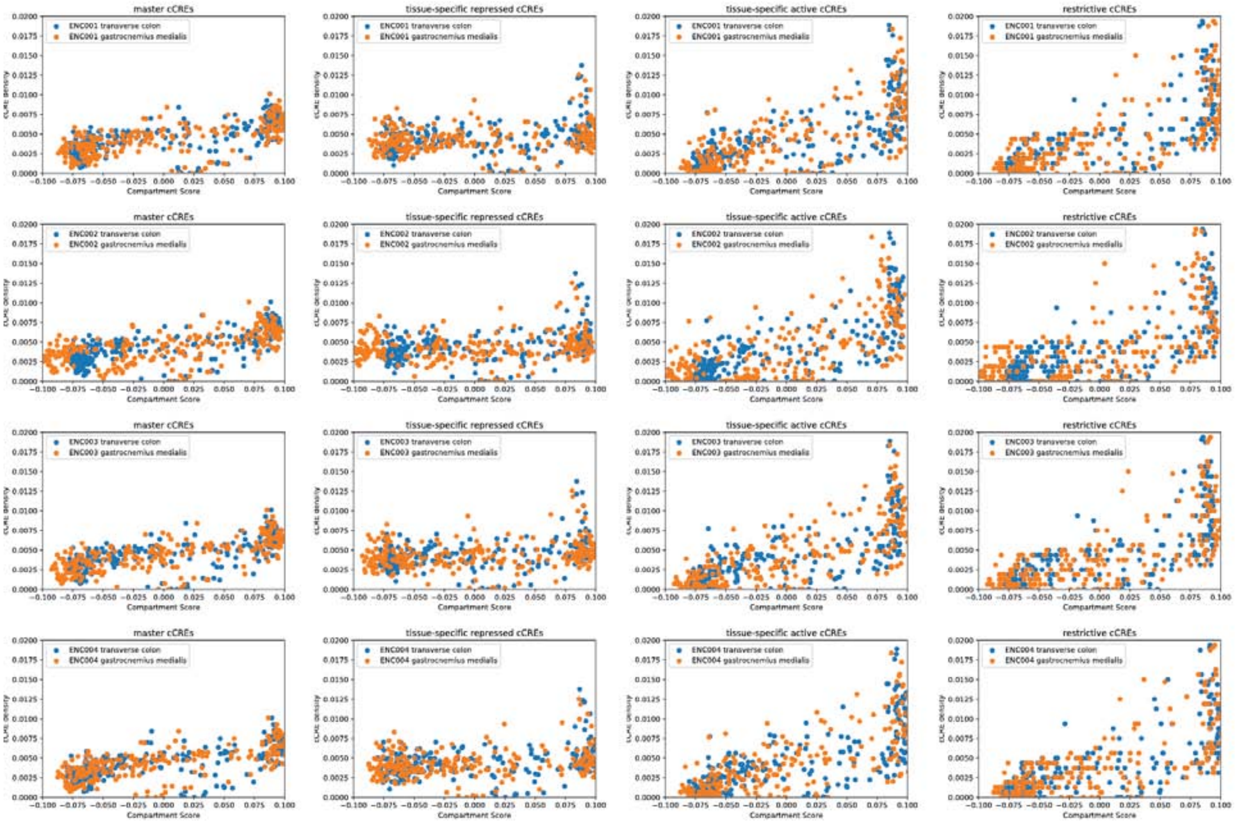

**Figure S7.1g. cCRE enrichment with respect to A/B compartments**

These plots show the cCRE enrichment in the A vs. B compartment of two different tissues. We show this for the master cCRE list from ENCODE, including both tissue-specific active and repressed cCREs. As the tissue specificity increases, the cCRE enrichment in the active A compartment increases compared with the inactive B compartment.

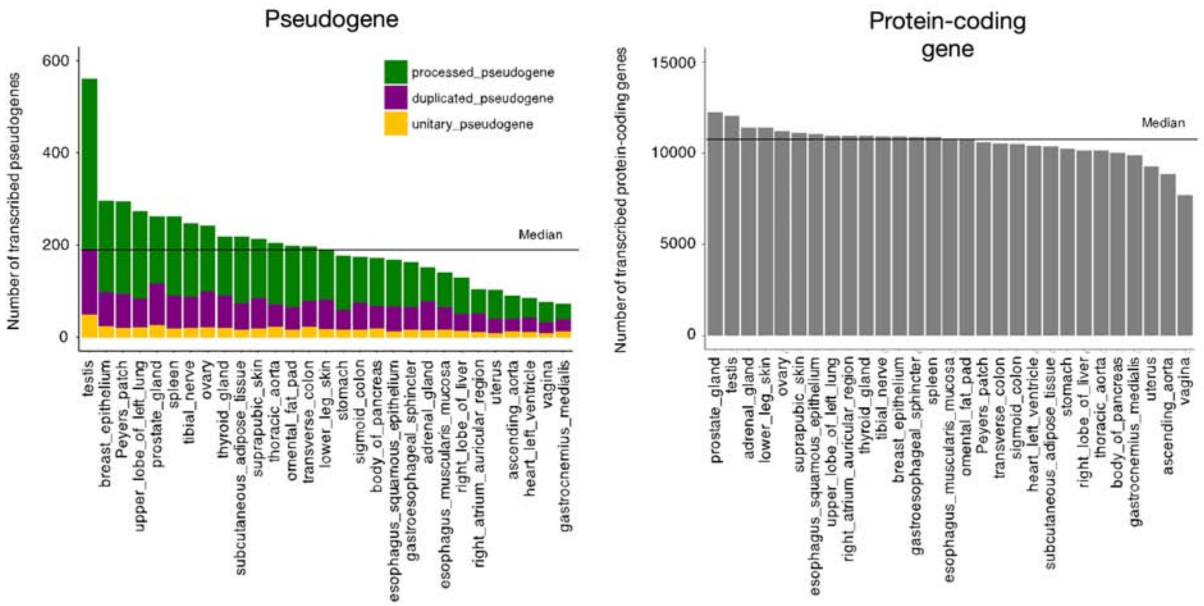

**Figure S7.2a. The number of transcribed genes in tissues**

This figure shows the number of transcribed pseudogenes (left) and protein-coding genes (right) across all tissue types. The median of transcribed pseudogenes and protein-coding genes across the tissues is 200 and ~11K, respectively.

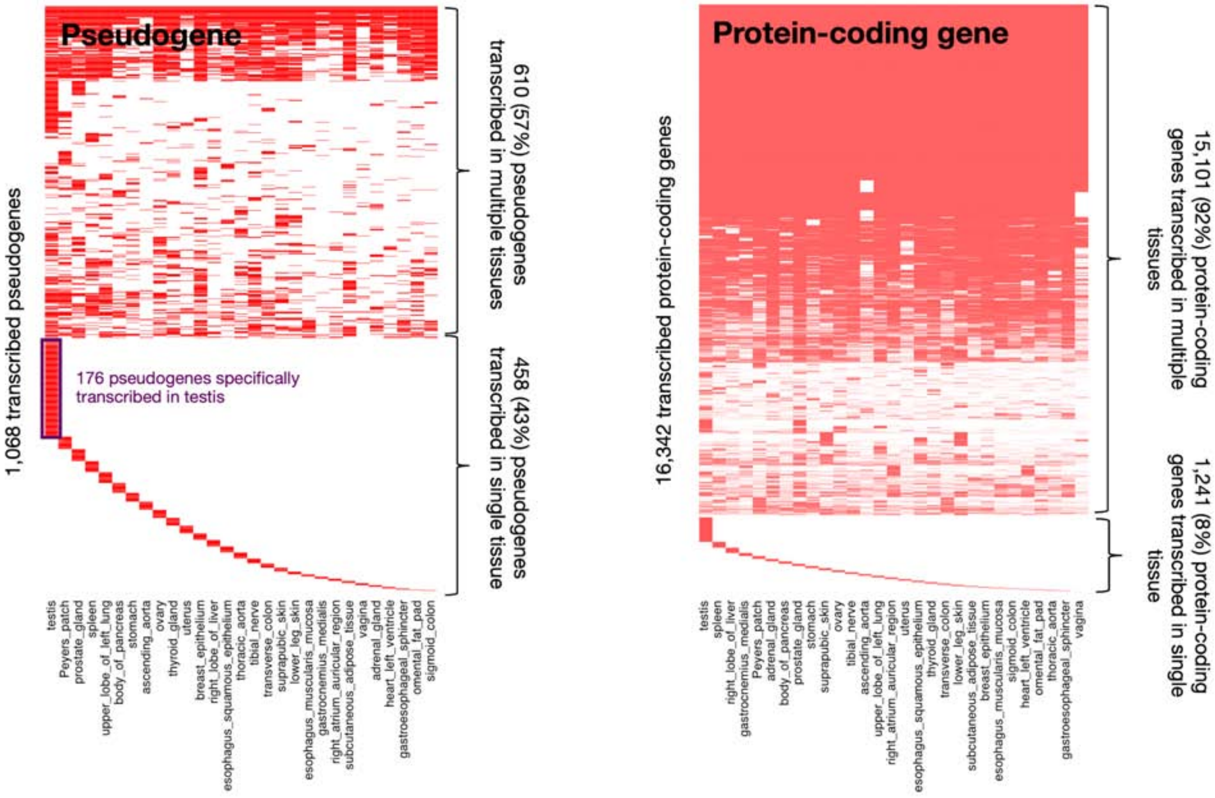

**Figure S7.2b. Tissue specificity of transcribed genes**

The heatmaps show the activity of pseudogenes (left) and protein-coding genes (right) across tissue types. In each tissue, the pseudogenes/protein-coding genes are classified as actively transcribed (shown in red) or not based on their expression level.

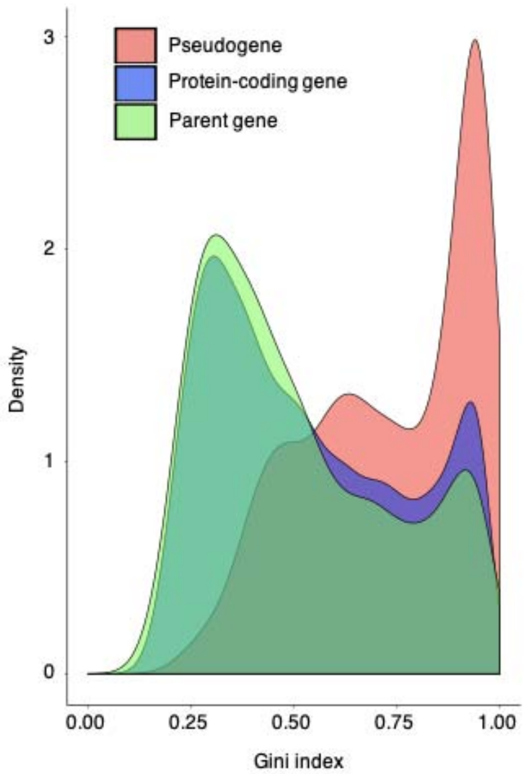

**Figure S7.2c. Gini index of gene expression level across tissues**
We applied the Gini index to quantify the tissue specificity of protein-coding genes, pseudogenes, and parent genes based on their expression level. The pseudogenes show higher Gini indexes than protein-coding genes, suggesting stronger tissue specificity of pseudogenes. The Gini index distribution of the pseudogenes is quite different from that of the parent genes, confirming that the multi-mapping bias from quantification of the pseudogene expression level has been minimized.

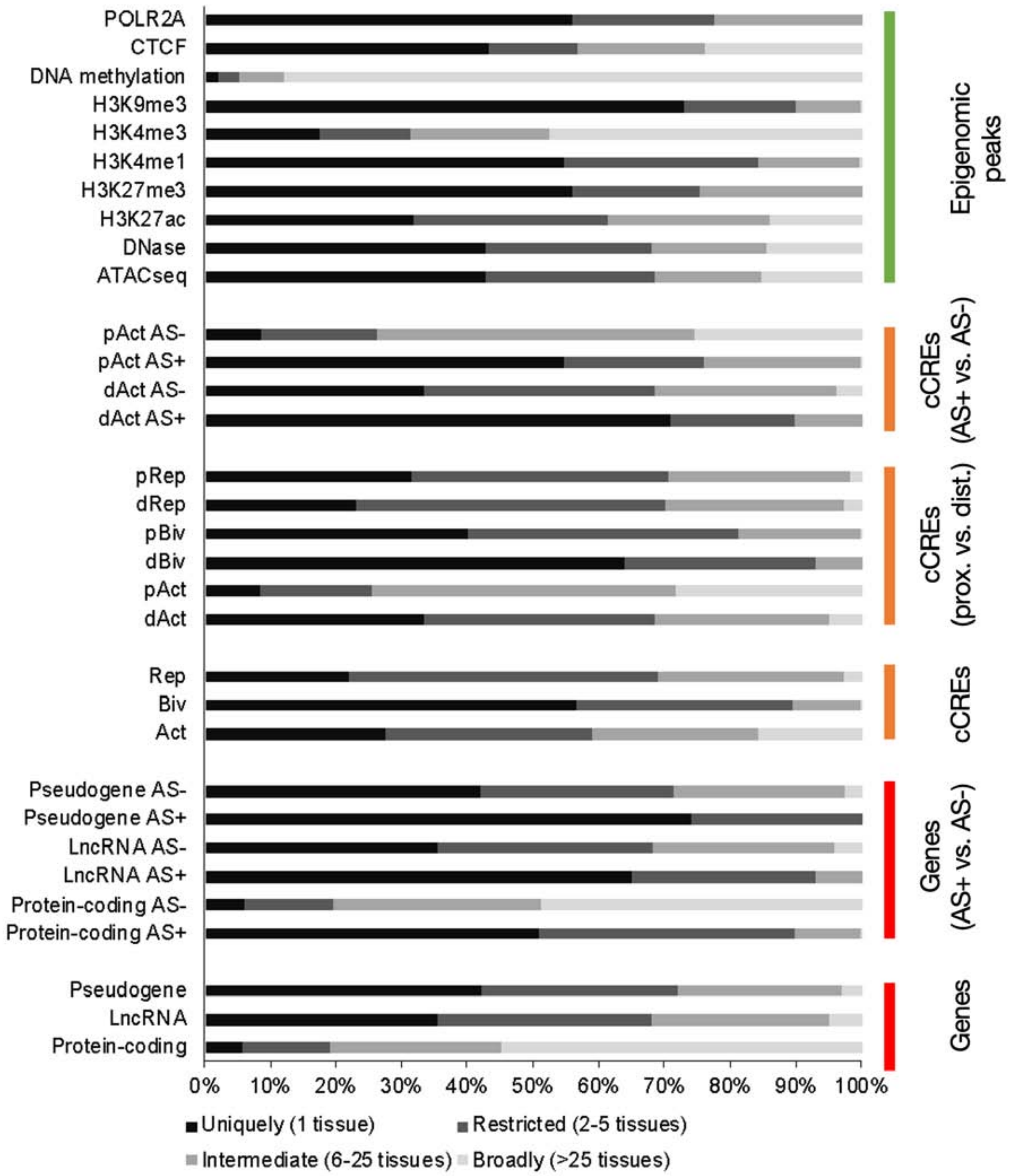

**Figure S7.2d. Tissue specificity of different subgroups of cCREs vs. genes and epigenomic peaks**

We compared the tissue specificity of protein-coding genes, non-coding genes, different subgroups of decorated cCREs, and various epigenomic peaks. The uniqueness of the activity across tissue types is shown in different colors. Note that the different decoration terms are defined in Figure S7.1c.

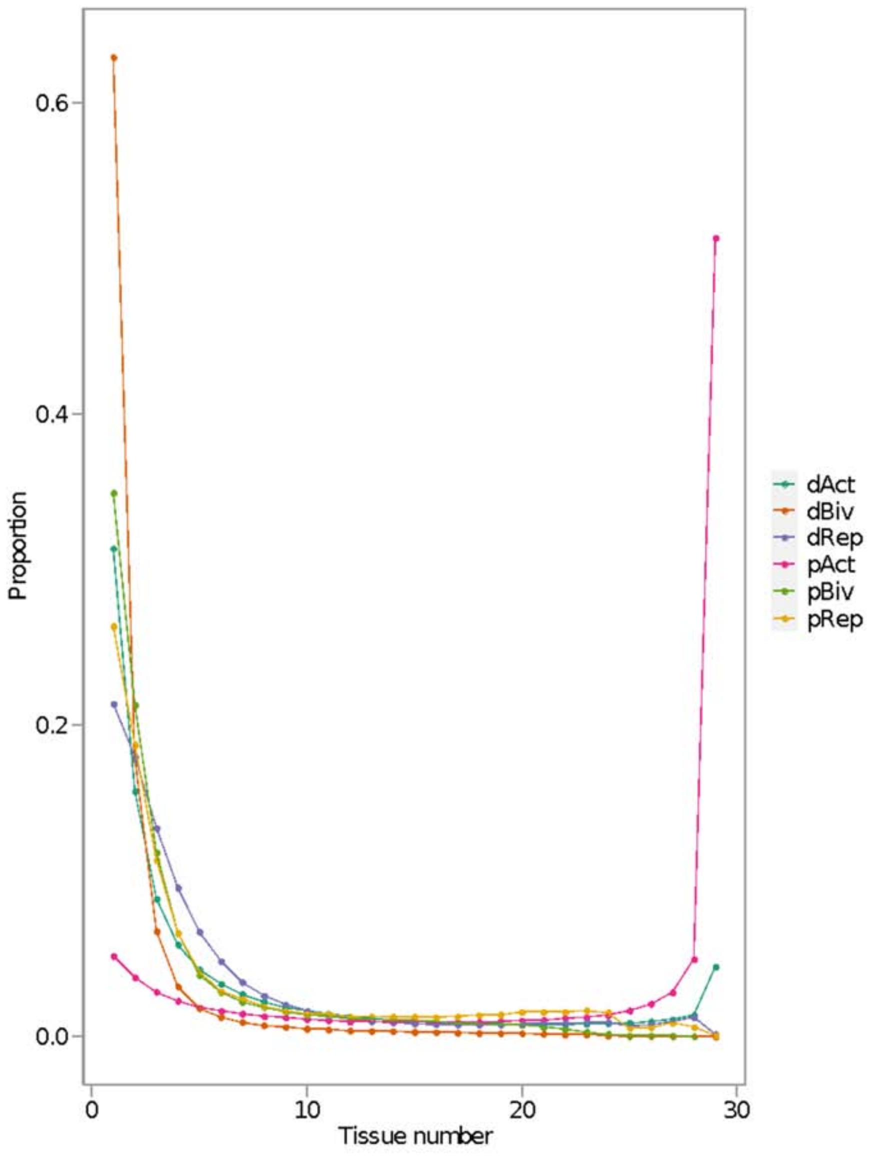

**Figure S7.2e. Tissue specificity of different subgroups of cCREs**
For each cCRE subgroup, we show the proportion of the cCREs that are defined as "active" across the different numbers of tissue types ranging from one (i.e., high tissue specificity) to all tissue types (i.e., low tissue specificity). Note that the different decoration terms are defined in Figure S7.1c.

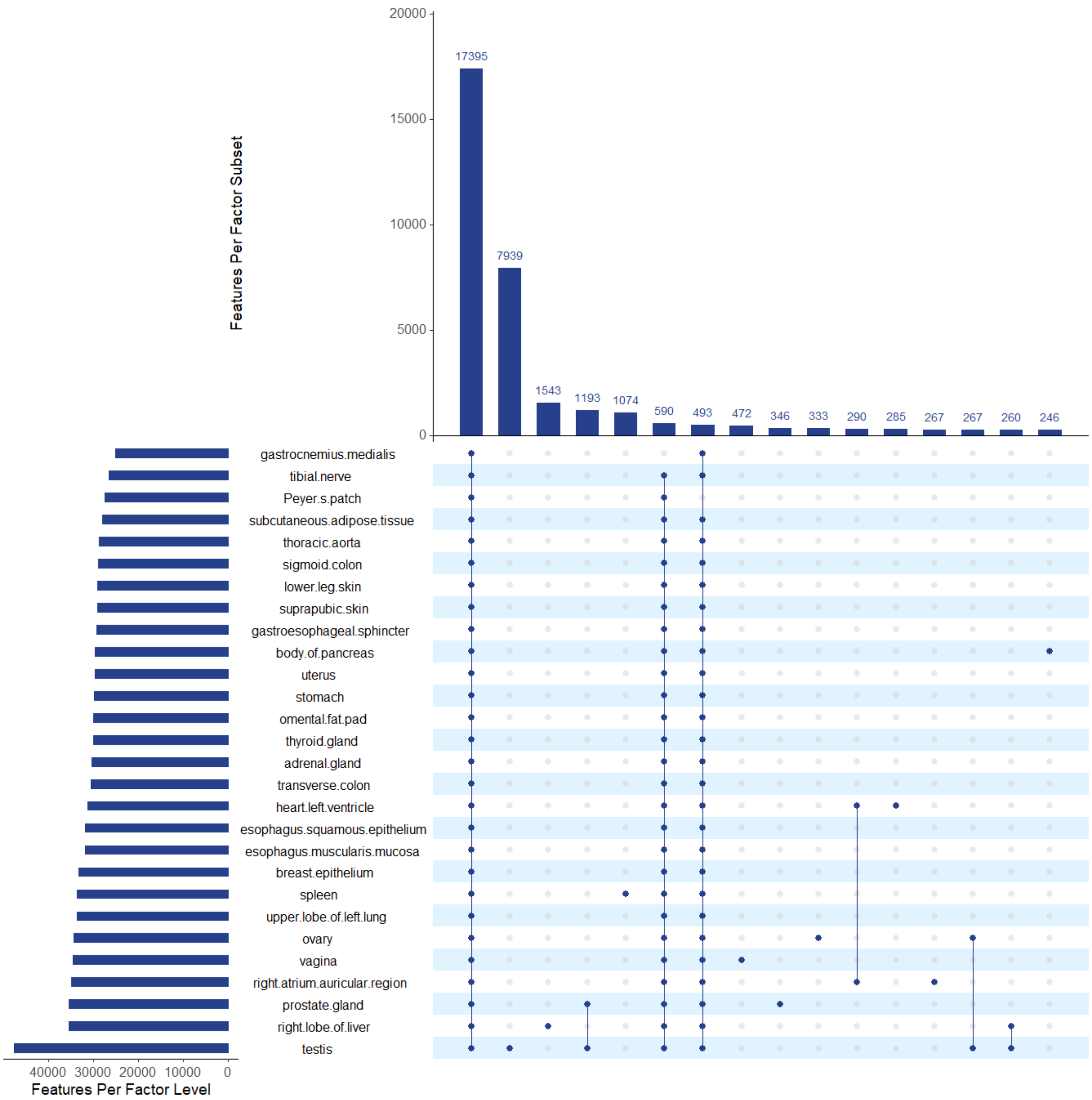

**Figure S7.2f. Tissue specificity of RAMPAGE data at TSSs of protein-coding genes**
This figure shows an UpSet plot of counts of GENCODE TSSs of genes (vertical bars), measured using RAMPAGE data in combinations of tissues (sets of dots), sorted by the number of TSSs. Bars on the left correspond to the number of TSSs in each tissue. Ubiquitously expressed TSSs using RAMPAGE are the most abundant.

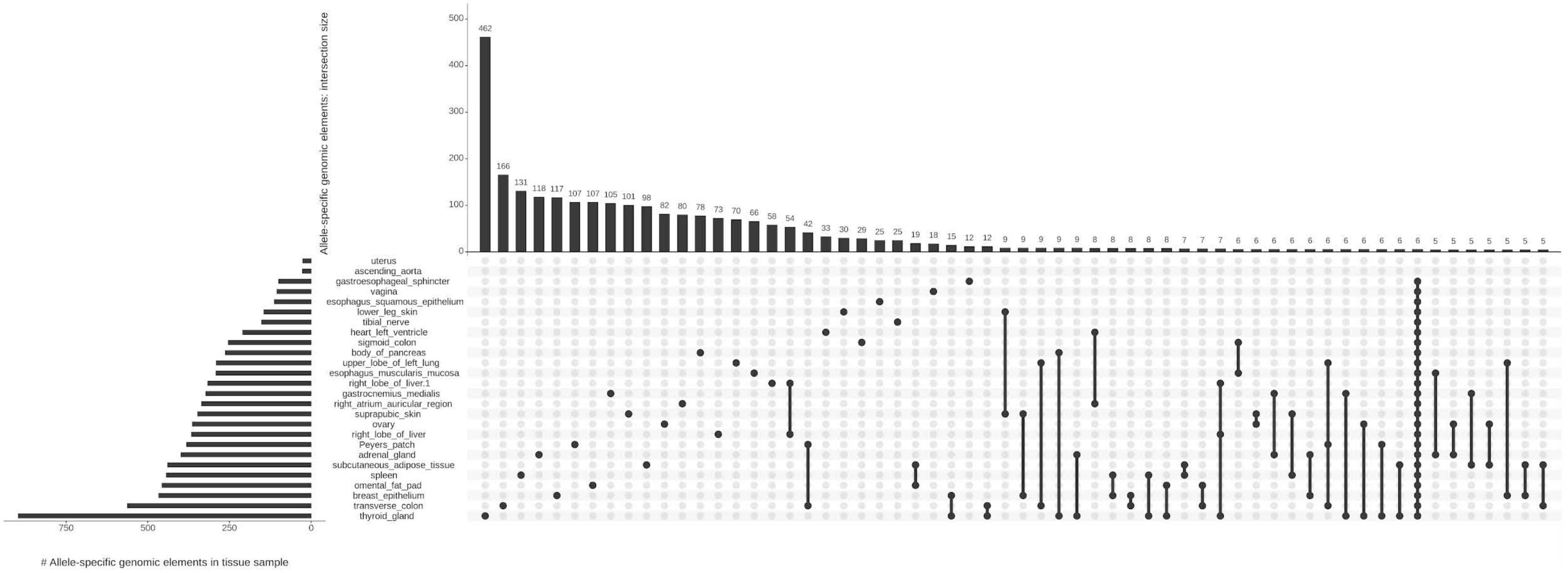

Allele-specific genomic elements: intersection size
### Allele-specific genomic elements in tissue sample
462
166
131
118
117
107
107
105
101
98
82
80
78
73
70
66
58
54
42
33
30
29
25
25
19
18
15
12
12
9
9
9
9
9
8
8
8
8
8
7
7
7
6
6
6
6
6
6
6
6
6
6
6
5
5
5
5
5
5
5
uterus
ascending_aorta
gastroesophageal_sphincter
vagina
esophagus_squamous_epithelium
lower_leg_skin
tibial_nerve
heart_left_ventricle
sigmoid_colon
body_of_pancreas
upper_lobe_of_left_lung
esophagus_muscularis_mucosa
right_lobe_of_liver.1
gastrocnemius_medialis
right_atrium_auricular_region
suprapubic_skin
ovary
right_lobe_of_liver
Peyers_patch
adrenal_gland
subcutaneous_adipose_tissue
spleen
omental_fat_pad
breast_epithelium
transverse_colon
thyroid_gland
750
500
250
0
500
400
300
200
100
0

**Figure S7.2g. ASE genes across different tissues of individual 3**

Counts of genes (bars) called ASE in the combinations of tissues (sets of dots) with the largest number of ASE genes. Bars on the left correspond to the number of ASE genes in each tissue.

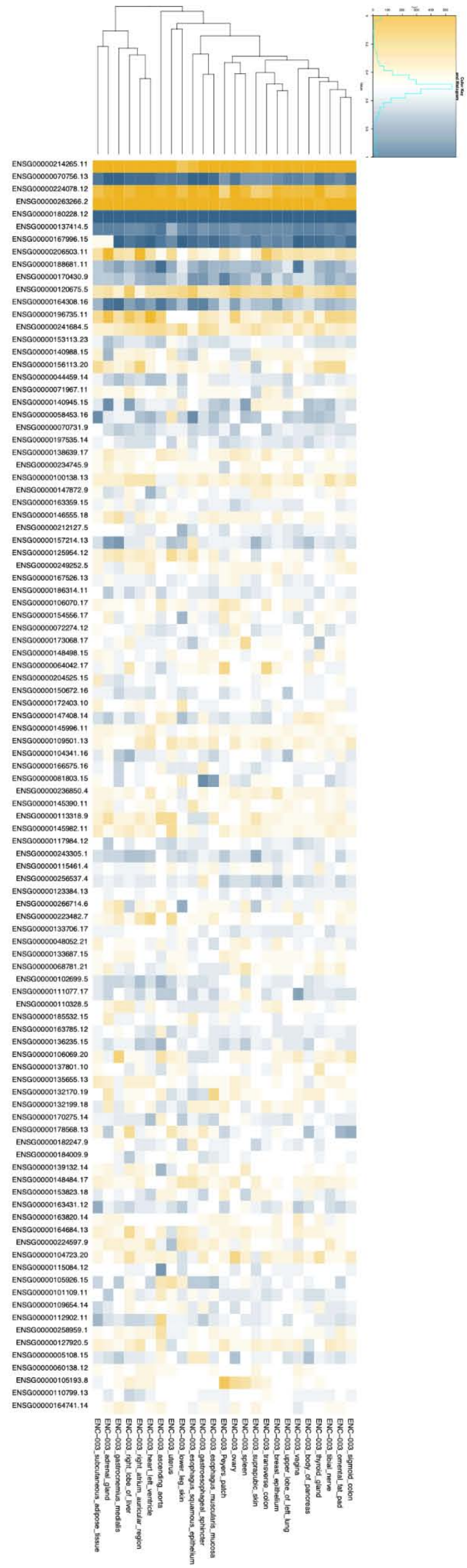

ENSG00000214265.11
ENSG00000070756.13
ENSG00000224078.12
ENSG00000263266.2
ENSG00000180228.12
ENSG00000137414.5
ENSG00000167996.15
ENSG00000206503.11
ENSG00000188681.11
ENSG00000170430.9
ENSG00000120675.5
ENSG00000164308.16
ENSG00000196735.11
ENSG00000241684.5
ENSG00000153113.23
ENSG00000140988.15
ENSG00000156113.20
ENSG00000044459.14
ENSG00000071967.11
ENSG00000140945.15
ENSG00000058453.16
ENSG00000070731.9
ENSG00000197535.14
ENSG00000138639.17
ENSG00000234745.9
ENSG00000100138.13
ENSG00000147872.9
ENSG00000163359.15
ENSG00000146555.18
ENSG00000212127.5
ENSG00000157214.13
ENSG00000125954.12
ENSG00000249252.5
ENSG00000167526.13
ENSG00000186314.11
ENSG00000106070.17
ENSG00000154556.17
ENSG00000072274.12
ENSG00000173068.17
ENSG00000148498.15
ENSG00000064042.17
ENSG00000204525.15
ENSG00000150672.16
ENSG00000172403.10
ENSG00000147408.14
ENSG00000145996.11
ENSG00000109501.13
ENSG00000104341.16
ENSG00000166575.16
ENSG00000081803.15
ENSG00000236850.4
ENSG00000145390.11
ENSG00000113318.9
ENSG00000145982.11
ENSG00000117984.12
ENSG00000243305.1
ENSG00000115461.4
ENSG00000256537.4
ENSG00000123384.13
ENSG00000266714.6
ENSG00000223482.7
ENSG00000133706.17
ENSG00000048052.21
ENSG00000133687.15
ENSG00000068781.21
ENSG00000102699.5
ENSG00000111077.17
ENSG00000110328.5
ENSG00000185532.15
ENSG00000163785.12
ENSG00000136235.15
ENSG00000106069.20
ENSG00000137801.10
ENSG00000135655.13
ENSG00000132170.19
ENSG00000132199.18
ENSG00000170275.14
ENSG00000178568.13
ENSG00000182247.9
ENSG00000184009.9
ENSG00000139132.14
ENSG00000148484.17
ENSG00000153823.18
ENSG00000163431.12
ENSG00000163820.14
ENSG00000164684.13
ENSG00000224597.9
ENSG00000104723.20
ENSG00000115084.12
ENSG00000105926.15
ENSG00000101109.11
ENSG00000109654.14
ENSG00000112902.11
ENSG00000258959.1
ENSG00000127920.5
ENSG00000005108.15
ENSG00000060138.12
ENSG00000105193.8
ENSG00000110799.13
ENSG00000164741.14
ENC−003_subcutaneous_adipose_tissue
ENC−003_adrenal_gland
ENC−003_gastrocnemius_medialis
ENC−003_right_lobe_of_liver
ENC−003_right_atrium_auricular_region
ENC−003_heart_left_ventricle
ENC−003_ascending_aorta
ENC−003_uterus
ENC−003_lower_leg_skin
ENC−003_esophagus_squamous_epithelium
ENC−003_gastroesophageal_sphincter
ENC−003_esophagus_muscularis_mucosa
ENC−003_Peyers_patch
ENC−003_ovary
ENC−003_spleen
ENC−003_suprapubic_skin
ENC−003_transverse_colon
ENC−003_breast_epithelium
ENC−003_upper_lobe_of_left_lung
ENC−003_vagina
ENC−003_body_of_pancreas
ENC−003_thyroid_gland
ENC−003_tibial_nerve
ENC−003_omental_fat_pad
ENC−003_sigmoid_colon

**Figure S7.2h. Haplotype 1 allele ratios (number of haplotype 1 reads over the total number of reads) for expression of genes that are accessible across all tissues and AS in at least one tissue of individual 3**

This figure parallels the allelic ratios for H3K27ac in Figure 9G and shows the same trend for expression as for histone modification.

| cCRE ID | cCRE Type | cCRE Coordinate | Regulatory Build | Associated Gene Name | Gene Type | Housekeeping Gene |
|---|---|---|---|---|---|---|
| EH38D2450505 | pELS_CTCF_bound | chr11_47642297_47642476 | Promoter | MTCH2 | Protein coding | Yes |
| EH38D2768300 | pELS_CTCF_bound | chr15_24956163_24956513 | Promoter | SNRPN | Protein coding | / |
| EH38D2900035 | PLS_CTCF_bound | chr17_1455938_1456100 | Promoter | CRK | Protein coding | Yes |
| EH38D2901207 | pELS_CTCF_bound | chr17_2401937_2402232 | Promoter | MNT | Protein coding | / |
| EH38D2916215 | dELS_CTCF_bound | chr17_19507807_19508157 | / | / | / | / |
| EH38D2933500 | pELS_CTCF_bound | chr17_43360380_43360725 | Promoter | LINC00910 | LncRNA | / |
| EH38D3043965 | pELS_CTCF_bound | chr19_14005711_14006060 | Promoter | RFX1 | Protein coding | / |
| EH38D3061913 | pELS_CTCF_bound | chr19_40425366_40425529 | Promoter | SERTAD1 | Protein coding | / |
| EH38D3112234 | pELS_CTCF_bound | chr2_39436701_39437019 | Promoter | MAP4K3 | Protein coding | / |
| EH38D3214874 | PLS_CTCF_bound | chr2_178451090_178451434 | Promoter | PRKRA | Protein coding | Yes |
| EH38D3218481 | PLS_CTCF_bound | chr2_183038344_183038694 | Promoter | NCKAP1 | Protein coding | / |
| EH38D3320686 | pELS_CTCF_bound | chr20_62652500_62652832 | Promoter | SLCO4A1 | Protein coding | / |
| EH38D3374502 | dELS_CTCF_bound | chr22_41414017_41414333 | / | / | / | / |
| EH38D3375755 | pELS_CTCF_bound | chr22_42614566_42614721 | Promoter | POLDIP3 | Protein coding | / |
| EH38D3448294 | PLS_CTCF_bound | chr3_75785373_75785718 | Promoter | ZNF717 | Protein coding | / |
| EH38D3802403 | pELS_CTCF_bound | chr6_291711_292043 | Promoter | DUSP22 | Protein coding | Yes |
| EH38D3802406 | pELS_CTCF_bound | chr6_292649_292999 | Promoter | DUSP22 | Protein coding | Yes |
| EH38D3819578 | pELS_CTCF_bound | chr6_17600685_17600980 | Promoter | FAM8A1 | Protein coding | Yes |
| EH38D3829720 | PLS_CTCF_bound | chr6_29888019_29888233 | Promoter | HLA-H | Pseudo | / |
| EH38D3829827 | pELS_CTCF_bound | chr6_29976507_29976854 | Promoter | HCG9 | LncRNA | / |
| EH38D3829829 | dELS_CTCF_bound | chr6_29977252_29977415 | Promoter | HCG9 | LncRNA | / |
| EH38D4038383 | pELS_CTCF_bound | chr7_139341640_139341807 | Promoter | FMC1-LUC7L2 | Protein coding | / |
| EH38D4168415 | pELS_CTCF_bound | chr9_6007913_6008223 | Promoter | KIAA2026 | Protein coding | / |

**Figure S7.2i. Annotation of pan-tissue H3K27ac AS+ cCREs of individual 3**

Among the 23 H3K27ac AS cCREs that were detected across all available tissues of individual 3, 21 cCREs are within promoter regions of known genes, including six promoters of housekeeping genes. Promoters and associated genes are based on Ensembl, and housekeeping genes are based on the HRT Atlas (Hounkpe *et al.*, 2021).

| GENCODE ID | Gene Name | Gene Type | Housekeeping Gene |
|---|---|---|---|
| ENSG00000070756.13 | PABPC1 | Protein coding | Yes |
| ENSG00000084623.11 | EIF3I | Protein coding | Yes |
| ENSG00000090372.14 | STRN4 | Protein coding | Yes |
| ENSG00000109919.9 | MTCH2 | Protein coding | Yes |
| ENSG00000119669.4 | IRF2BPL | Protein coding | Yes |
| ENSG00000122026.10 | RPL21 | Protein coding | Yes |
| ENSG00000122884.12 | P4HA1 | Protein coding | / |
| ENSG00000130844.16 | ZNF331 | Protein coding | / |
| ENSG00000137414.5 | FAM8A1 | Protein coding | Yes |
| ENSG00000151233.10 | GXYLT1 | Protein coding | / |
| ENSG00000167996.15 | FTH1 | Protein coding | / |
| ENSG00000180228.12 | PRKRA | Protein coding | Yes |
| ENSG00000187840.4 | EIF4EBP1 | Protein coding | / |
| ENSG00000204186.7 | ZDBF2 | Protein coding | / |
| ENSG00000214265.11 | RP11-701H24.9 | Protein coding | / |
| ENSG00000224078.12 | SNHG14 | ncRNA | / |
| ENSG00000227124.8 | ZNF717 | Protein coding | / |
| ENSG00000232653.8 | GOLGA8N | Protein coding | / |
| ENSG00000258186.2 | SLC7A5P2 | Pseudo | / |
| ENSG00000263266.2 | RPS7P1 | Pseudo | / |

**Figure S7.2j. Annotation of pan-tissue ASE genes of individual 3**

Among the 20 ASE genes that were detected across at least 90% of available tissues of individual 3, eight genes are annotated as housekeeping genes in the HRT Atlas (Hounkpe *et al.*, 2021).

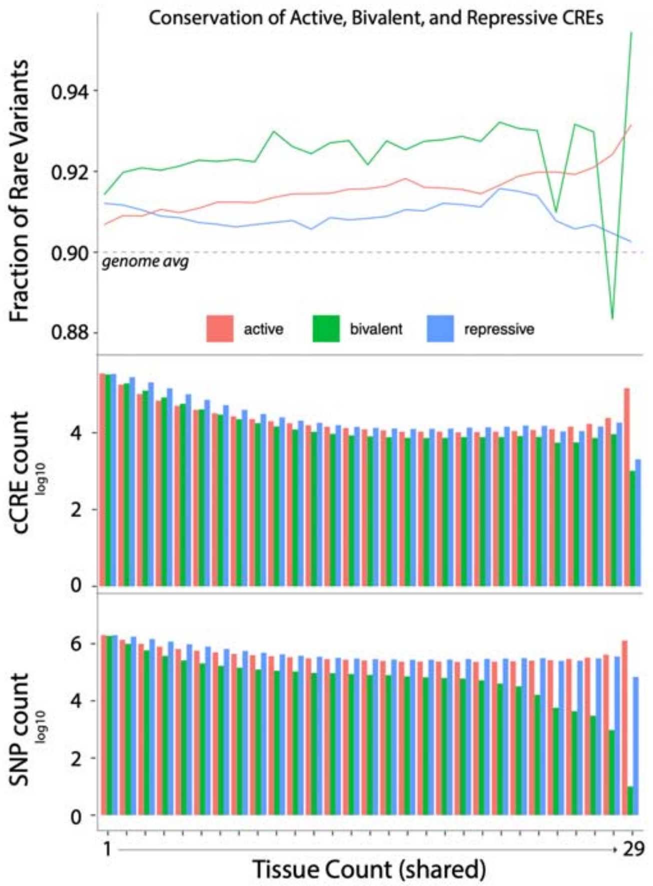

**Figure S7.2k. Rare DAF for active, bivalent, and repressive cCREs in increasing tissue count**

Fraction of rare variants was calculated as # rare variants / (# rare variants + # common variants). Total cCRE and SNP (taking into account all SNPS, common and rare) counts are shown for tissue count as well.

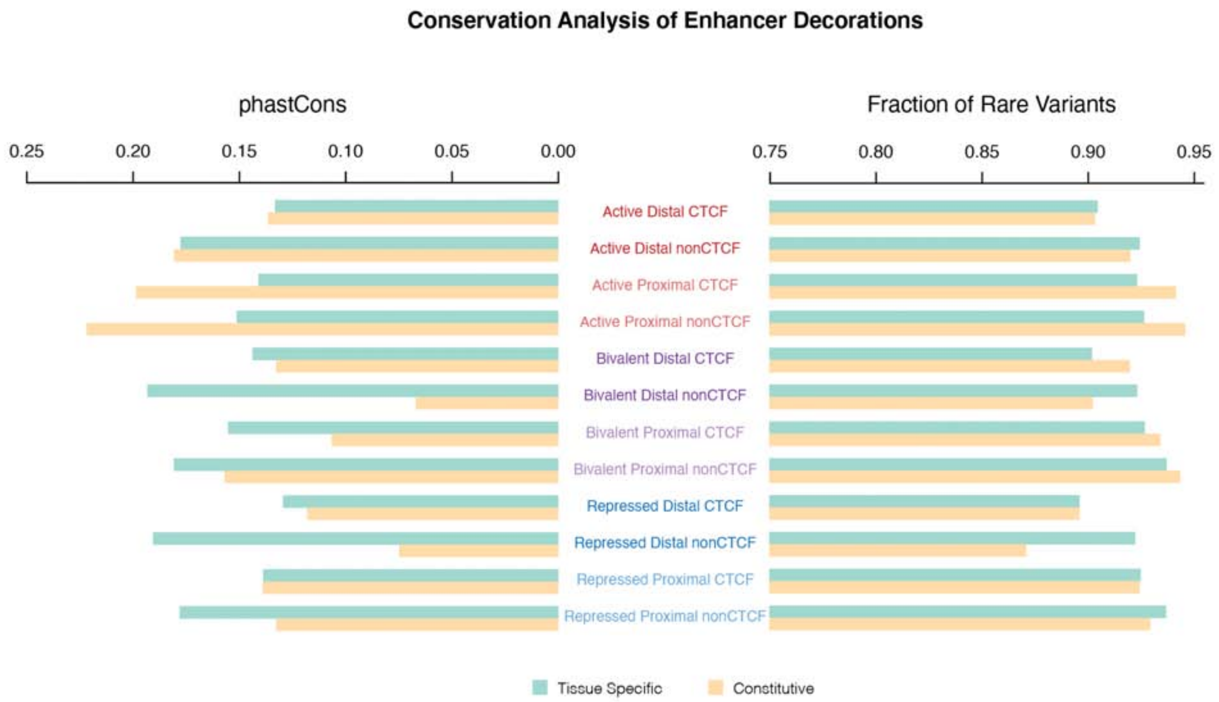

**Figure S7.2l. Conservation of enhancer decorations**

The conservation was calculated in terms of the phastCons score and fraction of rare variants, based on the DAF in the gnomAD database. The annotations are from Figure S7.1c.

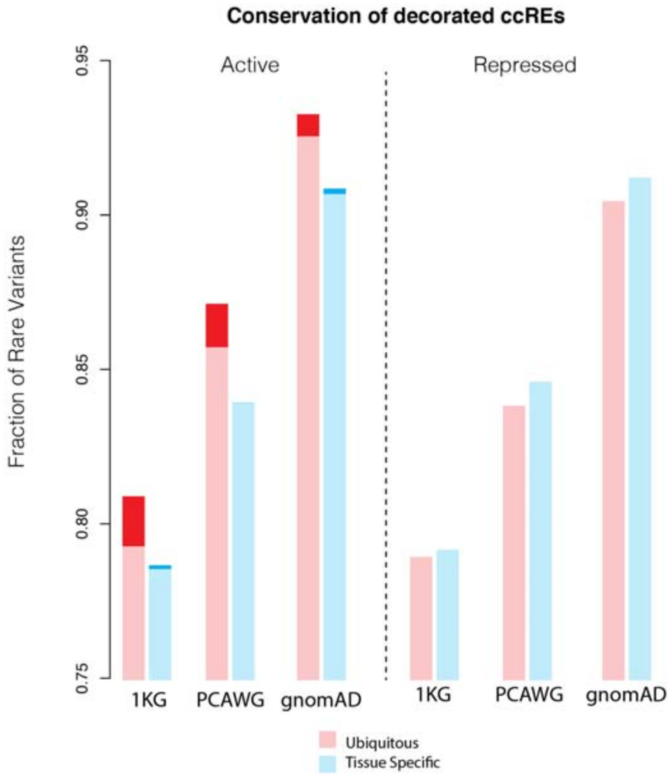

**Figure S7.2m. Conservation of active and repressed cCREs for tissue-specific and ubiquitous categories**

Dark red shows an increase in conservation for more stringently defined cCREs (selected via the top 1% of Matched Filter signals; See S7.2). The databases for this calculation include 1KG (1,000 Genomes), PCAWG, and gnomAD.

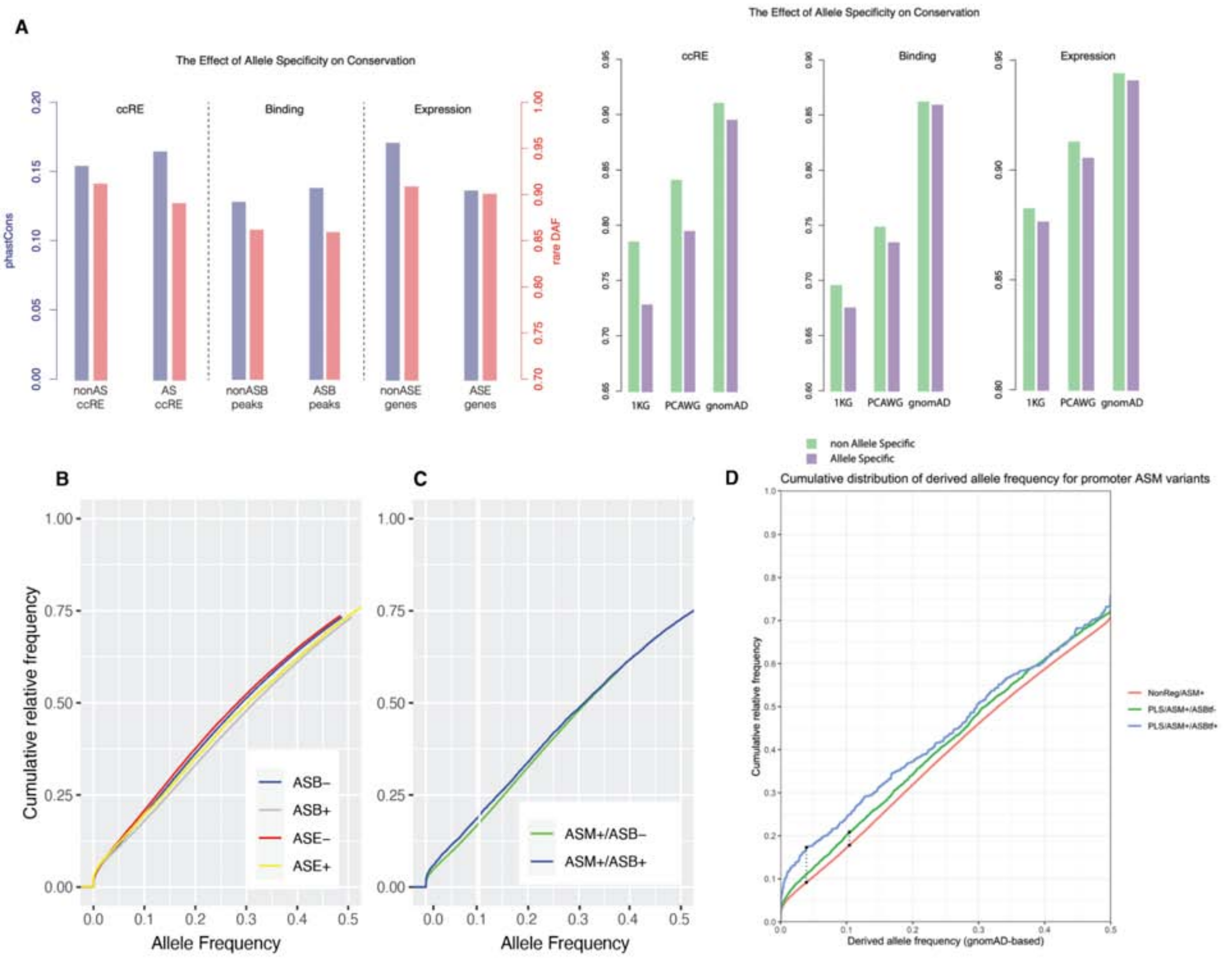

**Figure S7.2n. Conservation of regions exhibiting AS activity**

**(A)** The conservation of various AS annotations was calculated using phastCons and fraction of rare variants based on different population variants. Specifically, we considered AS/non-AS cCREs, AS/non-AS binding peaks from H3K27ac, and AS/non-AS genes. An alternate way to observe the same phenomena is to determine the cumulative relative frequency of variants, shown in **(B)**. Here, we see that non-AS events demonstrate stronger purifying selection than AS events, shown by the higher cumulative frequency curve. However, ASM/ASB variants demonstrate more consistency and higher purifying selection as compared to ASM/non-ASB events, shown in **(C).** Finally in **(D)** we show that the effect in **(C)** is amplified when exclusively considering promoter regions.

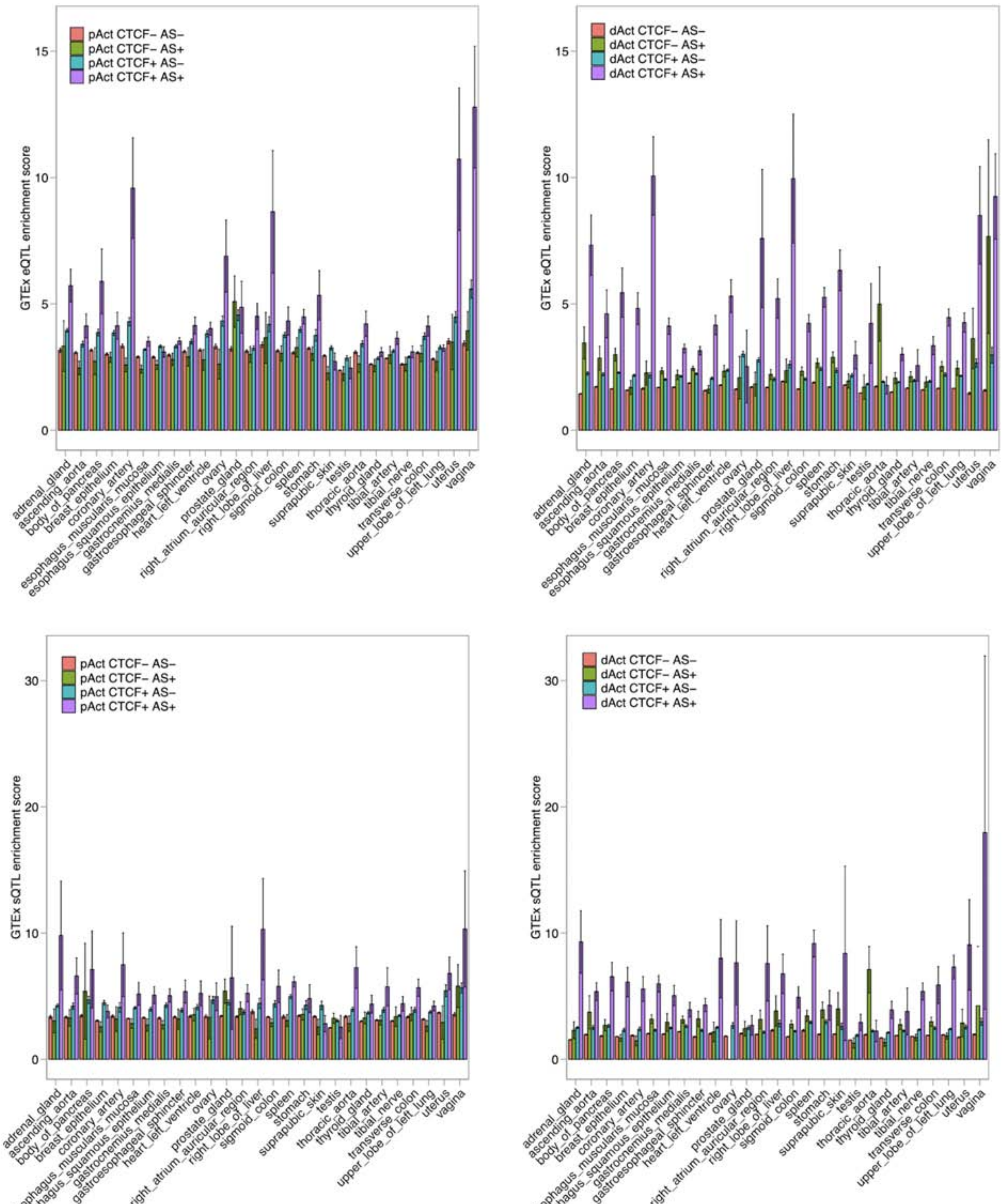

**Figure S7.3a. eQTL and sQTL enrichment in cCREs**

We computed odd ratios (ORs) to estimate the enrichment of the eQTL (upper panel) and sQTL (lower panel) SNPs identified from GTEx tissues in the cCREs from EN-TEx tissues. The ORs were calculated using the numbers of real QTL SNPs and the control SNPs located in the cCREs compared to those in the baseline regions. This procedure was repeated 30 times to calculate the standard deviation, and the values are indicated by the whiskers. In each panel, we show the QTL enrichment in the proximal active (left in each panel) and distal active (right in each panel) cCREs from each tissue type. In each figure, the cCREs are further separated into subgroups based on their CTCF binding and AS patterns. Note that the different decoration terms are defined in Figure S7.1c.

| Roadmap_ID | Roadmap_name | EN-Tex_name | GTEx_name |
|---|---|---|---|
| E065 | Aorta | ascending_aorta | Artery_Aorta |
| E098 | Pancreas | body_of_pancreas | Pancreas |
| E119 | HMEC Mammary Epithelial Primary C | breast_epithelium | Breast_Mammary_Tissue |
| E079 | Esophagus | esopahgus_muscularis_muco | Esophagus_Muscularis |
| E107 | Skeletal Muscle Male | gastrocnemius_medialis | Muscle_Skeletal |
| E095 | Left Ventricle | heart_left_ventricle | Heart_Left_Ventricle |
| E097 | Ovary | ovary | Ovary |
| E109 | Small Intestine | Peyers_patch | Small_Intestine_Terminal_Ileum |
| E104 | Right Atrium | right_atrium_auricular_region | Heart_Atrial_Appendage |
| E066 | Liver | right_lobe_of_liver | Liver |
| E106 | Sigmoid Colon | sigmoid_colon | Colon_Sigmoid |
| E113 | Spleen | spleen | Spleen |
| E094 | Gastric | stomach | Stomach |
| E096 | Lung | upper_lobe_of_lung | Lung |

**Figure S7.3b. Roadmap annotations**

We selected 14 tissue types that are matched across the EN-TEx, GTEx, and Roadmap projects to compare the QTL enrichment in the EN-TEx cCREs and Roadmap regulatory annotations. We used the 15-state Roadmap annotations in the analysis.

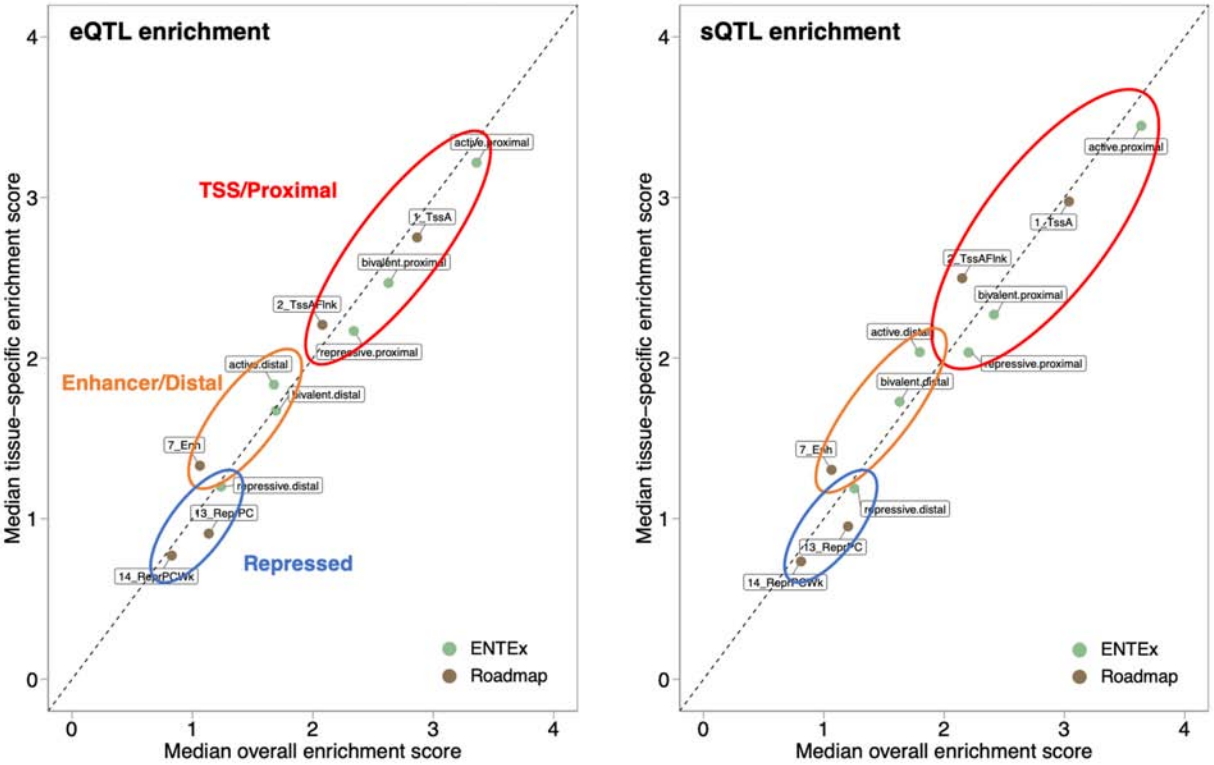

**Figure S7.3c. QTL enrichment in cCREs: EN-TEx vs. Roadmap**
We compared the enrichment of eQTL (left) and sQTL (right) SNPs in the TSS/proximal regions, enhancer/distal regions, and repressed regions. For this calculation, we matched the annotations between EN-TEx and Roadmap as shown in Figure S7.3b above.

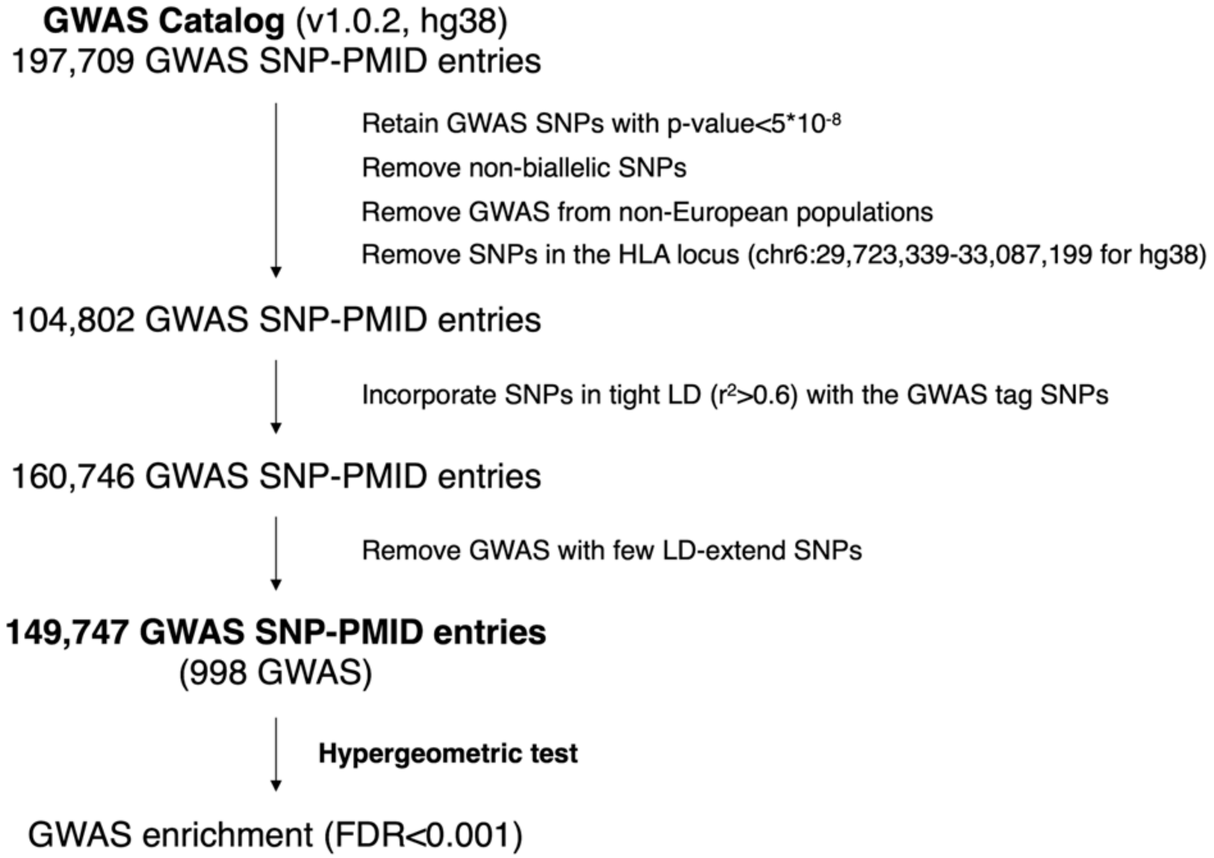

**Figure S7.3d. Framework of GWAS enrichment analysis**

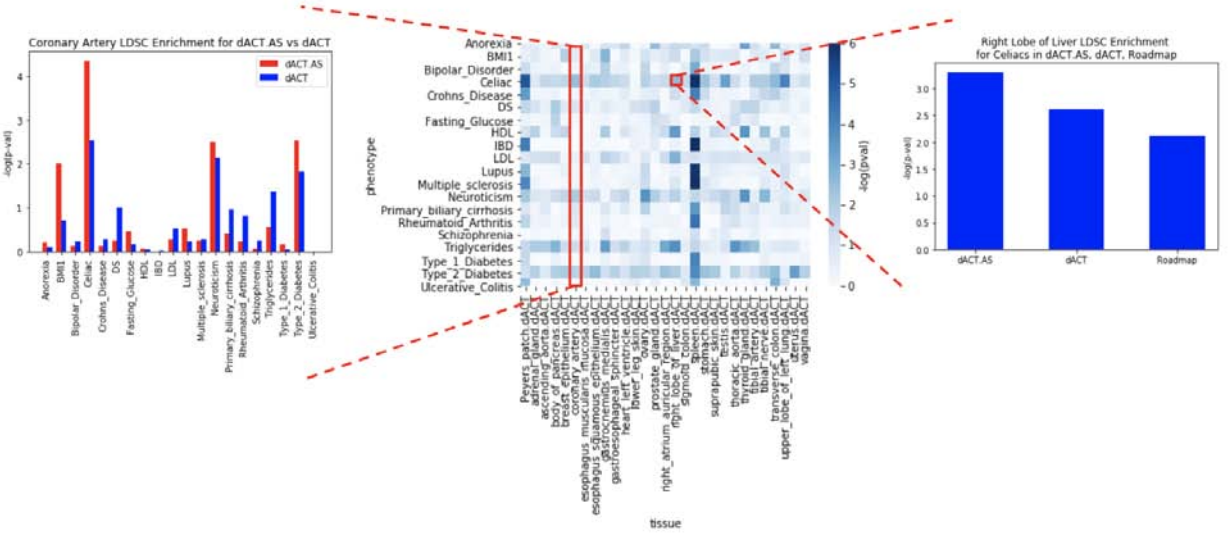

**Figure S7.3e. Stratified LDSC enrichment: Comparing EN-TEx AS, non-AS, and Roadmap annotations**

This is a shadow figure for Figure 10B in the main text. The central heatmap is the stratified LDSC enrichment of various GWAS traits over distal active elements of all EN-TEx tissues. In the left panel, we compare LDSC enrichment of distal active AS and non-AS over all traits for the coronary artery. In the right panel, we compare LDSC enrichment of distal active AS, non-AS, and Roadmap annotations in the right lobe of liver.

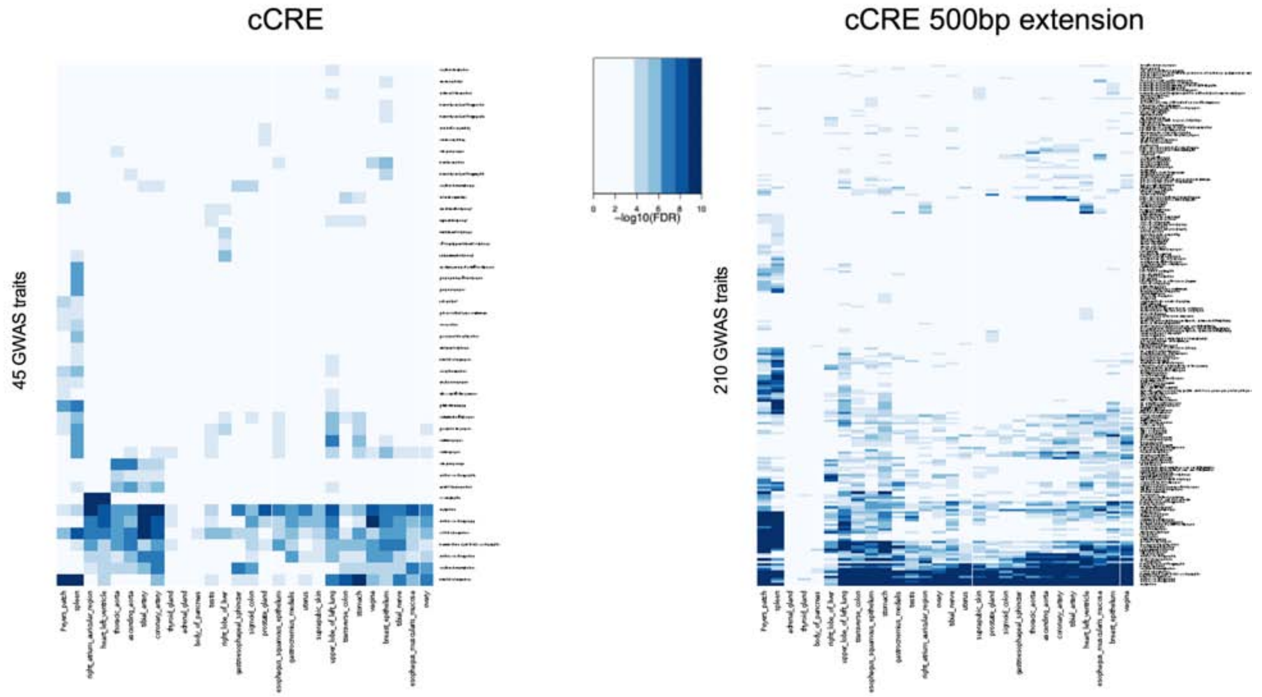

**Figure S7.3f. GWAS enrichment: cCREs vs. cCREs with 500 bp extensions**

We performed GWAS enrichment analysis on the original cCRE regions and the cCRE regions with a 500 bp extension on both sides. More significantly enriched GWAS traits can be identified on the cCRE regions with extensions, suggesting that it is necessary to include the flanking regions in the GWAS enrichment analysis.

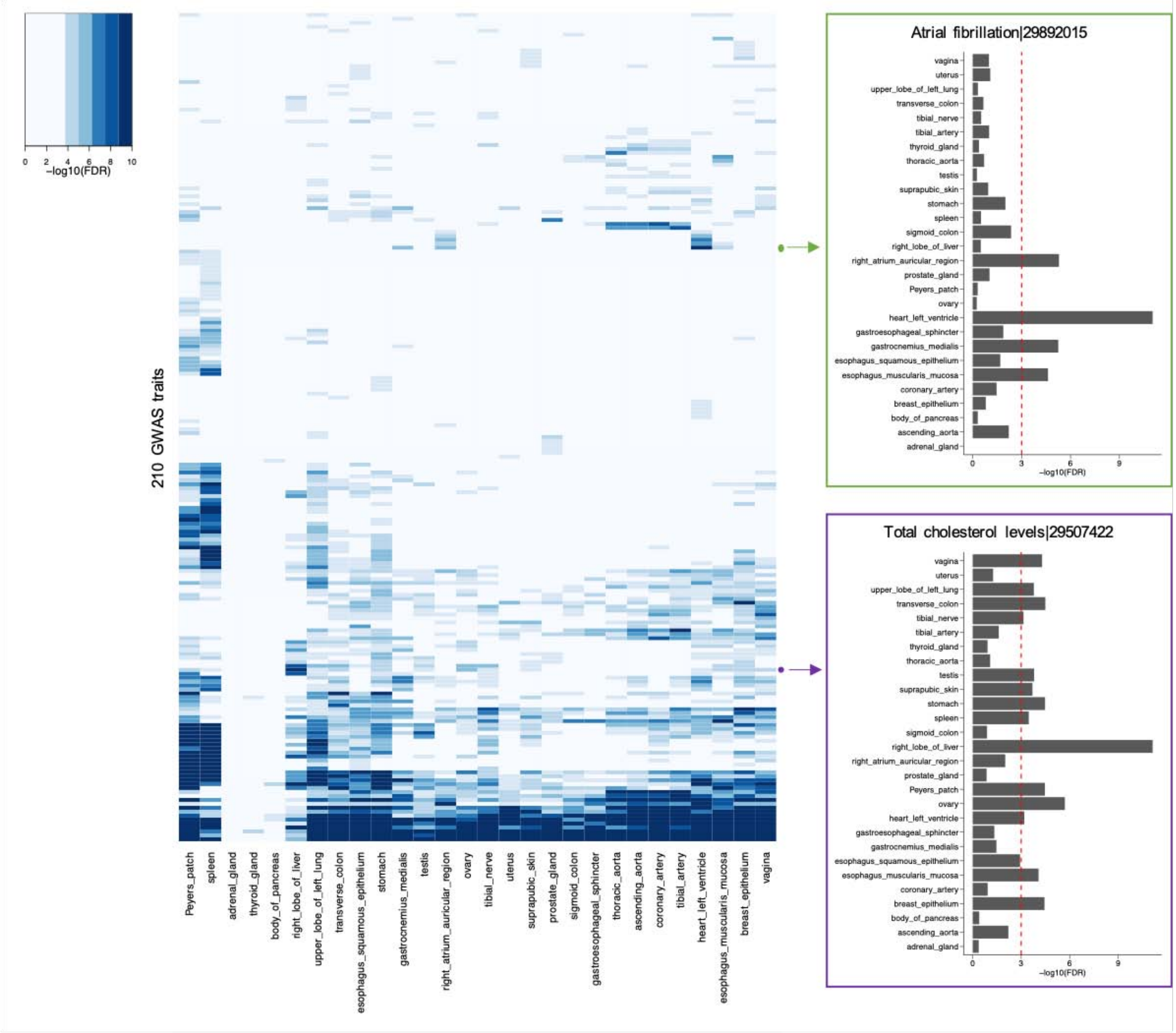

**Figure S7.3g. GWAS enrichment across tissues**

We selected two GWAS traits, atrial fibrillation and total cholesterol levels, to show their enrichment scores across all the tissue types.

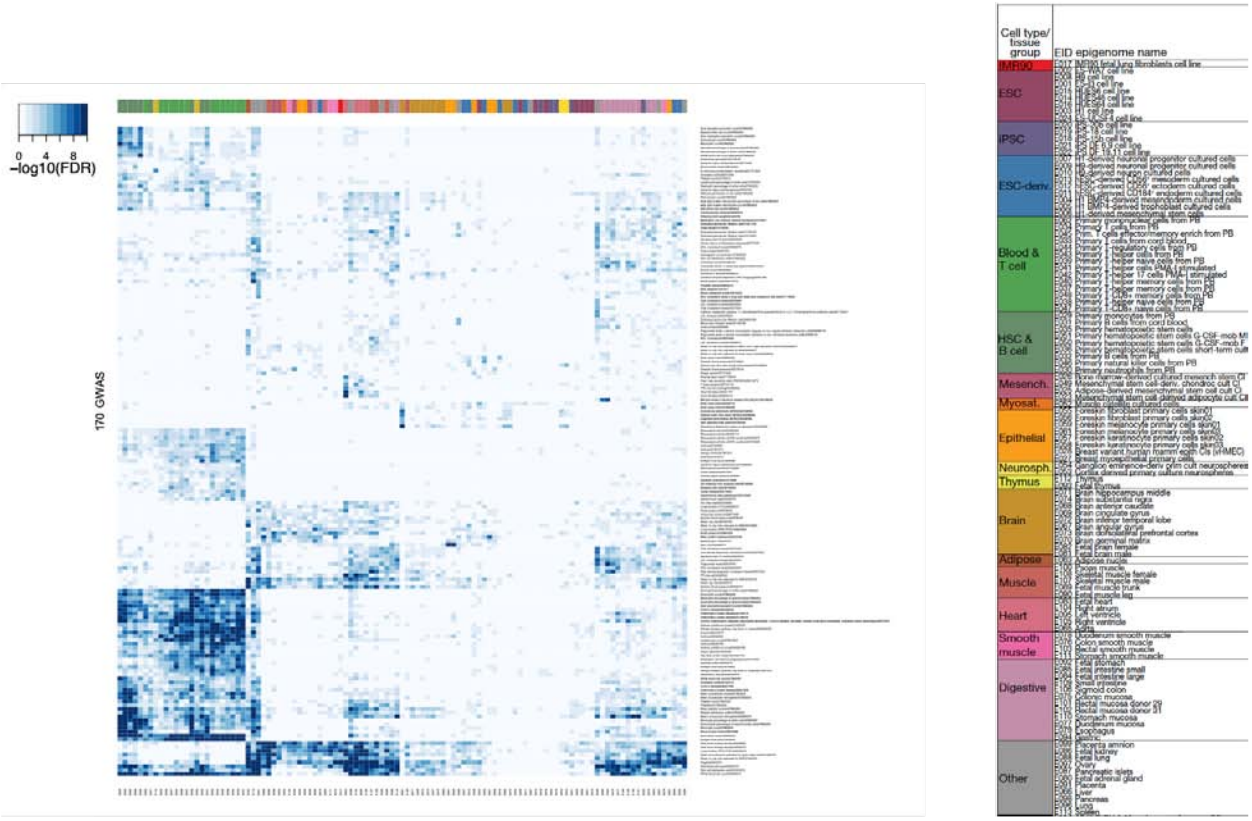

**Figure S7.3h. GWAS enrichment for Roadmap annotations**

We performed GWAS enrichment analysis on the enhancer annotations from the 127 cell and tissue types from the Roadmap Epigenomics Project. Tissue names are on the horizontal axis and traits are on the vertical axis. Simple clustering of this matrix reveals a blocky structure with sets of traits associated with groups of tissues.

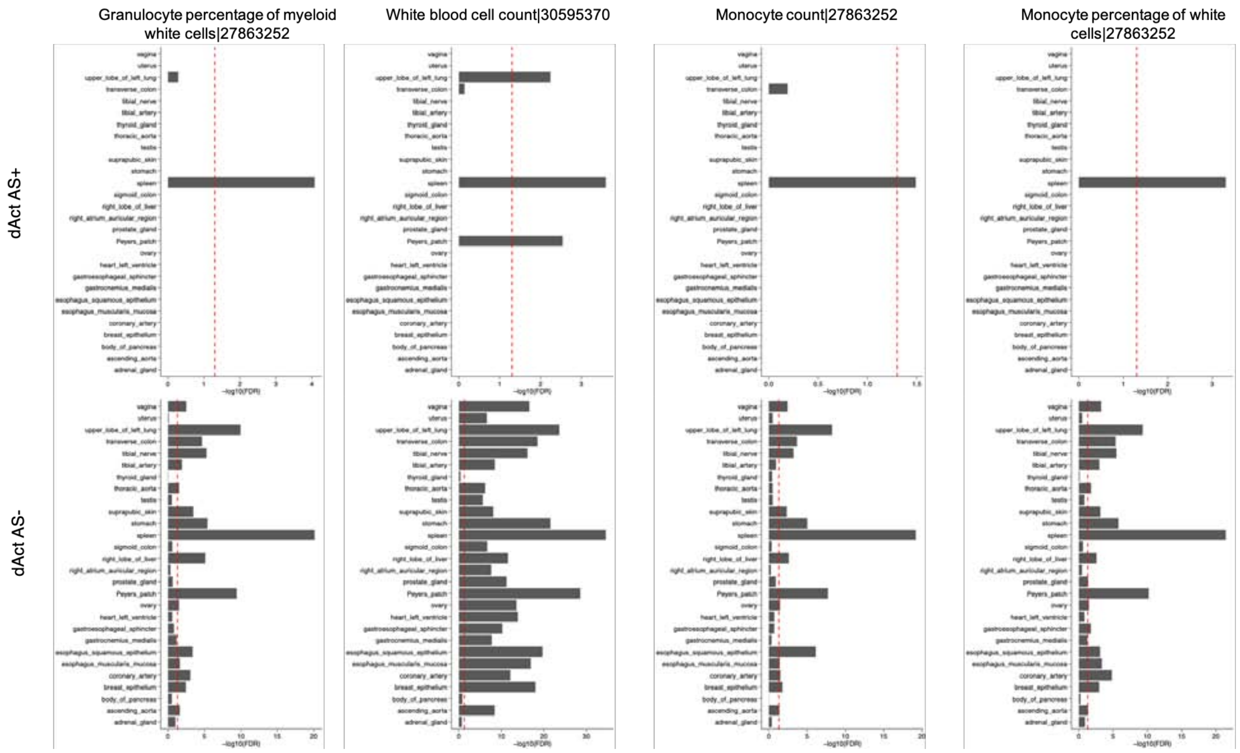

**Figure S7.3i. GWAS enrichment for AS+ vs. AS- cCREs**

We compared the GWAS enrichment scores on the distal active cCREs with (upper) and without (lower) AS signatures using the GWAS tag SNPs from blood-associated traits. Note that the different decoration terms are defined in Figure S7.1c.

**A** Motif+: the cCRE is overlapping any one of the **top 100** motifs

| | Motif + | Motif - |
|---|---|---|
| AS cCRE | 115934 | 149 |
| Non AS cCRE | 5203820 | 10432 |

Odd ratio=1.56
P<9.1e-9

**B** Motif+: the cCRE is overlapping any one of the **bottom 100** motifs

| | Motif + | Motif - |
|---|---|---|
| AS cCRE | 59252 | 56831 |
| Non AS cCRE | 3249317 | 1905683 |

Odd ratio=0.61
P≈0

**Figure S7.3j. Enrichment of AS sensitive or AS non-sensitive motifs in cCREs**
We intersect all cCREs with the **(A)** top 100 or **(B)** bottom 100 motifs from the motif ranking (see S4.1 for the ranking). AS cCREs (see S2.2) were significantly enriched with the top 100 motifs while non-AS cCREs contained more AS non-sensitive motifs (Fisher's exact test).

#### Figure S8. Supp. Figures for Main Text Section “Discussion”

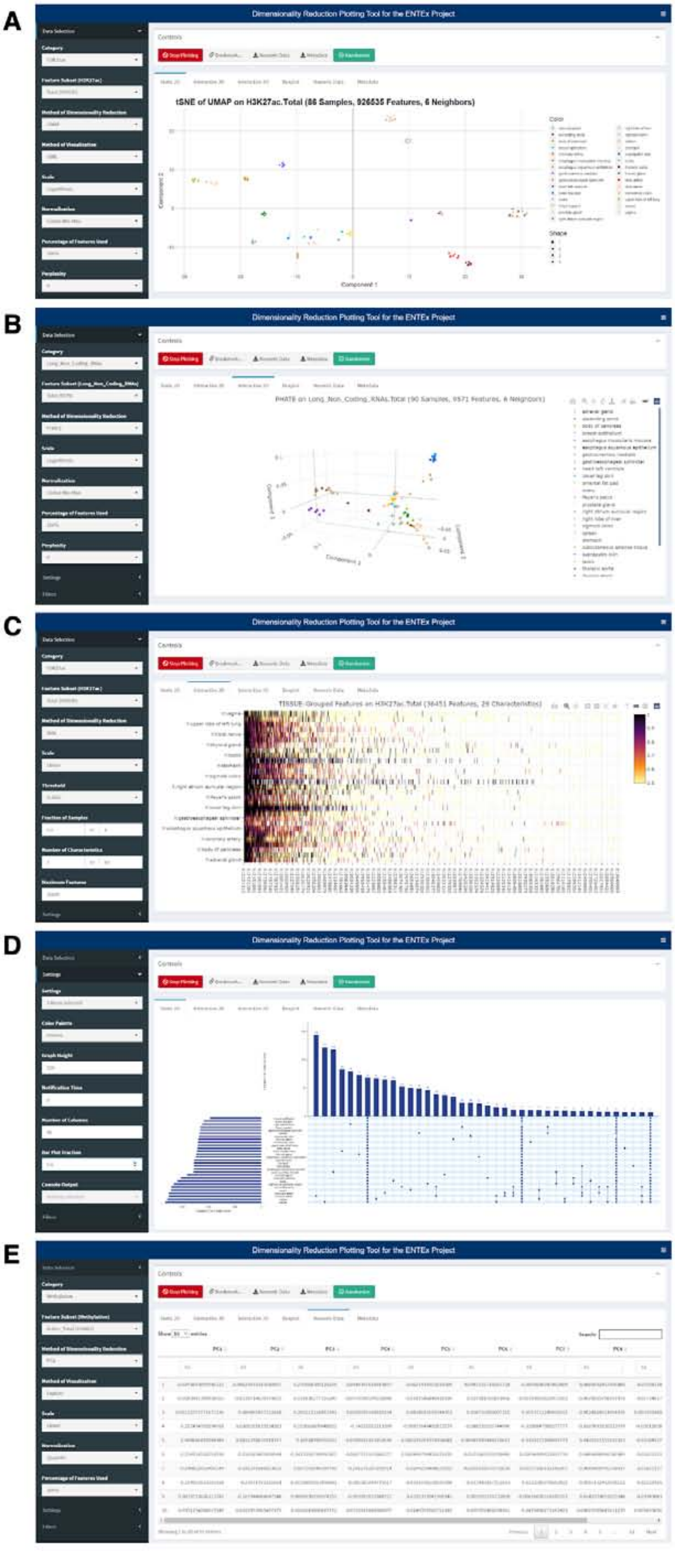
A
Dimensionality Reduction Plotting Tool for the ENTEx Project
tSNE of UMAP on H3K27ac.Total (86 Samples, 926535 Features, 6 Neighbors)
Component 1
B
Dimensionality Reduction Plotting Tool for the ENTEx Project
C
Dimensionality Reduction Plotting Tool for the ENTEx Project
D
Dimensionality Reduction Plotting Tool for the ENTEx Project
E
Dimensionality Reduction Plotting Tool for the ENTEx Project

**Figure S8.3. Explorer tool**

**(A)** Dimensionality reduction of the EN-TEx Explorer Tool allows for the generation of low-dimensional plots of several assays comprising cCREs, genomic expression, and proteomic expression. Data is primarily reduced to ten dimensions through PCA, VAE, UMAP, or PHATE. Components of the result can be plotted against each other (e.g., principal component 1 vs. principal component 2 on a scatter plot), summarized based on the reduction method, or reduced further with t-SNE. It is also possible to rapidly view different configurations of preprocessing parameters (scaling, normalization, feature variance) or hyperparameters through extensive precomputation. **(B)** Interactive reduction 2D and 3D visualizations are also included for intuitively exploring the data. **(C)** UpSetR plots visualize the intersection of genes in various tissues, replacing the traditional Venn diagram for larger sets. In the context of EN-TEx, these tools apply user-defined thresholds for each gene, consider the fraction of samples for which that gene is present in that tissue, and then calculate the UpSetR plot. **(D)** Heatmaps, which can also have dendrograms applied, visualize the data that are aggregated in the UpSetR plot. **(E)** The numeric data and metadata for all results can be bookmarked or downloaded for rapid sharing or analysis. Note, the input files for the explorer tool are available from the website. File: ENTEx.Explorer.cCRE.Combined.zip contains the cCREs, File: ENTEx.Explorer.Expression.Combined.zip contains the expressed genes, and File: ENTEx.Proteomics.cCRE.Combined.zip contains the mass spectrometry proteomic data.

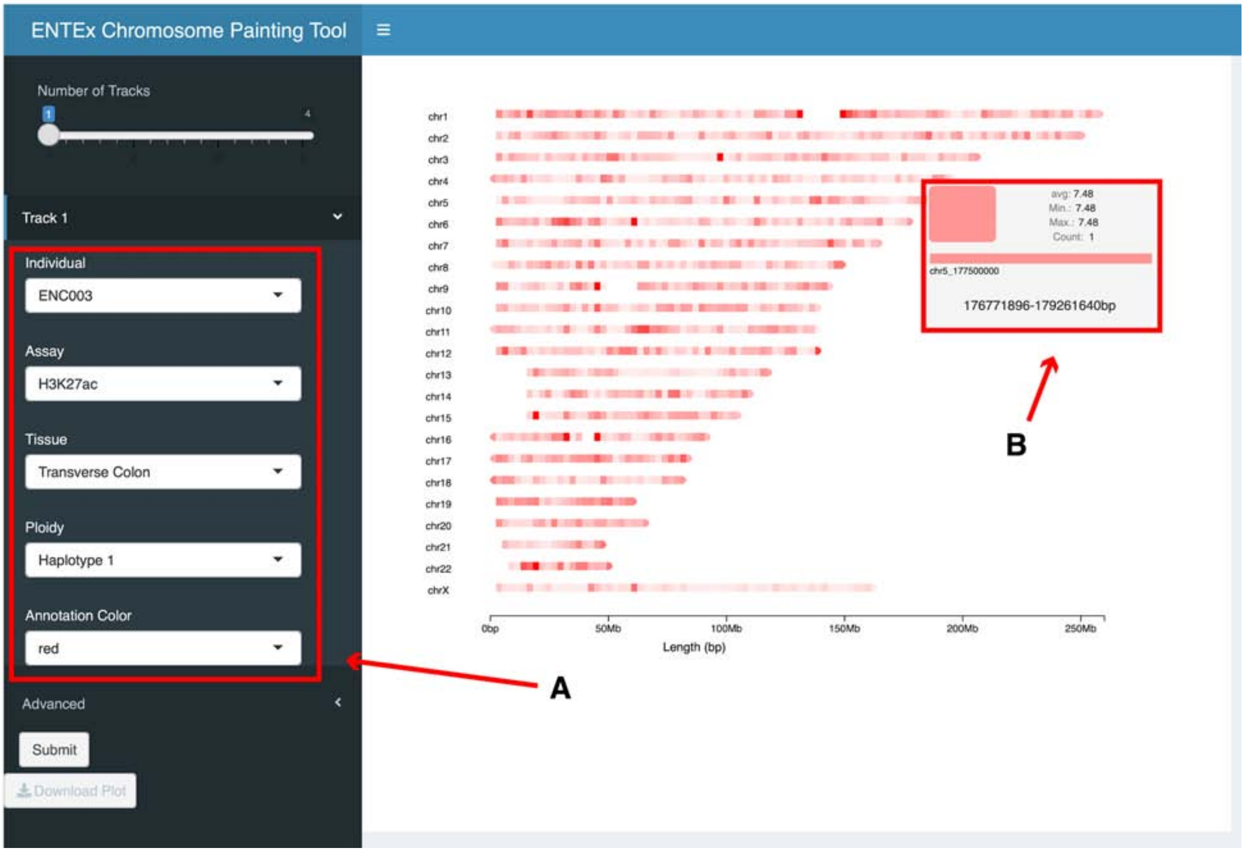

**Figure S8.4a. Screenshot of the EN-TEx chromosome painting tool**
(A) Parameters for data visualization of the EN-TEx data. (B) Plots generated by the chromosome painting tool are interactive.

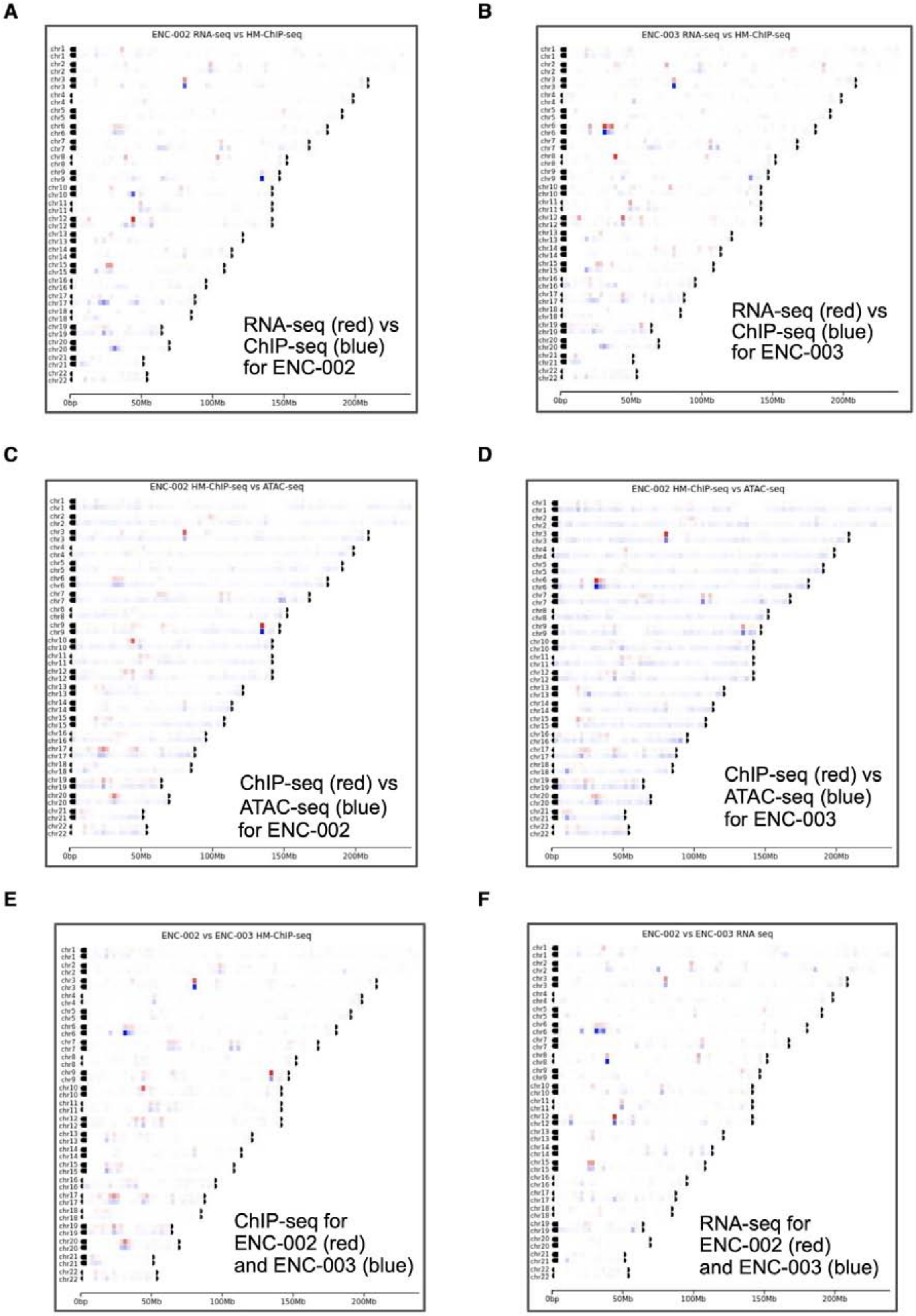
A
ENC-002 RNA-seq vs HM-ChIP-seq
RNA-seq (red) vs
ChIP-seq (blue)
for ENC-002
B
ENC-003 RNA-seq vs HM-ChIP-seq
RNA-seq (red) vs
ChIP-seq (blue)
for ENC-003
C
ENC-002 HM-ChIP-seq vs ATAC-seq
ChIP-seq (red) vs
ATAC-seq (blue)
for ENC-002
D
ENC-002 HM-ChIP-seq vs ATAC-seq
ChIP-seq (red) vs
ATAC-seq (blue)
for ENC-003
E
ENC-002 vs ENC-003 HM-ChIP-seq
ChIP-seq for
ENC-002 (red)
and ENC-003 (blue)
F
ENC-002 vs ENC-003 RNA seq
RNA-seq for
ENC-002 (red)
and ENC-003 (blue)
0bp
50Mb
100Mb
150Mb
200Mb

**Figure S8.4b. Examples of the chromosome painting tool**

**(A)**–**(B)** RNA-seq (red) vs. ChIP-seq (blue) for individuals 2 and 3. **(C)**–**(D)** RNA-seq and ChIP-seq for individual 2 (red) and individual 3 (blue). **(E)**–**(F)** ChIP-seq (red) vs. ATAC-seq (blue) for individuals 2 and 3.

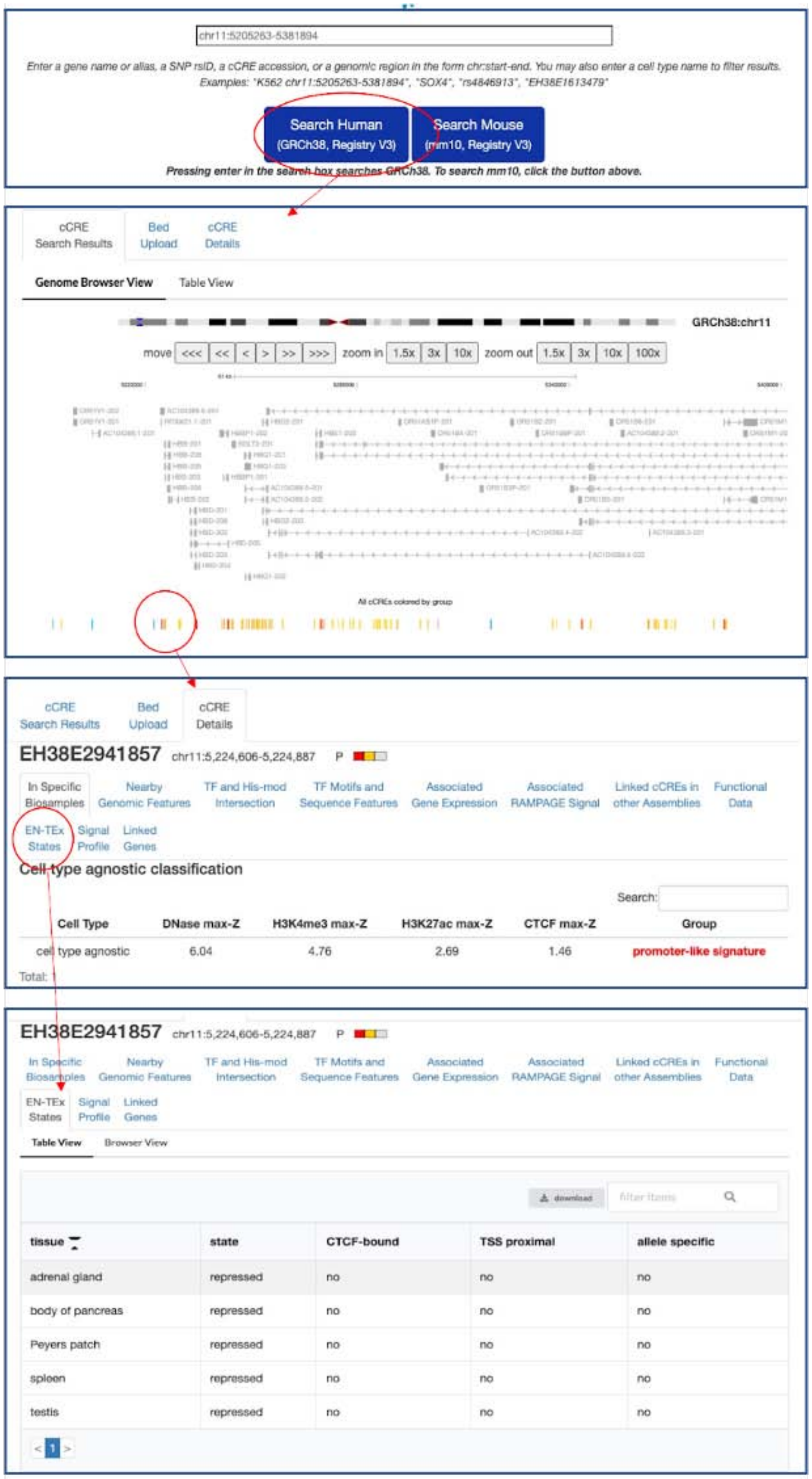

**Figure S8.5. Viewing EN-TEx decoration on the SCREEN website**
This shows views of the SCREEN website (screen.encodeproject.org) that indicate how to access the EN-TEx decoration of ENCODE Encyclopedia cCREs.

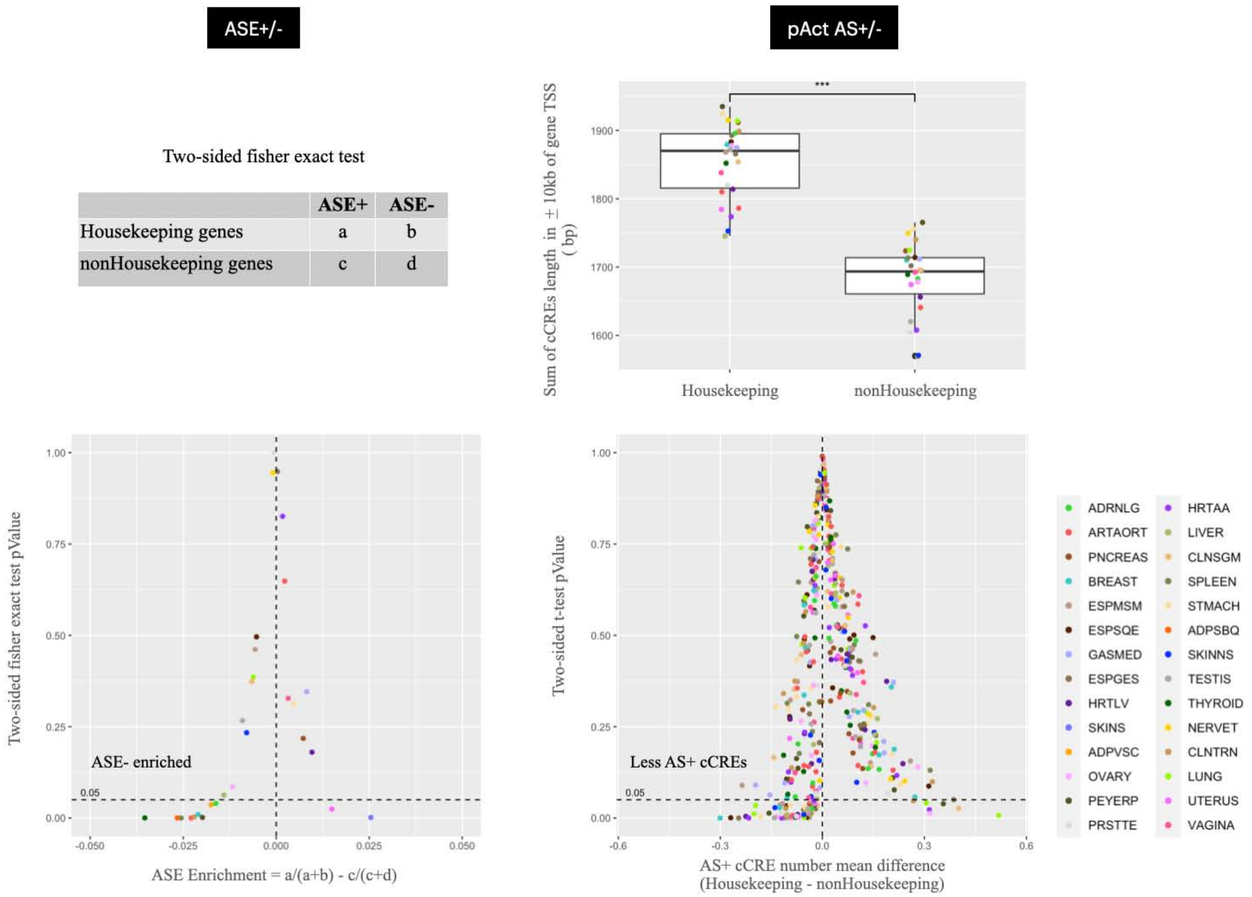

**Figure S8.6a. Allelic specificity of housekeeping genes**

Left: for each tissue, expressed protein-coding genes were split into housekeeping genes and non-housekeeping genes. Based on the two-sided Fisher's exact test, housekeeping genes are generally expressed in less of an AS fashion than non-housekeeping genes. Right: for each tissue, we examined the allele specificity of pAct cCREs flanking the TSS (defined by the gene starting site) of housekeeping genes. To eliminate the bias caused by significantly different cCRE lengths flanking the genes, we split genes into 20 bins based on the total length of the flanking cCREs. Within each bin, the number of pAct AS cCREs was compared between the housekeeping and non-housekeeping genes, and pAct cCREs flanking the housekeeping genes display relatively less allele specificity than the ones flanking non-housekeeping genes.

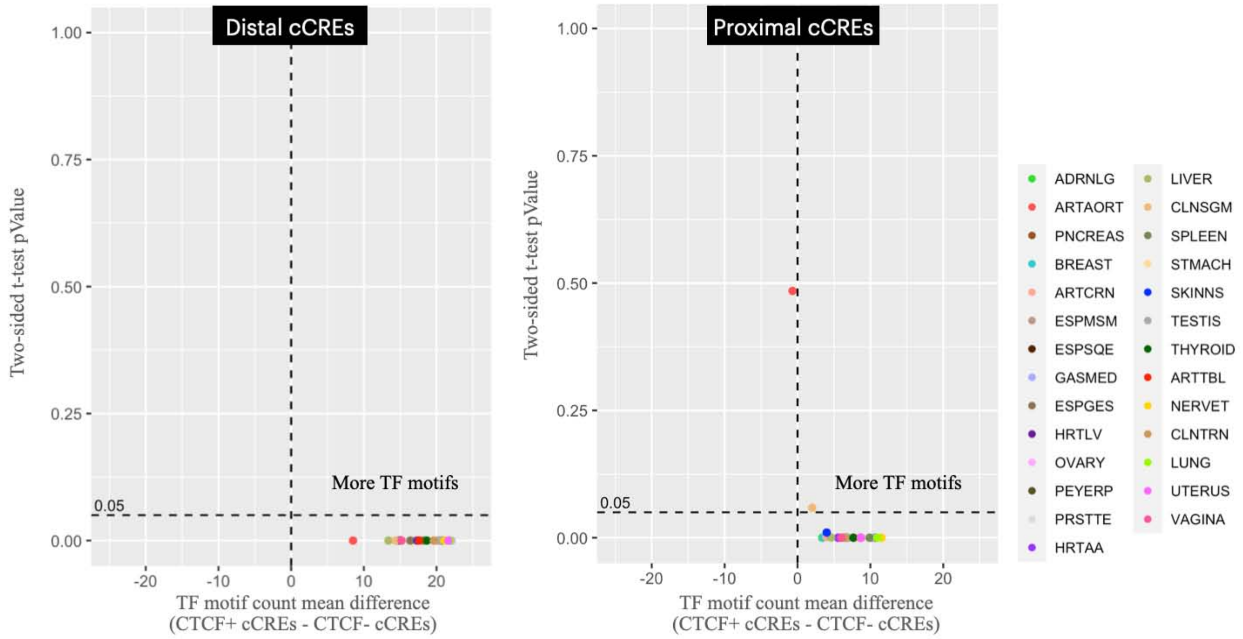

**Figure S8.6b. Enrichment of TF motifs in CTCF+ cCREs**

A list of 206 TF motifs (CTCF excluded) was used to count the total number of TF motifs that intersect with each CTCF+ and CTCF- cCRE in each tissue. For both distal and proximal cCREs, CTCF+ cCREs have significantly (paired-tissue two-sided t-test, p-value < 0.05) more TF motifs than CTCF- cCREs.

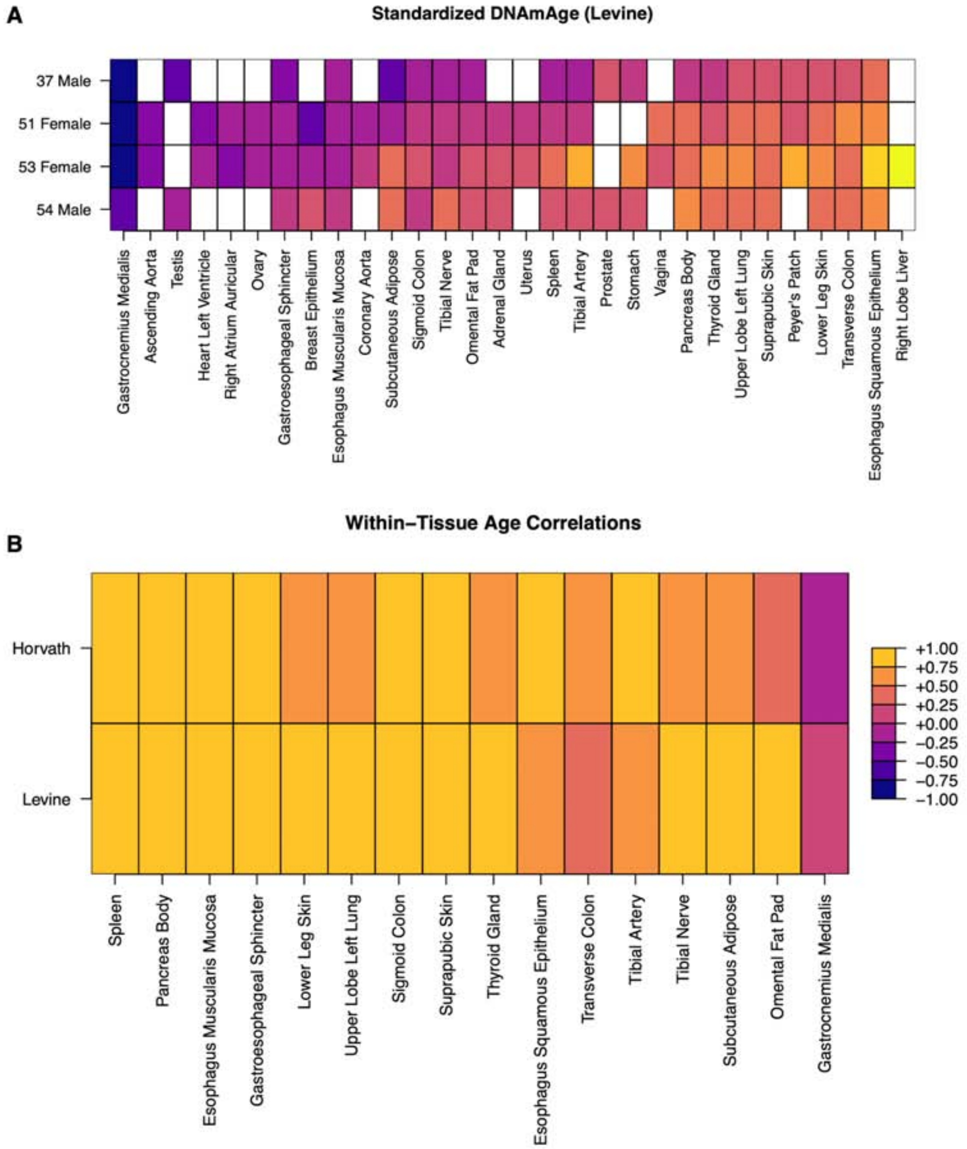

**Figure S8.7. Predicting the ages of tissues from their DNA methylation**
The statistical model developed by Levine et al. (Levine et al., 2018) was used to predict the ages of the different tissues from the four individuals. As a result, the different tissues of the same individuals have quite different predicted ages **(A)**. However, for each tissue type, the predicted ages and the actual ages of the four individuals tend to be highly correlated **(B),** suggesting that the model is accurate for capturing the changes in tissues with actual aging. The high correlation is also observed using other predictive models (Horvath, 2015). Taken together, these results suggest that the different tissues age at quite different speeds.

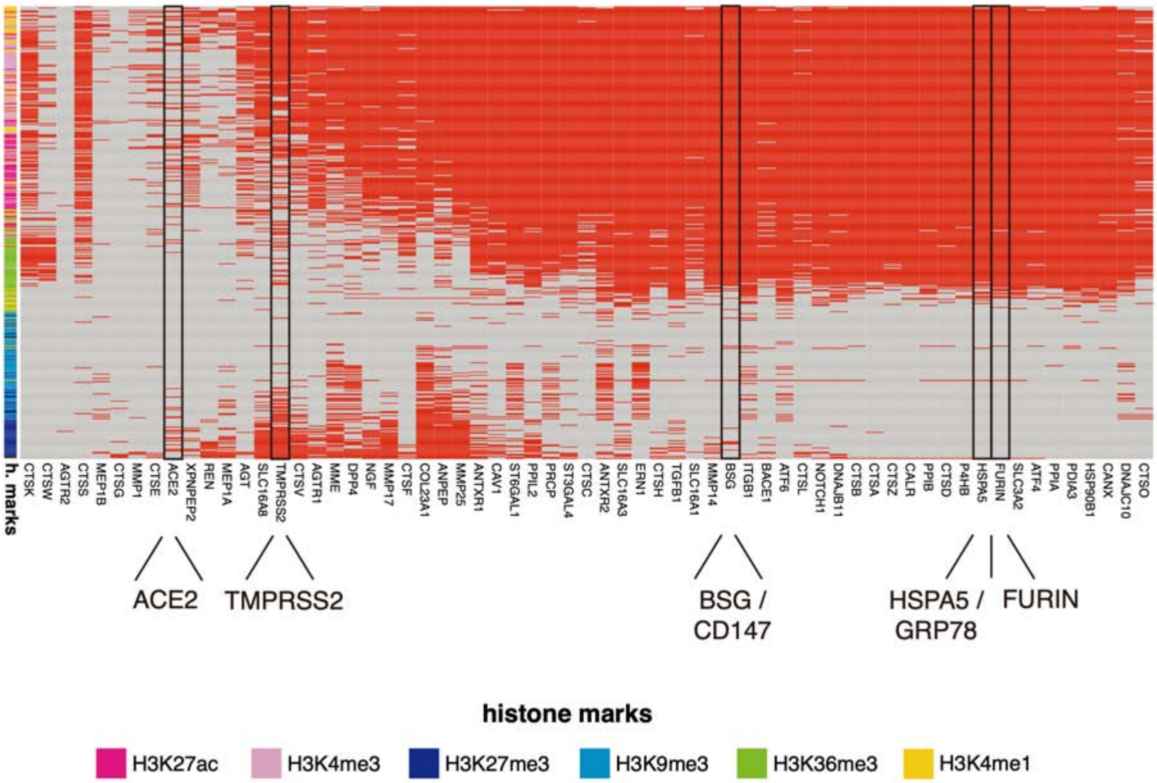

**Figure S8.8. Histone ChIP-seq data for COVID19-related genes**
Chromatin marking of COVID-19-related genes. The heatmap represents the presence/absence (red/gray) patterns of ChIP-seq peaks for the six histone marks assayed across the EN-TEx tissues. The list of 63 genes includes ACE2, CD147, FURIN, GRP78, and their protein interactors as retrieved from STRING (https://string-db.org/cgi/input?sessionId=bDjsdV72Wbsr&input_page_show_search=off)(Szklarczyk et al., 2019). Additional COVID-19/SARS-CoV-2 entry-associated genes proposed by the COVID19 Cell Atlas (https://www.covid19cellatlas.org/index.healthy.html), such as TMPRSS2, are also included (Sungnak et al., 2020).

**EN-TEx ENC002 hetSNVs VS GTEx ENC002**

- transverse_colon

| Match criteria | (chr, position, ref, alt, gt) | | | | (chr, position, ref, alt) |
|---|---|---|---|---|---|
| Type | Heterozygous | | Homozygous | | All |
| Source | EN-TEx | GTEx | GTEx | GTEx | GTEx |
| Genotype | 0\|1 or 1\|0 | 0\|1 | 0\|0 | 1\|1 | .\|., 0\|0, 0\|1, 1\|1 |
| ASE+ | 1,600 | 3,264,055 | 61,417,453 | 1,780,534 | 66,463,168 |
| | | 1,067 (66.69%) | 235 (14.69%) | 63 (3.94%) | 1365 (85.31%) |
| ASE- | 54,485 | 3,264,055 | 61,417,453 | 1,780,534 | 66,463,168 |
| | | 53,672 (98.51%) | 441 (0.81%) | 28 (0.05%) | 54,142 (99.37%) |
| ASB+ (HM+TF) | 4,498 | 3,264,055 | 61,417,453 | 1,780,534 | 66,463,168 |
| | | 2,491 (55.38%) | 756 (16.81%) | 310 (6.89%) | 3,557 (79.08%) |
| ASB- (HM+TF) | 259,038 | 3,264,055 | 61,417,453 | 1,780,534 | 66,463,168 |
| | | 246,713 (95.24%) | 6,756 (2.61%) | 477 (0.18%) | 253,946 (98.03%) |

- spleen

| Match criteria | (chr, position, ref, alt, gt) | | | | (chr, position, ref, alt) |
|---|---|---|---|---|---|
| Type | Heterozygous | | Homozygous | | All |
| Source | EN-TEx | GTEx | GTEx | GTEx | GTEx |
| Genotype | 0\|1 or 1\|0 | 0\|1 | 0\|0 | 1\|1 | .\|., 0\|0, 0\|1, 1\|1 |
| ASE+ | 315 | 3,264,055 | 61,417,453 | 1,780,534 | 66,463,168 |
| | | 175 (55.56%) | 53 (16.83%) | 12 (3.81%) | 240 (76.19%) |
| ASE- | 13,960 | 3,264,055 | 61,417,453 | 1,780,534 | 66,463,168 |
| | | 13,697 (98.12%) | 152 (1.09%) | 15 (0.11%) | 13,864 (99.31%) |
| ASB+ (HM+TF) | 14,645 | 3,264,055 | 61,417,453 | 1,780,534 | 66,463,168 |
| | | 8,477 (57.88%) | 2,423 (16.54%) | 986 (6.73%) | 11,886 (81.16%) |
| ASB- (HM+TF) | 838,756 | 3,264,055 | 61,417,453 | 1,780,534 | 66,463,168 |
| | | 815,653 (97.25%) | 12,574 (1.5%) | 852 (0.1%) | 829,080 (98.85%) |

**Figure S8.9. Coverage of EN-TEx AS hetSNVs in GTEx corresponding individual tissue**
The comparison was performed on two tissues from individual 2. For each of the four categories of hetSNVs called in EN-TEx, the number and percentage of EN-TEx hetSNVs detected in GTEx were calculated. For ASB hetSNVs, the call sets from all available histone marks (HM) and transcription factors (TFs) were integrated without duplications.
